# Salmonid gene expression biomarkers indicative of physiological responses to changes in salinity, temperature, but not dissolved oxygen

**DOI:** 10.1101/491001

**Authors:** Aimee Lee S. Houde, Arash Akbarzadeh, Oliver P. Günther, Shaorong Li, David A. Patterson, Anthony P. Farrell, Scott G. Hinch, Kristina M. Miller

## Abstract

An organism’s ability to respond effectively to environmental change is critical to their survival. Yet, life stage and overall condition can dictate tolerance thresholds to heightened environmental stressors, such that stress may not be equally felt across individuals within a species. Environmental changes can induce transcriptional responses in an organism, some of which reflect generalized responses, and others are highly specific to the type of change being experienced. Thus, if transcriptional biomarkers specific to a heightened environmental stressor, even under multi-stressor impacts, can be identified, the biomarkers could be then applied in natural environments to determine when and where individuals are experiencing such stressors. Here, we validate candidate gill gene expression biomarkers by experimentally challenging juvenile Chinook salmon *(Oncorhynchus tshawytscha*). A sophisticated experimental set-up (four trials) manipulated salinity (freshwater, brackish water, and seawater), temperature (10, 14, and 18°C), and dissolved oxygen (normoxia and hypoxia), in all 18 possible combinations, for up to six days during the pre-smolt, smolt, and de-smolt life stages. In addition, we also describe the changes in juvenile behaviour, plasma variables, gill Na^+^/K^+^- ATPase (NKA) activity, body size, body morphology, and skin pigmentation associated with salinity, temperature, dissolved oxygen, mortality, and smolt status. We statistically identified biomarkers specific to salinity and temperature treatments, as well as mortality across multiple stressors and life stages. Similar biomarkers for the dissolved oxygen treatment could not be identified in the data and we discuss our next steps using an RNA-seq study. This work demonstrates the unique power of gene expression biomarkers to identify a specific stressor even under multi-stressor conditions.

## 1. Introduction

Declining wild populations have been documented in association with environmental change in multitudes of species worldwide, suggesting that physiological limits of tolerance are being reached (Wikelski and Cooke, 2006). Notably, the early marine survival of Pacific salmon juveniles from many populations in southern British Columbia has diminished rapidly over the past twenty years, with hatchery smolt releases generally suffering the lowest survival (Beamish et al., 2009; Bradford and Irvine, 2000; Martins et al., 2012; Riddell et al., 2013). Factors responsible for the decline are beginning to emerge. For example, annual variability in the marine survival is associated, in part, with climate regime shifts, which can result in periods of heightened environmental stressors, including higher water temperatures and lower dissolved oxygen (Crozier, 2016). Although environmental monitoring provides information on the potential for heightened stressors, knowledge of the degree of animal exposure is typically lacking. Therefore, examining individual salmon is a more direct and integrative approach to assessing the magnitude of the effect different stressors are having on the fish (Wikelski and Cooke, 2006). For example, using panels of gene expression biomarkers associated with transcriptional responses to specific environmental stressors (Connon et al., 2018). A similar biomarker approach is being used for examining the relationship among Pacific salmon viral disease status and survival in the natural environment (Miller et al., 2017a). In the present study, our aim was to experimentally validate such biomarkers to a particularly critical development period of a salmon, smoltification, and overlay the interactive effects of temperature and hypoxia (low dissolved oxygen) challenges.

During smoltification, anadromous juvenile salmonids physiologically prepare themselves for the transition from freshwater to seawater (i.e. seawater tolerance). Inadequate smoltification or ill-timed seawater entry outside of their physiological window (as pre-smolts or de-smolts) can cause < 40% mortality or stunted growth for 1–2 months (Stien et al., 2013) beyond the well-documented physiological disturbances. Furthermore, the lower survival of hatchery-produced juveniles may also be associated, in part, with larger physiological stress during their freshwater to seawater transition relative to wild juveniles (e.g. Chittenden et al., 2008; Shrimpton et al., 1994). The smoltification window narrows at higher water temperatures (Bassett et al., 2018). Adult salmonids returning to rivers can similarly experience salinity stress if they are not adequately physiologically prepared (Cooke et al., 2006). The impact of hypoxia on smoltification is unknown. Yet, climate change is contributing to lower dissolved oxygen as well as higher temperature.

Critical benchmarks for hypoxic and thermal stress in salmonids for which most stocks cannot survive for prolonged periods of time are < 6 mg L^-1^ dissolved oxygen (DO) and 18°C (Lucas and Southgate, 2003). Thermal stress is associated with decreased survival and fitness-related traits (e.g. swim performance, body growth, and disease resistance) across all salmonid life stages (Quinn et al., 2011; Tomalty et al., 2015), although the effects on juveniles are much less studied. Furthermore, excessive nutrient loading, lower water discharge, and increased prevalence of algal blooms mean that billions of juvenile salmon annually experience hypoxia as they pass through estuaries on the North American Pacific coasts (Birtwell and Kruzynski, 1989). Eutrophic lakes can also become hypoxic (Conley et al., 2009), which may be a concern as juveniles begin the smoltification process, e.g. Sockeye salmon in Cultus Lake, BC (Putt, 2014). Hypoxic stress, due to repeated or prolonged exposure to low DO, is detrimental for fish activity, feeding, growth rates, and other normal biological functions (Closs et al., 2016).

As a non-lethal tool that uses gill tissue for measuring gene expression, we have already developed 93 candidate salinity (*n* = 37, Houde et al., 2018), temperature (*n* = 33, Akbarzadeh et al., 2018a), and dissolved oxygen (*n* = 23, Table S1) biomarkers for all Pacific salmon species. The candidate salinity biomarkers were originally developed and examined for broadly measuring the degree of smoltification (Houde et al., 2018). In the present study, we specifically examine if the regulation of some of these genes may be directly influenced by salinity. The candidate temperature biomarkers were previously examined as a single stressor (Akbarzadeh et al., 2018a). In the present study, we specifically examine these biomarkers under multi-stressor conditions. The candidate dissolved oxygen biomarkers were derived from a few studies on fish relative to the salinity and temperature biomarkers that were derived from a review of transcriptomic studies. Beyond these candidate biomarkers, there are currently validated biomarkers for viral disease development (Miller et al., 2017a), smoltification (Houde et al., 2018), and general stress and imminent mortality (Evans et al., 2011; Jeffries et al., 2014a; Jeffries et al., 2012; Jeffries et al., 2014b). We used the Fluidigm BioMark^TM^ platform, a high throughput microfluidics technology that can independently measure the gene expression of 96 assays by 96 samples at once, and has been used to measure pathogen loads (Miller et al., 2016). Consequently, this salmon ‘Fit-Chip’ has the potential to be the first tool of its kind to use rapid characterization of biomarkers to comprehensively assess physiological health and condition of cultured and wild fauna.

Our first objective was to identify the biomarkers specific to the salinity, temperature, and dissolved oxygen treatments across multi-stressor conditions and life stages. Therefore, we experimentally challenged ocean-type Chinook salmon *(Oncorhynchus tshawytscha*) to three salinities, three temperatures, and two dissolved oxygen values in all 18 possible combinations for six days using four trials that spanned the smoltification period (i.e. pre-smolt, smolt, and de-smolt, between March and August). For the pre-smolt and de-smolt trials, we expected some juvenile mortality (dead or moribund) in seawater. Thus, our second objective was to identify biomarkers associated with mortality and physiological imbalance in seawater. Few studies have specifically documented the association between gene expression and other physiological measures or fitness-related traits important to the conservation of fishes (e.g. Connon et al., 2018; Oomen and Hutchings, 2017). Hence, beyond measurements of mortality, we also collected measures of salmon fitness-related traits such as behaviour, skin pigmentation, body morphology, as well as physiological biomarkers of stress and seawater tolerance such as plasma lactate, glucose, chloride concentrations (Barton, 2002) and gill Na^+^/K^+^ ATPase activity (McCormick, 1993). These variables were then associated to gene expression patterns and differences among groups within treatments.

## 2. Materials and Methods

### 2.1. Study species

The experiment was approved by the Fisheries and Oceans (DFO) Pacific Region Animal Care Committee (2017–002 and 2017–020) abiding to the Canadian Council of Animal Care Standards. Sub-yearling ocean-type Chinook salmon *(Oncorhynchus tshawytscha*) juveniles were transported from Big Qualicum River Hatchery, Qualicum Beach, British Columbia, Canada to the Pacific Biological Station, Nanaimo, BC on May 15 and 29, 2017, and February 26, 2018. For transport, juveniles were sedated with Aquacalm (0.1 mg L^-1^, Syndel) within containers having portable aerators and Vidalife mucous protectant (15 mL per L, Syndel). Juveniles were reared in communal circular tanks supplied with dechlorinated municipal freshwater (10 to 14°C), aeration, and artificial light set at the natural cycle until used in the experimental trials. Juveniles were feed 2% body weight ration of pellets (Bio-Oregon) every 1–2 days.

### 2.2. Experimental set-up

Juveniles were exposed to 18 groups (Figure 1), composed of all possible combinations of treatments for salinity (freshwater at 0 PSU, brackish at 20 PSU, and seawater at 28 or 29 PSU), temperature (10, 14, and 18°C), and dissolved oxygen (hypoxia at 4–5 mg L^-1^ and normoxia at > 8 mg L^-1^). Each group was represented by two (replicate) 30 L pot tanks with tight fitting lids to limit gas exchange for a total of 36 tanks. Seawater was pumped from nearby Departure Bay and disinfected using ultra-violet light. Both freshwater and seawater were provided at ambient temperatures as well as chilled and heated for a total of six available water sources. The experimental salinities and temperatures were achieved using combinations of these water sources; the brackish groups were a two-step process of water passing through a mixer to achieve 20 PSU then divided into metal coils within three water baths to achieve the temperatures. Water was then divided into an aeration column containing plastic media and another column containing a ceramic air stone to supply nitrogen. Both columns had additional stones for supplemental air if required. Water from the bottom of the columns was gravity fed to the tanks using 1.27 cm internal diameter PVC pipe or tubing. For the hypoxic group, nine temperature-dissolved oxygen probes were each connected to a Point Four RIU3monitor-controller (Pentair); monitor-controllers were collectively connected to a Point Four LC3central water system (Pentair). One probe was placed into one of the two hypoxic replicate tanks to maintain DO within 4–5 mg L^-1^ by continuously monitoring DO and turning the nitrogen regulator on or off as required. Nitrogen was supplied to the set-up using portable liquid units or compressed gas bottles (Praxair). Greater technical details on the experimental set-up are provided in Appendix 1.

**Figure 1:**
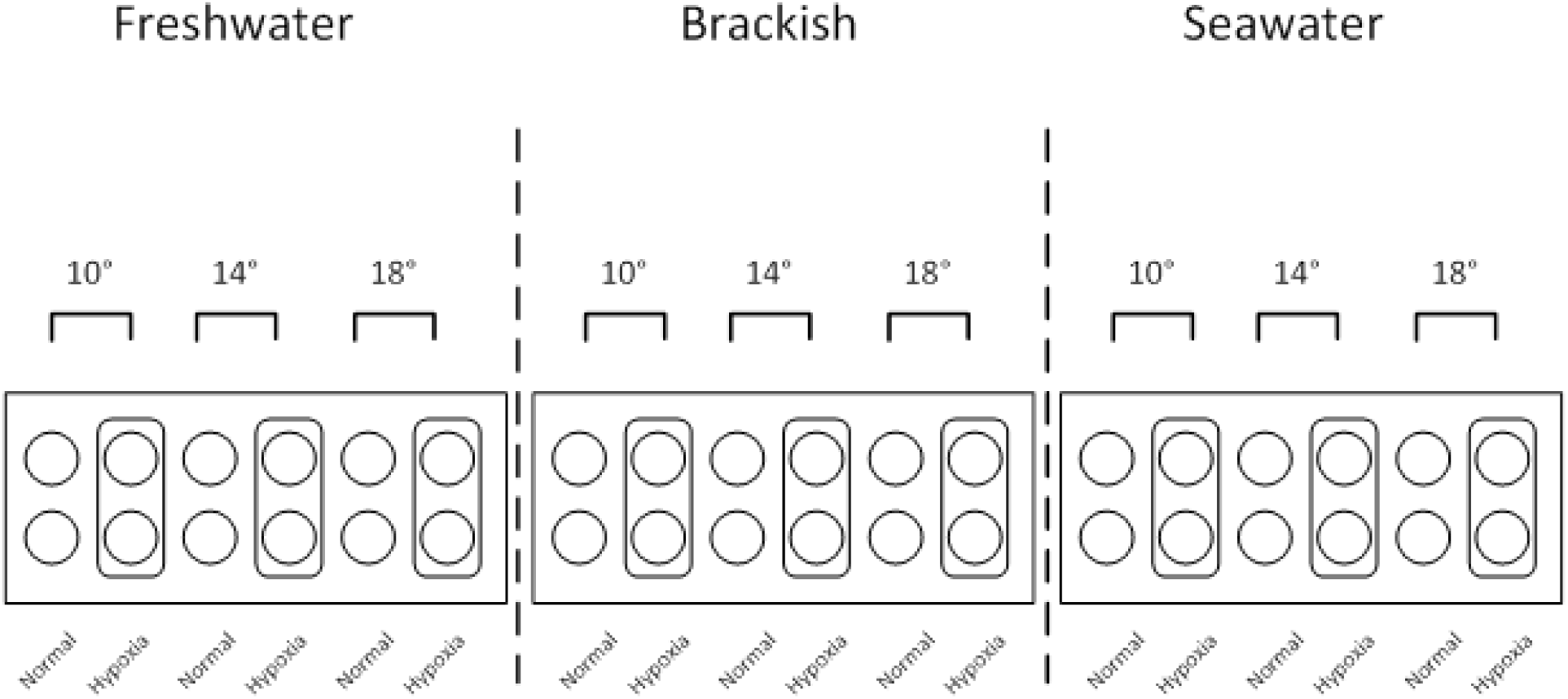
Experimental set-up and tank layout of the 18 groups for three treatments: three salinities by three temperatures by two dissolved oxygen concentrations. More technical details on the experimental set-up are provided in Appendix 1.

### 2.3. Trial set-up and welfare monitoring

Four trials were conducted spanning the pre-smolt, smolt, and de-smolt life stages based on timing (Table 1). The trial smolt statuses were also supported by expected differences in seawater mortality, skin pigmentation, body morphology, and NKA activity (see results). Six days before starting a treatment, juveniles were moved to the experimental tanks for acclimation. All juveniles in the communal tank were starved for 24 or 48 h, sedated with Aquacalm, and the water was treated with Vidalife. Juveniles were haphazardly selected; lightly anaesthetized in buffered TMS (100 mg L^-1^, Syndel); measured for fork length (± 0.1 cm) and mass (± 0.01 g); digitally photographed on their left side (Nikon Coolpix AW110) with a camera stand containing size and colour standards for later skin pigmentation and body morphology measurements; and allowed to recover with aeration before being moved to an experimental tank. Surfaces in contact with juveniles were sprayed liberally with concentrated Vidalife and care was taken so that juveniles were not out of water for more than 30 s. Each tank contained 12 (2017) or 16 (2018) juveniles that were supplied freshwater (13–14°C and > 8 mg L^-1^ DO) and fed 2% body weight ration per day (acclimation conditions). Body condition was calculated as 100 × mass ÷ length^3^ (Fulton, 1904).

**Table 1:** Summary of trial set-up using juvenile Chinook salmon *(Oncorhynchus tshawytscha*). Presented are the dates for changes in water conditions, mean body length and mass ± SD, and smolt status.

| Trial | Date | Length (cm) | Mass (g) | Status |
| --- | --- | --- | --- | --- |
| 1 | 12–13 March, 2018 | $4.8 \pm 0.3$ | $1.14 \pm 0.18$ | pre-smolt |
| 2 | 29 May, 2017 | $7.3 \pm 0.4$ | $4.14 \pm 0.70$ | smolt |
| 3 | 19–20 June, 2017 | $7.6 \pm 0.4$ | $4.67 \pm 0.78$ | de-smolt |
| 4 | 14–15 August, 2017 | $9.6 \pm 0.5$ | $9.40 \pm 1.66$ | de-smolt |

Juveniles were starved for 24 or 48 h again prior to changing the water conditions, which occurred over one or two days. On day 1, seawater was introduced over 2–3 h in the morning, temperature was changed by 2°C in the early afternoon, and DO was set to 6.5–8 mg L^-1^ at 3 pm using the LC3 central water system. On day two, temperature was changed by the remaining 2°C and DO was set to 4–5 mg L^-1^ in the morning. Juveniles were fed a ration the next morning. Juveniles were exposed to the full treatment conditions for six days and their welfare was checked at least every 4 h between 8 am and 8 pm. During visual checks, any dead juveniles (no gill ventilation) were removed and moribund juveniles (loss of equilibrium but gill ventilation) were euthanized in an overdose of TMS (250 mg L^-1^, buffered for freshwater groups) using water of the same salinity and temperature. These juveniles were lethally sampled as described below.

### 2.4. Data collection

After two days of full treatments, 4 of 12 juveniles (2017) were anaesthetized or 4 of 16 juveniles (2018) were euthanized per tank to examine short-term gill gene expression. Juveniles were anaesthetized or euthanized using water of the same salinity and temperature, measured, and photographed. The anaesthetized juveniles were placed on their back in a cushioned trough under a lamp light and then a small piece of gill tissue (1 mm 1 mm) was removed from the tip of filaments using sterilized scissors. Juveniles were then tagged below the dorsal fin with visible implant elastomer (Northwest Marine Technology) and allowed to recover in aerated treatment water before being returned to their tank. Care was taken so that juveniles were not out of aerated treatment water for more than 30 s. Surfaces in contact with juveniles were sprayed liberally with Vidalife. Tools were disinfected between juveniles using 3–5 min of 10% bleach and immersion in 95% ethanol and flame, with tools being allowed to cool before use on the next juvenile. The gill tissue for all juveniles was placed in RNAlater (Invitrogen) for 24 h in a 4°C fridge then stored in a -80°C freezer until used for gene expression measurements.

After four days of full treatments, juveniles were fed a ration in the morning and examined for behaviour related to performance between 11 am and 3 pm. For the first chronological trial (trial 2 in May 2017), behavioural observations were collected for each tank by a trained observer that removed the tank cover and watched for 3 min. The observer paid attention to relative startle response (i.e. swimming speed after disturbance) and gill ventilation rate. For the later trials (chronologically trial 3, 4, and 1), 10 min video clips were collected for each tank using small cameras within weighted underwater cases (GoPro Hero or Monster Digital Villain). Raw video clips were brightened and trimmed (first and last 30 s removed) using GoPro Studio 2.5. Edited video clips were renamed to hide the tank identity and examined by two trained observers. The observers calculated the gill ventilation per min for three individuals per tank.

After six days of full treatments, remaining juveniles were sampled for tissues to examine long-term effects. Juveniles were euthanized using water of the same salinity and temperature, measured, and photographed (Nikon Coolpix AW110 or Olympus Tough TG-3). Blood was collected within 5 min using caudal severance and a capillary tube. Blood was centrifuged (2,000 RCF) for 5 min to separate plasma and red blood cells then immediately frozen using dry ice. Gill tissue from the right side was placed into a cryovial then immediately frozen using liquid nitrogen for Na^+^/K^+^-ATPase activity. For examining long-term gene expression, the remaining gill tissue from the left side was placed in RNAlater as described above. The rest of the body was individually placed into a plastic bag then frozen immediately with dry ice. Tissues were stored in a -80°C freezer until used for measurements.

### 2.5. Gene expression

A total of 441 individuals were examined for gill gene expression. The data were composed of 2–3 live individuals per tank for each of the four trials (316 juveniles) that had been exposed for six days and 125 dead or moribund juveniles (mostly from trials 1, 3, and 4, i.e. pre-smolts and de-smolts). Gill samples were also collected after two days, but we focussed our detailed analysis on six day samples because the expression of a subset of these two day samples (representing 33 individuals of trial 1) was generally similar to the six day samples (Appendix 2). Gill tissue was homogenized in TRIzol (Ambion) and BCP reagent using stainless steel beads on a MM301 mixer mill (Retsch Inc.). RNA was extracted from the homogenate using the ‘No-Spin Procedure’ of MagMAX-96 Total RNA Isolation kits (Ambion) and a Biomek FXP automation workstation (Beckman-Coulter). RNA yield was quantified using the A_260_ value and extracts were normalized to 62.5 ng mL^-1^. Normalized RNA was reverse transcribed to cDNA using SuperScript VILO synthesis kits (Invitrogen). Normalized RNA and cDNA were stored at -80°C between steps.

Gene expression was examined for three housekeeping genes, i.e. Coil-P84, 78d16.1, MrpL40 (Miller et al., 2017a), and 93 candidate genes, i.e. 37 for salinity, 33 for temperature, and 23 for dissolved oxygen. Each gene expression chip contained these 96 assays, dilutions of a cDNA pool (five-fold serial 1 to 1/3,125), and an inter-chip calibrator sample. Following Fluidigm prescribed methods, target cDNA sequences were enriched using a specific target amplification (STA) method that included small concentrations of the 96 assay primer pairs. Specifically, for each reaction, 3.76 μL 1X TaqMan PreAmp master mix (Applied Biosystems), 0.2 μM of each of the primers, and 1.24 μL of cDNA. Samples were run on a 14 cycle PCR program, with excess primers removed with EXO-SAP-IT (Affymetrix), and diluted 1/5 in DNA suspension buffer. The diluted samples and assays were run in singleton following the Fluidigm platform instructions. For sample reactions, 3.0 μL 2X TaqMan mastermix (Life Technologies), 0.3 μL 20X GE sample loading reagent, and 2.7 μL STA product. For assay reactions, 3.3 μL 2X assay loading reagent, 0.7 μL DNA suspension buffer, 1.08 μL forward and reverse primers (50 μM), and 1.2 μL probe (10 μM). The PCR was 50°C for 2 min, 95°C for 10 min, followed by 40 cycles of 95°C for 15 s, and then 60°C for 1 min. Data were extracted using the Real-Time PCR Analysis Software (Fluidigm) using Ct thresholds set manually for each assay.

For determining the optimal normalization gene(s) from the three housekeeping (HK) candidates, gene expression of each gene was first linearly transformed (efficiency ^minimum Ct - sample Ct^). Values were then used in the NormFinder R function (Andersen et al., 2004) with assemblages for the 18 groups to identify the gene or gene pair with the lowest stability (standard deviation). Sample gene expression was normalized with the ∆∆Ct method (Livak and Schmittgen, 2001) using the mean (for single gene) or geometric mean (for pair of genes) and the species-specific calibrator sample. Gene expression was then log transformed: log_2_(2^-∆∆Ct^).

Eighty-seven out of 93 candidate genes were used in analyses. The gene assays for glu_2, LDH_3, Myo_1 (dissolved oxygen) and CIRBP_10 (temperature) were removed because of poor efficiency using the present study samples. Also removed were Tuba1a_16 (temperature) because the assay did not work for half of the samples and HIF1A_4 (dissolved oxygen) because this assay was not represented for smolts in trial 2 and there was no pattern with the treatment for the remaining trials (data not shown).

### 2.6. Na^+^/K^+^-ATPase activity and plasma variables

Juveniles were also measured for physiological variables and infectious agent presence and loads. All individuals used for gene expression analysis were also examined for gill Na^+^/K^+^-ATPase enzyme (NKA) activity; a subset of individuals from trials 2, 3, and 4 were examined for plasma lactate, glucose, and chloride concentrations. NKA activity was measured using the method of McCormick (1993). Plasma lactate, glucose, and chloride concentrations were measured using the methods of Farrell et al. (2001). Although plasma cortisol is associated with stress, we did not examine this hormone because it is naturally elevated during seawater acclimation alone (McCormick, 2001; Young et al., 1995). For 79 dead or moribund juveniles from trials 2, 3 and 4, we examined a mixed-tissue (i.e. gill, liver, heart, kidney, and spleen) for the presence and load of 47 infectious agents known or suspected to cause diseases in salmon, using the methods of Miller et al. (2016).

### 2.7. Skin pigmentation and body morphology

A subset of individuals was examined for skin pigmentation and body morphology using the photographs and methods described by Houde et al. (2015): half the individuals from initial set-up and half the individuals after six days of treatment (including all those examined for gene expression) per tank for each trial. Briefly, for skin pigmentation, LAB colour space values of the anterior region, posterior region (both covering the lateral line), and caudal fin were subjected to principal component analysis. For body morphology, 21 landmarks were subjected to relative warp analysis using *tpsRelw32* software (Rohlf, 2017).

### 2.8. Statistical analysis of variables

Analyses were performed using R 3.3.3 (R Core Team) at a significance level of *α* = 0.05. To identify any potential treatment effects, the expression of each gene was subjected to forward model selection using AIC criteria, including *salinity*, *temperature*, and *oxygen* with their interactions. Mortality (proportion of dead and euthanized moribund juveniles) per tank was examined using binomial generalized linear models with a quasi link. The significance of effects for mortality was examined using analysis of deviance (ANODEV) F-tests and ANOVA for remaining variables. Tukey’s post hoc tests examined the significance of contrasts. Gill ventilation rate, NKA activity, plasma, body size, skin pigmentation, and body morphology variables were examined for simple Pearson correlations with gene expression patterns (PC1 and PC2) described below.

### 2.9. Statistical analysis to identify biomarkers

We aimed to identify clusters containing 8 to 12 genes (biomarkers) that collectively co-vary in response to salinity, temperature, and dissolved oxygen treatments because such clusters may be conserved across species and studies (e.g. in viral disease development, Miller et al., 2017a). To this end, the entire dataset was divided into a two-thirds training set and a one-third testing set. The training set was subjected to supervised gene shaving of the candidate genes for each treatment (Hastie et al., 2000) using GeneClust JS (Do et al., 2003). We selected the first, first two, or first three clusters to be in range of 8 to 12 identified biomarkers. Two validation approaches were implemented. First, identified genes were subjected to principal component analysis (PCA) for the training set. This PCA was then applied to the testing set for visualization of unsupervised group separation within treatment using the *fviz-pca* function of the *factoextra* R package (Kassambara and Mundt, 2017). The function provided 95% confidence ellipses for groups of the training set. Second, classification ability of the groups examined by subjecting the identified biomarkers to linear discriminant analysis (LDA) using the training set, followed by determining classification performance on the testing set. A similar approach was used for examining mortality (dead or moribund vs. live) using all the candidate genes.

## 3. Results

### 3.1. Mortality

During trial 3, the two seawater-18°C-hypoxia tanks were terminated (11 of 24 juveniles were dead or moribund); it was discovered that the DO sensor was still at a freshwater setting causing DO to be lower than intended for seawater. Remaining 13 juveniles were ethically euthanized. This test group was then restarted with new juveniles 10 days after the original start date and was included in the analysis of trial 3.

Mortality data were analyzed separately for non-biopsied and biopsied juveniles because gill biopsy after two days increased overall mortality by 2.3 times compared to non-biopsied juveniles (ANODEV, *p* = 0.003; Table 2). As expected for different smolt statuses in seawater, overall mortality was low for non-biopsied smolts in trial 2 (*n* = 2) relative to pre-smolts in trial 1 (46), de-smolts in trial 3 (28), and de-smolts in trial 4 (28), and significantly different from the remaining trials (Tukey’s poc hoc test, *p <* 0.005 for all). In particular, 11 of 16 pre-smolts in trial 1 were dead or moribund in one replicate tank for seawater-18°C-hypoxia; this tank was terminated and remaining five juveniles were ethically euthanized. Consequently, mortality of non-biopsied juveniles was associated with salinity only for trials 1, 3, and 4 based on model selection (ANODEV, *p* <0.001 for all; Table 2); mortality was significantly higher in seawater (trial 1 = 46 juveniles, trial 3 = 22 juveniles, and trial 4 = 23 juveniles) than freshwater or brackish (Tukey, *p <* 0.016).

**Table 2:** Summary of juvenile Chinook salmon mortality by trial and groups within treatments. Presented are the non-biopsied (X) and biopsied (G) mortality. The juvenile counts were combined for both replicate tanks of a group. Groups had a total of 32 juveniles for trial 1 and 24 juveniles for trials 2 to 4. Gill biopsies were conducted on four anaesthetized juveniles per group of trials 2 to 4 that were later returned to their tanks, whereas four juveniles were euthanized in trial 1 for the biopsies. Salinity treatment symbols are SW, BW, and FW for seawater, brackish, and freshwater groups. Temperature treatment symbols are 10, 14, and 18 for °C groups. Dissolved oxygen treatment symbols are N for normoxia and H for hypoxia groups.

| Group | Trial 1 | Trial 2 |  | Trial 3 |  | Trial 4 |  |
| --- | --- | --- | --- | --- | --- | --- | --- |
|  | X | X | G | X | G | X | G |
| FW10N |  |  |  | 1 | 2 |  | 1 |
| FW10H |  | 1 | 1 |  | 3 | 1 | 1 |
| FW14N |  |  |  |  | 3 | 1 | 4 |
| FW14H |  |  |  |  | 5 |  | 2 |
| FW18N |  |  | 1 | 1 | 3 |  | 2 |
| FW18H |  |  |  |  | 3 |  |  |
| BW10N |  |  |  |  | 2 |  |  |
| BW10H |  |  |  |  | 2 |  |  |
| BW14N |  |  |  | 1 |  |  |  |
| BW14H |  |  |  | 2 | 2 | 1 | 1 |
| BW18N |  |  | 1 |  | 1 |  | 1 |
| BW18H |  |  |  | 1 | 2 | 2 | 1 |
| SW10N | 6 |  |  | 2 | 2 | 3 |  |
| SW10H | 6 |  |  | 1 | 3 | 6 | 1 |
| SW14N | 2 | 1 | 1 | 3 |  | 5 | 2 |
| SW14H | 7 |  |  | 3 | 4 | 2 | 2 |
| SW18N | 7 |  |  | 6 | 2 | 7 | 2 |
| SW18H | 18 |  |  | 7 | 2 | 2 |  |

No gill biopsies were taken for pre-smolts in trial 1. Mortality in the three other trials with biopsy were significantly different from each other (Tukey, *p <* 0.005): smolts in trial 2 (*n* = 4); de-smolts in trial 3 (41); and de-smolts in trial 4 (20). Mortality of biopsied juveniles was associated with oxygen only for trial 3 (ANODEV, *p* = 0.043); mortality was higher for hypoxia (26 juveniles) than normoxia (15 juveniles). In terms of the pathogen loads for the 79 dead or moribund juveniles, infectious agents were limited to detections of only 2 out of 47 candidates in only some individuals at loads > 1,000 copies per μg RNA: bacteria *Candidatus Branchiomonas cysticola* and *Flavobacterium psychrophilum* (data not shown).

### 3.2. Trial differences in body variables and NKA activity

All body variables in all trials differed significantly relative to initial values (ANOVA, *p <* 0.001 for all; beanplots in Appendix 3). Not surprisingly, body length and mass increased with the trial date (Figure 2). As expected for different smolt statuses, there were differences in skin pigmentation and body morphology across trials. Anterior and posterior brightness and caudal fin darkness and yellowness increased from pre-smolts in trial 1 to smolts in trial 2; anterior brightness and caudal fin yellowness then decreased for de-smolts in trials 3 and 4 with smaller changes for caudal fin darkness and posterior brightness. Body condition, elongation and thickness, back roundness, and caudal peduncle length increased from pre-smolts in trial 1 to smolts in trial 2; body thickness then decreased for de-smolts in trials 3 and 4 with smaller changes for remaining variables.

**Figure 2:**
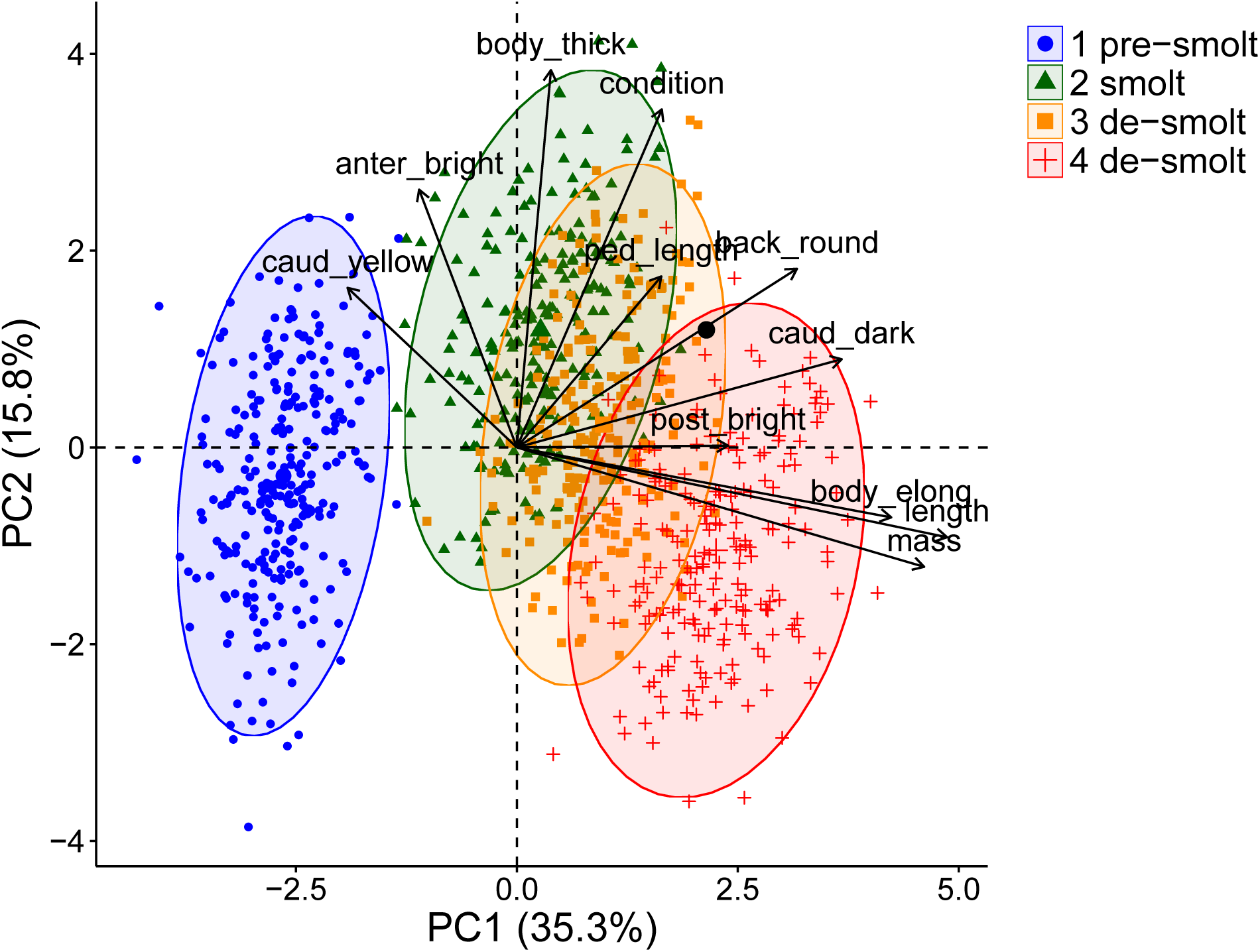
Canonical plots of the first two principal components of initial body size, skin pigmentation, and body morphology variables for the four trials. Trials are presented in the order of smolt development, and smolt statuses are based on seawater survival. Beanplots of initial trial differences and statistics for these variables are presented in Appendix 3.

We considered the first four skin pigmentation principal components (PCs) and first four body morphology relative warp axes (RWs) that were biologically meaningful. Skin pigmentation was represented by anterior brightness (PC1, 45.8%), caudal fin darkness (PC2, 25.0%), posterior brightness (PC3, 13.7%), and caudal fin yellowness (PC4, 9.7%). Body morphology was represented by body elongation or distance between fins (RW1, 19.6%), back roundness (RW2, 12.4%), caudal peduncle length (RW4, 8.4%), and body thickness or distance of the mid-section from top to bottom (RW5, 5.9%); RW3 (11.5%) was not considered because it represented body flexing during anaesthesia.

As expected in freshwater, NKA activity of smolts in trial 2 was higher than pre-smolts in trial 1 (one-sided Student’s t-test, *p* = 0.037; Table 3). However, this was not the case for smolts compared to both de-smolts trials (trial 3 *p* = 0.155 and trial 4 p = 0.312). Interestingly, in seawater, NKA activity of smolts was higher than the remaining trials (trial 1 *p* = 0.001, trial 3 *p* = 0.045, and trial 4 *p* = 0.029). There was a similar pattern in brackish except for the last de-smolt trial (trial 1 *p* = 0.047, trial 3 *p* = 0.005, and trial 4 *p* = 0.173).

**Table 3:** Summary of gill Na^+^ /K^+^-ATPase activity by trial and salinity. Presented are the mean ± SD. Na^+^/K^+^-ATPase activity units are μmol ADP (mg protein)^-1^ h^-1^.

| Salinity | Trial 1 | Trial 2 | Trial 3 | Trial 4 |
| --- | --- | --- | --- | --- |
| freshwater | $3.9 \pm 1.5$ | $5.0 \pm 2.5$ | $4.3 \pm 2.2$ | $4.7 \pm 2.9$ |
| brackish | $5.5 \pm 2.4$ | $6.7 \pm 2.3$ | $4.8 \pm 2.6$ | $6.0 \pm 2.8$ |
| seawater | $7.3 \pm 2.5$ | $9.6 \pm 2.8$ | $8.0 \pm 4.3$ | $8.1 \pm 3.3$ |

### 3.3. Salinity, temperature, and dissolved oxygen biomarkers

Results of forward model selection and figures of gene expression for the 87 candidate genes that remained after quality assurance checks are presented in Table 4 and Appendix 4. Salinity primarily influenced the expression of 20 of 37 candidate salinity genes and temperature also influencing 10 of these 20 genes. Temperature primarily influenced the expression of 25 of 31 candidate temperature genes and salinity influence 19 of these 25 genes. Oxygen primarily influenced only one (i.e. HIF1A_6) of 19 candidate dissolved oxygen genes, and oxygen followed after salinity or temperature for three genes (i.e. ALD_1, Enolase_2, and HemA1_1). Including the candidate temperature genes, oxygen primarily influenced one gene (i.e. COX6B1_19) and followed temperature and salinity for one other gene (i.e. HSP90alike_6). The biomarkers specific to each treatment under multi-stressor conditions and smolt status are described below.

**Table 4:** Summary of model selection for 87 candidate genes associated with responses to salinity, temperature, and dissolved oxygen after six days. Based on AIC forward selection using linear models and significance of *p <* 0.05. Variables considered were salinity, temperature, oxygen, and their interactions.

| Gene name | Assay name | Model |
| --- | --- | --- |
| *Salinity* |  |  |
| Beta actin | ACTB_v1 | temperature + salinity |
| Carbonic anhydrase 4 | CA4_v1 | salinity |
| C-C chemokine 19 | CCL19_v1 | salinity |
| C-C chemokine 4 | CCL4_v1 | salinity |
| Cystic fibrosis transmembrane conductance regulator I | CFTR-I_v1 | salinity + temperature |
| C-type lectin domain family 4 member M | CLEC4M_v1 | temperature |
| Cytochrome P450 2K1 | CYP2K1_v2 | salinity |
| Elongation factor 2 | EEF2_v1 | salinity + temperature |
| Exonuclease 1 | EXO1_v1 | temperature + salinity |
| FK506-binding protein 5 | FKBP5_v1 | salinity |
| Formin-like protein 1 | FMNL1_v1 | temperature + salinity |
| Growth hormone receptor 1 | GHR1_v1 | temperature + salinity |
| Hemoglobin subunit $\alpha$ | HBA_v1 | temperature + salinity + oxygen |
| Hemoglobin subunit $\alpha$(true HBA) | HBA_t_v1 | temperature |
| Interferon-induced protein 44 | IFI44_v1 | salinity |
| Interleukin-12 beta | IL12B_v1 | temperature |
| DNA replication licensing factor MCM4 | MCM4_v1 | temperature |
| Mitochondrial pyruvate carrier 1 | MPC1_v1 | temperature + salinity + temperature × salinity |
| Membrane-spanning 4-domains A-4A | MS4A4A_v1 | nothing |
| Nicotinamide phosphoribosyltransferase | NAMPT_v1 | temperature + salinity |
| NADH dehydrogenase 1 beta subcomplex subunit 2 | NDUFB2_v1 | salinity + temperature |
| NADH dehydrogenase 1 beta subcomplex subunit 4 | NDUFB4_v1 | salinity + temperature |
| Na/K ATPase α-1a (freshwater) | NKAa1-a_v2 | salinity + temperature |
| Na/K ATPase α-1b (seawater) | NKAa1-b_v2 | salinity + temperature + salinity × temperature |
| Glucocorticoid receptor 1 | NR3C1_v1 | salinity |
| Serine/threonine-protein kinase PLK2 | PLK2_v2 | salinity + temperature + oxygen |
| Prolactin receptor | PRLR_v1 | salinity + temperature |
| Regulator of G-protein signaling 21 | RGS21_v1 | salinity |
| Rhesus blood group-associated glycoprotein | RHAG_v1 | salinity + temperature |
| 60S ribosomal protein L31 | RPL31_v1 | salinity + temperature |
| Monocarboxylate transporter 10 | SLC16A10_v1 | nothing |
| Thyroid hormone receptor B | THRB1_v2 | temperature + salinity |
| T-cell receptor alpha | TRA_v1 | nothing |
| Translocator protein | TSPO_v2 | temperature + salinity |
| Tubulin, alpha 8 like 2 | TUBA8L2_v1 | temperature + salinity |
| Ubiquitin-like modifier-activating enzyme 1 X | UBA1_v1 | salinity |
| Wiskott-Aldrich Syndrome protein | WAS_v1 | salinity |
| *Temperature* |  |  |
| AP-3 complex subunit sigma-1 | AP3S1_24 | temperature + salinity |
| Cold-inducible RNA-binding protein | CIRBP_16_v2 | temperature |
| Cytochrome C oxidase | COX6B1_19 | oxygen + salinity + temperature + salinity × temperature |
| Elongation factor 2 | ef2_14 | temperature + salinity |
| Elongation factor 2 | ef2_3 | temperature + salinity |
| Eukaryotic initiation factor 4A-II | EIF4A2_14_v1 | temperature + salinity |
| Eukaryotic translation initiation factor | Eif4enif1_12 | temperature + salinity |
| FK506-binding protein 10 precursor | FK506_19_v2 | temperature |
| FK506-binding protein 10 precursor | FK506_3&6_v2 | temperature |
| Heat shock cognate 70 kDa protein | hsc71_9 | temperature + salinity |
| Heat shock 70 kDa protein | hsp70_6 | salinity + temperature |
| Heat shock protein 90 alpha | hsp90a_15_v2 | temperature + salinity + temperature × salinity |
| Heat shock protein 90 beta | HSP90ab1_15_v1 | temperature + salinity |
| Heat shock protein 90 alpha like | HSP90alike_6 | temperature + salinity + oxygen + temperature × salinity |
| Isocitrate dehydrogenase [NAD] subunit beta, mitochondrial precursor | IDH3B_12_v2 | salinity |
| Keratinocytes-associated transmembrane protein 2 precursor | KCT2_13 | temperature + salinity |
| Mitogen-activated protein kinase kinase kinase 14 | Map3k14_3 | temperature + salinity |
| Mannose-P-dolichol utilization defect 1 protein | MPDU1_7 | salinity |
| Serine/threonine-protein kinase Nek4 | Nek4_12 | temperature + salinity |
| Protein DJ-1 | park7_22 | nothing |
| Protein disulfide-isomerase A4 precursor | PDIA4_19_v1 | nothing |
| Sec1 family domain-containing protein 1 | SCFD1_1 | temperature + salinity |
| Selenoprotein W | sepw1_11_v1 | temperature + salinity |
| Serpin H1 precursor (HSP47) | SERPIN_20_v1 | temperature + salinity |
| Serpin H1 precursor (HSP47) | SERPIN_9 | temperature + salinity |
| Splicing factor, arginine/serine-rich 2 | SFRS2_3_v2 | temperature + salinity |
| Splicing factor, arginine/serine-rich 9 | Sfrs9_1_v2 | temperature + salinity |
| Tubulin alpha chain, testis-specific | Tuba1a_11 | temperature |
| Ubiquitin-conjugating enzyme E2 Q2 | UBE2Q2_26 | temperature |
| Transmembrane protein 185-like | zgc:63572_11 | temperature + salinity |
| Zinc finger MYND domain-containing protein 11 | Zmynd11_3 | temperature |
| *Dissolved oxygen* |  |  |
| Fructose-bisphosphate aldolase A1 | ALD_1 | temperature + oxygen |
| Fructose-bisphosphate aldolase A1 | ALD_4_v1 | salinity |
| Enolase | Enolase_2 | temperature + salinity + oxygen |
| Glycogen phosphorylase | GIPh_1 | temperature + salinity |
| Glycogen phosphorylase B | GIPh_4 | temperature |
| Solute carrier family 2, facilitated glucose transporter member 1 | glu1 | temperature + salinity |
| Hemoglobin | HemA1_1 | salinity + temperature + oxygen |
| Heme oxygenase 1 | HemOxi1_2 | temperature + salinity |
| Heme oxygenase 1 | HemOxi1_3 | salinity |
| Heme oxygenase 2 | HemOxi2_1 | temperature |
| Hypoxia-inducible factor 1-alpha-like | HIF1A_6 | oxygen<br>+ salinity + temperature |
| Hypoxia inducible factor 1 alpha | HIF1A_7 | temperature + salinity |
| Insulin-like growth factor binding protein-1 | IGFBP_10_v1 | nothing |
| Insulin-like growth factor binding protein-like 1 | IGFBP1 | nothing |
| L-lactate dehydrogenase B-A chain-like | LDH_1 | temperature |
| Neuroglobin | Ngb1_2 | nothing |
| Phosphoglycerate kinase | PgK_5 | temperature |
| Phosphoglycerate kinase | PgK3_v1 | temperature + salinity |
| Vascular endothelial growth factor A | VEGFa_1 | temperature + salinity |

Salinity biomarkers had two clusters, totalling 11 genes. Cluster one contained CA4 and CFTR-I; cluster two contained CCL19, CCL4, FKBP5, IFI44, NAMPT, NDUFB2, NKAa1-a, PRLR, and RGS21 (Figure 3a,b). Examining the PCA on the training set applied to the testing set, a few dead or moribund individuals fell outside the 95% confidence ellipses. Most of the separation for the salinity groups was along PC2. Freshwater was more separated from brackish and seawater, but brackish and seawater were similar. Mortality and smolt status were better associated with PC1: smolt at one extreme, pre-smolt and de-smolt in between, and mortality at the other extreme. The direction of mortality in seawater was associated with higher expression of FKBP5, NKAa1-a, PRLR, and RGS21, as well as decreased expression of CFTR-I. Using all 11 biomarkers in LDA on the training set, testing set classification ability for freshwater was 100%, seawater was 81%, and brackish was 57% (Table 5). Classification ability was almost perfect (99%) if seawater and brackish groups were combined.

**Figure 3:**
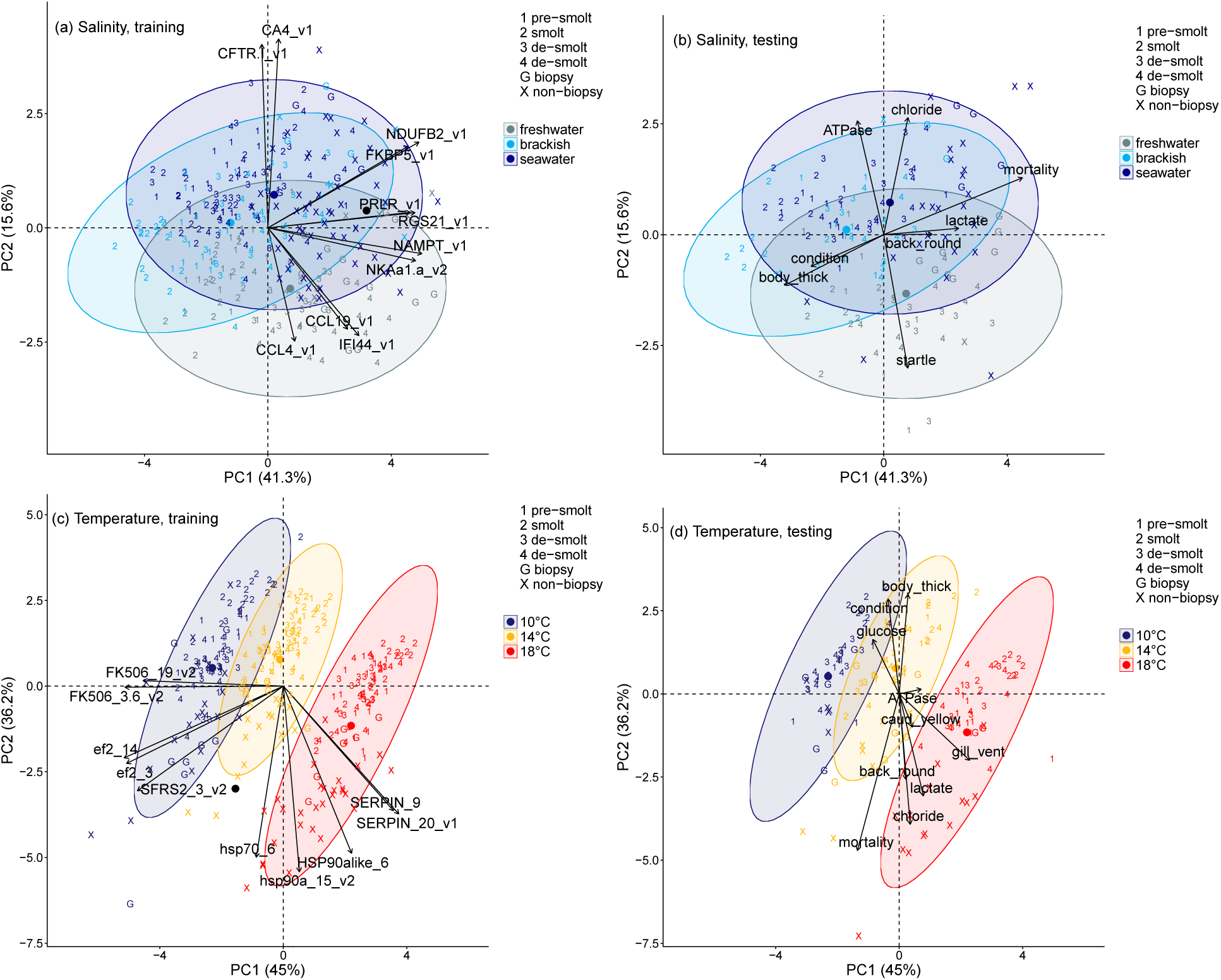
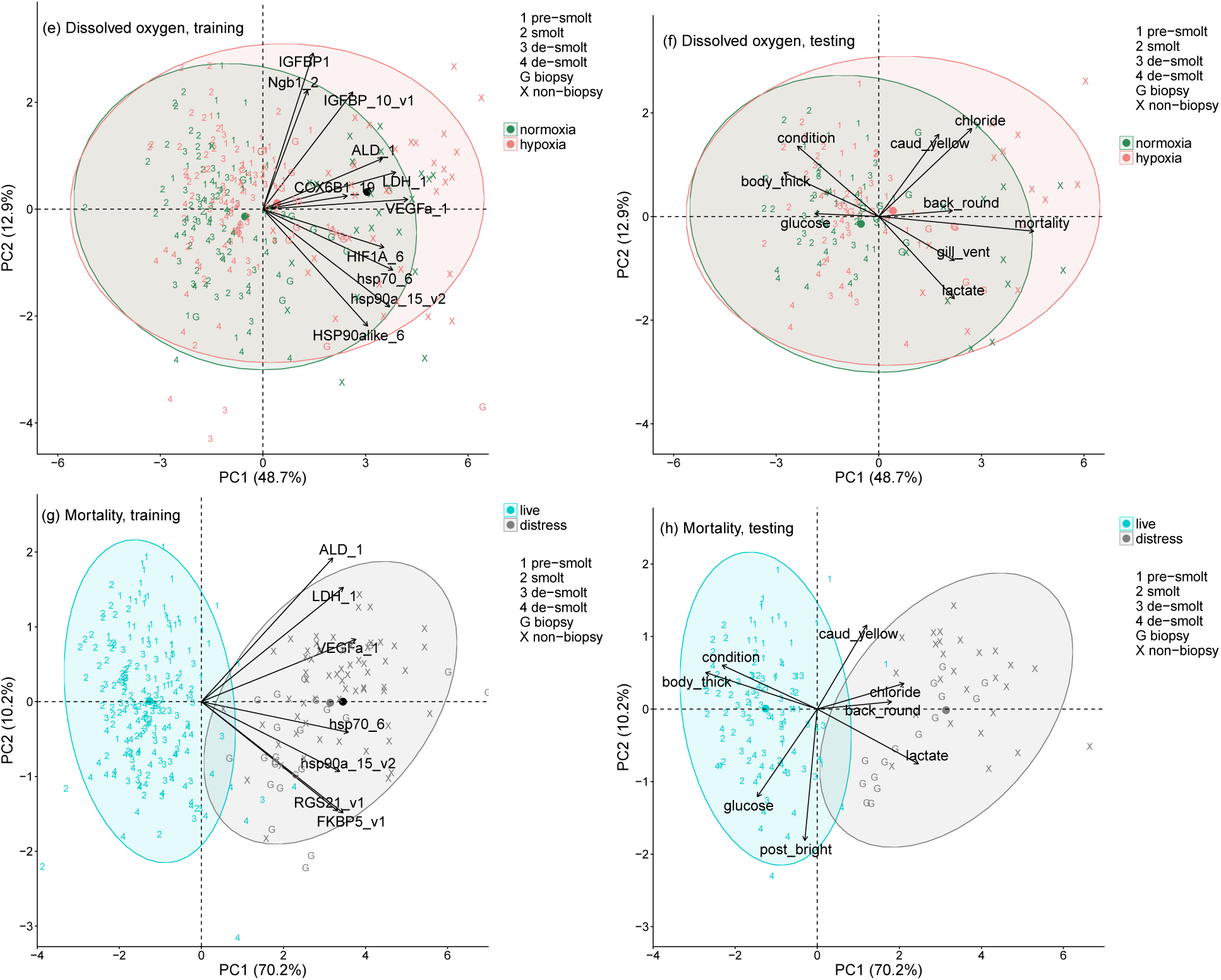
Canonical plots of the first two principal components of the identified biomarkers for salinity, temperature, dissolved oxygen, and mortality. Principal component analysis was performed on the training set (left panel) and then applied to the testing set (right panel). Ellipses represent 95% confidence areas for the groups within treatments using the training set; centroids are represented by the largest point of the same colour. Numbers are live juveniles and X and G are dead or moribund juveniles. Arrows (left panel) represent loading vectors of the biomarkers using the training set; centroid is a black circle. Arrows (right panels) represent loading vectors of variables using the subset of available data across the training and testing sets; statistics of treatment and mortality effects on variables are presented in Appendix 3, and variable correlations with gene expression in Table S2.

**Table 5:**
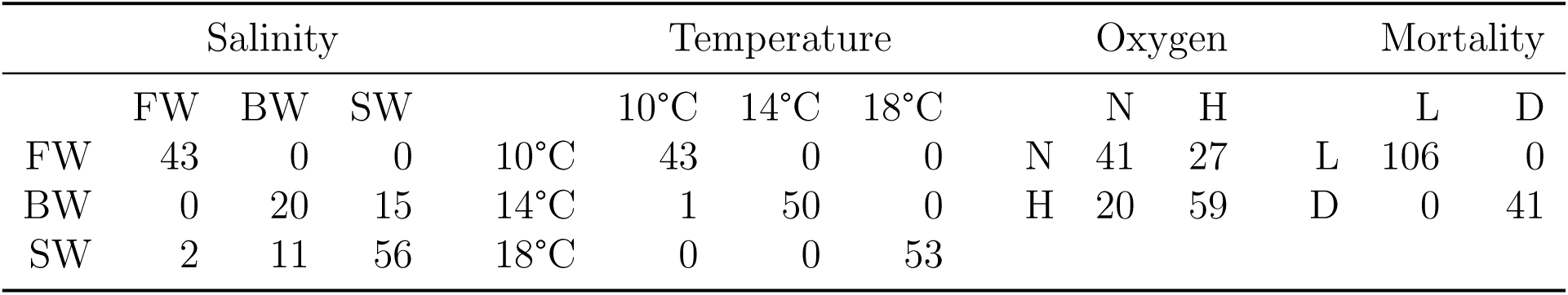
Classification ability of the groups within treatments and mortality using the identified biomarkers. Biomarkers identified by gene shaving were placed into a linear discriminant analysis (LDA) for classifying groups within each treatment and mortality. LDA training set used two-thirds of the entire dataset. Presented are the classifications on the remaining one-third testing set. Columns are the LDA classification groups and rows are the real group. Salinity symbols are FW for freshwater, BW for brackish, and SW for seawater groups. Dissolved oxygen symbols are N for normoxia and H for hypoxia groups. Mortality symbols are L for live (persisted six days) and D for distress (dead or moribund) groups.

Temperature biomarkers had three clusters, totalling 10 genes. Cluster one contained HSP90alike6, SERPIN_20, and SERPIN_9; cluster two contained ef2_14, ef2_3, hsp70_6, hsp90a_15, and SFRS2_3; cluster three contained FK506_19 and FK506_3.6 (Figure 3c,d). The PCA displayed visible separation of the temperature groups with little overlap of the ellipses, and there were a few dead or moribund individuals falling outside the ellipses. The highest temperature (18°C) separated strongly from 10 and 14°C. Classification ability approached perfection at all three temperatures: lowest was 98% for 14°C (Table 5).

Dissolved oxygen biomarkers had two clusters, totalling 11 genes. For this analysis, we considered all the candidate temperature and dissolved oxygen genes because forward model selection indicated issues separating normoxia and hypoxia using only the candidate dissolved oxygen genes. Cluster one contained ALD_1, COX6B1_19, HIF1A_6, hsp70_6, IGFBP_10, IGFBP1, LDH_1; and cluster two contained hsp90a_15, HSP90alike_6, Ngb1_2, and VEGFa_1 (Figure 3e,f). Regardless, there was a poor separation between normoxia and hypoxia. Classification ability for normoxia was 60.3% and hypoxia was 74.7% (Table 5).

Across the 18 treatment groups, average classification ability (using the three treatment classifiers) was 55.2%, with primary classification issues for normoxia and hypoxia (Appendix 5). Removing the dissolved oxygen treatment (regrouping as nine salinity by temperature groups), the average classification ability increased to 79.5%, with secondary classification issues for brackish and seawater. Combining brackish with seawater, the average classification ability increased to 98.0% (freshwater-brackish/seawater and three temperatures).

### 3.4. Mortality biomarkers

Dead and moribund juveniles had visually similar gene expression (Appendix 4) and were combined. Mortality biomarkers across all 18 groups had one cluster, totalling 7 genes. This cluster contained: ALD_1, FKBP5, hsp70_6, hsp90a_15, LDH_1, RGS21, and VEGFa_1 (Figure 3g,h). The PCA showed ellipses with little overlap and a few individuals falling outside the ellipses. There was good separation for live (persisted six days) and distress. Classification ability was 100% for both live and distressed juveniles (Table 5).

### 3.5. Relationship to physiological and body variables

The gene expression patterns displaying the largest separation for groups within treatments, i.e. salinity (PC2), temperature (PC1), and mortality (PC1), were generally not strongly correlated with the body variables (Figure 3a,c, correlation values which were *r >* -0.3 or *r <* 0.3 and significance in Table S2). Salinity (PC2) had significant positive correlations with both NKA activity and chloride concentrations, as expected, and a negative correlation with relative startle response. Temperature (PC1) was positively correlated gill ventilation rate, as expected, and with NKA activity. Mortality (PC1) was positively correlated with plasma lactate and chloride concentrations, caudal fin yellowness, and back roundness; and negatively correlated with plasma glucose concentrations, body condition, and thickness (Figure 3g).

In contrast to salinity, temperature, and mortality, gene expression patterns for dissolved oxygen (PC1 and PC2) did not separate groups well, and there was a larger number of correlations with body variables. Dissolved oxygen (PC1) was correlated with mortality, gill ventilation rate, plasma lactate, glucose, and chloride concentrations, as well as body condition, thickness, back roundness, and caudal fin yellowness. We also examined each variable using model selection, including salinity, temperature, oxygen, and their interaction in Appendix 3. Juveniles from hypoxia had higher gill ventilation rate (ANOVA, *p <* 0.001) and anterior region brightness (p = 0.039) than normoxia.

## 4. Discussion

Through laboratory challenges whereby juvenile ocean-type Chinook salmon were exposed to multi-stressor conditions for six days, we resolved gill gene expression biomarkers specific to salinity and temperature treatments, but not the dissolved oxygen treatment. These biomarkers showed very high classification ability (99%) for freshwater vs. saline (brackish or seawater) and three temperatures (10, 14, and 18°C) using live, moribund, and dead individuals. As expected, most of the non-biopsied dead or moribund individuals were mainly from seawater (88%) and the pre-smolt and de-smolt trials (98%) over the six days of treatment exposure. Similarly, we identified biomarkers that were associated with mortality across the 18 groups (live vs. dead or moribund). Here, we describe the changes in other physiological variables, body size, skin pigmentation, and body morphology associated with smolt status, as well as the salinity, temperature, and dissolved oxygen treatments.

### 4.1. Smolt status effects

Juveniles placed into seawater outside of the smoltification physiological window (pre-smolt or de-smolt) can experience high mortality (Björnsson et al., 2011; McCormick et al., 1998; McCormick et al., 2013). As expected, the four trials (March to August) differed in the mortality of non-biopsied juveniles, such that juveniles that were not optimally smolted (pre-smolt or de-smolt) experienced higher mortality in seawater than smolt. Consequently, a physiological condition (because of life stage) affected the severity of a common stressor (salinity), with mortality indicating a severely maladaptive or negative consequence (Schreck and Tort, 2016). These results support other studies (e.g. Stich et al., 2015; Stich et al., 2016), that show the degree of smoltification matters for survival in seawater, and likely early marine survival.

As expected for candidate salinity genes initially developed for measuring the degree of smoltification (see Houde et al., 2018), the gene expression patterns using the salinity biomarkers were affected by smolt status (PC1) and salinity groups (PC2). That is, some of these smoltification genes did respond directly to salinity. Also, the patterns for temperature and dissolved oxygen biomarkers did not have as clear a relationship with smolt status, nor was one expected. Salinity (PC1) displayed a pattern of smolt at one extreme, pre-smolt and de-smolt in between, with mortality at the other extreme. We describe the gene expression pattern in the direction of mortality, as well as the pattern for the salinity groups below.

Typically during smoltification, the body silvers, caudal fin margins darken, and the caudal peduncle elongates, morphological changes that possibly help with camouflage and swim performance in marine habitats (Björnsson et al., 2011; McCormick et al., 1998). Correspondingly, the smolts had brighter anterior and posterior regions, darker caudal fins, and longer caudal peduncles than pre-smolts, but the anterior region brightness decreased in de-smolts. Thus, increased caudal fin darkness and peduncle length were common smoltification changes in the present and previous study (Houde et al., 2018), but changes in body brightness were not detected in the previous study.

Although we detected significant differences in NKA activity in freshwater between pre-smolt and smolts, as may be expected for different degrees of smoltification (e.g. Kiilerich et al., 2007; Lemmetyinen et al., 2013; Piironen et al., 2013), we did not detect similar differences between smolts and de-smolts. However, we can classify these freshwater individuals as pre-smolt, smolt, and de-smolt using the same biomarkers that were examined for salinity treatment (Houde et al., 2018). Our data suggest that gene expression may be a more sensitive indicator of smoltification or seawater preparedness than freshwater NKA activity. Similarly, high NKA activity in freshwater prior to seawater entry may not be necessary if juveniles can rapidly increase NKA activity in seawater during the physiological window (Bassett et al., 2018; Madsen and Naamansen, 1989). After seawater transfer, the NKA activity over six days was the highest in smolts relative to pre-smolt and de-smolt. These results suggest that smolts may also be able to achieve higher NKA activity in seawater than pre-smolts and de-smolts.

### 4.2. Salinity biomarkers

There were 11 salinity biomarkers, with gene expression pattern (PC2) associated with separation among the salinity groups. Supporting that PC2 was associated with salinity groups were the positive correlations with plasma chloride concentrations and NKA activity, another physiological biomarker of salinity acclimation (Björnsson et al., 2011; McCormick et al., 1998).

In seawater, fish excrete excess ions primarily via the gills (Evans et al., 2005; Hwang and Lee, 2007). Two of 11 biomarkers were ion regulation genes (i.e. CA4 and CFTR-I) that were tightly linked to the positive end of PC2. Similarly, others have found a higher expression of ion regulation genes with transfer from freshwater to higher salinity (Flores and Shrimpton, 2012; Havird et al., 2013; Singer et al., 2002). Although the third ion regulation gene (i.e. NKAa1-b) is highly supported for its association with smoltification and salinity transfer (e.g. Björnsson et al., 2011; Nilsen et al., 2007), this gene may not be a good biomarker because it was not specific to salinity. NKAa1-b and NKA activity were secondarily influenced by temperature, as seen previously (Bassett et al., 2018). On the other end of PC2, three genes were related to immunity (i.e. CCL4, CCL19, and IFI44). Immunity genes are generally suppressed during seawater transfer because of a proposed energetic trade-off between immunity and acclimation to seawater (Johansson et al., 2016; Makrinos and Bowden, 2016). Interestingly, IFI44 is a validated biomarker for the detections of viral disease state (Miller et al., 2017a). Collectively, specific ion regulation and immunity genes were linked to a change in salinity, with a strong ability to distinguish fish in freshwater from those in either seawater or brackish.

### 4.3. Mortality in seawater biomarkers

Juveniles can die in seawater from internal ionic and osmotic disturbances, which can be detected from higher plasma ion concentrations (Blackburn and Clarke, 1987) and associated with lower gill NKA activity (Kennedy et al., 2007; Stich et al., 2015; Stich et al., 2016). Indeed, juveniles that died or became moribund shortly after seawater exposure had higher plasma chloride concentrations, lower NKA activity, smaller body size (length, mass, and elongation), and lower condition (including thickness) than juveniles that lived. Handeland et al. (1996) earlier observed shorter predator escape distance and higher piscine predation during seawater transfer, which may correspond with the lower startle response of live juveniles in seawater than freshwater and brackish. Although there was higher NKA activity and plasma chloride concentrations than freshwater, there may have been little mortality and body changes in brackish (20 PSU) because it is closer to internal isoosmotic salinity with less energy used for homeostasis than seawater (Morgan and Iwama, 1991; Stien et al., 2013; Webster and Dill, 2006).

Mortality was associated with gene expression pattern (PC1) of the 11 salinity biomarkers. Dead or moribund juveniles also had higher expression of two freshwater ion regulation genes (i.e. NKAa1-a and PRLR) and lower expression of a seawater ion regulation gene (i.e. CFTR-I) than live juveniles. These results further emphasize that mortality in seawater is associated with a mismatch of ion regulation gene expression causing internal ionic and osmotic disturbances after transfer from freshwater.

### 4.4. Temperature biomarkers

Our results showed that five of the 10 upregulated genes at 18°C, including two paralogs of SERPINH and two paralogs of hsp90a and hsp70, were heat shock proteins. These genes are known to be the most frequent heat shock proteins (HSPs) in response to higher temperature across salmonids and other species studied (Akbarzadeh et al., 2018a). The importance of molecular chaperoning of macromolecules during higher temperature has been well established. In particular, the heat shock response (HSR), defined as the ability of the HSPs to re-fold thermally damaged proteins and prevent their cytotoxic aggregation, is a well-described phenomenon across taxa (Logan and Buckley, 2015). The upregulation of many paralogs relating to HSP genes including SERPIN, HSP70, and HSP90a were also observed in adult Sockeye salmon in response to chronic elevated temperature (Akbarzadeh et al., 2018a; Jeffries et al., 2012; Jeffries et al., 2014b). Our literature review revealed that the above HSP genes are upregulated in more than 16 different fish species belonging to different taxa, including salmonids in response to high temperature (Akbarzadeh et al., 2018a). Therefore, HSP genes seem to be robust temperature biomarkers across all salmonids and likely many fish species.

Two paralogs of FKBP10 were significantly downregulated at the highest temperature compared to 10 and 14°C. FK506-binding protein 10 is localized in the lumen of the endoplasmic reticulum (ER), and it encompasses four peptidylprolyl isomerase domains (PPIase) which accelerate protein folding by catalyzing the cis-trans isomerization of proline imidic peptide bonds in oligopeptides (Silvestre et al., 2010). Downregulation of FKBP10 in fish under higher temperature may be related to the remarkable increase of SERPIN expression. Although FKBP10 has been proposed to be a collagen molecular chaperone responsible for the stabilization of collagen triple helices cooperatively with SERPIN, this chaperone function in the ER may actually be predominantly fulfilled by SERPIN (Ishikawa et al., 2017). Under certain conditions like higher temperature or loss of Hsp47, FKBP10 may provide a functional redundancy for molecular chaperone activity (Ishikawa et al., 2017). Downregulation of FKBP10 was also observed in adult Sockeye salmon (Akbarzadeh et al., 2018a; Jeffries et al., 2014b), catfish (Liu et al., 2013) and white sturgeon (Silvestre et al., 2010) in response to high temperature. Differential expression of FKBP10 gene seems to be specific for temperature change, hence could be a strong contributor towards a biomarker panel to predict chronic exposure to higher temperature in fish.

Genes involved in protein biosynthesis including EEF2 and SFRS2 were also among the significant downregulated genes in fish held in higher temperature. Eukaryotic elongation factor 2 (EEF2, assays ef2_14 and ef2_3), is a member of the GTPase superfamily of proteins, catalyzes the movement of the ribosome along the mRNA and is essential for translocation of pep-tidy1 tRNA from the A to P site of the ribosome (Shastry et al., 2001). Serine/arginine-rich splicing factor 2 (SFRS2), which is a specific well known serine/arginine-rich (SR) protein family member, is a mediator of genome stability, pre-mRNA splicing, mRNA nuclear export, and translational control (Wang et al., 2016). Downregulation of both EEF2 and SFRS2 suggests that a decrease in protein biosynthesis may be an energy saving mechanism at the cellular level in response to higher temperature (Jeffries et al., 2014b). Indeed, exposure to high temperatures can inhibit transcription and protein synthesis, probably reflecting the suppression of noncritical activities (Buckley et al., 2006; Kassahn et al., 2007). Previous studies have shown that exposure to chronic elevated water temperatures decreases the expression of EEF2 and SFRS2 genes in adult Pacific salmon (Akbarzadeh et al., 2018a; Jeffries et al., 2012; Jeffries et al., 2014b).

Changes in behavioural and physiological variables indicated higher metabolism and energy utilization during higher temperature. As expected (Heath, 1973; Zhao et al., 2017), gill ventilation rate at 14°C and 18°C was higher compared to that at 10°C. Also, plasma lactate concentrations increased with higher temperatures, as seen earlier with adult Sockeye salmon (Jeffries et al., 2012; Steinhausen et al., 2008), which is indicative of the need for anaerobic metabolism (Han et al., 2017; Pankhurst, 2011). Furthermore, the juveniles had higher NKA activity in warmer than colder water. NKA activity is an energy consuming cell membrane ion pump (Mobasheri et al., 2000). Higher NKA activity may be a mechanism to cope with these increased fluxes for salmon in warmer temperatures (Handeland et al., 2000) due to higher cell membrane permeability (Vernberg and Silverton, 1979). Moreover, plasma glucose concentrations decreased with higher temperatures perhaps because hepatic glycogen stores were being depleted with temperature (Chadwick and McCormick, 2017). Overall, these results of increased metabolism and energy utilization may also explain the smaller body size (length, mass, and elongation) and lower condition (including thickness) for juveniles kept in warmer temperatures. Similar changes in growth in response to higher temperature have been observed for juvenile Chinook salmon (Marine and Cech, 2004) and Artic charr (Lyytikainen et al., 2002).

### 4.5. Dissolved oxygen biomarkers

The 11 dissolved oxygen (and temperature) biomarkers identified by gene shaving could not strongly separate normoxia and hypoxia. This may not be surprising given that the background candidate dissolved oxygen gene expression information was limited in fish, especially salmonid species and gill tissue. Currently, using the freshwater samples of the present study, we are undertaking an RNA-seq study to discover additional candidate dissolved oxygen genes, which will then be validated with other samples (Akbarzadeh et al., 2018b In Prep). Regardless, two genes (HIF1A_6 and COX6B1) were primarily influenced by dissolved oxygen using model selection. We discuss the hypoxia-related reasoning behind these two genes.

Exposure to hypoxia causes upregulation of HIF-α mRNA expression in tetrapods and teleost fishes (Rahman and Thomas, 2017). HIF1-A is suggested as a reliable fish biomarker of hypoxia exposure (Wenger, 2002). The upregulation of HIF1-A in response to hypoxia has been also observed in Atlantic croaker (Rahman and Thomas, 2017), goby (Gracey et al., 2001), ruffe and flounder (Tiedke et al., 2014) and Pacific herring (Froehlich et al., 2015). COX6B1 is a non-transmembrane subunit of COX, which could be upregulated in response to the higher oxidation during hypoxia. It is involved in oxidation-reduction process, proton transport, and ion transmembrane transport (Long et al., 2015). The upregulation of COX6B1 has been observed in both hypoxia (Long et al., 2015) and higher temperature (Garvin et al., 2015; Jeffries et al., 2012; Jeffries et al., 2014b).

Hypoxia alone did not influence juvenile mortality except during handling; however, juveniles showed adaptive responses that modulated their behavioural and physical phenotype. Gill biopsied mortality was higher for hypoxia than normoxia in one of the de-smolt trials but not the second de-smolt trial. It is possible that some gill filaments were damaged during juvenile handling, providing less efficient oxygen exchange in hypoxia eventually causing mortality. Regarding behaviour, juveniles showed higher ventilation rate for hypoxia compared to normoxia. One of the most evident physiological adjustments to hypoxia is increased ventilation in an effort to compensate for lower dissolved oxygen (Itazawa and Takeda, 1978; Steffensen et al., 1982). Juveniles in hypoxia also showed paler skin pigmentation compared to normoxia. This result may be associated with the effects of skin pigmentation controlling hormones including α-melanophore stimulating hormone (αMSH) and melanin concentrating hormone (MCH). These hormones are pleiotropic by not only controlling skin pigmentation but also regulating the response to other stressors (Burton and Vokey, 2000).

### 4.6. General mortality biomarkers

Across the 18 groups, distress (dead or moribund) non-biopsied and biopsied juveniles showed an upregulation of seven genes associated with the physiological stress response. Changes in heat shock proteins (e.g. hsp70_6 and hsp90a_15 genes), metabolite (e.g. fructose-bisphosphate aldolase, ALD_1 and lactate, LDH_1 genes), and immune function (e.g. FKBP5, RGS21, VEGFa_1 genes) are secondary in the stress response of fishes, after the primary release of stress hormones (Barton, 2002). Plasma lactate concentrations were also higher for distressed than live juveniles in seawater. The increase in heat shock response is interesting given its prevention of apoptosis (Beere, 2005). One potential explanation is this response may not completely protect organisms from mortality. When the ability of protective functions fails, organisms may face drastic oxidative or necrotic tissue damage and finally enter in a death phase (Dutta et al., 2018).

Tertiary effects of stress can be the enhancement of disease susceptibility through the breakdown of immune barriers of defense (Aich et al., 2009; Marsland et al., 2002). The fish were screened for diseases by DFO prior to the commencement of our study. Although the fish passed the screening, this may not necessarily mean that the fish were not carrying agents that can cause disease. We examined the presence and loads of 47 salmon infectious agents in 79 distressed juveniles, and detected only two bacteria (i.e. *Candidatus Branchiomonas cysticola* and *Flavobacterium psychrophilum*) carried at elevated loads (within the range that can be associated with disease) in only a few fish. Both bacteria are common in juvenile Chinook salmon, and their presence alone is not necessarily indicative of a disease state (Bass et al., 2017; Miller et al., 2017b; Tucker et al., 2018). If an outbreak of disease by either bacterium had occured, and contributed to mortality, we would have expected a general elevation of the bacterium in most or all dying fish in the tank; instead, we observed only sporadic fish with elevated levels of either bacterium amongst the tanks. These data suggest that there were no outbreaks of stress-induced disease during the six days for the trials, and instead the distressed juveniles of our study were the result of stress from seawater and gill biopsy.

### 4.7. Conclusion

We identified salmonid gill gene expression biomarkers specific to salinity and temperature treatments across multi-stressor conditions and smolt statuses using a sophisticated experimental set-up. Salinity biomarkers were associated with ion regulation and immunity functions. Temperature biomarkers were associated with heat shock proteins and protein biosynthesis functions. Similar biomarkers were not identified for dissolved oxygen. However, we are discovering and validating additional candidate dissolved oxygen biomarkers using RNA-seq on samples from the present study (Akbarzadeh et al., 2018b In prep). We also identified biomarkers associated with general mortality, with links to secondary protein products of the fish stress response. The changes in behaviour, plasma variables, NKA activity, body size, body morphology, and skin pigmentation in response to the three treatments, as well as mortality and smolt status are also described. Most of the mortality was in seawater, where juveniles not optimally smolted experienced osmotic and ionic disturbances, e.g. lower condition from dehydration and higher plasma ions. We highlight that moribund or dead juveniles may have a mismatch in ion regulation gene expression patterns expected for seawater acclimation. The general and seawater mortality biomarkers are useful for gauging general physiological state and stress response. Importantly, these salinity, temperature, and eventually dissolved oxygen biomarkers can be used in natural environments to identify a specific stressor even under multi-stressor conditions.

## Supporting information

## Acknowledgements

This research was supported by the Natural Sciences and Engineering Research Council of Canada through a Postdoctoral Fellowship and by Mitacs/Pacific Salmon Foundation through an Accelerate Internship to ASH. Funding for the research was provided by Genome British Columbia, the Pacific Salmon Commission, and Fisheries and Oceans Canada (DFO) Genomic Research and Development Fund to KMM. APF holds a Canada Research Chair. We thank DFO Aquarium services for help in the design and assembly of the experiment. We also thank E. Di Cicco, C. Rycroft, B. Sutherland, N. Ginther, K. Mohns, D. Johnson, A. Yao, K. Halvorson, B. White, C. Lefebvre, R. Shearer, D. Moulton, L. Elmer, S. Esenkulova, R. Greiter, A. Duguid, A. Burton, X. He, J. Campbell, A. McMillan, C. Webb, for help with sample collection.

## Supplementary Tables

**Table S1:**
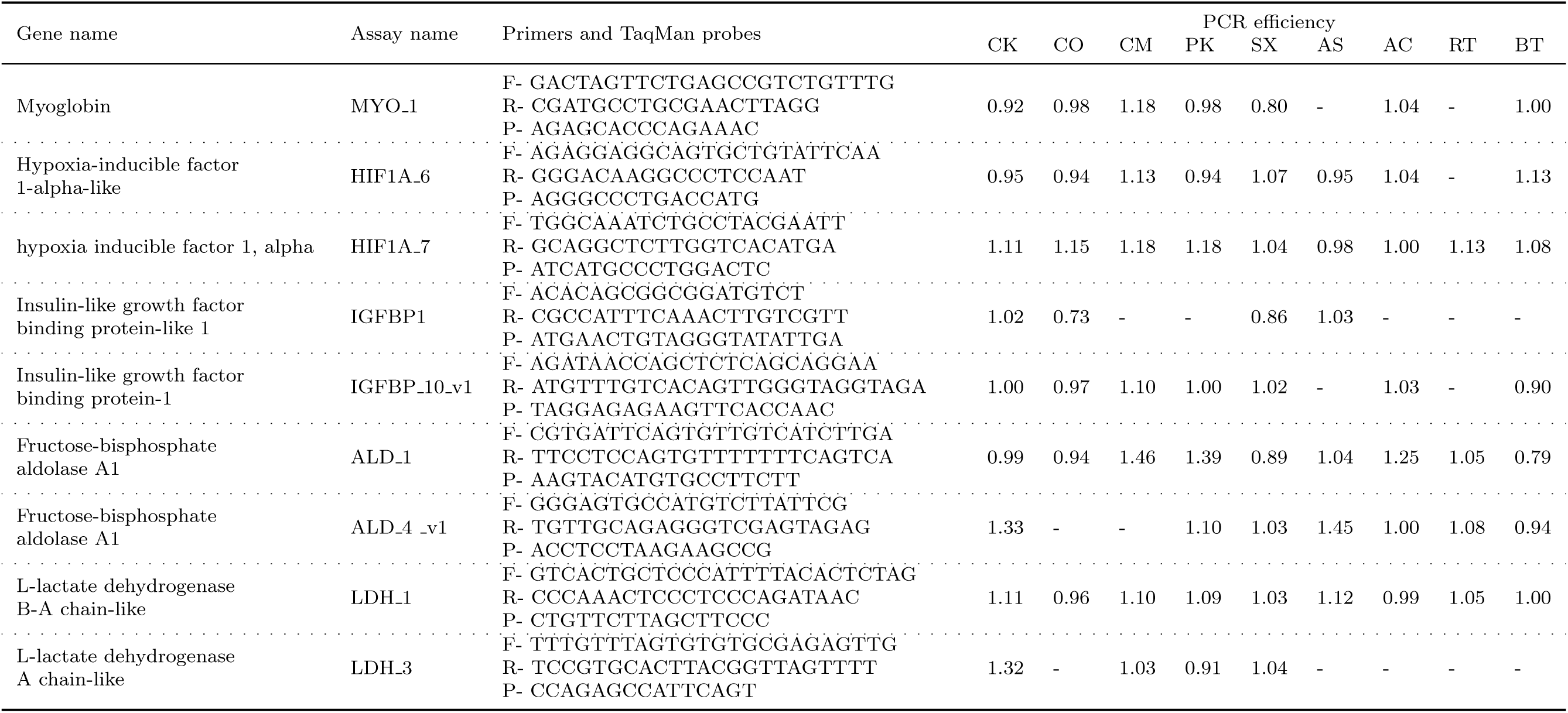
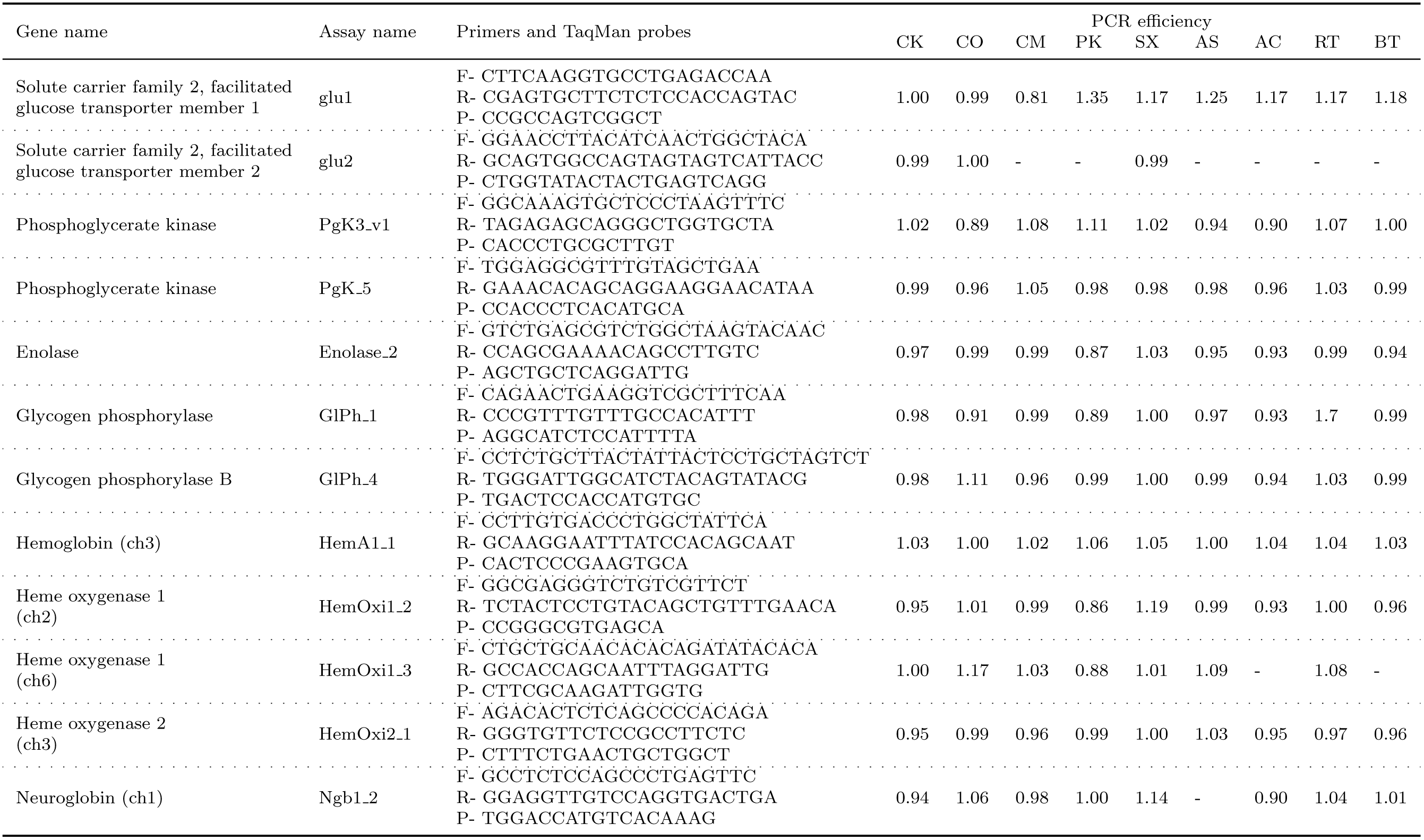
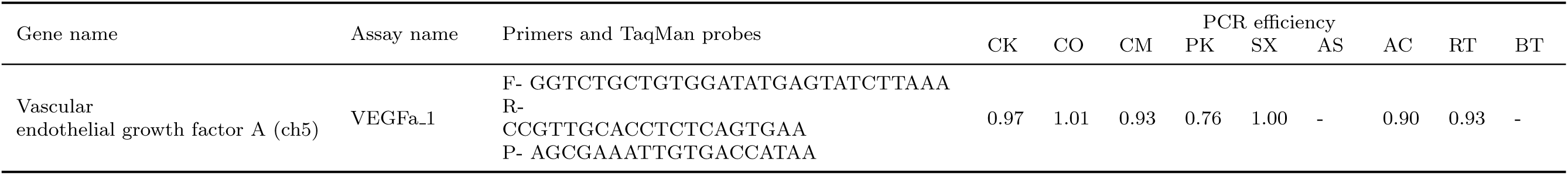
Summary of qPCR TaqMan assay designs and efficiencies for candidate dissolved oxygen genes. Presented are the forward (F), reverse (R), and probe sequences (P). Species abbreviations for efficieny: CK= Chinook salmon *(Oncorhynchus tshawytscha*), CO= Coho salmon (*O. kisutch)*, CM= Chum salmon (*O. keta*), PK= Pink salmon (*O. gorbuscha*), SX= Sockeye salmon (*O. nerka*), AS= Atlantic salmon *(Salmo salar*), AC= Artic charr *(Salvelinus alpinus*), RT= Rainbow trout (*O. mykiss*), and BT= Bull trout *(Salvelinus confluentus*).

**Table S2:** Correlations between gene expression patterns and physiological, body size, skin pigmentation, and body morphology variables. Displayed are the variable Pearson correlations (and p-value) with the genes expression patterns (PC1 and PC2) for salinity, temperature, dissolved oxygen, and mortality biomarkers. Mortality correlations coded live individuals as 0 and moribund or dead individuals as 1. Relative startle response correlations are at the level of the 18 groups, which coded freshwater as 3, brackish as 2, and seawater as 1. Gill ventilation correlations are also at the level of the groups; gill ventilation values were corrected for trial differences (see Appendix 3), and are the mean for trials 1, 3, and 4 combined; there was no video to analyze for trial 2.

| variable | salinity |  | temperature |  | dissolved oxygen |  | mortality |  |
| --- | --- | --- | --- | --- | --- | --- | --- | --- |
|  | PC1 | PC2 | PC1 | PC2 | PC1 | PC2 | PC1 | PC2 |
| *Physiological* |  |  |  |  |  |  |  |  |
| mortality | 0.63 (<0.001) | 0.18 (<0.001) | -0.19 (<0.001) | -0.66 (<0.001) | 0.80 (<0.001) | -0.05 (0.272) | 0.90 (<0.001) | -0.00 (0.982) |
| relative startle response | 0.23 (0.360) | -0.89 (<0.001) | 0.11 (0.658) | 0.48 (0.044) | -0.68 (0.002) | 0.01 (0.979) | -0.58 (0.012) | -0.45 (0.059) |
| gill ventilation rate | -0.11 (0.659) | 0.03 (0.913) | 0.32 (0.200) | -0.28 (0.262) | 0.39 (0.113) | -0.15 (0.547) | 0.21 (0.412) | -0.02 (0.953) |
| Na<sup>+</sup>/K<sup>+</sup>-ATPase activity | -0.12 (0.010) | 0.36 (<0.001) | 0.10 (0.045) | 0.02 (0.737) | -0.10 (0.049) | -0.18 (<0.001) | -0.10 (0.036) | -0.09 (0.050) |
| lactate concentrations | 0.34 (<0.001) | 0.02 (0.828) | 0.11 (0.115) | -0.43 (<0.001) | 0.39 (<0.001) | -0.28 (<0.001) | 0.49 (<0.001) | -0.15 (0.034) |
| glucose concentrations | 0.01 (0.941) | -0.02 (0.772) | -0.12 (0.091) | 0.23 (0.001) | -0.33 (<0.001) | 0.01 (0.909) | -0.29 (<0.001) | -0.24 (0.001) |
| chloride concentrations | 0.11 (0.367) | 0.37 (0.001) | 0.05 (0.685) | -0.55 (<0.001) | 0.48 (<0.001) | -0.30 (0.011) | 0.42 (<0.001) | 0.07 (0.571) |
| *Body size* |  |  |  |  |  |  |  |  |
| length | 0.19 (<0.001) | 0.04 (0.419) | -0.03 (0.508) | 0.09 (0.058) | -0.32 (<0.001) | -0.47 (<0.001) | -0.19 (<0.001) | -0.61 (<0.001) |
| mass | 0.21 (<0.001) | 0.00 (0.938) | -0.05 (0.280) | 0.08 (0.078) | -0.30 (<0.001) | -0.42 (<0.001) | -0.18 (<0.001) | -0.57 (<0.001) |
| condition | -0.33 (<0.001) | -0.10 (0.032) | -0.05 (0.334) | 0.40 (<0.001) | -0.42 (<0.001) | 0.24 (<0.001) | -0.46 (<0.001) | 0.12 (0.012) |
| *Skin pigmentation* |  |  |  |  |  |  |  |  |
| anterior brightness | 0.02 (0.710) | -0.09 (0.083) | -0.01 (0.812) | 0.00 (0.985) | 0.06 (0.195) | 0.09 (0.058) | 0.06 (0.195) | 0.05 (0.307) |
| caudal fin darkness | 0.10 (0.050) | 0.00 (0.929) | -0.05 (0.332) | -0.09 (0.063) | 0.07 (0.167) | -0.10 (0.046) | 0.10 (0.044) | -0.11 (0.032) |
| posterior brightness | 0.12 (0.017) | 0.01 (0.900) | 0.03 (0.524) | 0.02 (0.718) | -0.12 (0.020) | -0.17 (<0.001) | -0.06 (0.248) | -0.36 (<0.001) |
| caudal fin yellowness | 0.01 (0.801) | -0.01 (0.906) | 0.06 (0.223) | -0.14 (0.004) | 0.31 (<0.001) | 0.28 (<0.001) | 0.24 (<0.001) | 0.23 (<0.001) |
| *Body morphology* |  |  |  |  |  |  |  |  |
| elongation | 0.06 (0.208) | 0.05 (0.323) | -0.07 (0.170) | 0.21 (<0.001) | -0.39 (<0.001) | -0.29 (<0.001) | -0.30 (<0.001) | -0.50 (<0.001) |
| back roundness | 0.22 (<0.001) | 0.00 (0.947) | 0.03 (0.566) | -0.36 (<0.001) | 0.38 (<0.001) | 0.02 (0.714) | 0.36 (<0.001) | 0.02 (0.653) |
| caudal peduncle length | -0.07 (0.150) | -0.06 (0.204) | 0.08 (0.104) | 0.00 (0.948) | 0.07 (0.174) | 0.08 (0.107) | 0.02 (0.636) | 0.14 (0.006) |
| thickness | -0.46 (<0.001) | -0.16 (0.001) | 0.04 (0.446) | 0.42 (<0.001) | -0.49 (<0.001) | 0.15 (0.003) | -0.54 (<0.001) | 0.10 (0.043) |

## Supplementary Appendices

*Available online at BioRxiv.

Appendix 1. Technical details on the experimental set-up of the challenge study manipulating salinity, temperature, and dissolved oxygen.

Appendix 2. Individual-based expression for 87 candidate genes after two and six days of treatment. Data are for live trial 1 (pre-smolt) and are mostly represented by individuals from the brackish treatments. Salinity treatment symbols are SW, BW, and FW for seawater, brackish, and freshwater groups. Temperature treatment symbols are 10, 14, and 18 for °C groups. Dissolved oxygen treatment symbols are N for normoxia and H for hypoxia groups.

Appendix 3. Treatment and mortality effects for physiological and body variables.

Appendix 4. Individual-based expression for 87 candidate genes after six days of treatment. Salinity treatment symbols are SW, BW, and FW for seawater, brackish, and freshwater groups. Temperature treatment symbols are 10, 14, and 18 for °C groups. Dissolved oxygen treatment symbols are N for normoxia and H for hypoxia groups.

Appendix 5. Classification ability of different combinations of the groups using the identified treatment biomarkers.

