## Supplementary material for "Salmonid gene expression biomarkers indicative of physiological responses to changes in salinity, temperature, but not dissolved oxygen"

### Low Oxygen Fish Holding System

*Technical Design and Drawings*

By ASP

August 30, 2018

Aquarium Services Program was tasked with designing and building a fish holding system for Chinook Salmon Fry as part of a smoltification experiment and gene expression. The three factor experiment (Salinity, Temperature, and Oxygen Levels), allows the fish to placed in 40 liter tanks supplied with freshwater at 100 % saturation at a standard temperature and over a few days transition to the required experiment matrix end points.

#### Terms

ppt: Parts per thousand

mg/l: Milligrams per Litre

Hypoxia: A low oxygen level (> 40 % saturation)

Brackish: A salinity midway between Freshwater (0 ppt) and Saltwater (28 ppt)

AUP: Animal Use Permit; Limits on the use of experimental animals as defined by the animal care committee

TGP: Total Gas Pressure; the amount dissolved gas in water, 100 % is the amount gas present when in equilibrium with the atmosphere.

#### Experimental Matrix

| Salinity | 10 C | | 14 C | | 18C | |
|---|---|---|---|---|---|---|
| 0 ppt | Normal | Hypoxia | Normal | Hypoxia | Normal | Hypoxia |
| 20 ppt | Normal | Hypoxia | Normal | Hypoxia | Normal | Hypoxia |
| 28 ppt | Normal | Hypoxia | Normal | Hypoxia | Normal | Hypoxia |

For each experimental treatment there was a control tank, and each control and treatment group was replicated; for a total of 36 tanks. See Figure 1 for the tank layout, three aluminum pot table tank tables were used for this trial. A pot tank is a fiberglass tank that holds 40 liters of water, water enters through a hole in the Plexiglas tank lid and drains out the bottom of tank via a central standpipe. (find a photo)

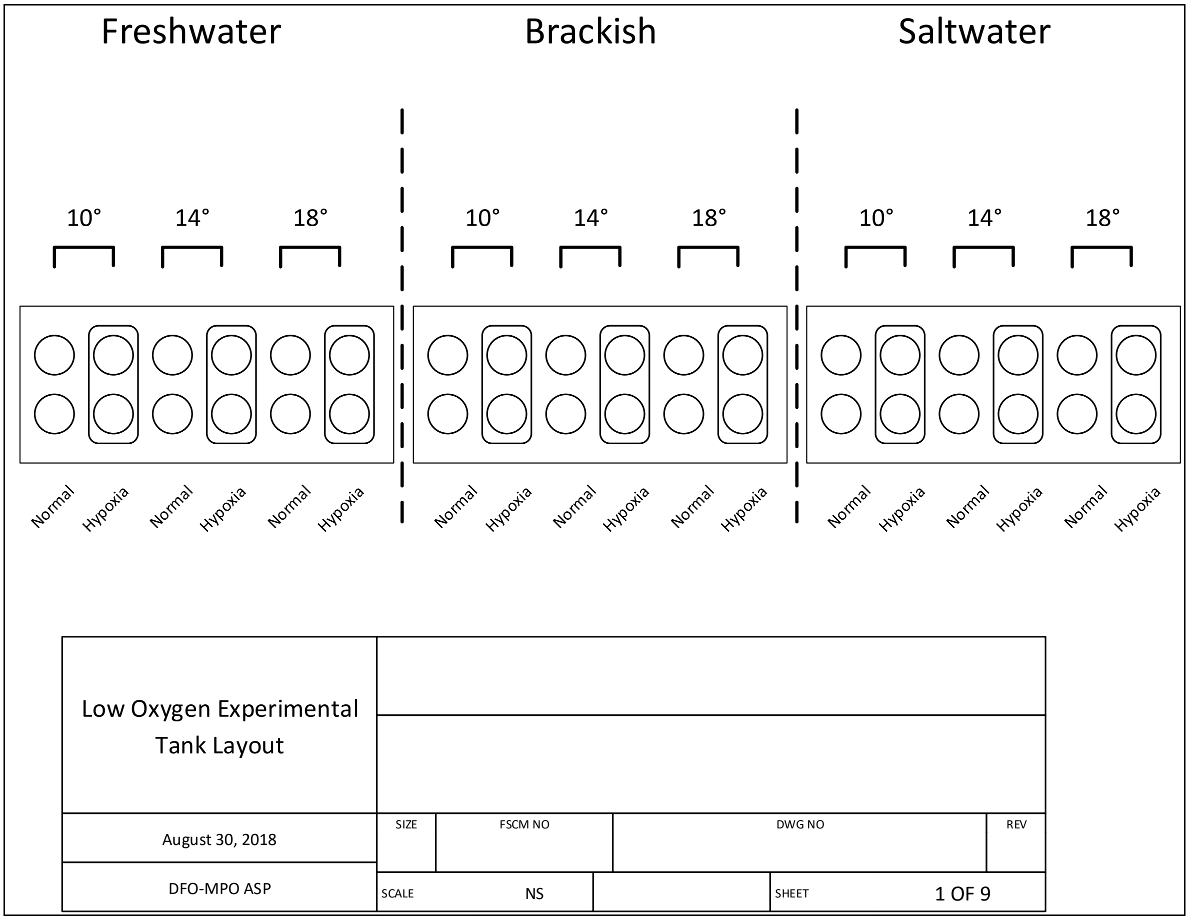


*Figure 1 Tank layout*

#### Salinity and Temperature Mixers

All water used in this experiment was supplied from the Brett Building supply headers. The Brett building can supply both fresh and saltwater at three water temperatures: chilled (3 C), ambient (10-14 C), and heated( 25 C).

Brackish water (20 ppt) was mixed first before it was adjusted to final water temperature, see figure 2 for the details.

#### Salinity Mixer

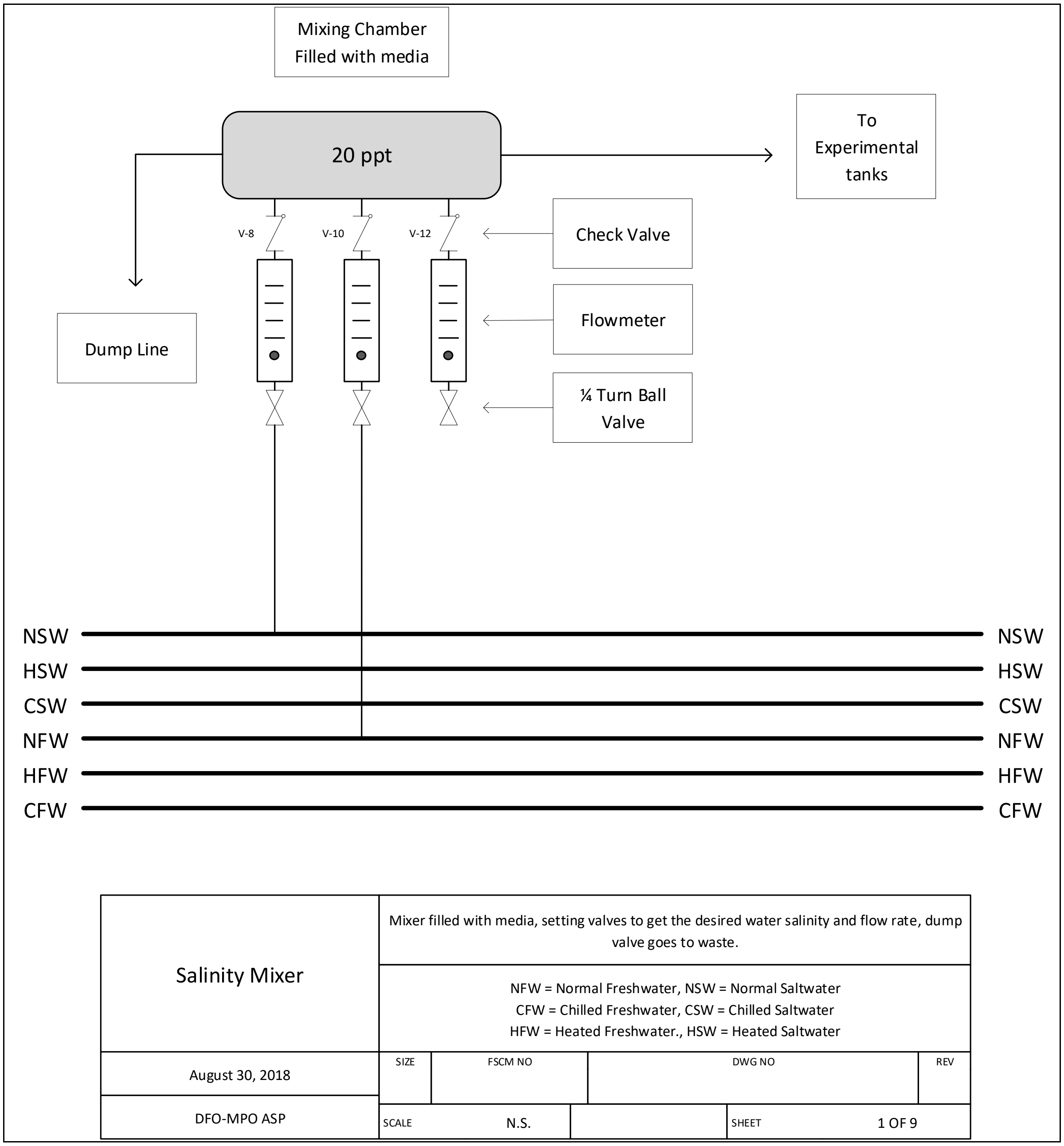


*Figure 2 Salinity Mixer*

By using the flowmeters and valves the salinity can be set, the dump valve allows the mixer to run at a optimum mix without restrictions of the temperature coil systems. Sch 80 PVC threaded pipe was used through, with vinyl tubing to allow flexible pipe routing. Most valves and flowmeters were  a mixture of ½ and ¾ inch fittings, the cooling coils were 3/8 inch ID Stainless Tubing.

#### Temperature Mixer

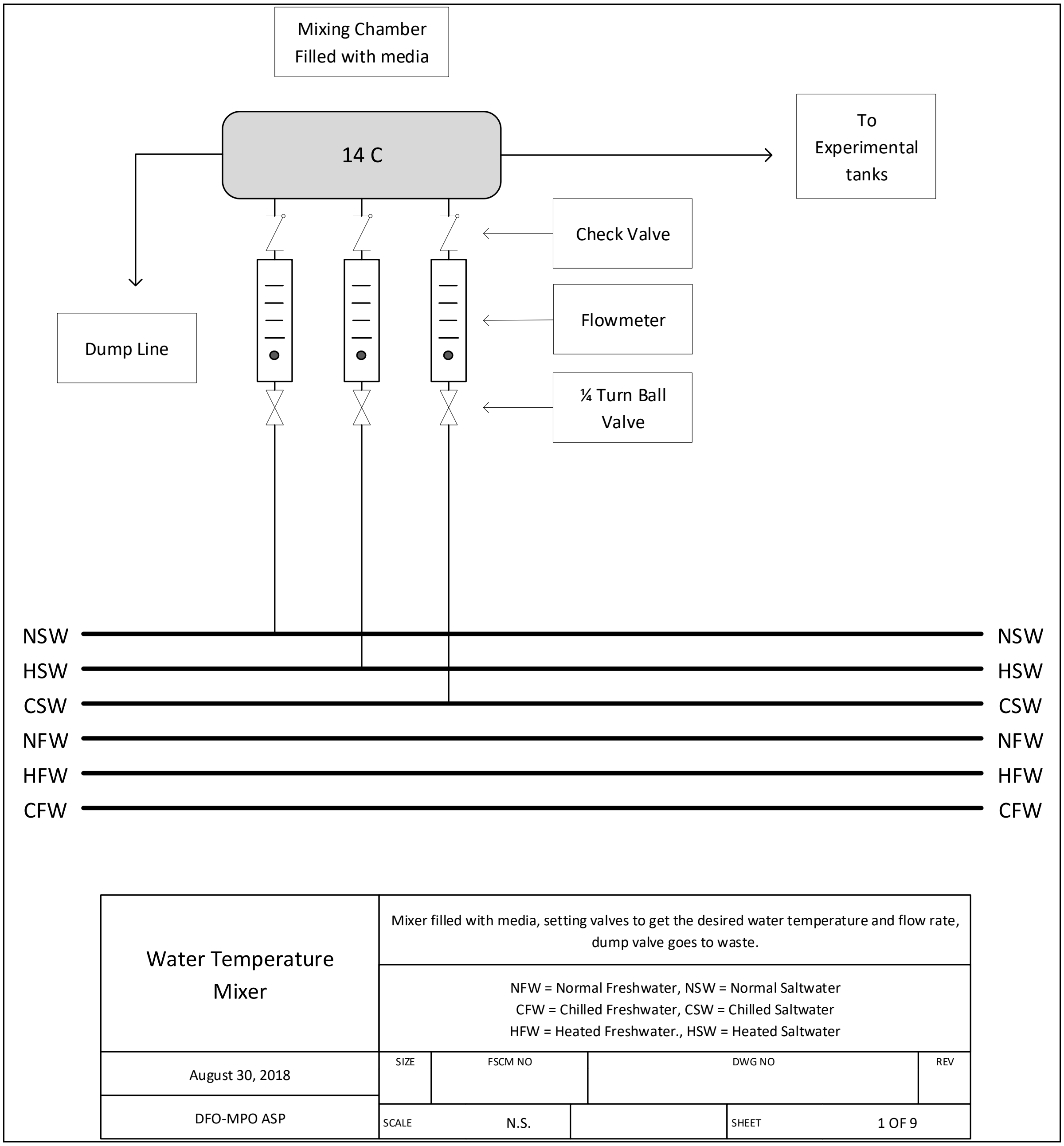


*Figure 3 Temperature Mixer*

Water temperature is controlled by adjusting the flow rate by noting the water flows and adjusting the valves.

The brackish temperature was more complex due to requirement change the water temperature without changing the salinity.

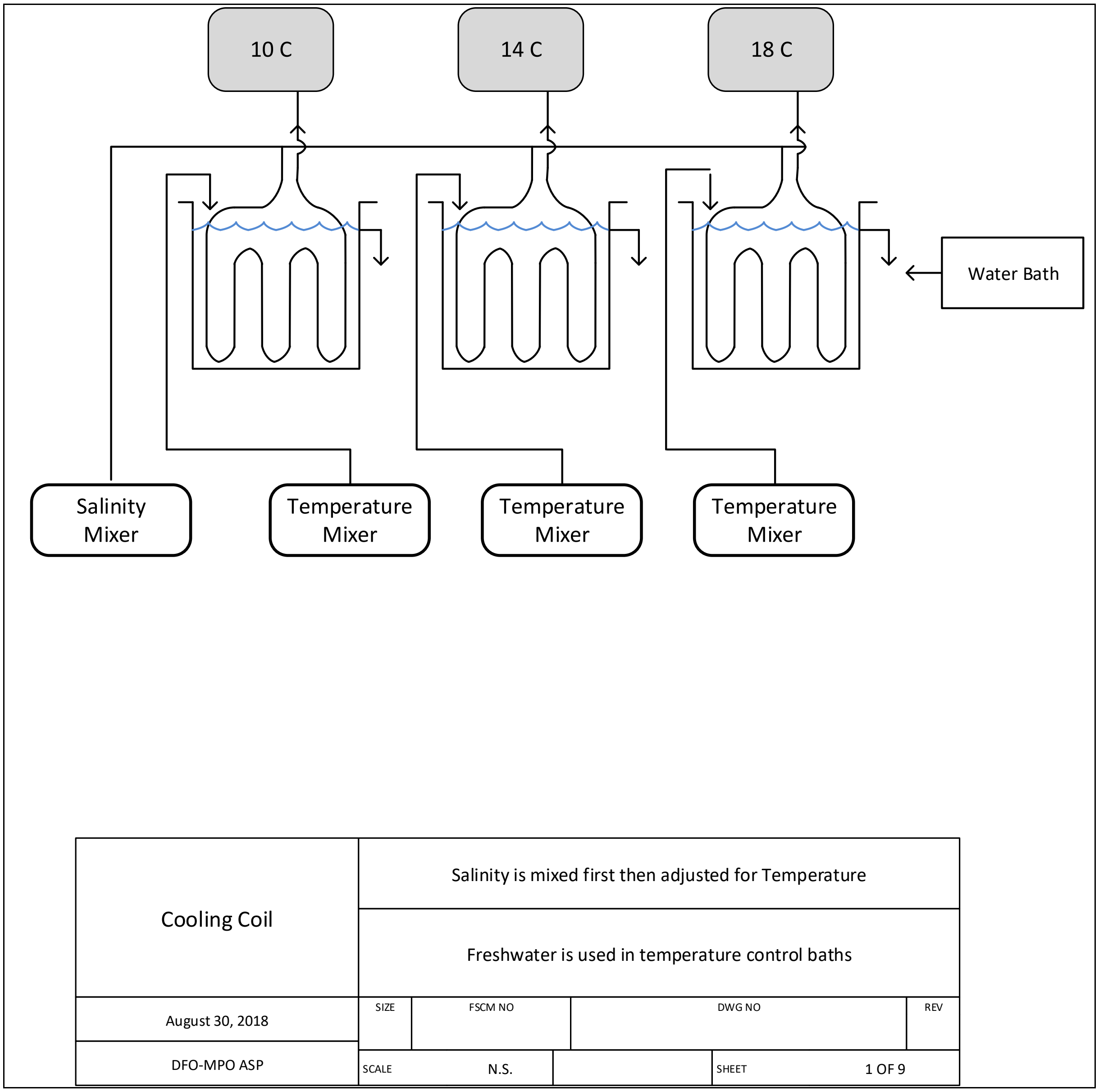


*Figure 4 Salinity Temperature adjustment using Cooling Coils*

Three different temperature mixers (similar to Figure 2) were used to control the water baths in which the coils for the salinity adjustment occurred.

#### Gas Level Control

Oxygen Levels in this experiment were at two levels; Normal and Hypoxic.

Normal is defined as a oxygen level equal 100 % saturation( a temperature and salinity neutral term), water is allowed to break up into a thin film as it pass over aeration media, this allow the water gas pressure to become equal to the atmosphere (Figure 5).

Hypoxic water was produced by passing water though a column using nitrogen to displace the dissolved oxygen. To control the level of stripping of oxygen, and dissolved oxygen meter and controller was to monitor the oxygen levels (Figure 6).

Nitrogen was supplied at first by high pressure K size bottles, and then later by Liquid Nitrogen tanks to a supply manifold with high precision flow meters. The high pressure bottles could be switch in the gas supply manifold when the liquid supply was being changed out. The nitrogen was injected in to the water column by ceramic airstone (Point 4 type), producing a bubble in the 5-10 micron size range. 3/8 inch braided PVC gas line with brass fittings (oxygen B-nut for connectors) was used to supply gas from the high pressure gas supply down to the airstones. Point 4 ceramic air stones were used through experimental setup, as they can produce 5-10 micron gas bubble.

The dissolved oxygen levels were controller by a Point 4 controller. The controller used a Oxyguard Oxygen Probe to monitor both oxygen and water temperature; this data was recorded by the controller every 5 minutes starting at the top of the hour. A gas solenoid in the nitrogen supply line controlled the tank oxygen levels. In this non standard control setup, the control logic as inverted to get required levels.

The controller was programed to present an alarm when the tank parameters (Temperature and oxygen levels) stepped out of bounds. The AUP limited how low the oxygen level could go (35% saturation) as not kill the fish, and the thermal limits.

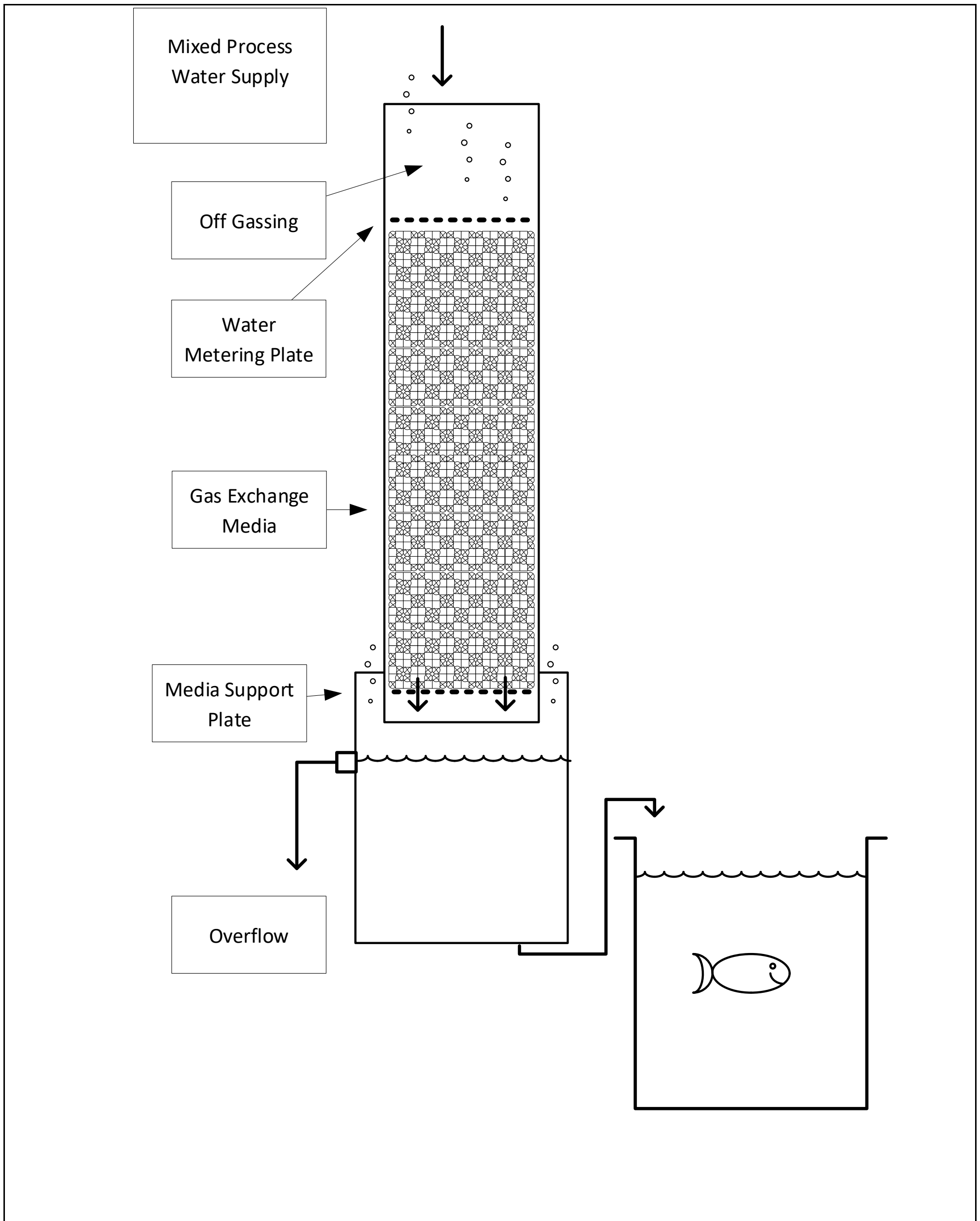


| | | | | | | |
|---|---|---|---|---|---|---|
| Aeration Column | Column Media ¾ inch to 1 ½ inch Biorings | | | | | |
| August 30, 2018 | SIZE | FSCM NO | | DWG NO | | REV |
| DFO-MPO ASP | SCALE | N.S. | | SHEET | 1 OF 9 | |

*Figure 5 Aeration Column*

Column made from 4 inch ABS pipe, use of Stainless Steel fasteners, and ¾ to 1 ½ inch PVC BioRings

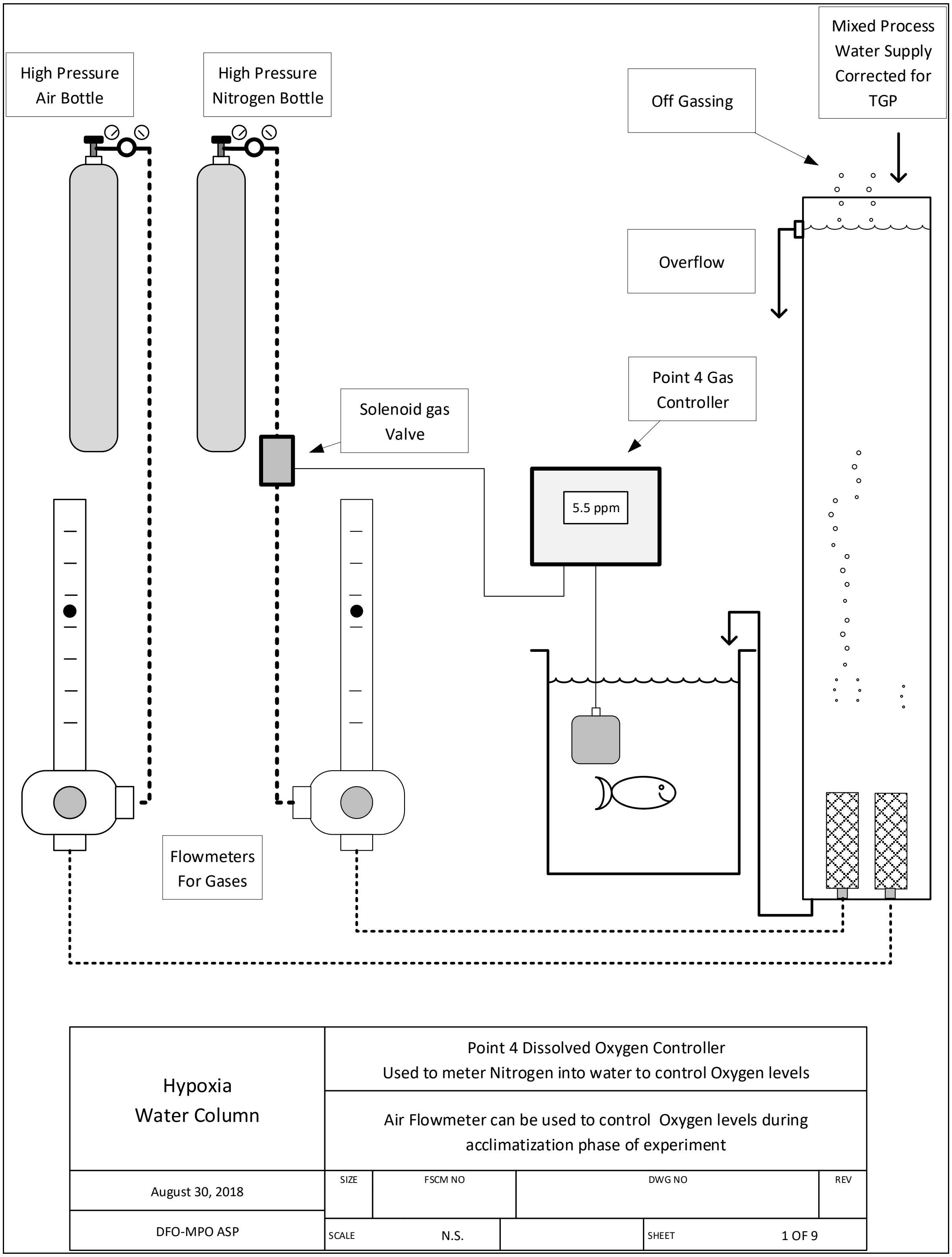


*Figure 6 Hypoxia Water Column*

The Point 4 Controller monitors the tank dissolved oxygen levels and used a solenoid valve on the nitrogen gas supply to control the gas level. Column make from 4 inch ABS pipe, used a ceramic airstone and braided PVC 3/8 inch airline.

#### Water Quality Meters

A number of water quality meters were used to check the output of this experimental system

- YSI salinity meter used to check water salinity and temperature
- Onset Tidbit v2 used to record the tank water temperatures (every 15 minutes start at top of the hour)
- Oxyguard DO meter, used check oxygen and temperatures (different models used)
- Point 4 TGP Meter, used to check TGP levels in the gas columns
