## Supplementary material for "Salmonid gene expression biomarkers indicative of physiological responses to changes in salinity, temperature, but not dissolved oxygen"

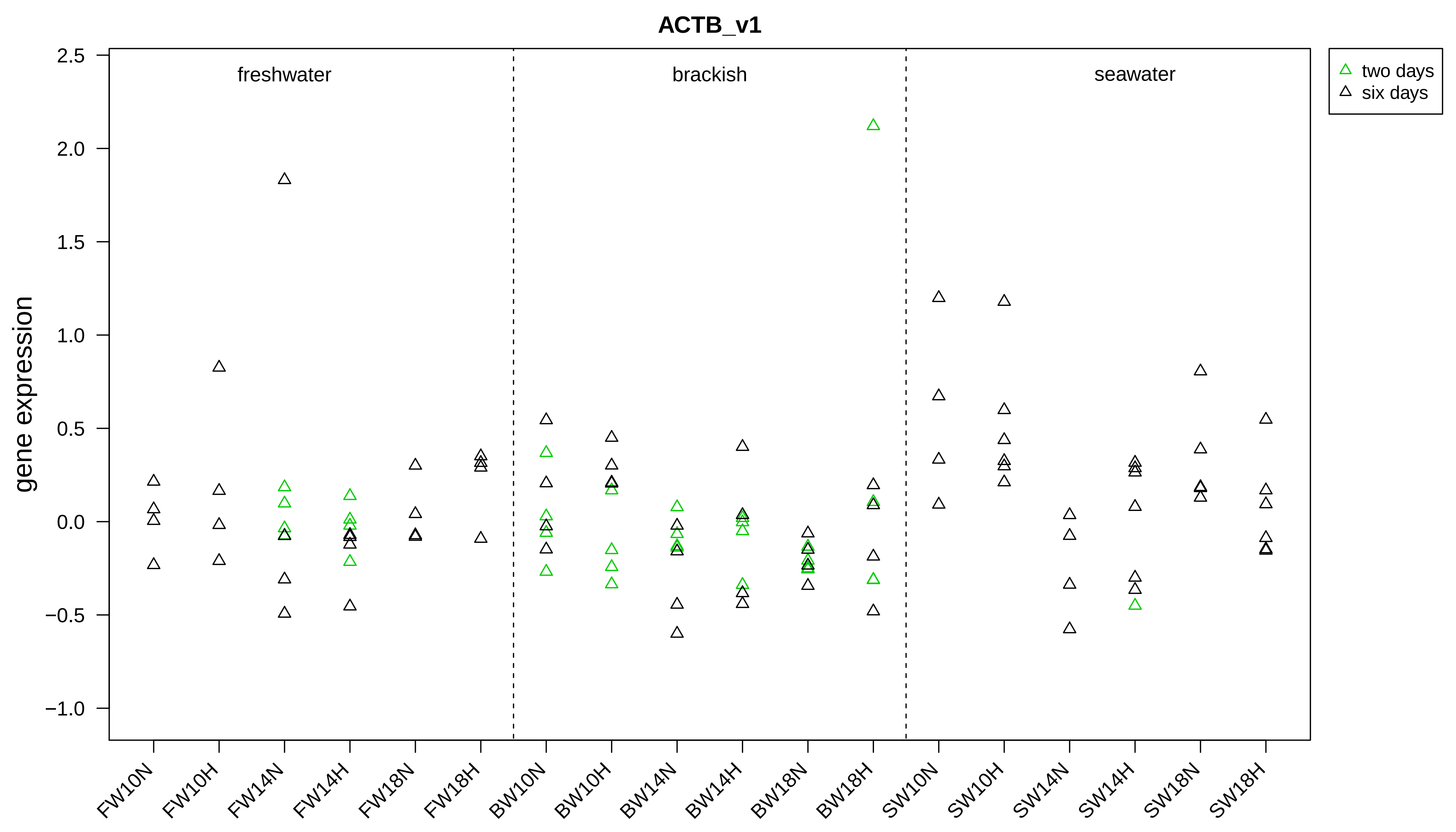
ACTB_v1
gene expression
2.5
2.0
1.5
1.0
0.5
0.0
−0.5
−1.0
freshwater
brackish
seawater
two days
six days
FW10N
FW10H
FW14N
FW14H
FW18N
FW18H
BW10N
BW10H
BW14N
BW14H
BW18N
BW18H
SW10N
SW10H
SW14N
SW14H
SW18N
SW18H

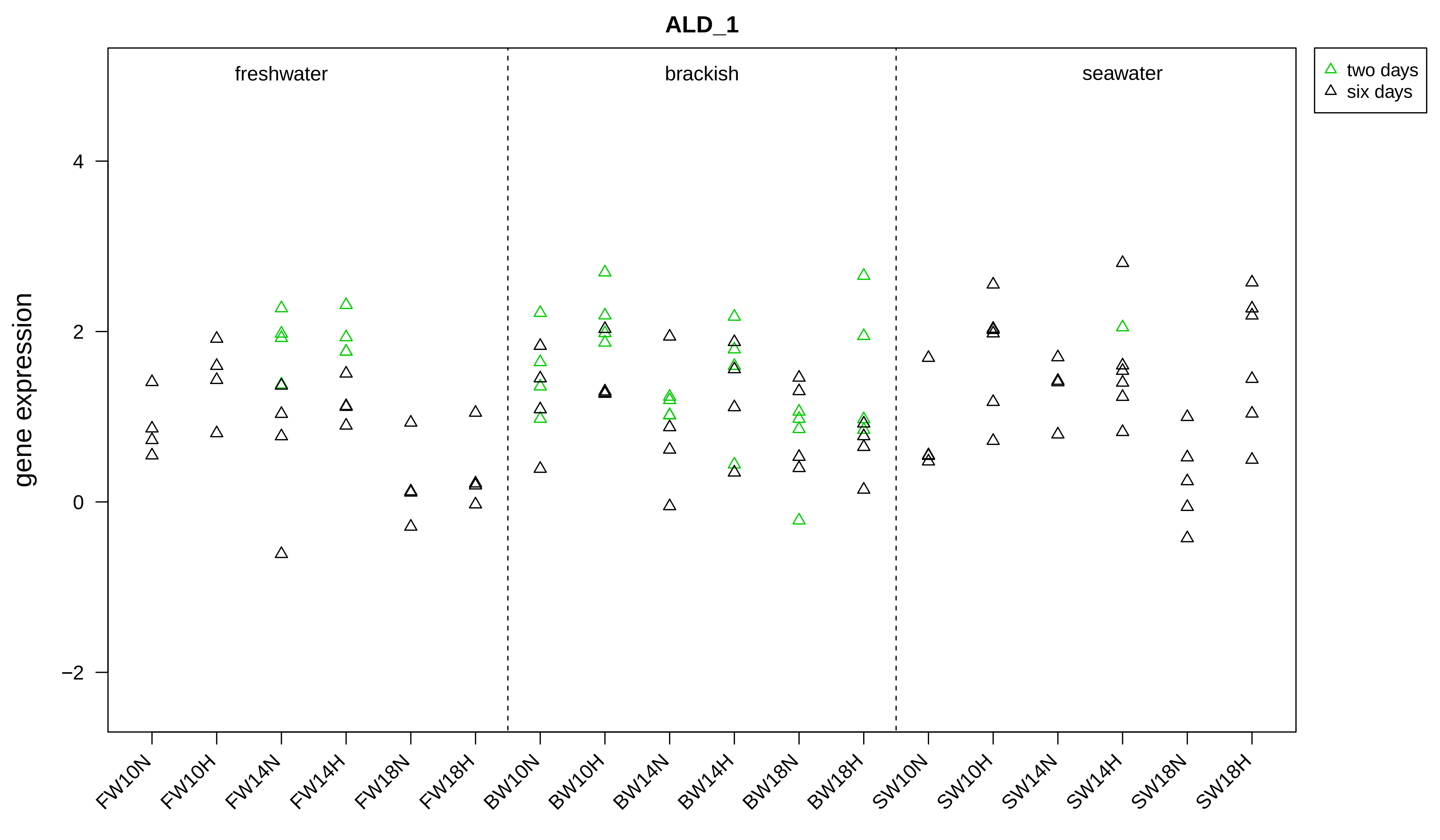
ALD_1
freshwater
brackish
seawater
two days
six days
gene expression
4
2
0
−2
FW10N
FW10H
FW14N
FW14H
FW18N
FW18H
BW10N
BW10H
BW14N
BW14H
BW18N
BW18H
SW10N
SW10H
SW14N
SW14H
SW18N
SW18H

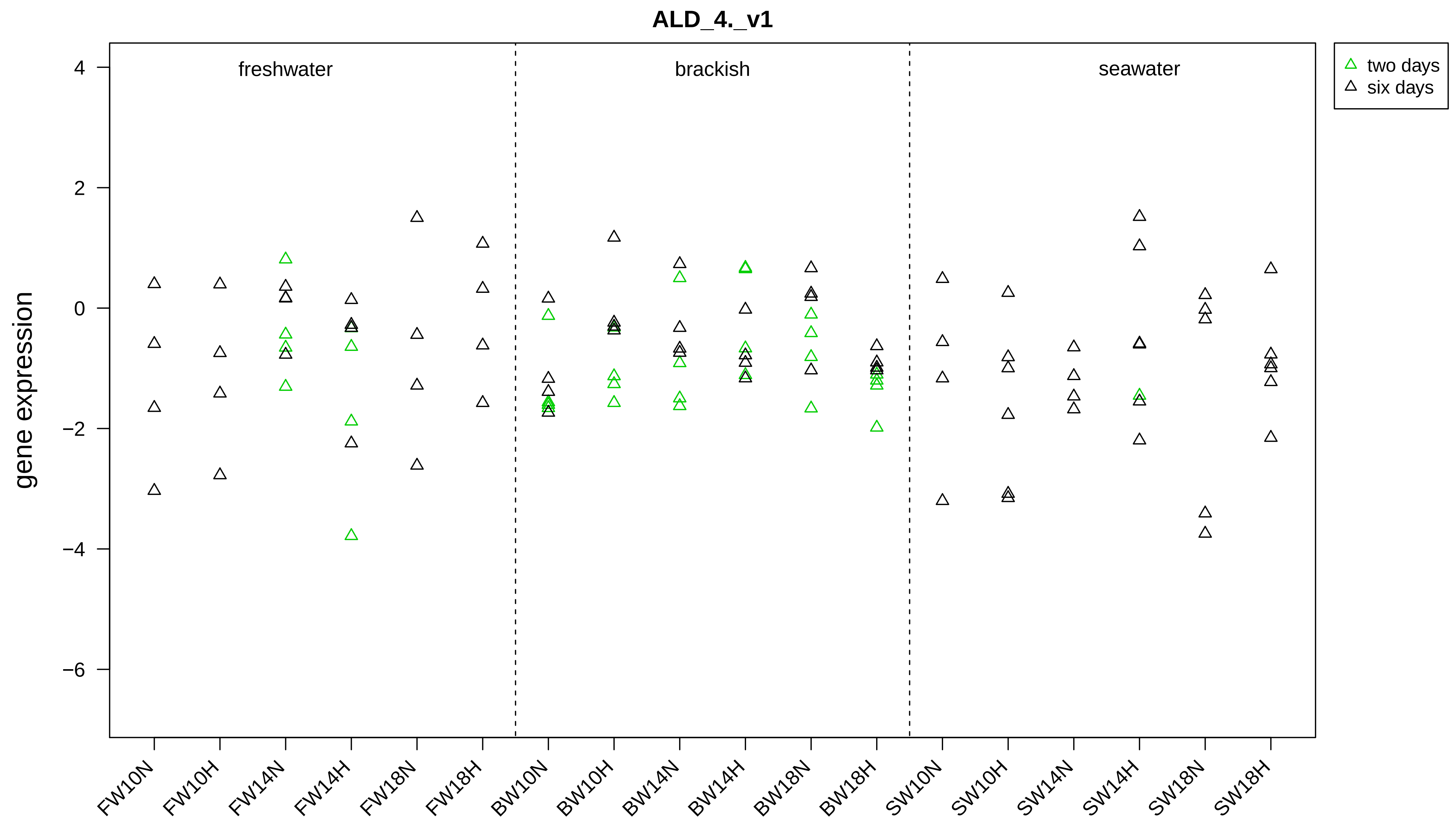

ALD_4._v1
gene expression
freshwater
brackish
seawater
two days
six days
4
2
0
−2
−4
−6
FW10N
FW10H
FW14N
FW14H
FW18N
FW18H
BW10N
BW10H
BW14N
BW14H
BW18N
BW18H
SW10N
SW10H
SW14N
SW14H
SW18N
SW18H

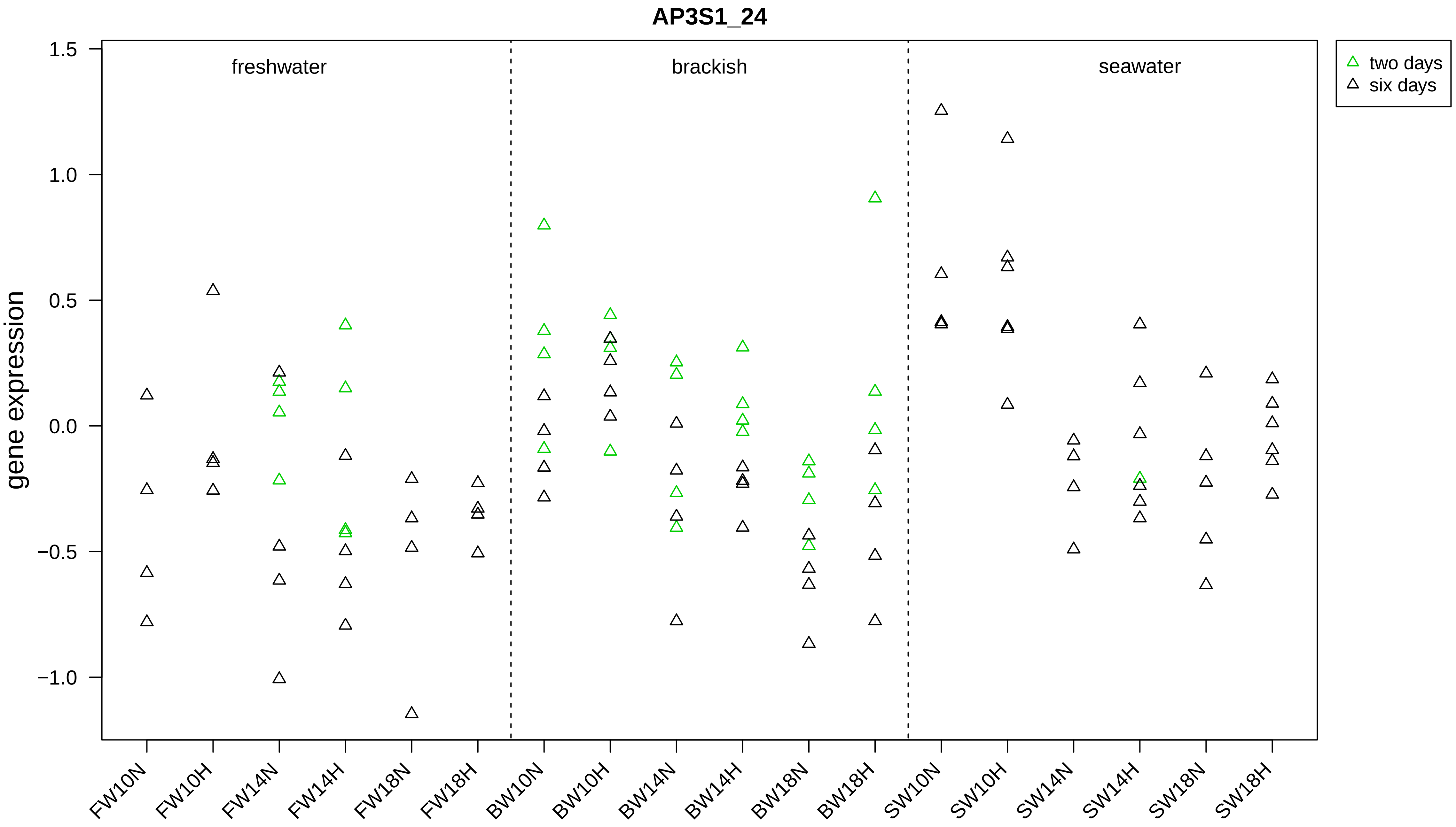

AP3S1_24
gene expression
freshwater
brackish
seawater
two days
six days
1.5
1.0
0.5
0.0
−0.5
−1.0
FW10N
FW10H
FW14N
FW14H
FW18N
FW18H
BW10N
BW10H
BW14N
BW14H
BW18N
BW18H
SW10N
SW10H
SW14N
SW14H
SW18N
SW18H

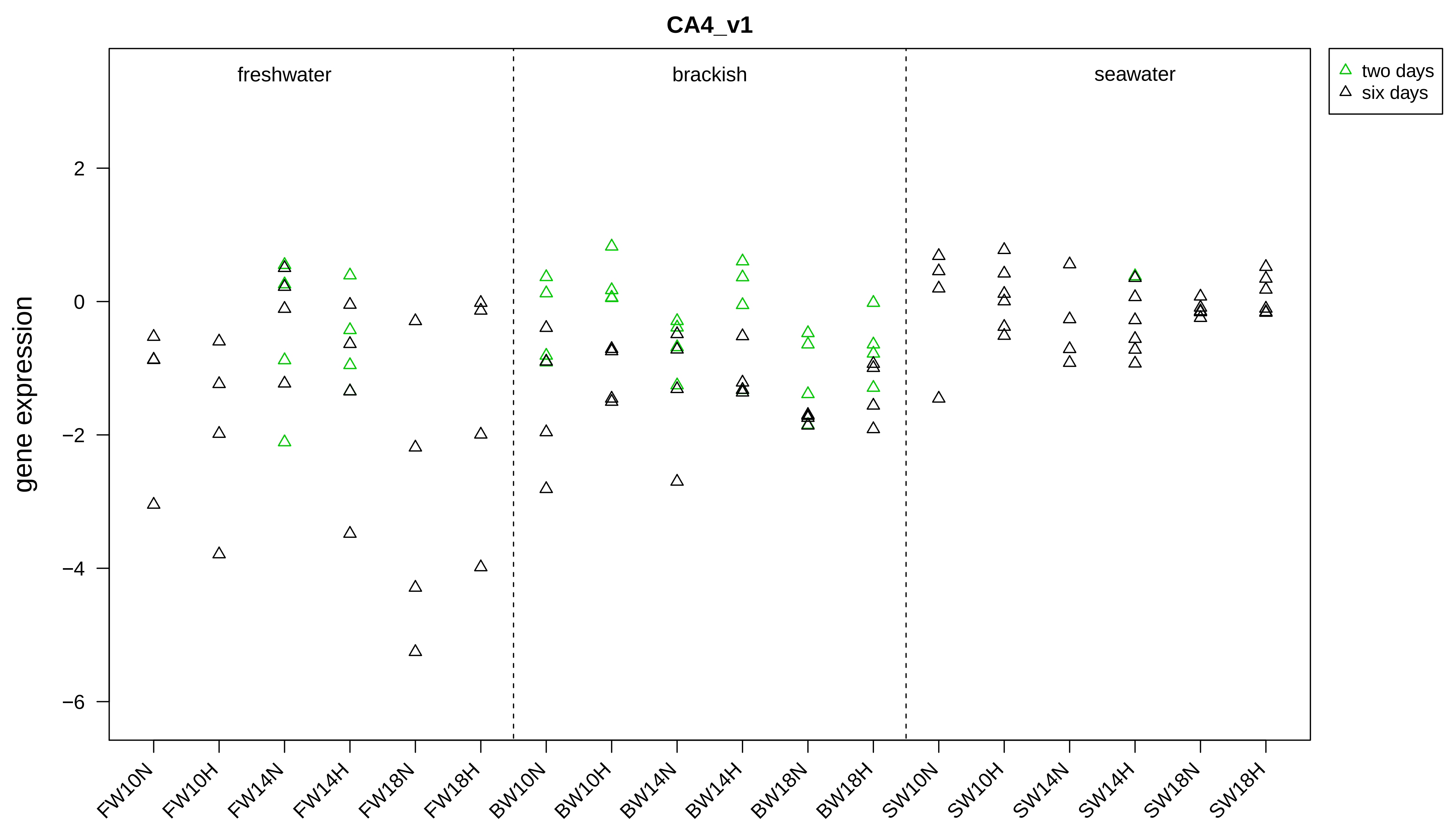
CA4_v1
freshwater
brackish
seawater
two days
six days
gene expression
2
0
−2
−4
−6
FW10N
FW10H
FW14N
FW14H
FW18N
FW18H
BW10N
BW10H
BW14N
BW14H
BW18N
BW18H
SW10N
SW10H
SW14N
SW14H
SW18N
SW18H

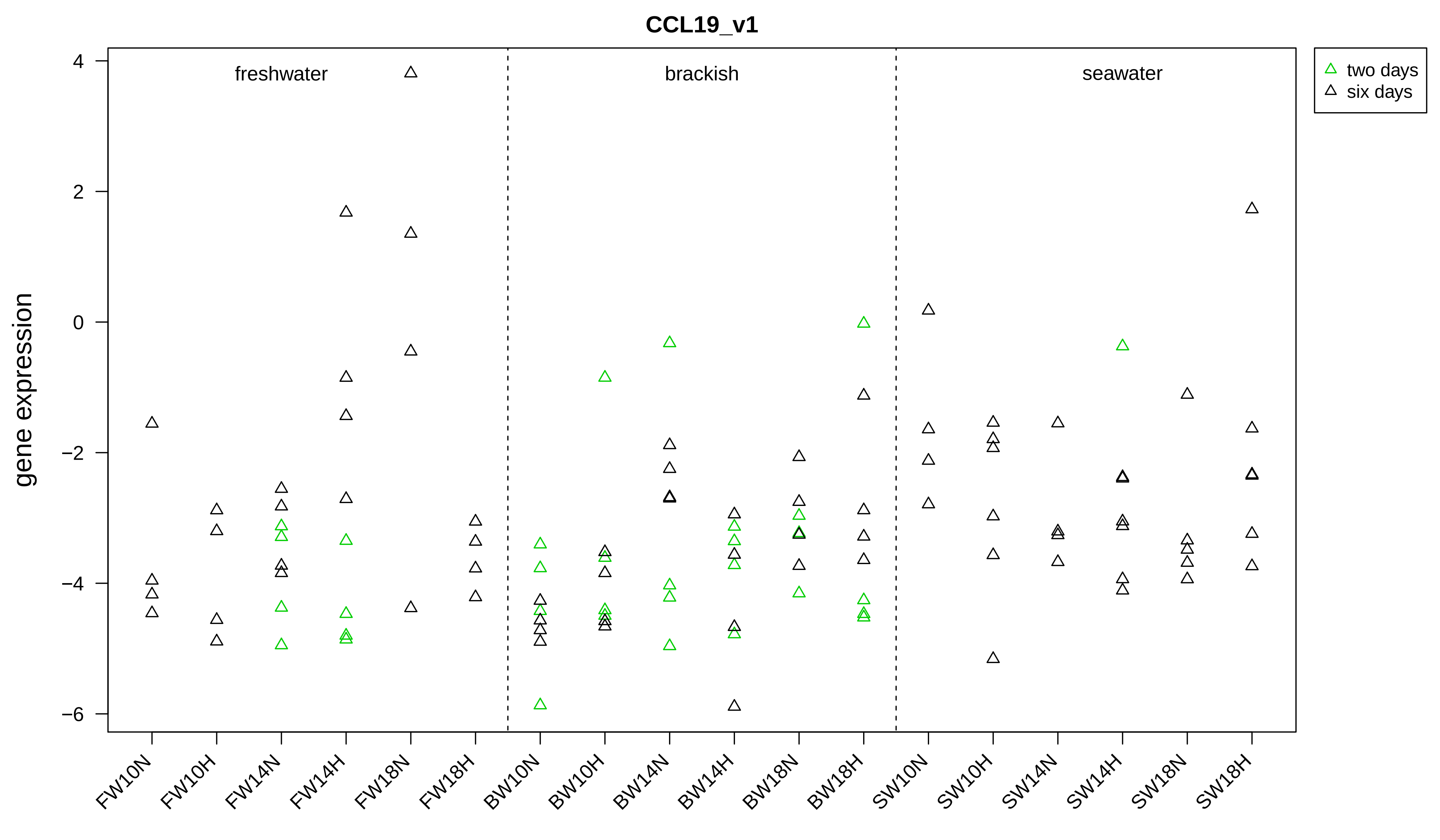
CCL19_v1
gene expression
freshwater
brackish
seawater
two days
six days
4
2
0
−2
−4
−6
FW10N
FW10H
FW14N
FW14H
FW18N
FW18H
BW10N
BW10H
BW14N
BW14H
BW18N
BW18H
SW10N
SW10H
SW14N
SW14H
SW18N
SW18H

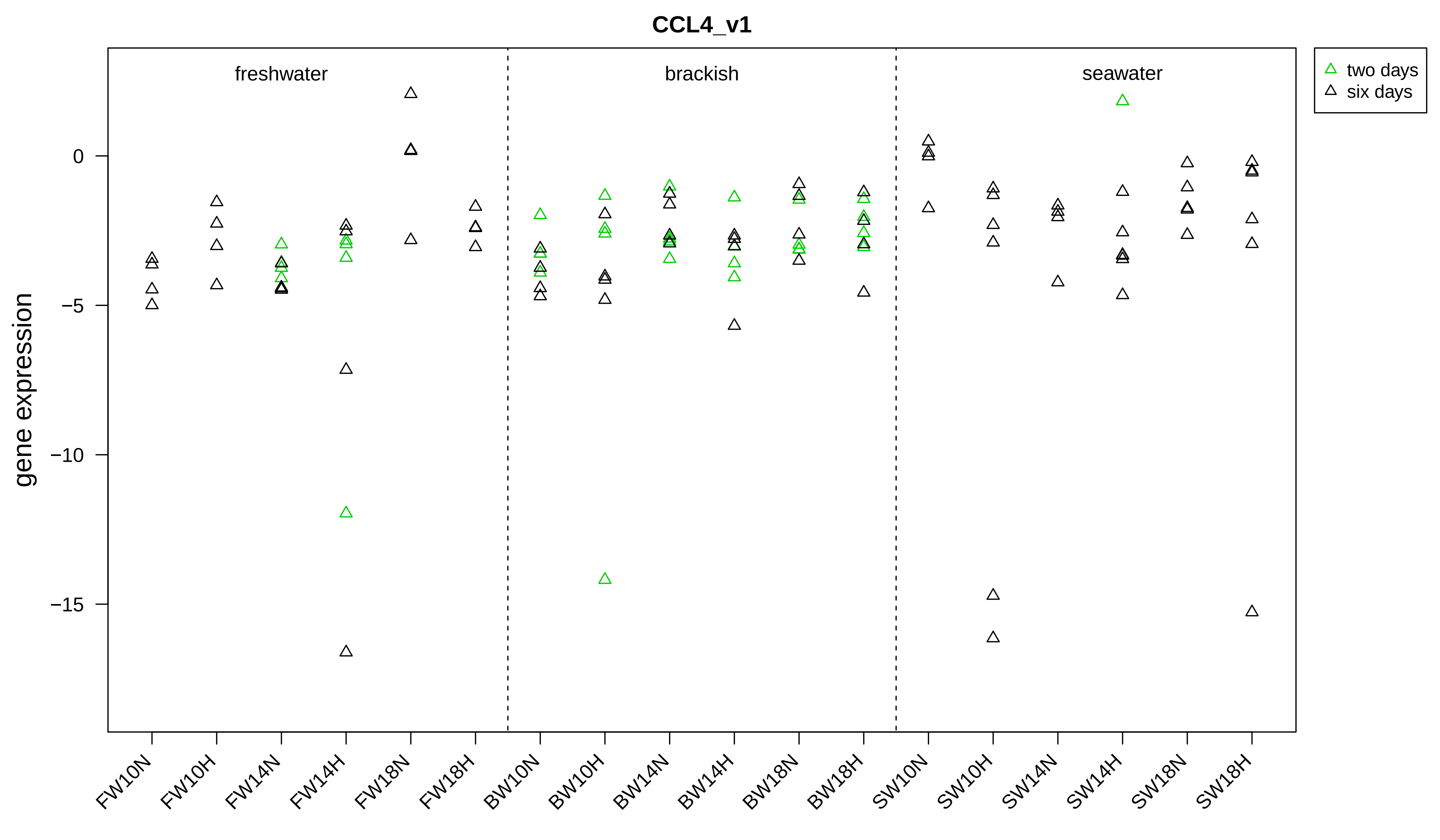
CCL4_v1
freshwater
brackish
seawater
two days
six days
gene expression
0
−5
−10
−15
FW10N
FW10H
FW14N
FW14H
FW18N
FW18H
BW10N
BW10H
BW14N
BW14H
BW18N
BW18H
SW10N
SW10H
SW14N
SW14H
SW18N
SW18H

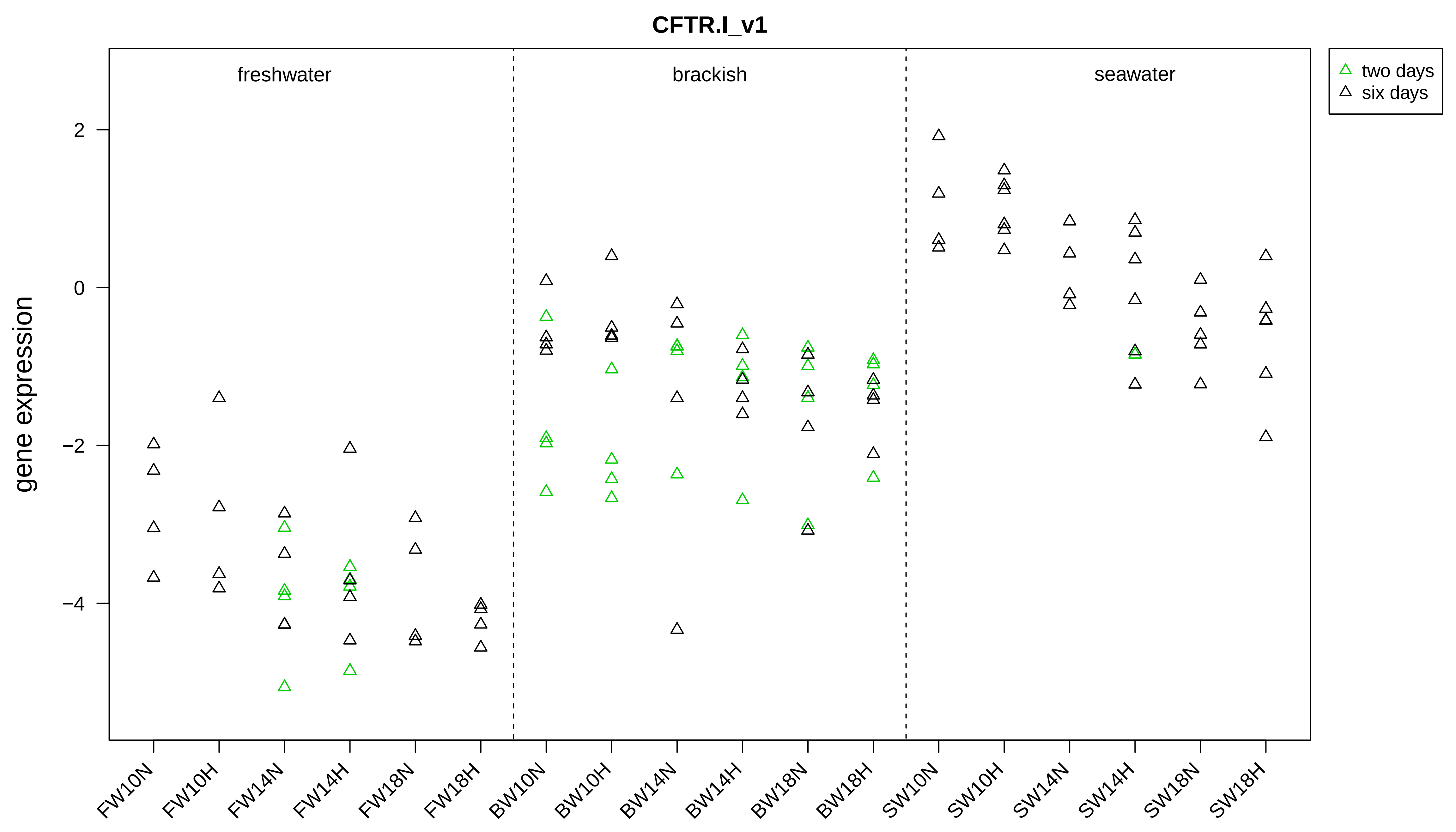
CFTR.I_v1
freshwater
brackish
seawater
two days
six days
gene expression
2
0
−2
−4
FW10N
FW10H
FW14N
FW14H
FW18N
FW18H
BW10N
BW10H
BW14N
BW14H
BW18N
BW18H
SW10N
SW10H
SW14N
SW14H
SW18N
SW18H

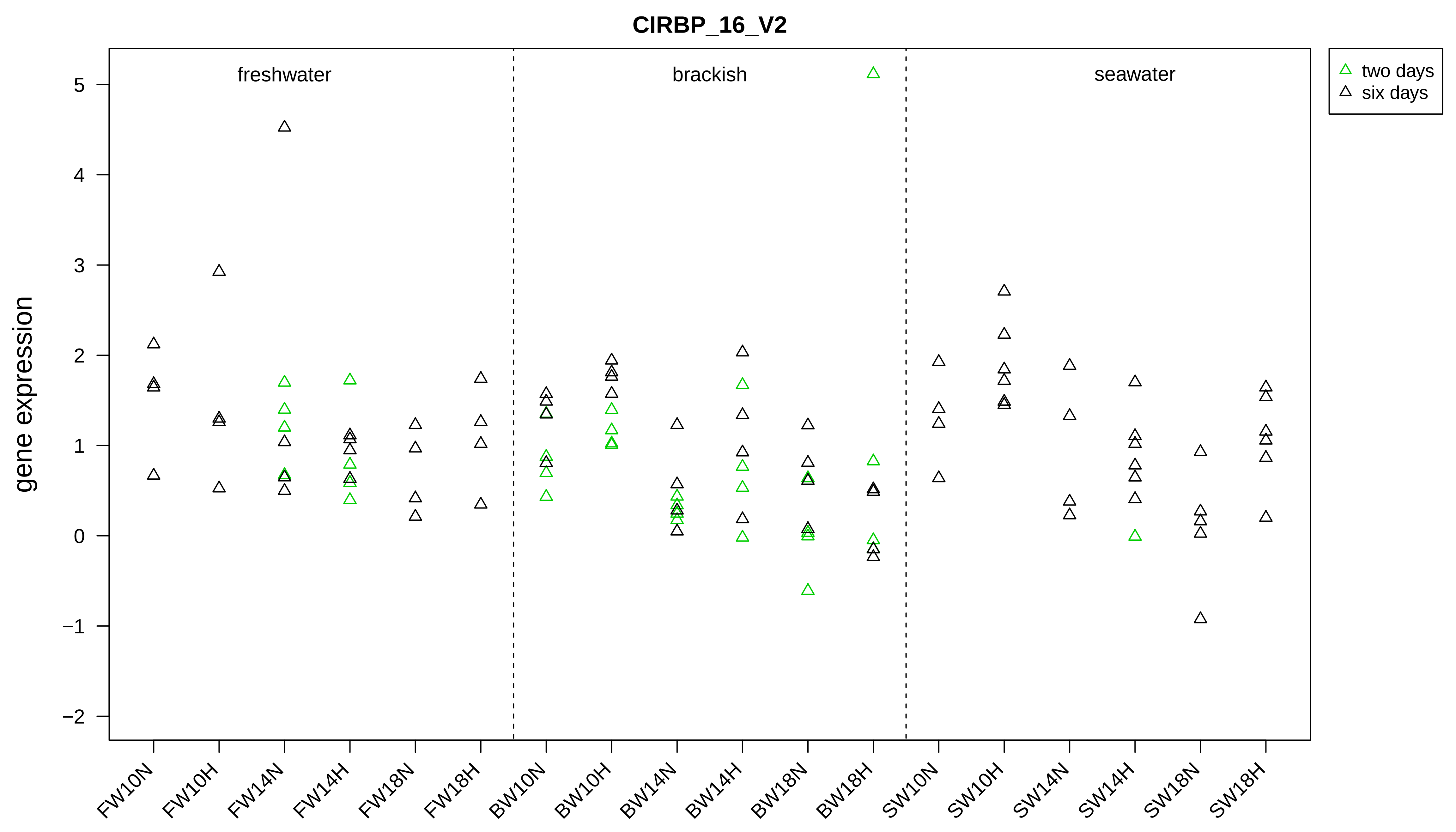
CIRBP_16_V2
freshwater
brackish
seawater
two days
six days
gene expression
5
4
3
2
1
0
−1
−2
FW10N
FW10H
FW14N
FW14H
FW18N
FW18H
BW10N
BW10H
BW14N
BW14H
BW18N
BW18H
SW10N
SW10H
SW14N
SW14H
SW18N
SW18H

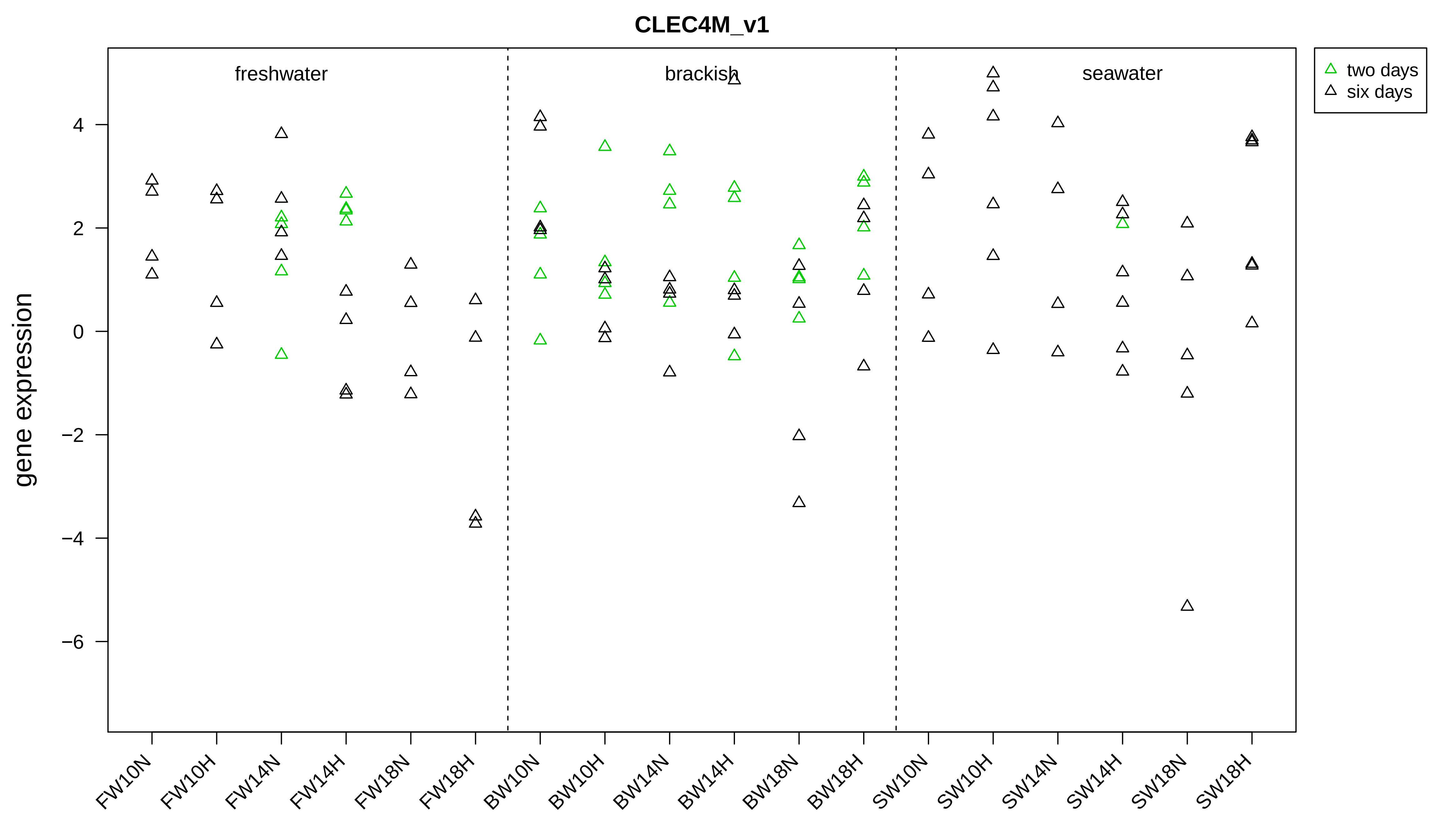

CLEC4M_v1
freshwater
brackish
seawater
two days
six days
gene expression
4
2
0
−2
−4
−6
FW10N
FW10H
FW14N
FW14H
FW18N
FW18H
BW10N
BW10H
BW14N
BW14H
BW18N
BW18H
SW10N
SW10H
SW14N
SW14H
SW18N
SW18H

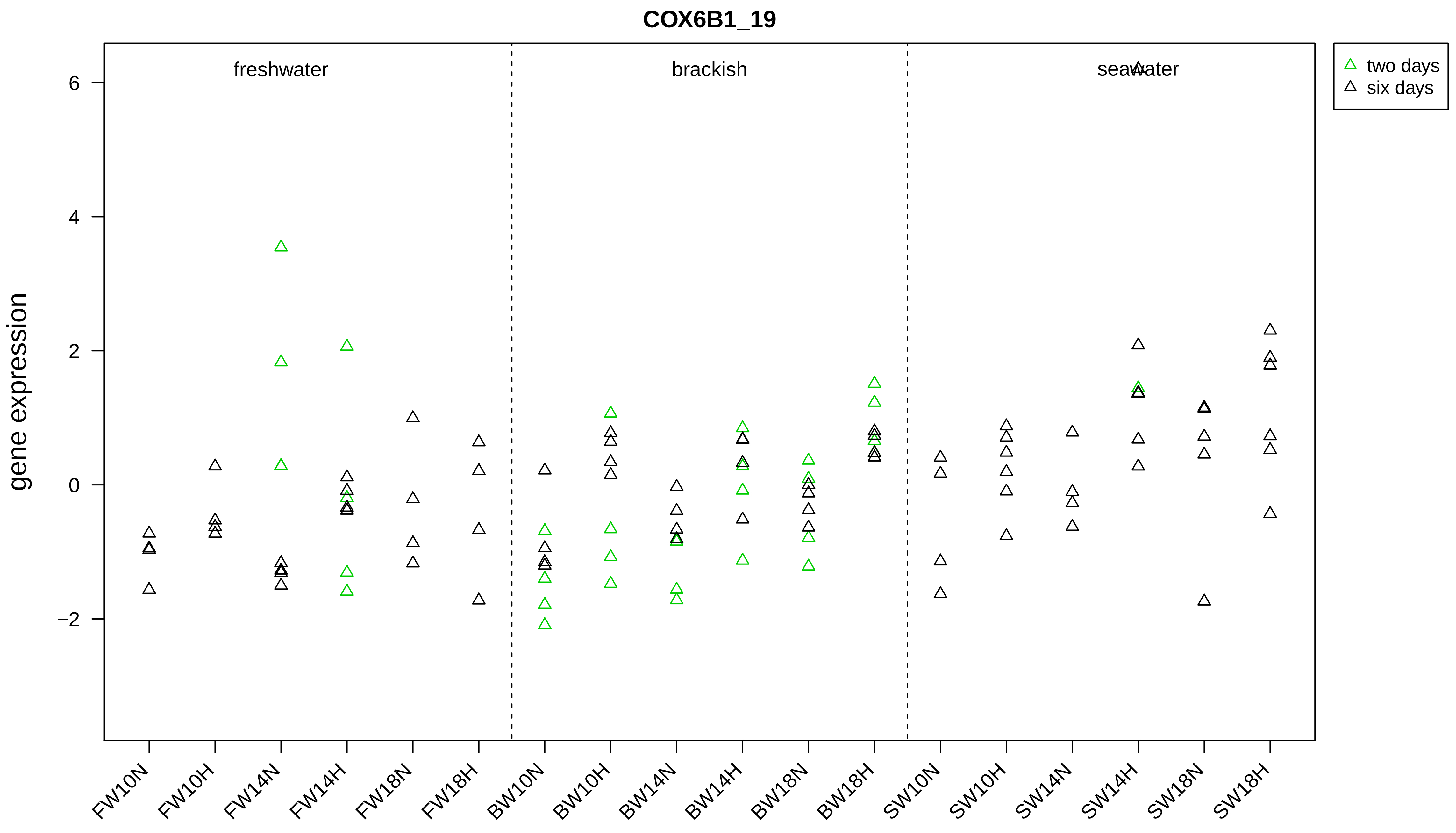

COX6B1_19
gene expression
freshwater
brackish
seawater
two days
six days
6
4
2
0
−2
FW10N
FW10H
FW14N
FW14H
FW18N
FW18H
BW10N
BW10H
BW14N
BW14H
BW18N
BW18H
SW10N
SW10H
SW14N
SW14H
SW18N
SW18H

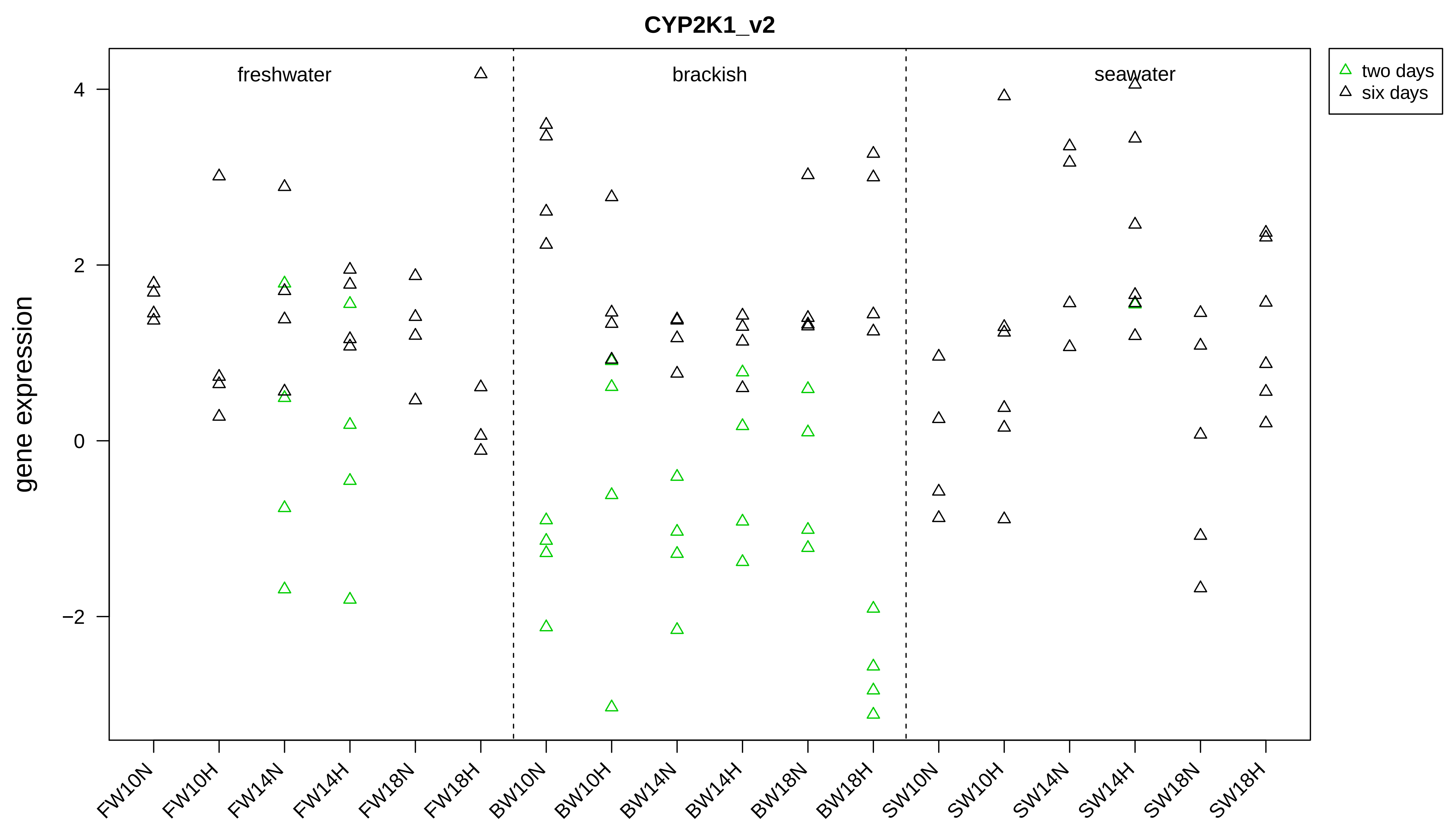
CYP2K1_v2
two days
six days
freshwater
brackish
seawater
gene expression
4
2
0
−2
FW10N
FW10H
FW14N
FW14H
FW18N
FW18H
BW10N
BW10H
BW14N
BW14H
BW18N
BW18H
SW10N
SW10H
SW14N
SW14H
SW18N
SW18H

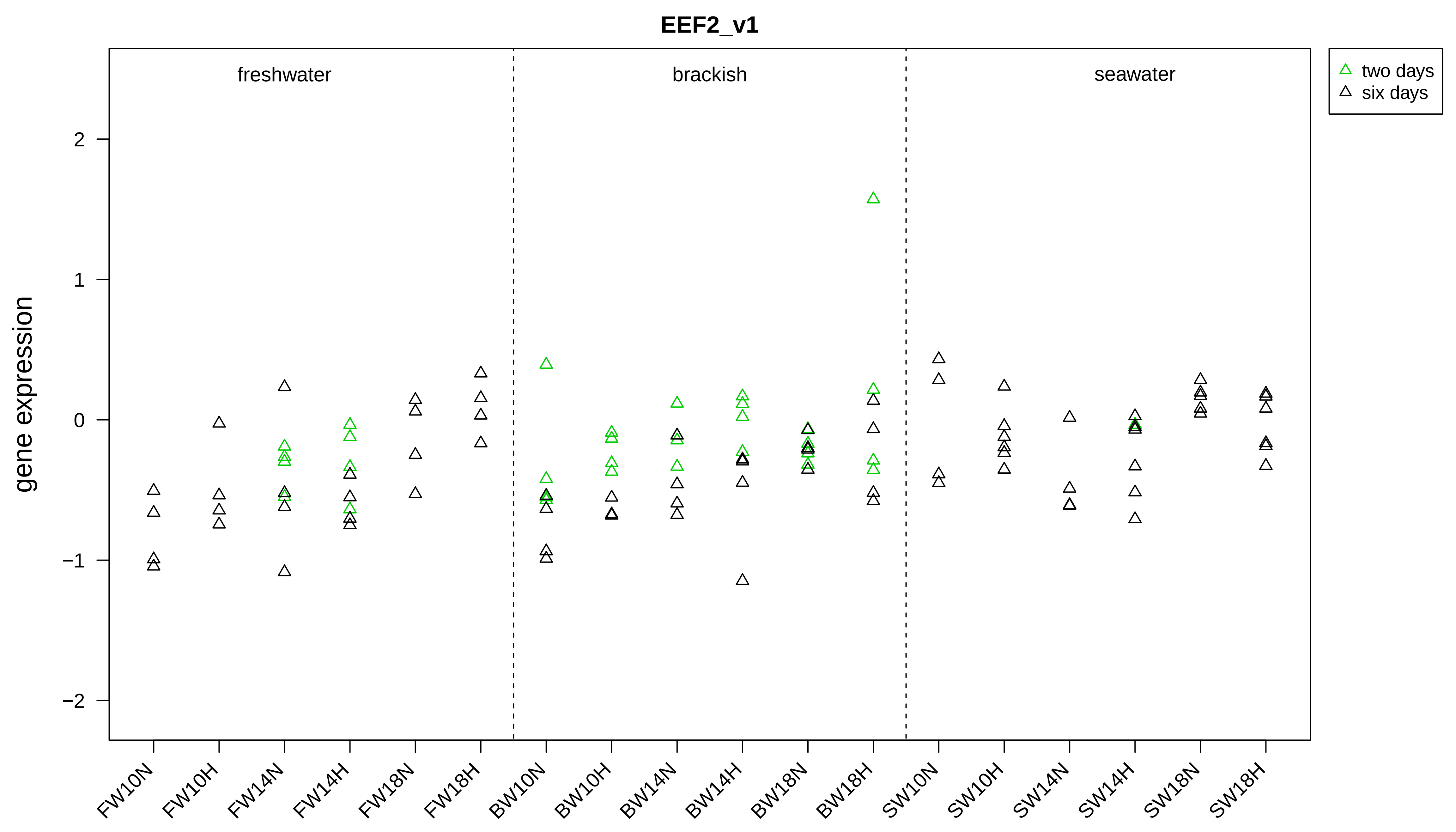

EEF2_v1
freshwater
brackish
seawater
two days
six days
gene expression
2
1
0
−1
−2
FW10N
FW10H
FW14N
FW14H
FW18N
FW18H
BW10N
BW10H
BW14N
BW14H
BW18N
BW18H
SW10N
SW10H
SW14N
SW14H
SW18N
SW18H

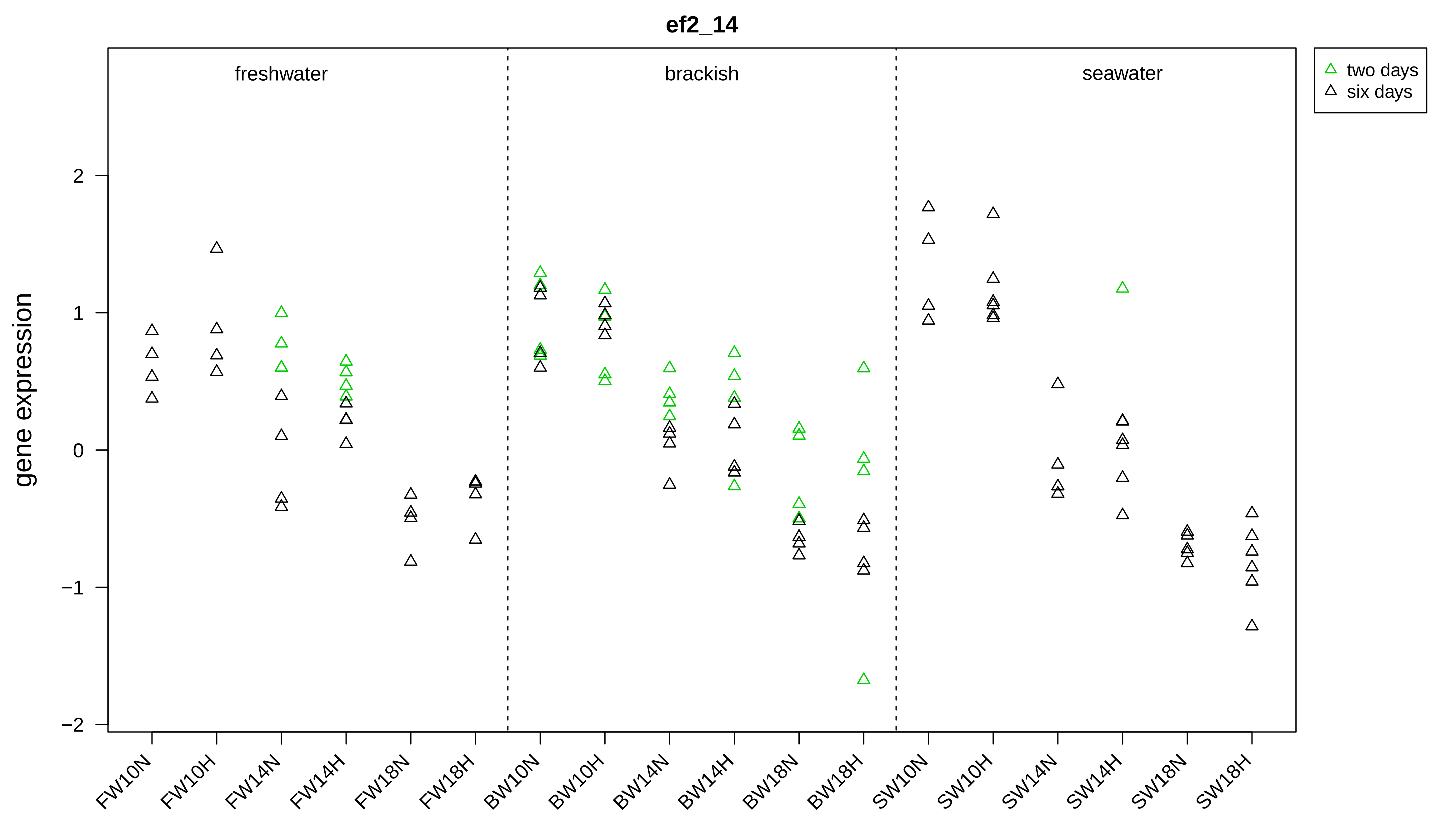
ef2_14
freshwater
brackish
seawater
two days
six days
gene expression
2
1
0
−1
−2
FW10N
FW10H
FW14N
FW14H
FW18N
FW18H
BW10N
BW10H
BW14N
BW14H
BW18N
BW18H
SW10N
SW10H
SW14N
SW14H
SW18N
SW18H

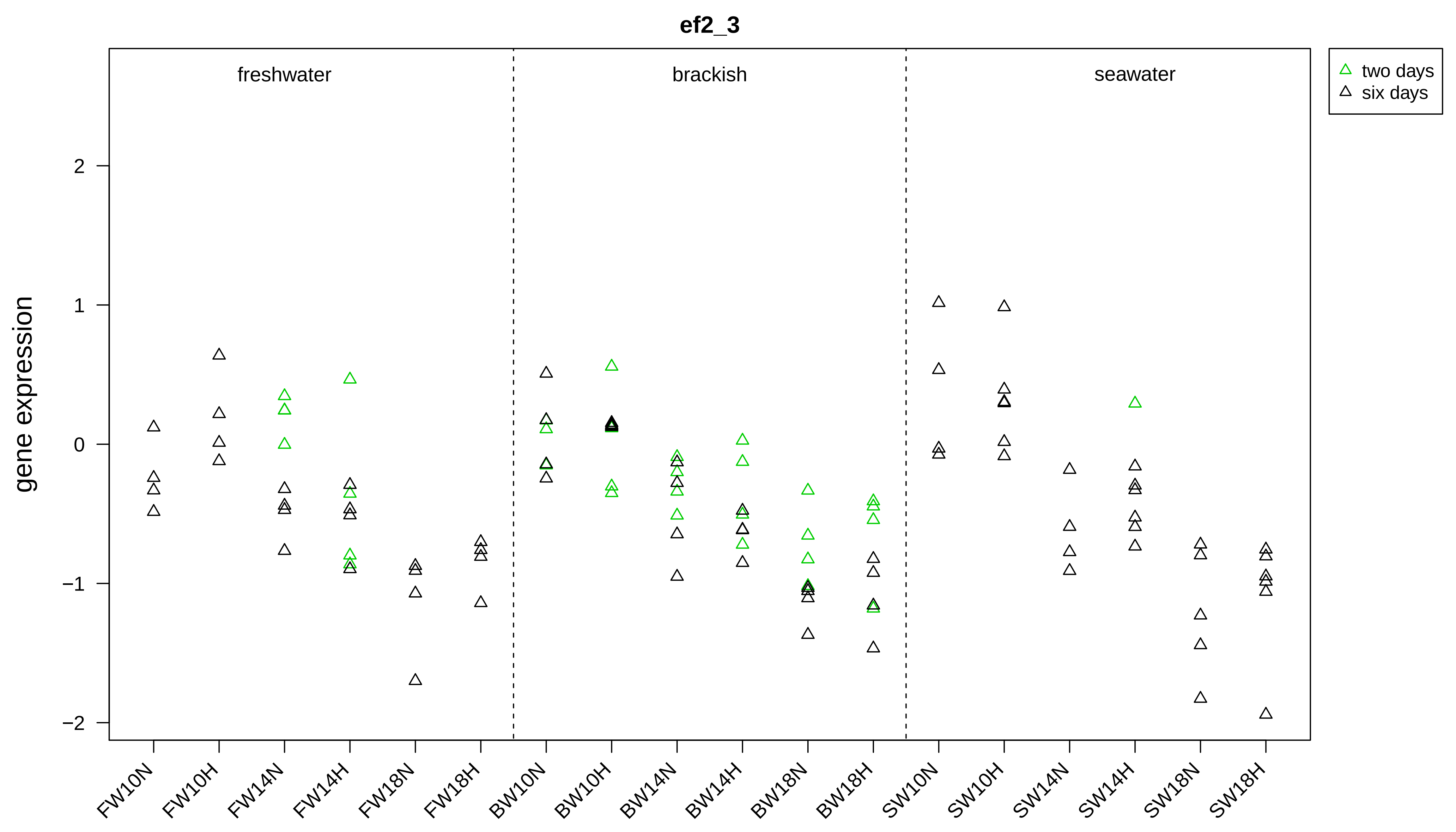

ef2_3
freshwater
brackish
seawater
two days
six days
gene expression
2
1
0
−1
−2
FW10N
FW10H
FW14N
FW14H
FW18N
FW18H
BW10N
BW10H
BW14N
BW14H
BW18N
BW18H
SW10N
SW10H
SW14N
SW14H
SW18N
SW18H

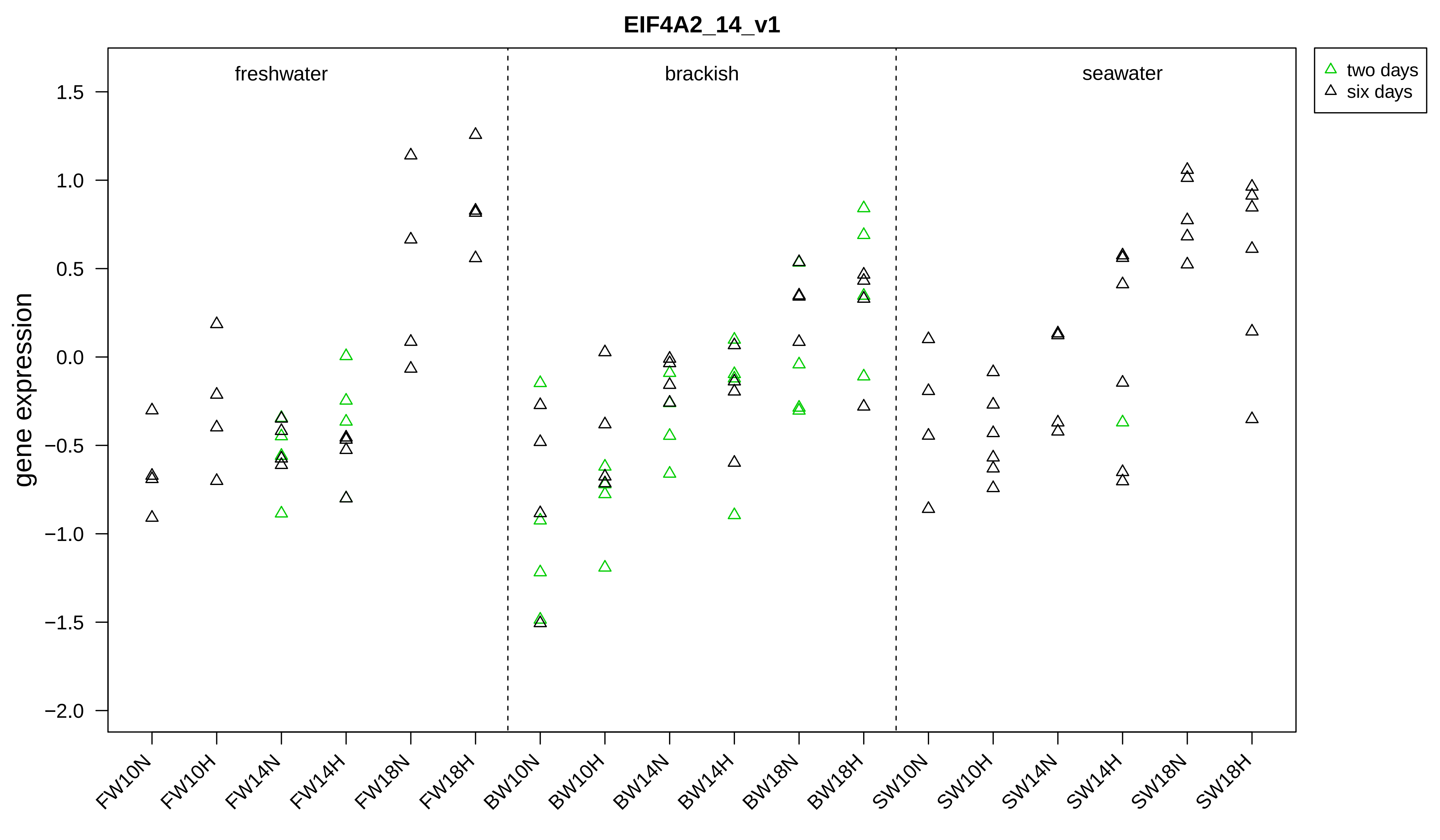

EIF4A2_14_v1
freshwater
brackish
seawater
two days
six days
gene expression
1.5
1.0
0.5
0.0
−0.5
−1.0
−1.5
−2.0
FW10N
FW10H
FW14N
FW14H
FW18N
FW18H
BW10N
BW10H
BW14N
BW14H
BW18N
BW18H
SW10N
SW10H
SW14N
SW14H
SW18N
SW18H

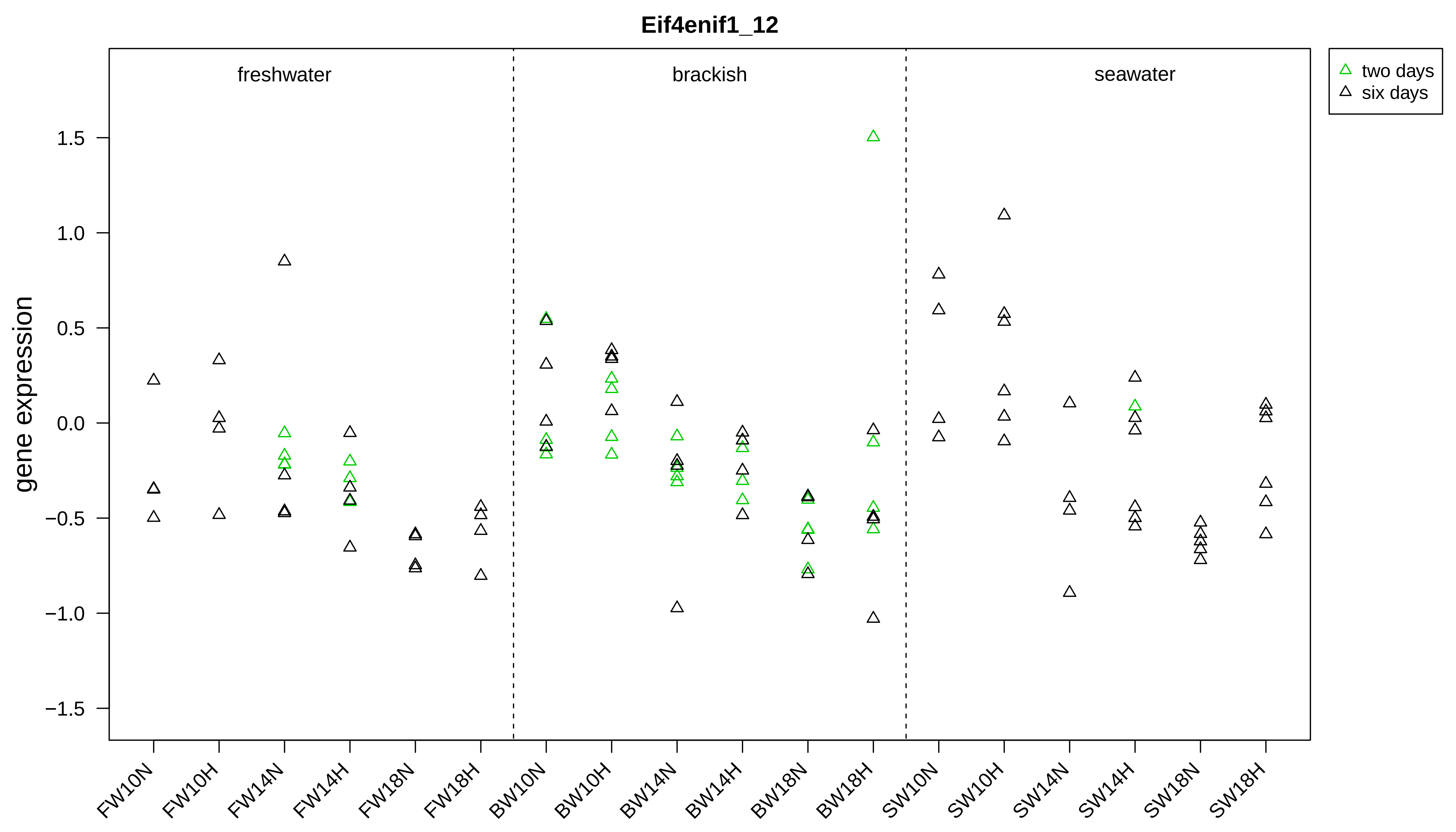

Eif4enif1_12
freshwater
brackish
seawater
two days
six days
gene expression
1.5
1.0
0.5
0.0
−0.5
−1.0
−1.5
FW10N
FW10H
FW14N
FW14H
FW18N
FW18H
BW10N
BW10H
BW14N
BW14H
BW18N
BW18H
SW10N
SW10H
SW14N
SW14H
SW18N
SW18H

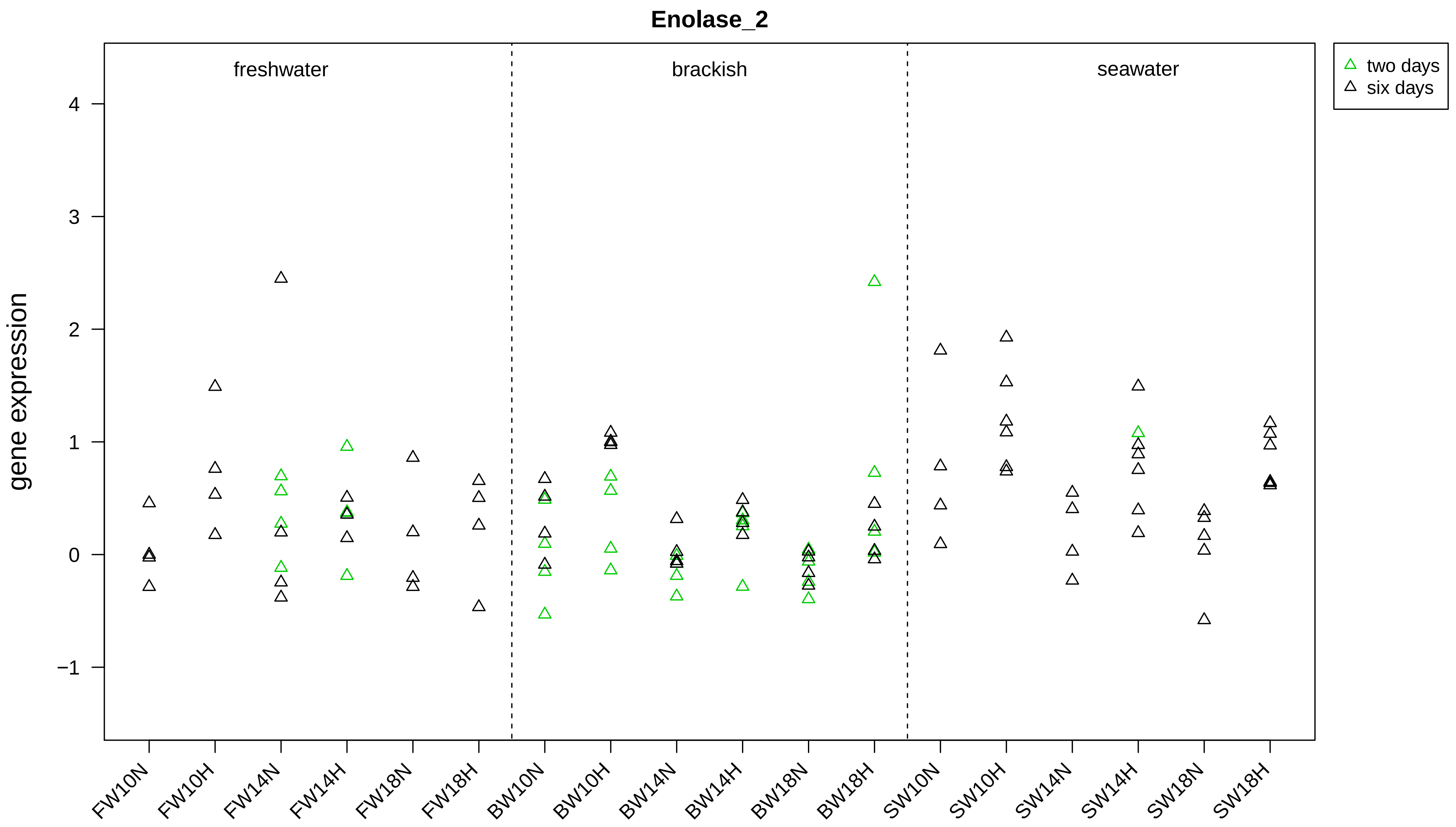
Enolase_2
freshwater
brackish
seawater
two days
six days
gene expression
4
3
2
1
0
−1
FW10N
FW10H
FW14N
FW14H
FW18N
FW18H
BW10N
BW10H
BW14N
BW14H
BW18N
BW18H
SW10N
SW10H
SW14N
SW14H
SW18N
SW18H

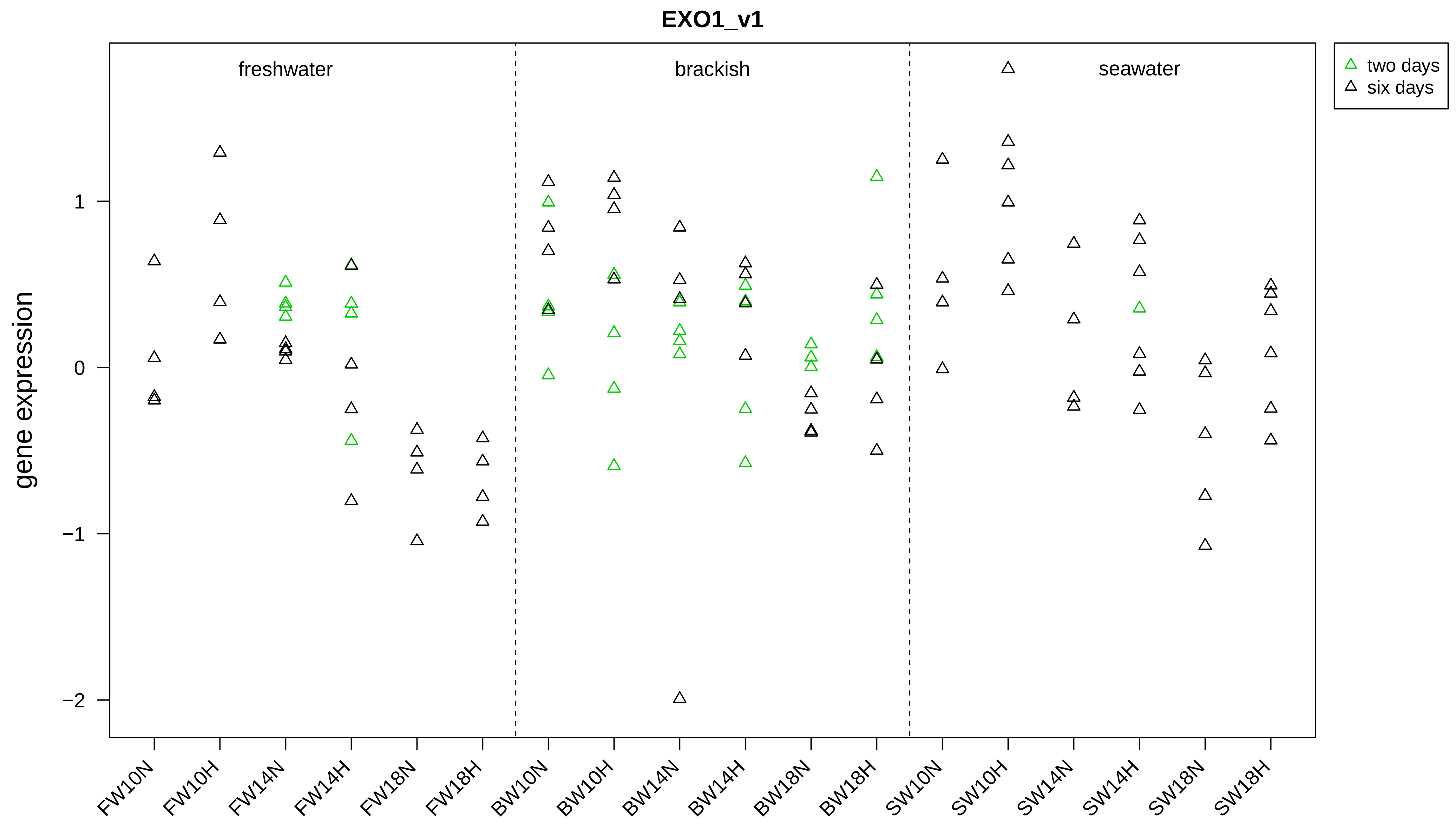
EXO1_v1
freshwater
brackish
seawater
two days
six days
gene expression
1
0
−1
−2
FW10N
FW10H
FW14N
FW14H
FW18N
FW18H
BW10N
BW10H
BW14N
BW14H
BW18N
BW18H
SW10N
SW10H
SW14N
SW14H
SW18N
SW18H

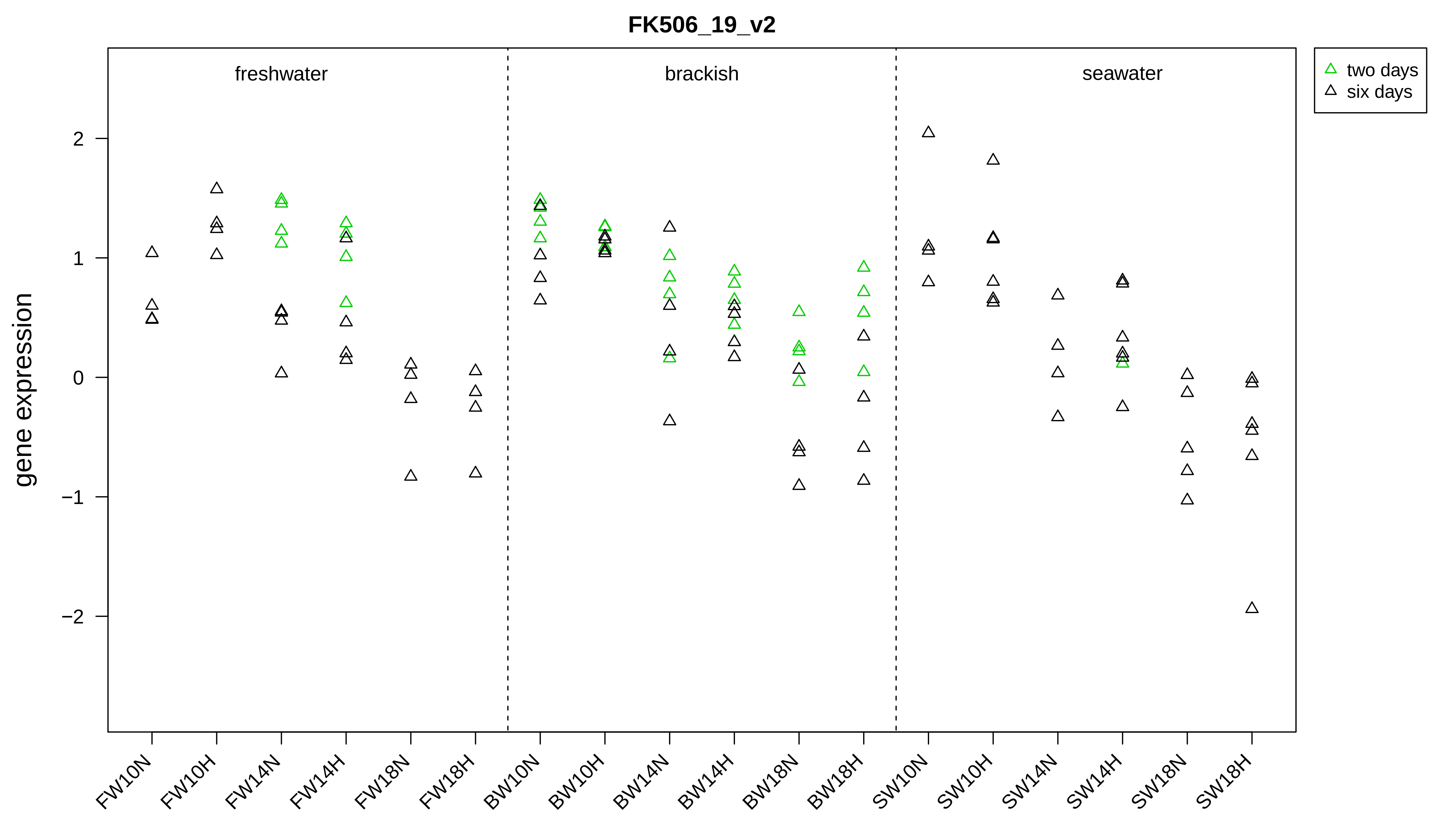
FK506_19_v2
freshwater
brackish
seawater
two days
six days
gene expression
2
1
0
−1
−2
FW10N
FW10H
FW14N
FW14H
FW18N
FW18H
BW10N
BW10H
BW14N
BW14H
BW18N
BW18H
SW10N
SW10H
SW14N
SW14H
SW18N
SW18H

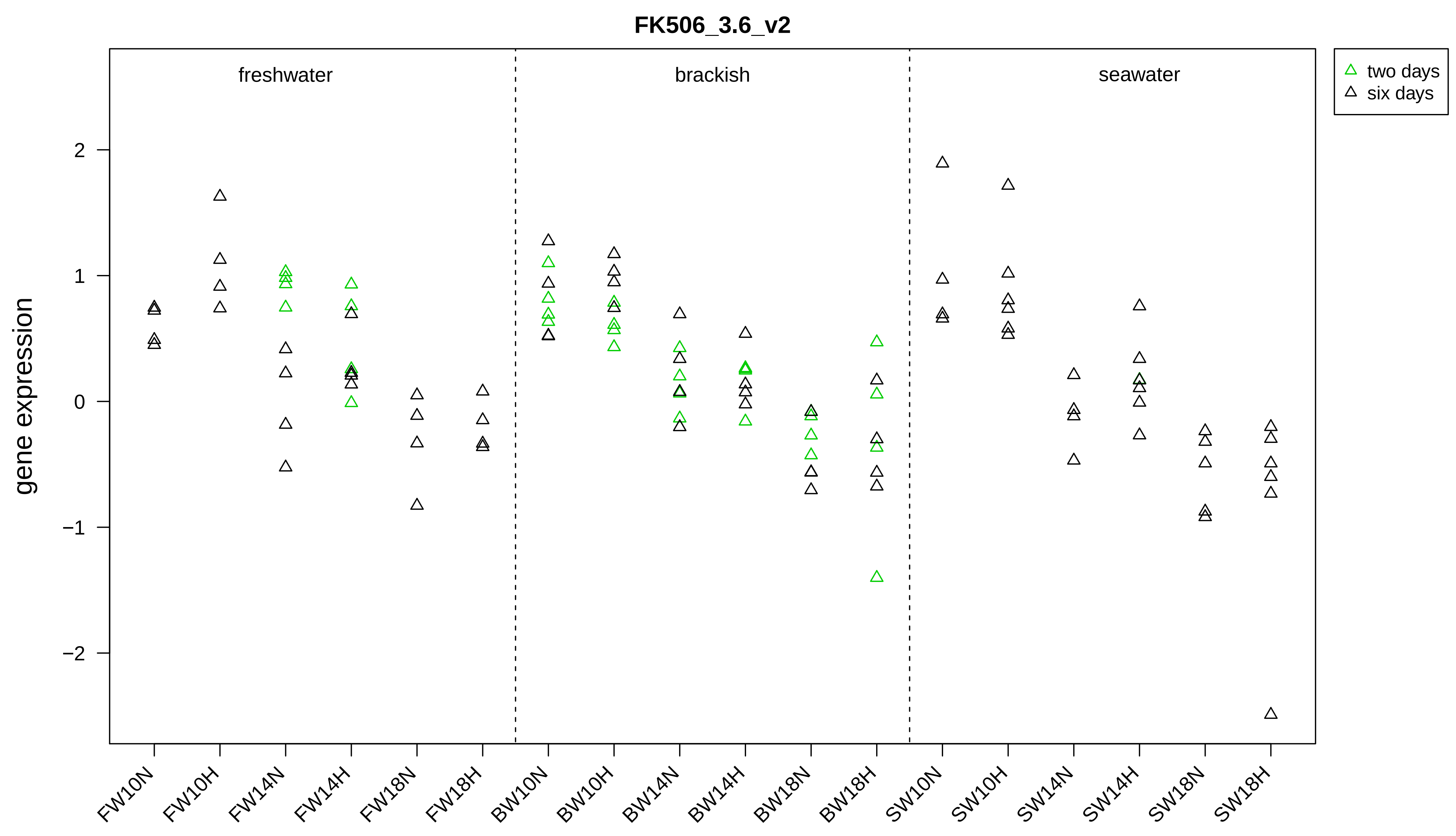
FK506_3.6_v2
freshwater
brackish
seawater
two days
six days
gene expression
2
1
0
−1
−2
FW10N
FW10H
FW14N
FW14H
FW18N
FW18H
BW10N
BW10H
BW14N
BW14H
BW18N
BW18H
SW10N
SW10H
SW14N
SW14H
SW18N
SW18H

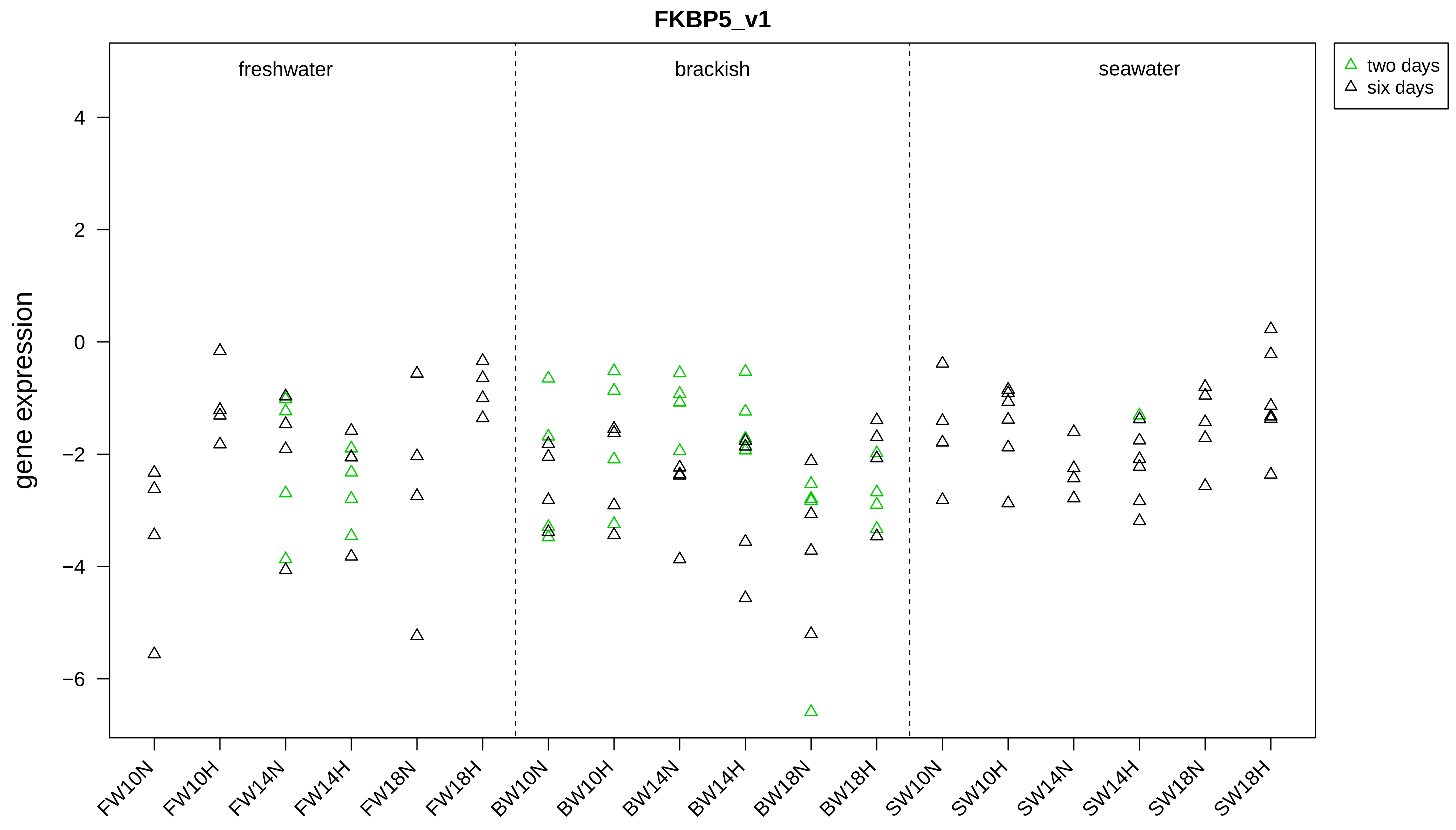

FKBP5_v1
freshwater
brackish
seawater
two days
six days
gene expression
4
2
0
−2
−4
−6
FW10N
FW10H
FW14N
FW14H
FW18N
FW18H
BW10N
BW10H
BW14N
BW14H
BW18N
BW18H
SW10N
SW10H
SW14N
SW14H
SW18N
SW18H

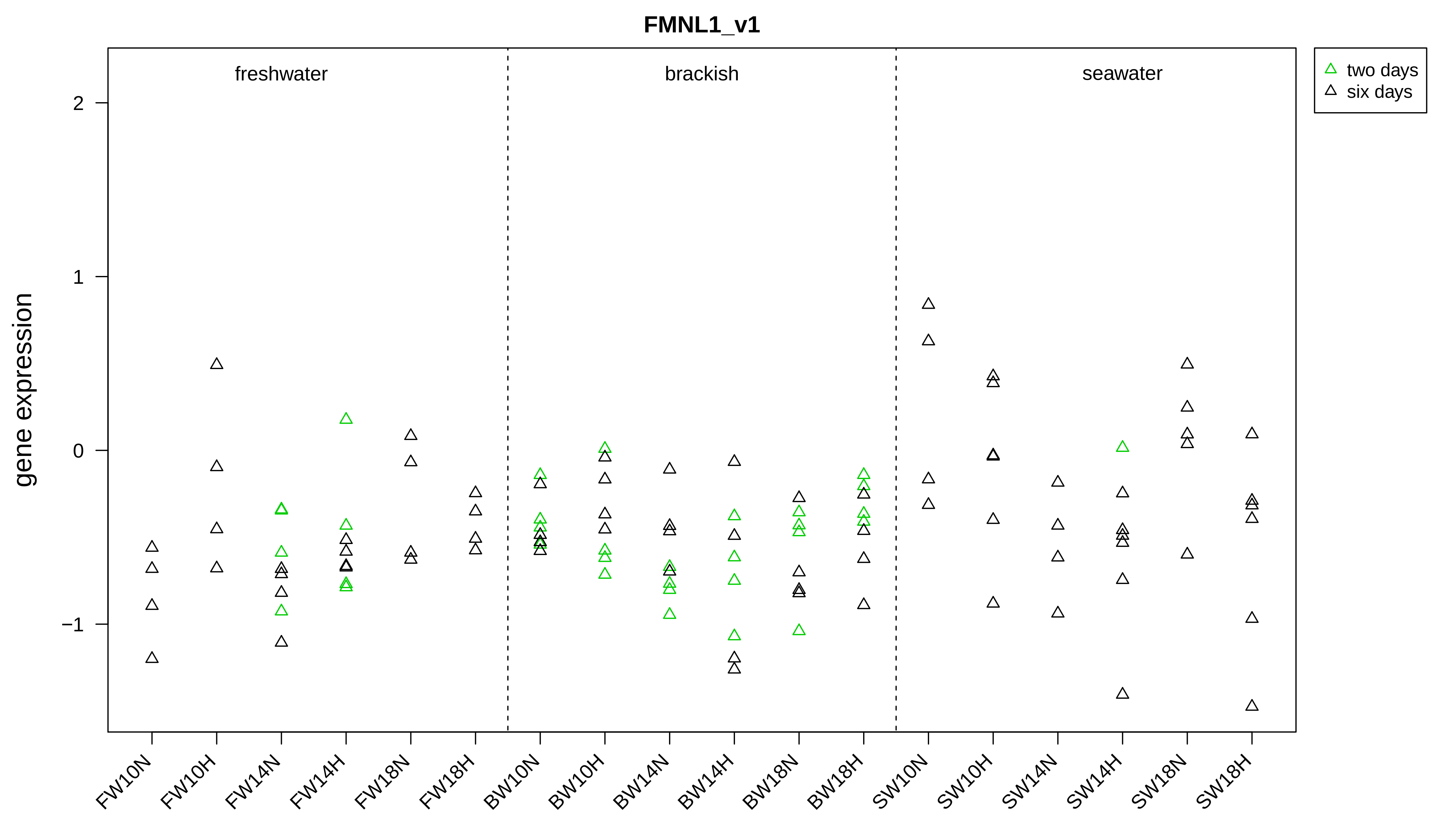

FMNL1_v1
freshwater
brackish
seawater
two days
six days
gene expression
2
1
0
−1
FW10N
FW10H
FW14N
FW14H
FW18N
FW18H
BW10N
BW10H
BW14N
BW14H
BW18N
BW18H
SW10N
SW10H
SW14N
SW14H
SW18N
SW18H

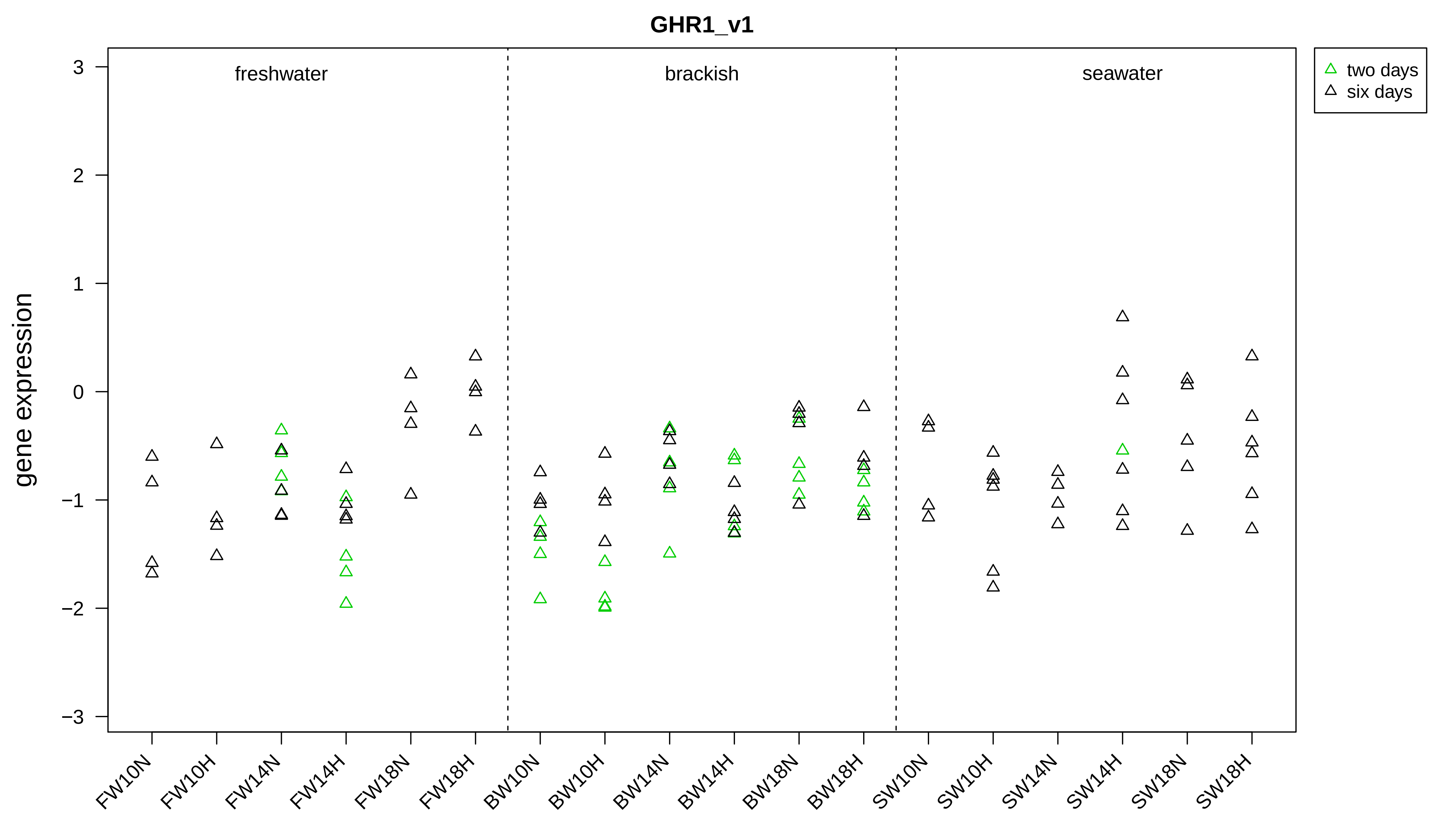

GHR1_v1
gene expression
freshwater
brackish
seawater
two days
six days
3
2
1
0
−1
−2
−3
FW10N
FW10H
FW14N
FW14H
FW18N
FW18H
BW10N
BW10H
BW14N
BW14H
BW18N
BW18H
SW10N
SW10H
SW14N
SW14H
SW18N
SW18H

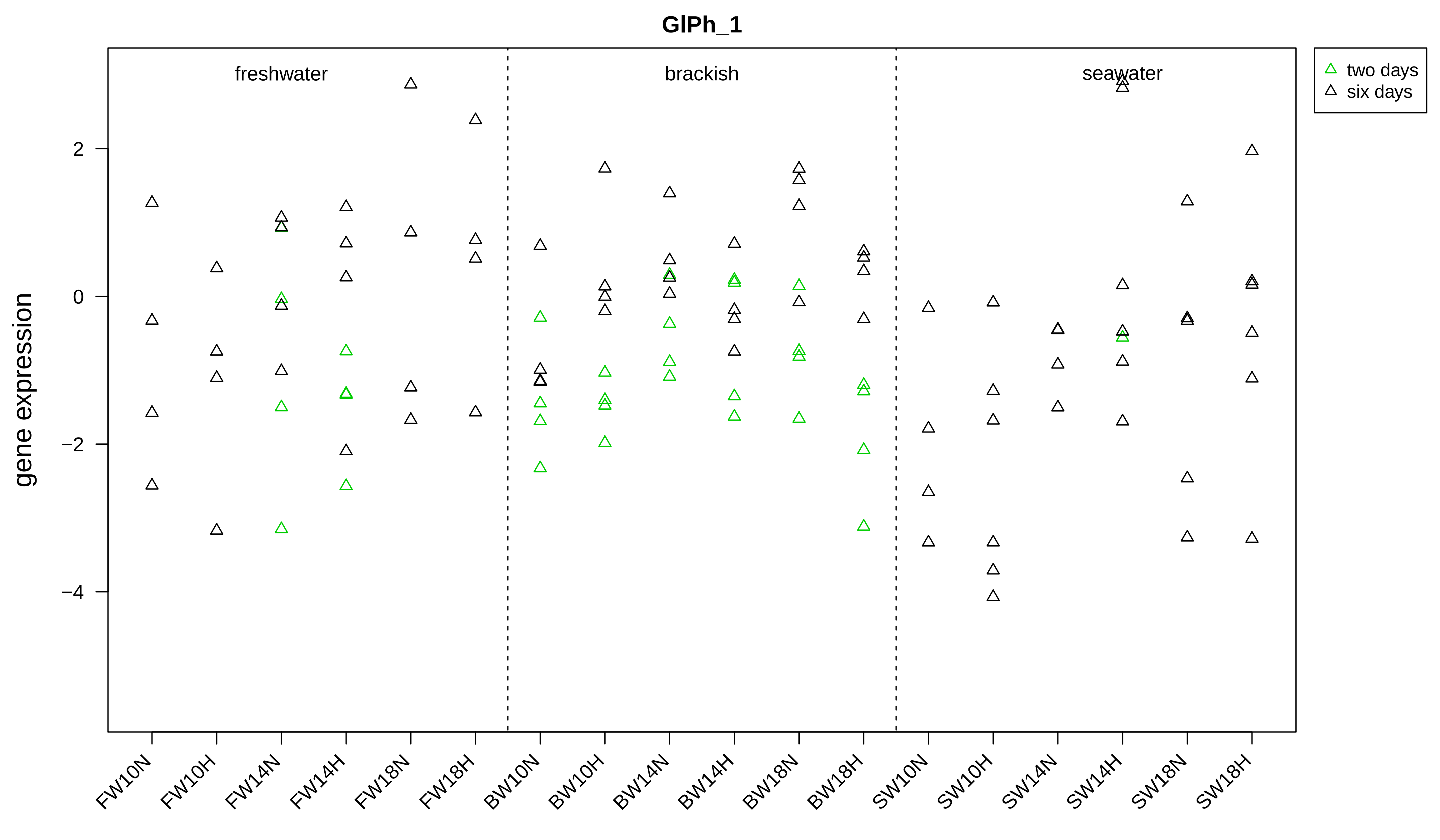
GlPh_1
freshwater
brackish
seawater
two days
six days
gene expression
2
0
−2
−4
FW10N
FW10H
FW14N
FW14H
FW18N
FW18H
BW10N
BW10H
BW14N
BW14H
BW18N
BW18H
SW10N
SW10H
SW14N
SW14H
SW18N
SW18H

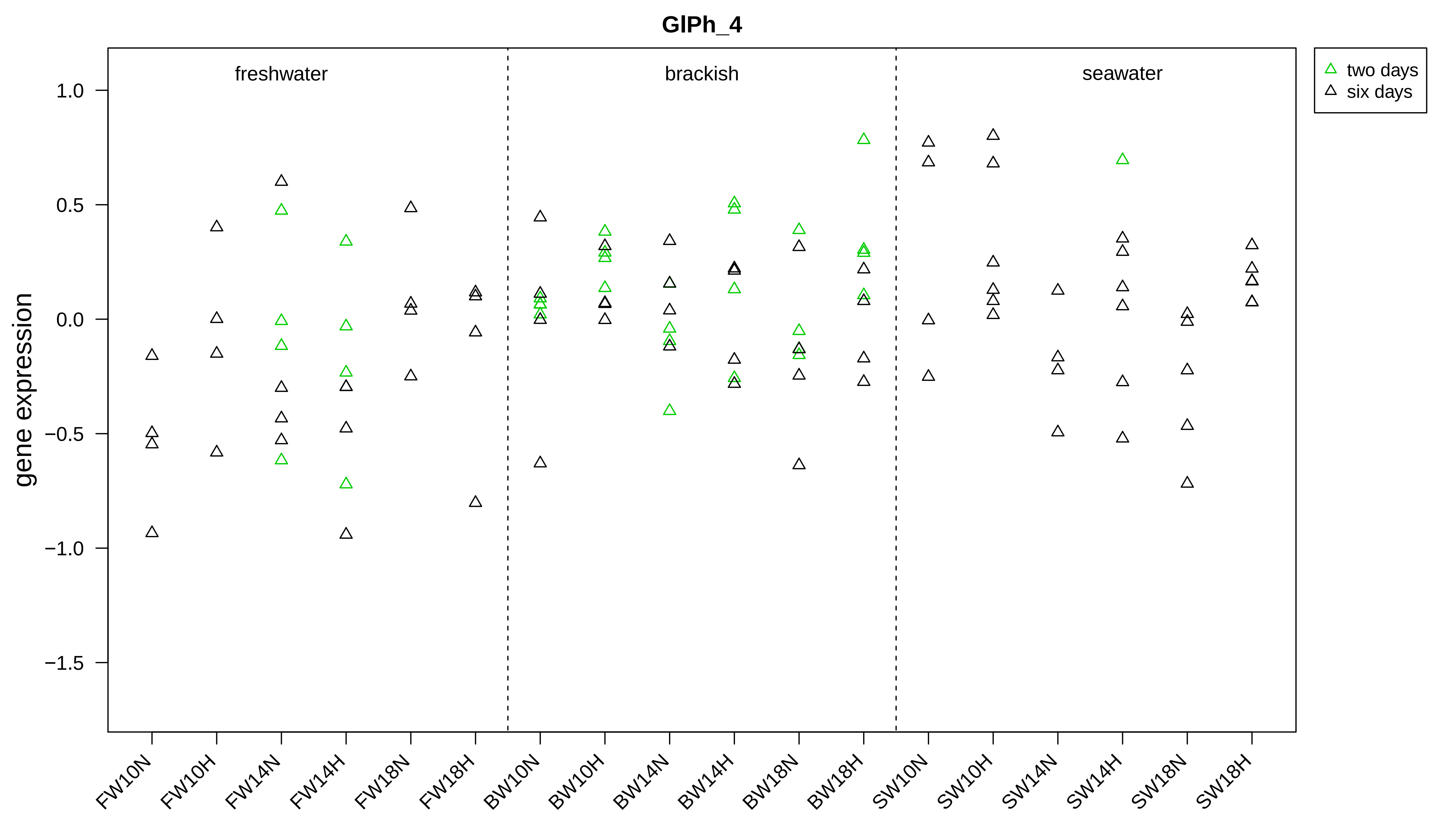
GlPh_4
gene expression
1.0
0.5
0.0
−0.5
−1.0
−1.5
freshwater
brackish
seawater
two days
six days
FW10N
FW10H
FW14N
FW14H
FW18N
FW18H
BW10N
BW10H
BW14N
BW14H
BW18N
BW18H
SW10N
SW10H
SW14N
SW14H
SW18N
SW18H

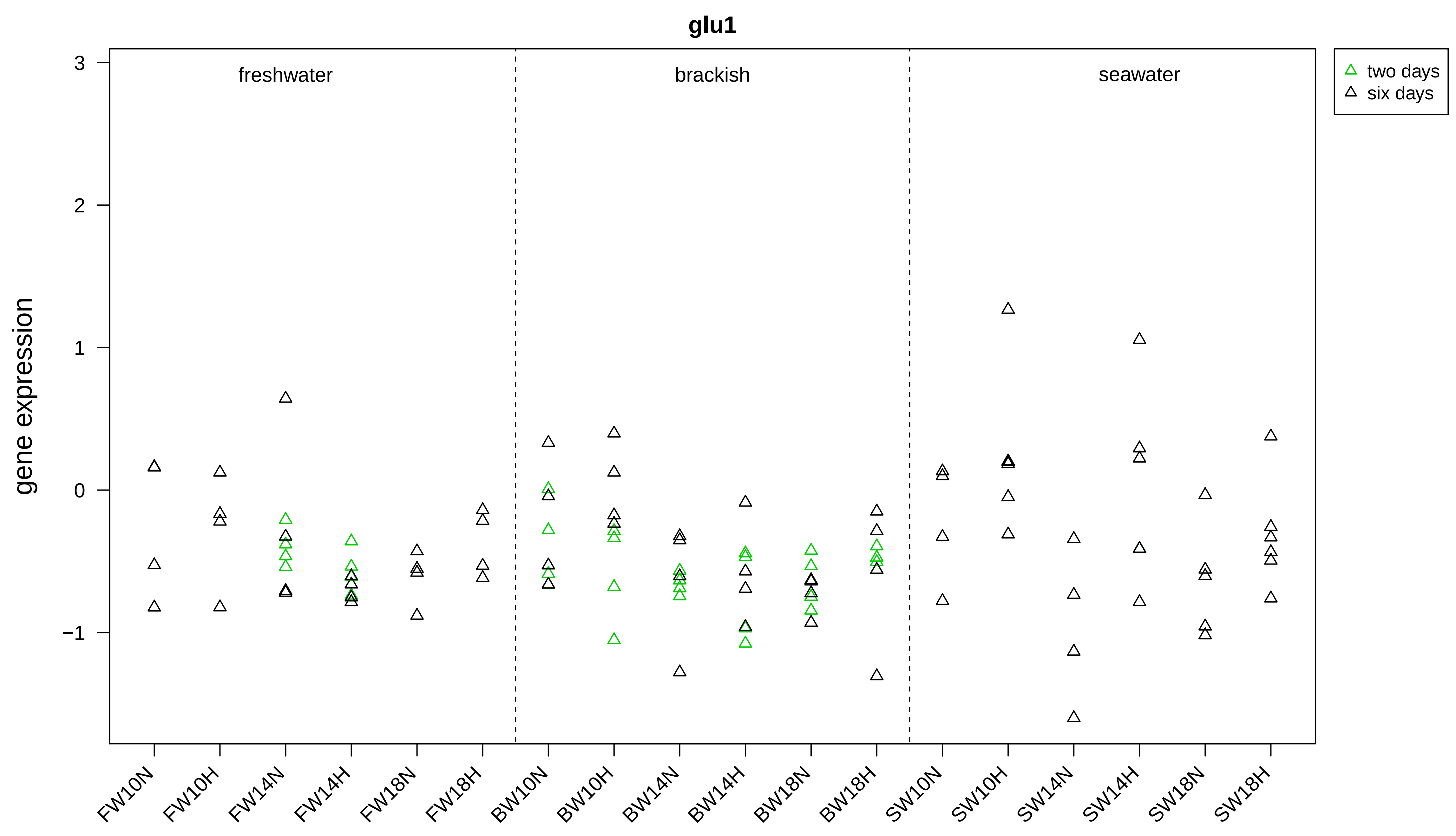

glu1
freshwater
brackish
seawater
two days
six days
gene expression
3
2
1
0
−1
FW10N
FW10H
FW14N
FW14H
FW18N
FW18H
BW10N
BW10H
BW14N
BW14H
BW18N
BW18H
SW10N
SW10H
SW14N
SW14H
SW18N
SW18H

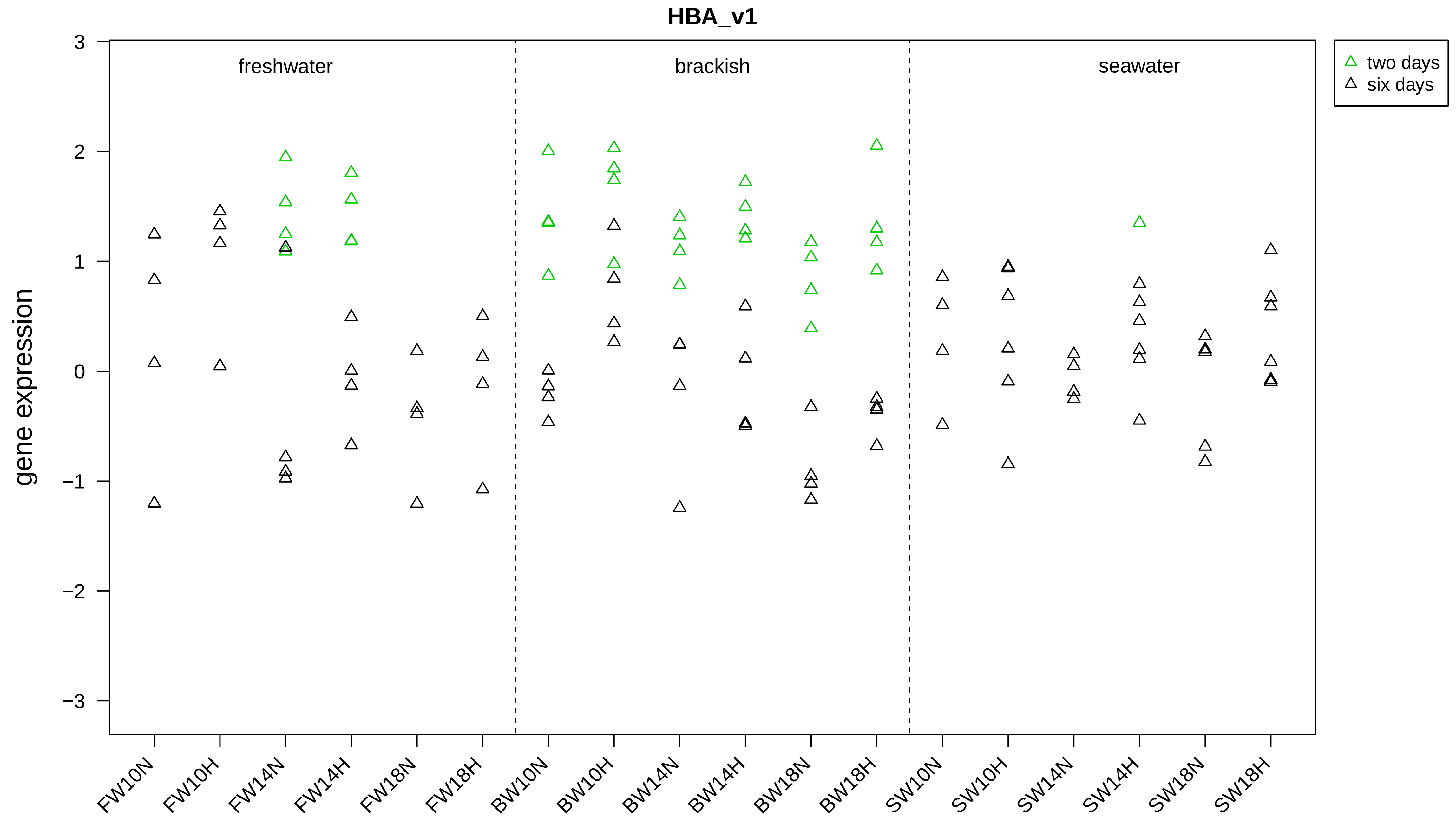

HBA_v1
freshwater
brackish
seawater
two days
six days
gene expression
3
2
1
0
−1
−2
−3
FW10N
FW10H
FW14N
FW14H
FW18N
FW18H
BW10N
BW10H
BW14N
BW14H
BW18N
BW18H
SW10N
SW10H
SW14N
SW14H
SW18N
SW18H

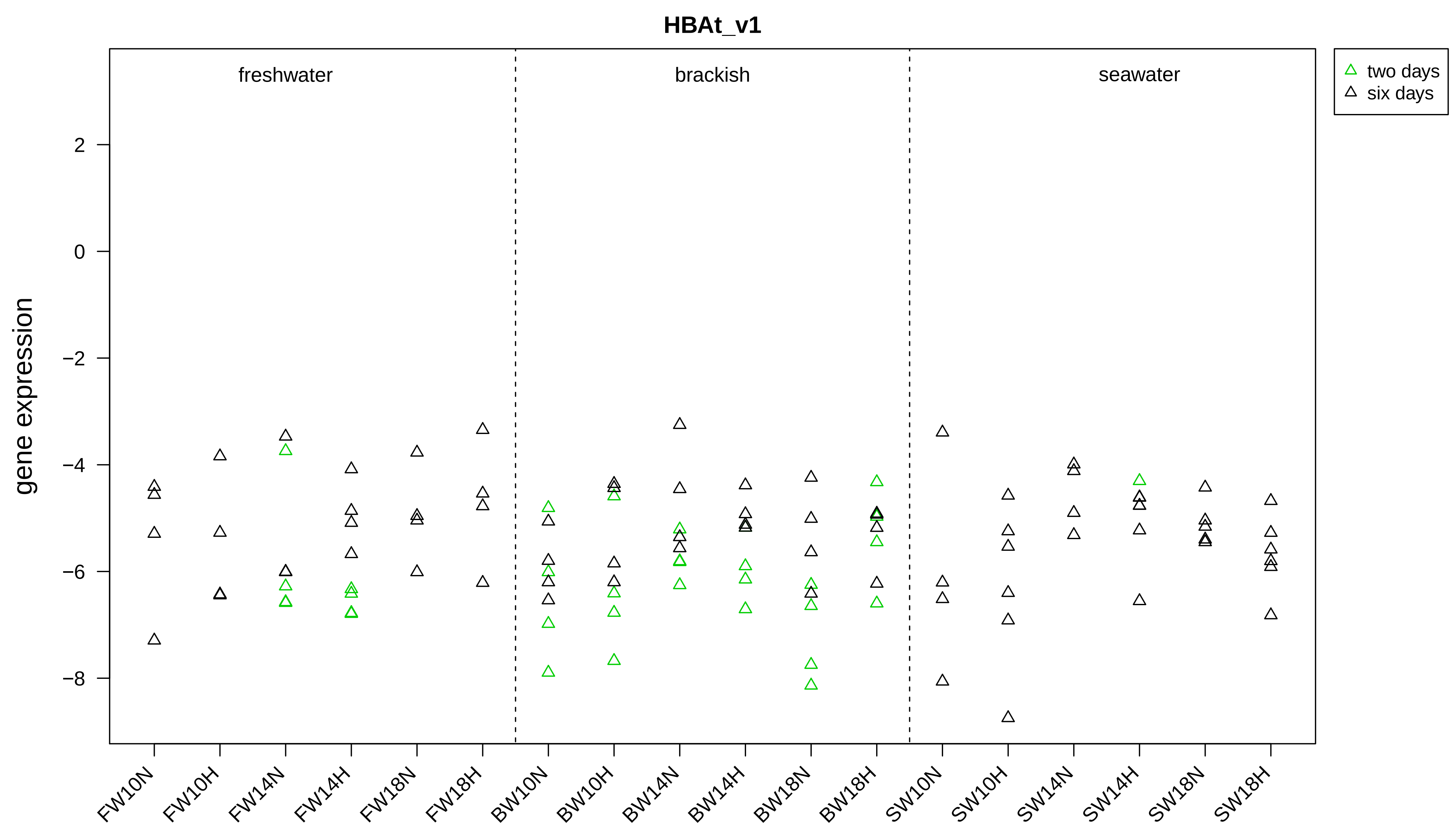

HBAt_v1
freshwater
brackish
seawater
two days
six days
gene expression
2
0
−2
−4
−6
−8
FW10N
FW10H
FW14N
FW14H
FW18N
FW18H
BW10N
BW10H
BW14N
BW14H
BW18N
BW18H
SW10N
SW10H
SW14N
SW14H
SW18N
SW18H

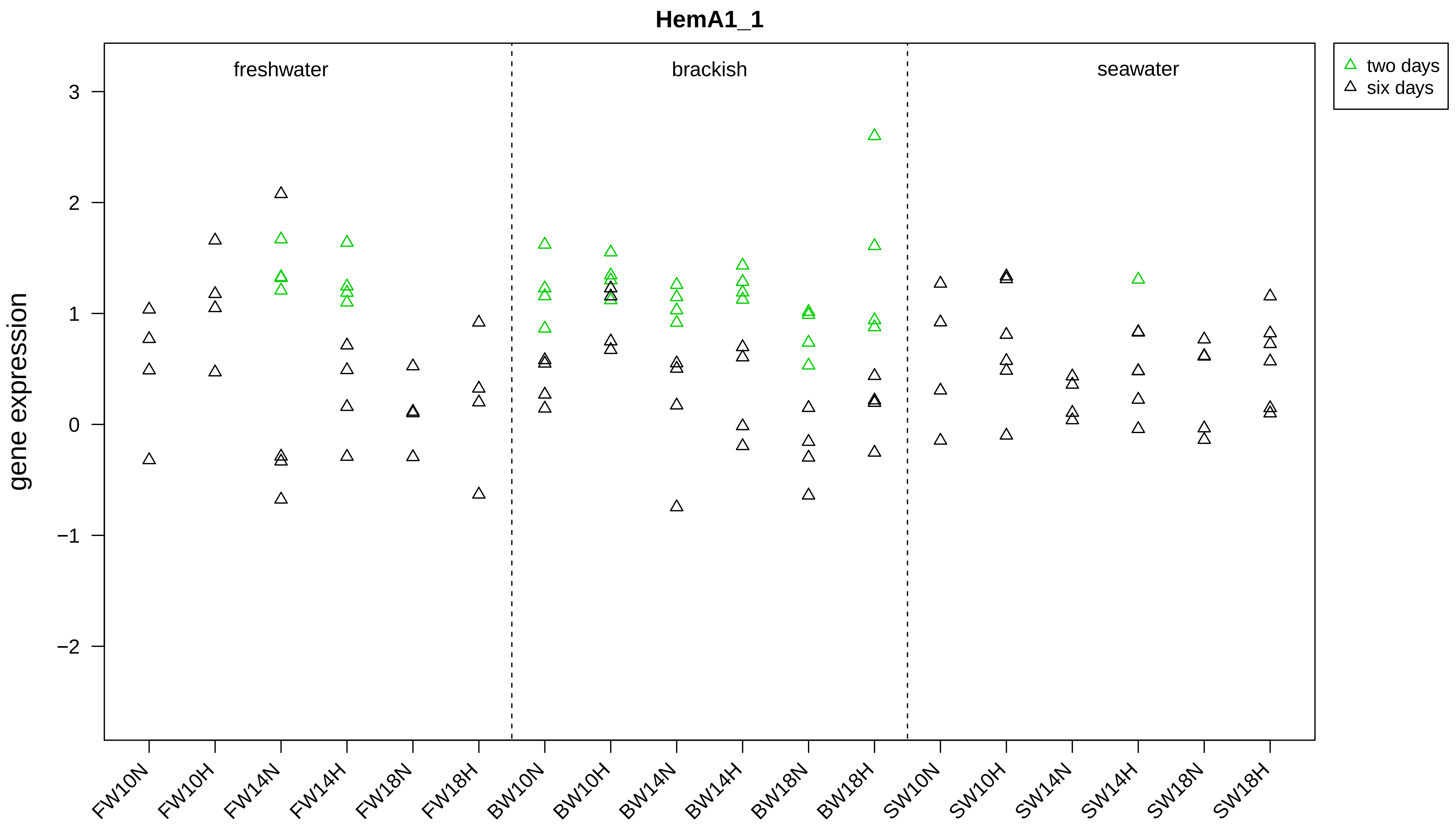
HemA1_1
gene expression
freshwater
brackish
seawater
two days
six days
3
2
1
0
−1
−2
FW10N
FW10H
FW14N
FW14H
FW18N
FW18H
BW10N
BW10H
BW14N
BW14H
BW18N
BW18H
SW10N
SW10H
SW14N
SW14H
SW18N
SW18H

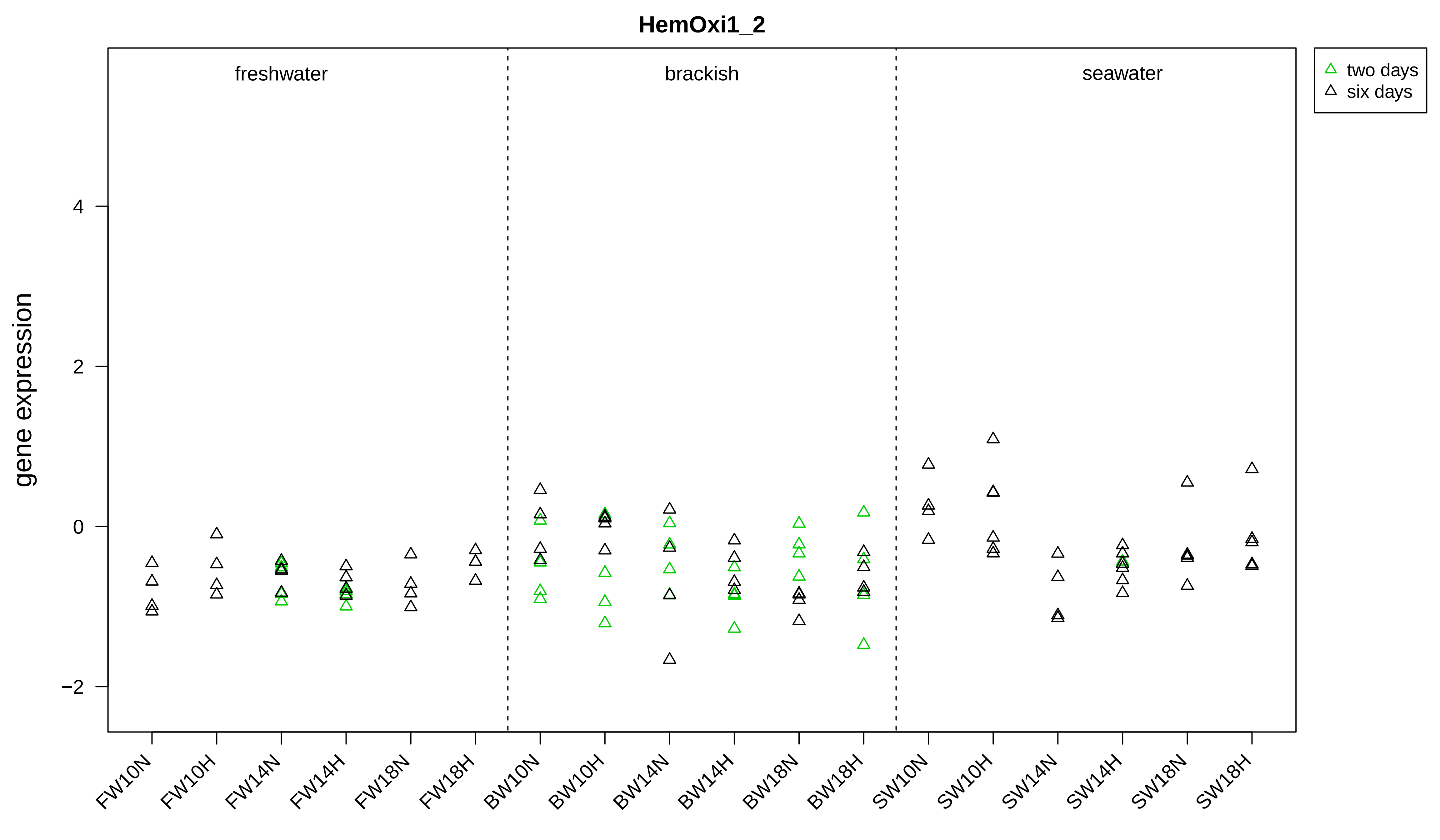

HemOxi1_2
freshwater
brackish
seawater
two days
six days
gene expression
4
2
0
−2
FW10N
FW10H
FW14N
FW14H
FW18N
FW18H
BW10N
BW10H
BW14N
BW14H
BW18N
BW18H
SW10N
SW10H
SW14N
SW14H
SW18N
SW18H

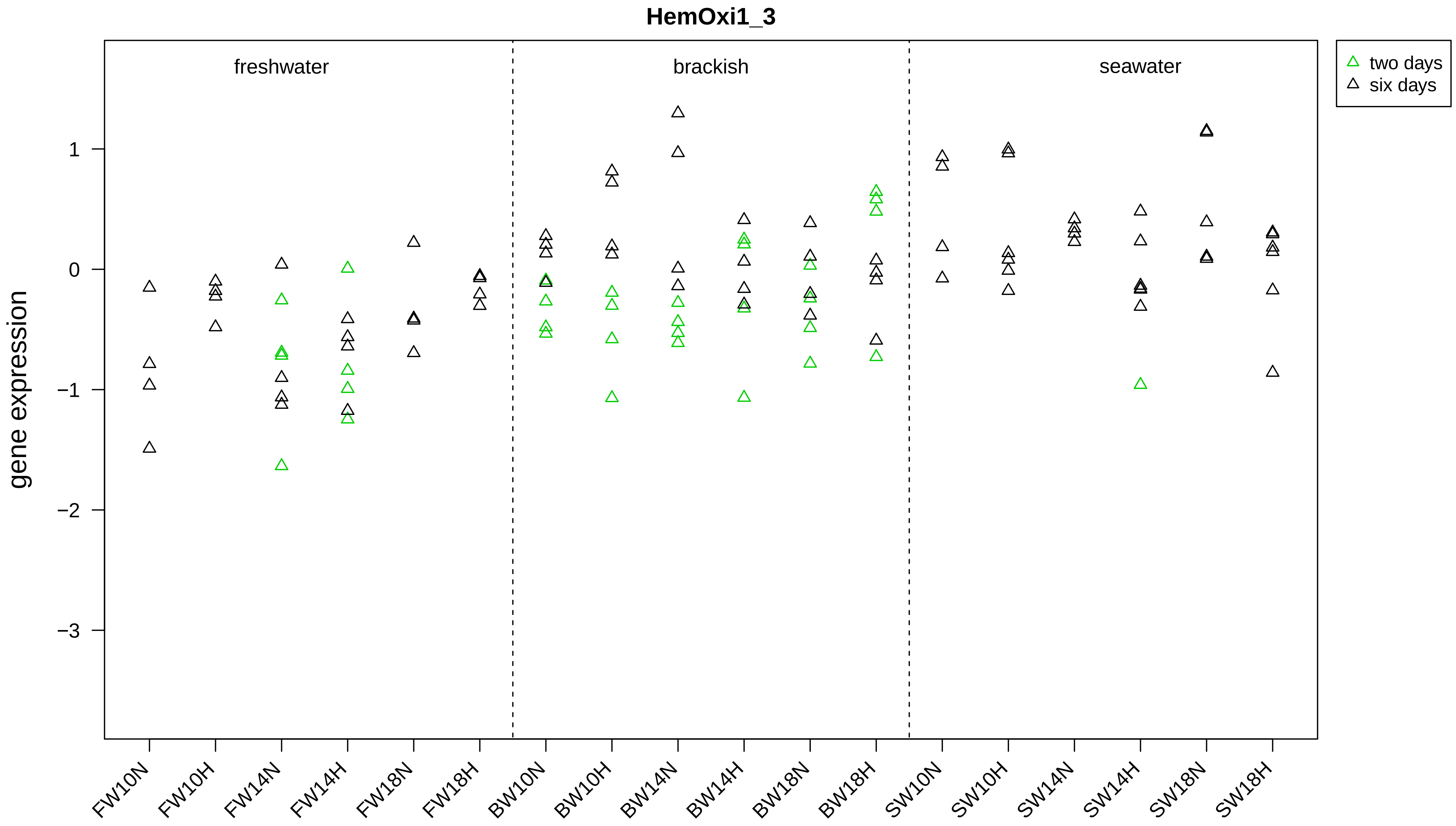

HemOxi1_3
gene expression
freshwater
brackish
seawater
two days
six days
1
0
−1
−2
−3
FW10N
FW10H
FW14N
FW14H
FW18N
FW18H
BW10N
BW10H
BW14N
BW14H
BW18N
BW18H
SW10N
SW10H
SW14N
SW14H
SW18N
SW18H

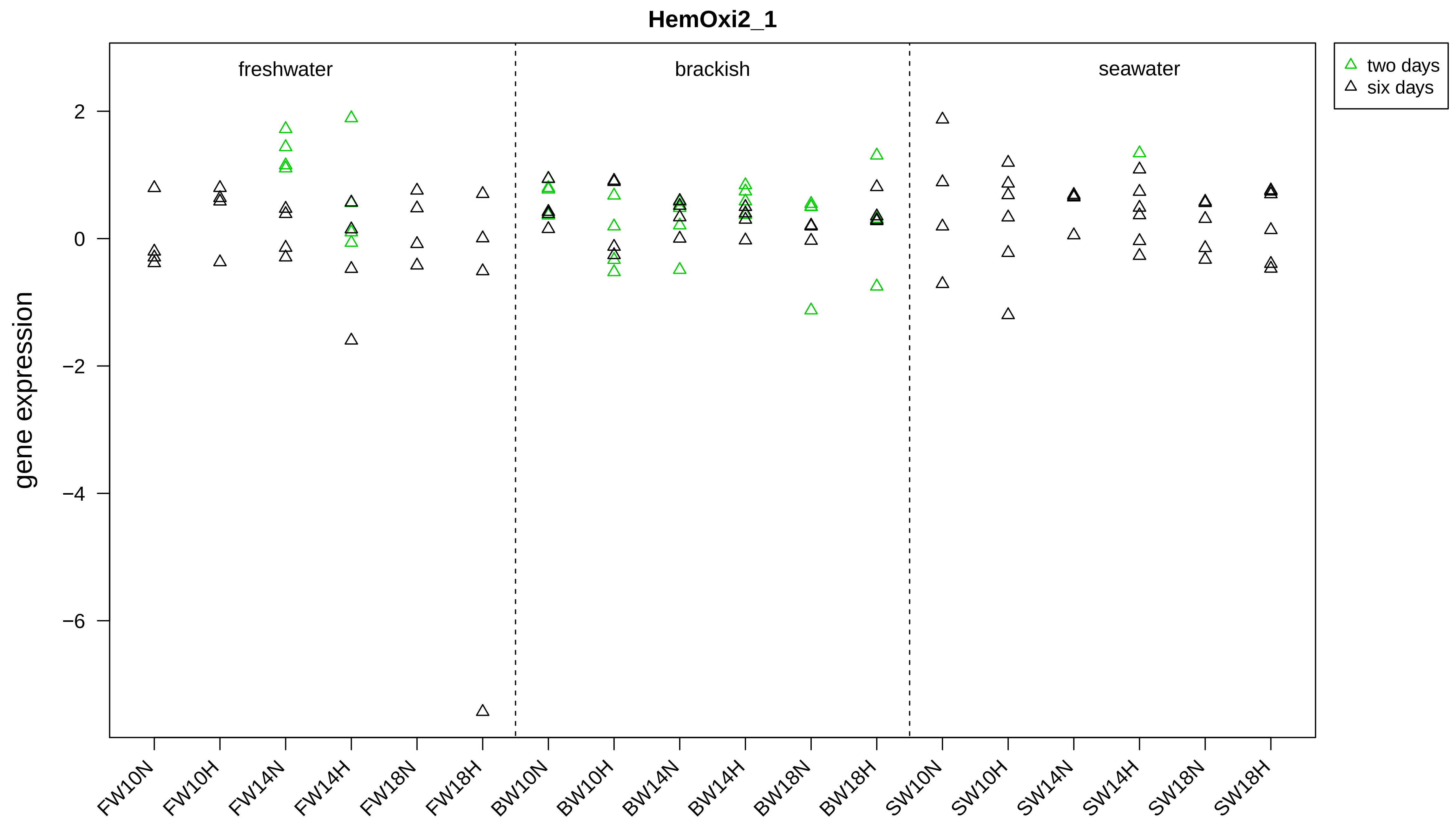

HemOxi2_1
freshwater
brackish
seawater
two days
six days
gene expression
2
0
−2
−4
−6
FW10N
FW10H
FW14N
FW14H
FW18N
FW18H
BW10N
BW10H
BW14N
BW14H
BW18N
BW18H
SW10N
SW10H
SW14N
SW14H
SW18N
SW18H

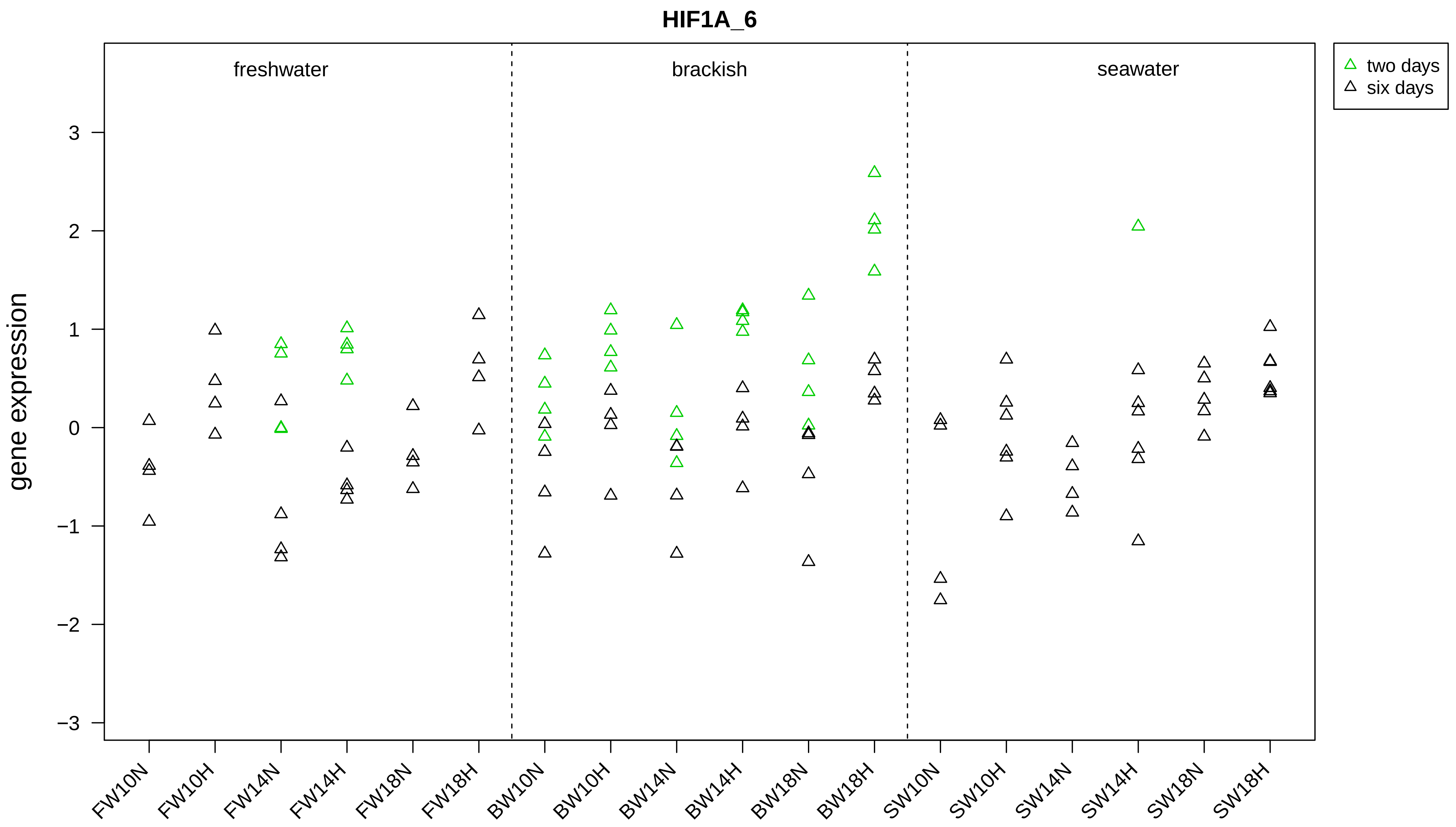
HIF1A_6
freshwater
brackish
seawater
two days
six days
gene expression
3
2
1
0
−1
−2
−3
FW10N
FW10H
FW14N
FW14H
FW18N
FW18H
BW10N
BW10H
BW14N
BW14H
BW18N
BW18H
SW10N
SW10H
SW14N
SW14H
SW18N
SW18H

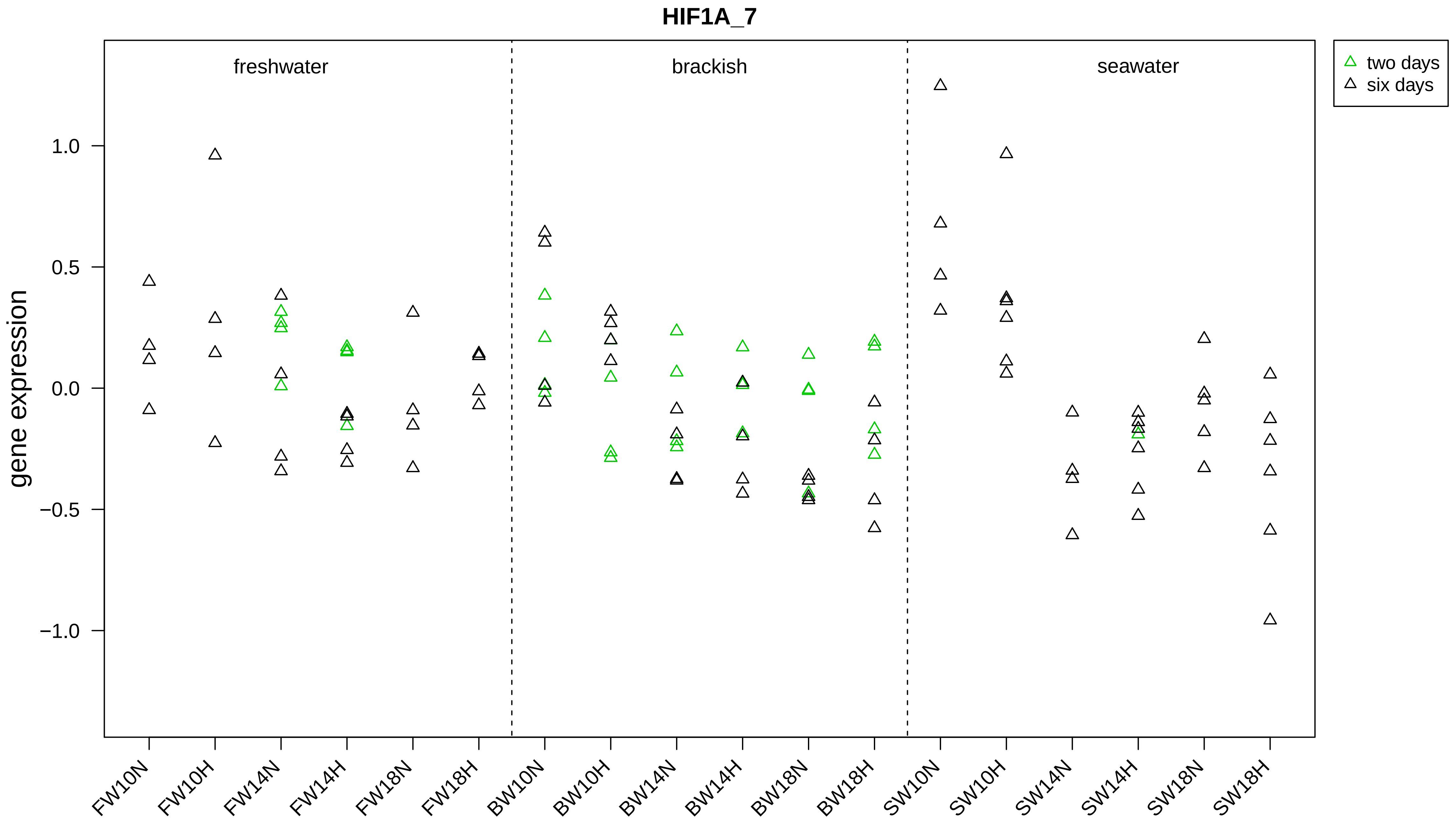

HIF1A_7
freshwater
brackish
seawater
two days
six days
gene expression
1.0
0.5
0.0
−0.5
−1.0
FW10N
FW10H
FW14N
FW14H
FW18N
FW18H
BW10N
BW10H
BW14N
BW14H
BW18N
BW18H
SW10N
SW10H
SW14N
SW14H
SW18N
SW18H

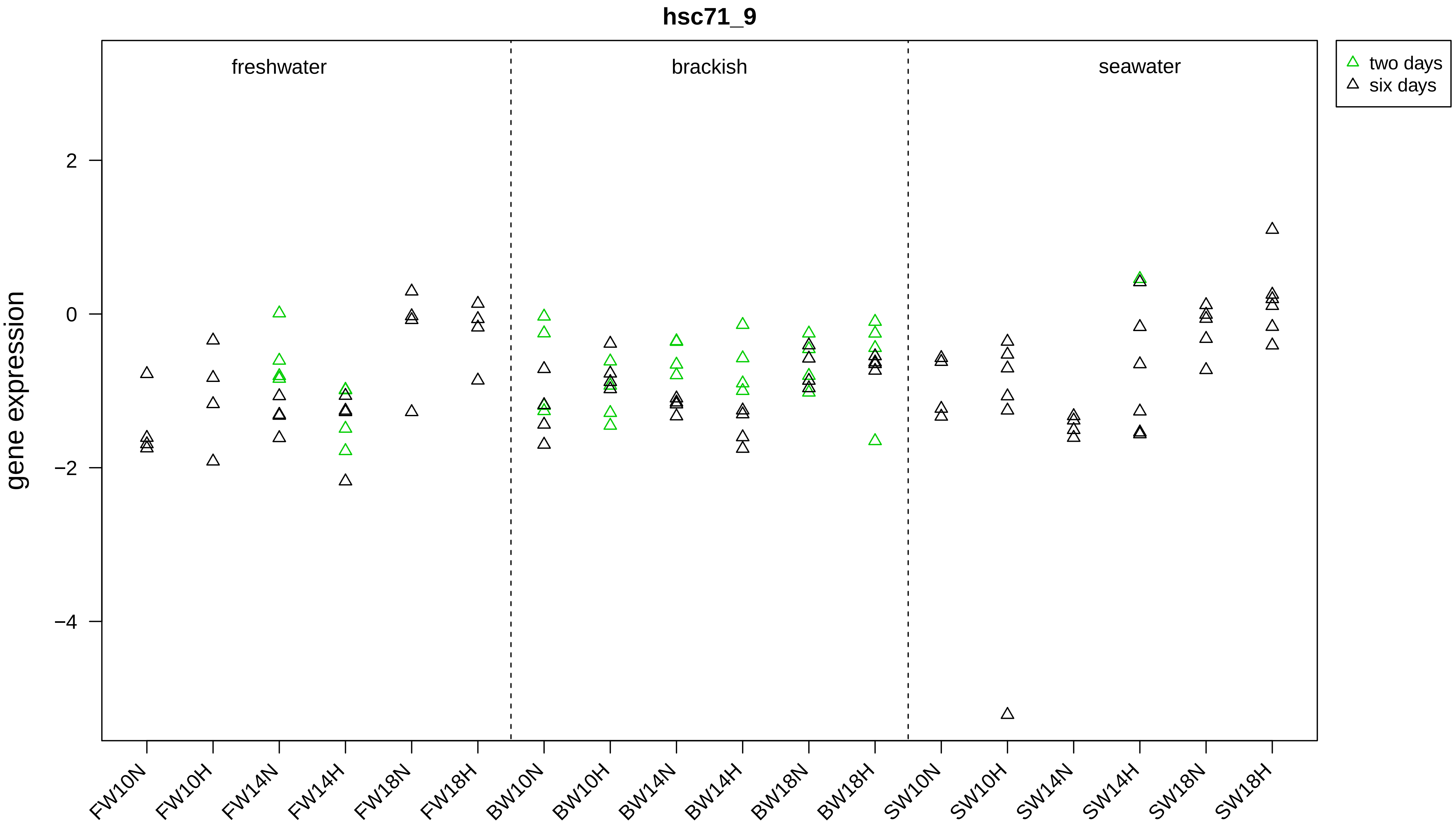

hsc71_9
freshwater
brackish
seawater
two days
six days
gene expression
2
0
−2
−4
FW10N
FW10H
FW14N
FW14H
FW18N
FW18H
BW10N
BW10H
BW14N
BW14H
BW18N
BW18H
SW10N
SW10H
SW14N
SW14H
SW18N
SW18H

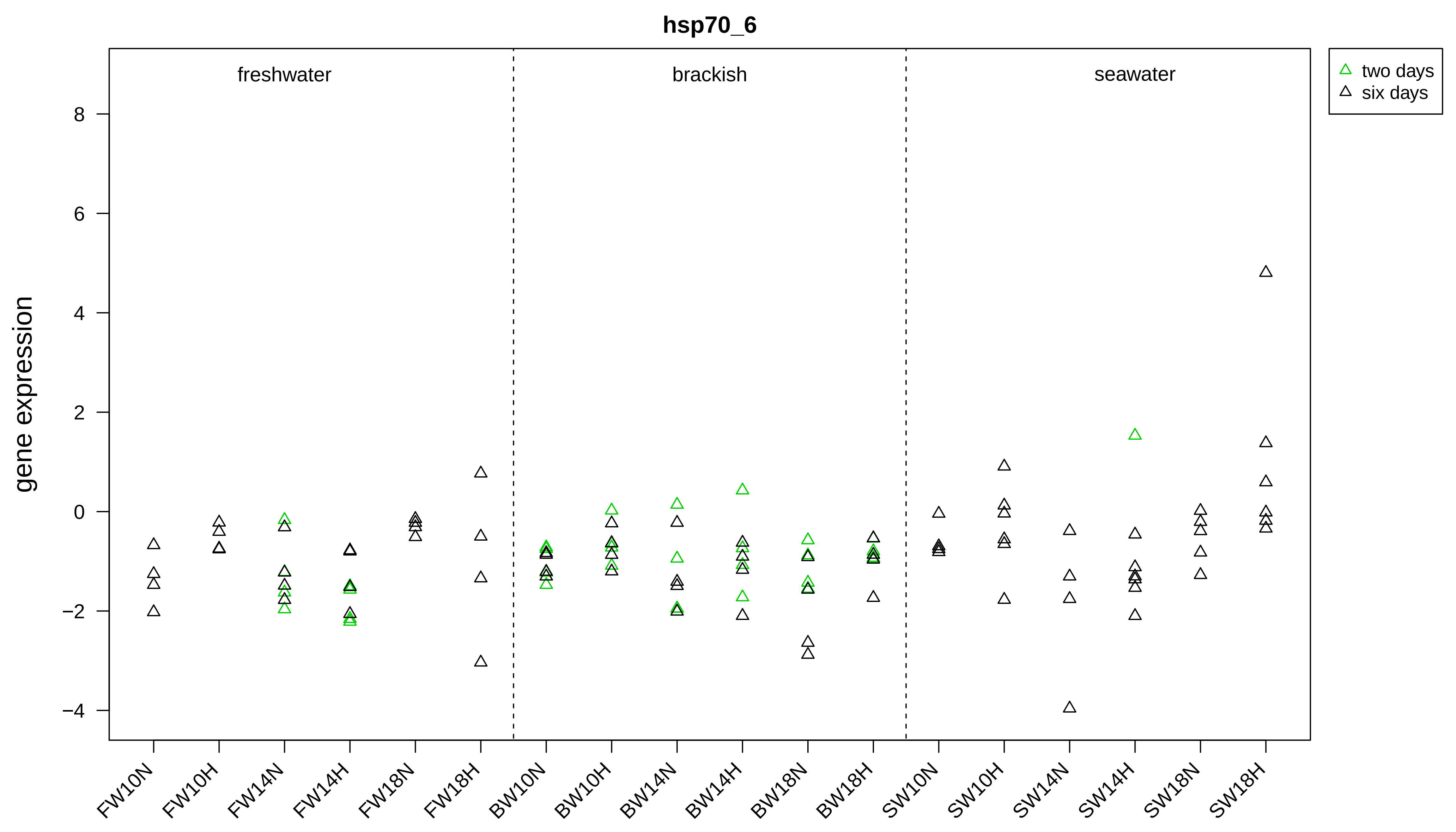

hsp70_6
freshwater
brackish
seawater
two days
six days
gene expression
8
6
4
2
0
−2
−4
FW10N
FW10H
FW14N
FW14H
FW18N
FW18H
BW10N
BW10H
BW14N
BW14H
BW18N
BW18H
SW10N
SW10H
SW14N
SW14H
SW18N
SW18H

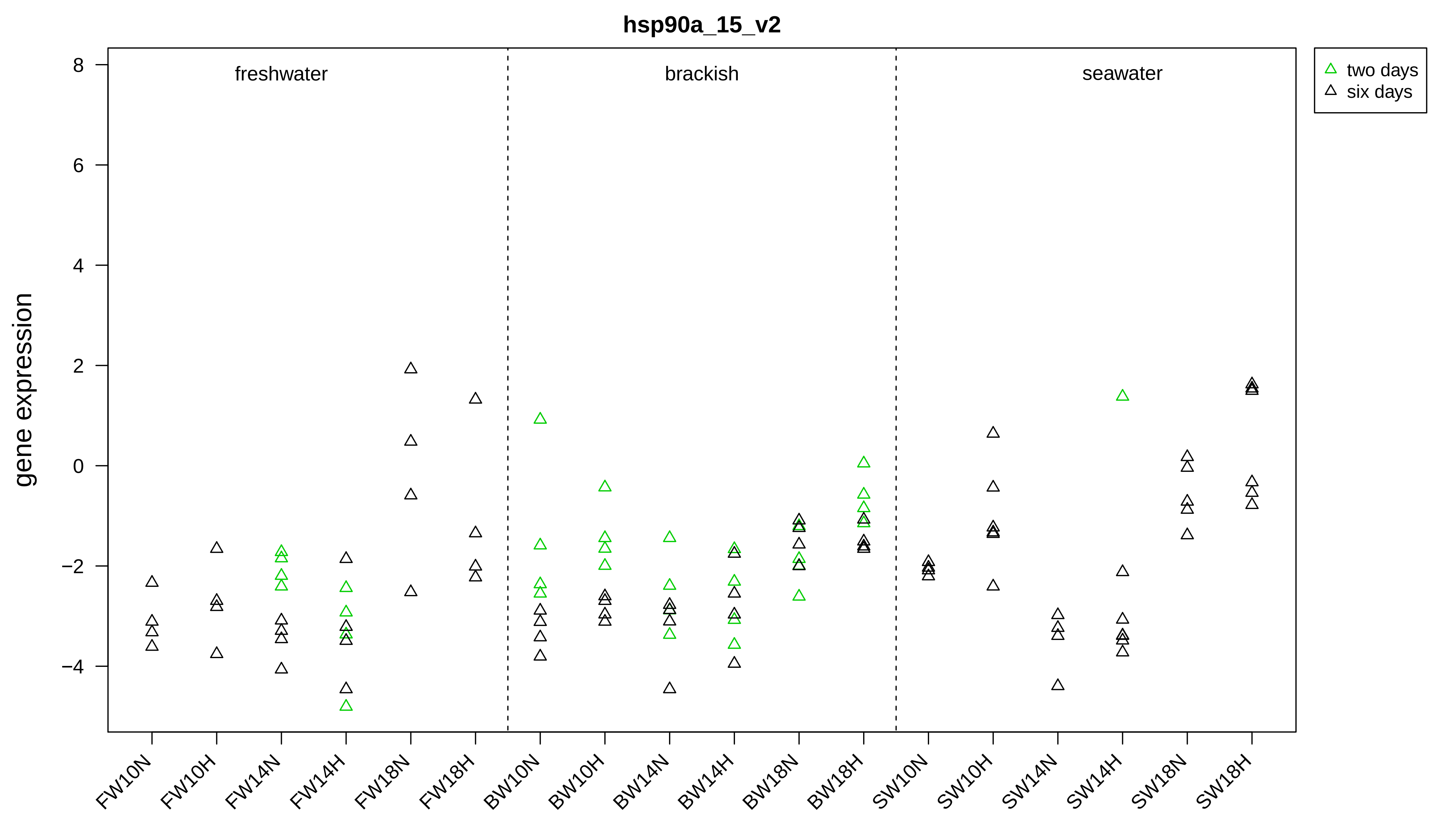
hsp90a_15_v2
gene expression
freshwater
brackish
seawater
two days
six days
8
6
4
2
0
−2
−4
FW10N
FW10H
FW14N
FW14H
FW18N
FW18H
BW10N
BW10H
BW14N
BW14H
BW18N
BW18H
SW10N
SW10H
SW14N
SW14H
SW18N
SW18H

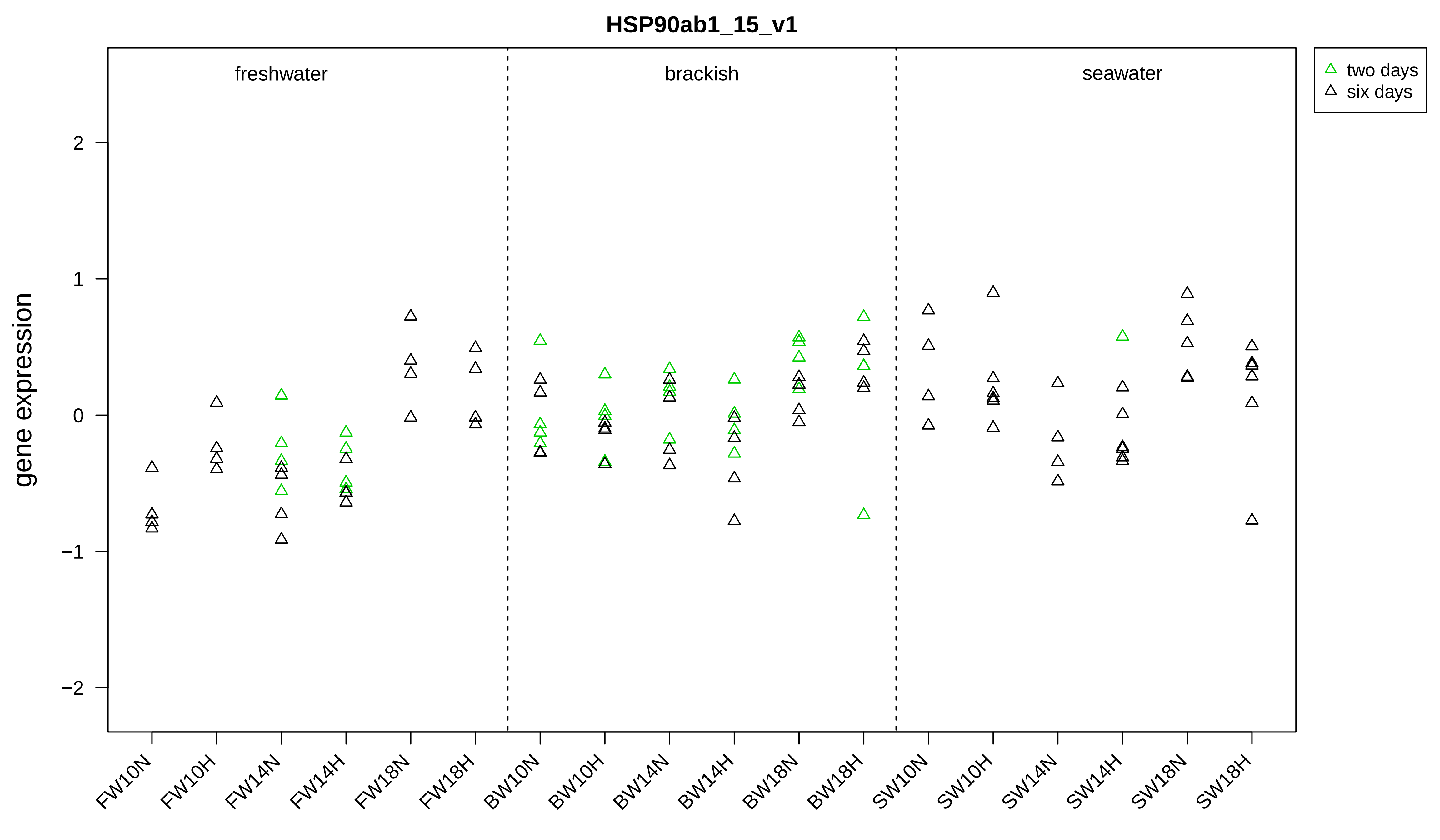
HSP90ab1_15_v1
freshwater
brackish
seawater
two days
six days
gene expression
2
1
0
−1
−2
FW10N
FW10H
FW14N
FW14H
FW18N
FW18H
BW10N
BW10H
BW14N
BW14H
BW18N
BW18H
SW10N
SW10H
SW14N
SW14H
SW18N
SW18H

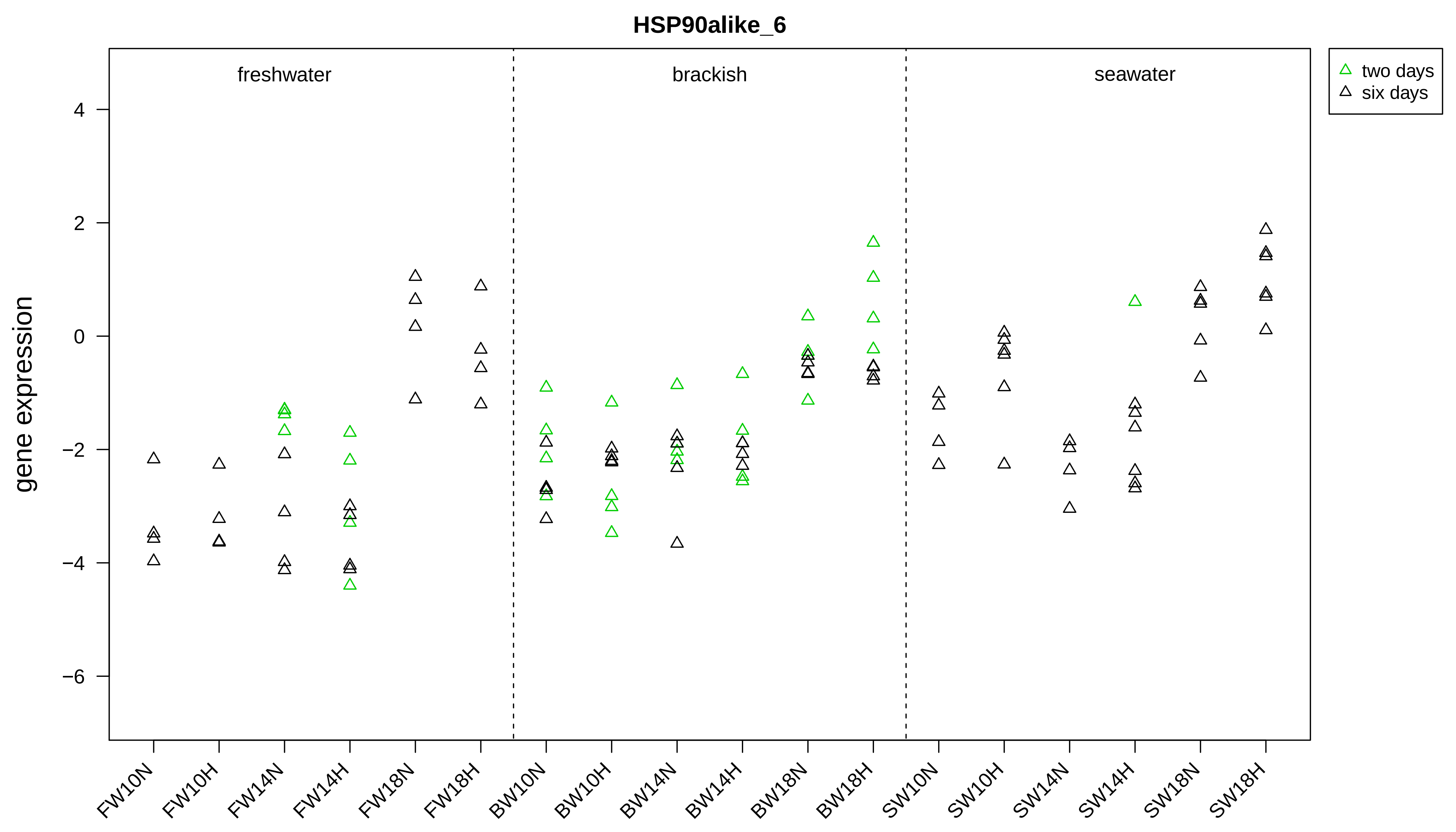

HSP90alike_6
freshwater
brackish
seawater
two days
six days
gene expression
4
2
0
−2
−4
−6
FW10N
FW10H
FW14N
FW14H
FW18N
FW18H
BW10N
BW10H
BW14N
BW14H
BW18N
BW18H
SW10N
SW10H
SW14N
SW14H
SW18N
SW18H

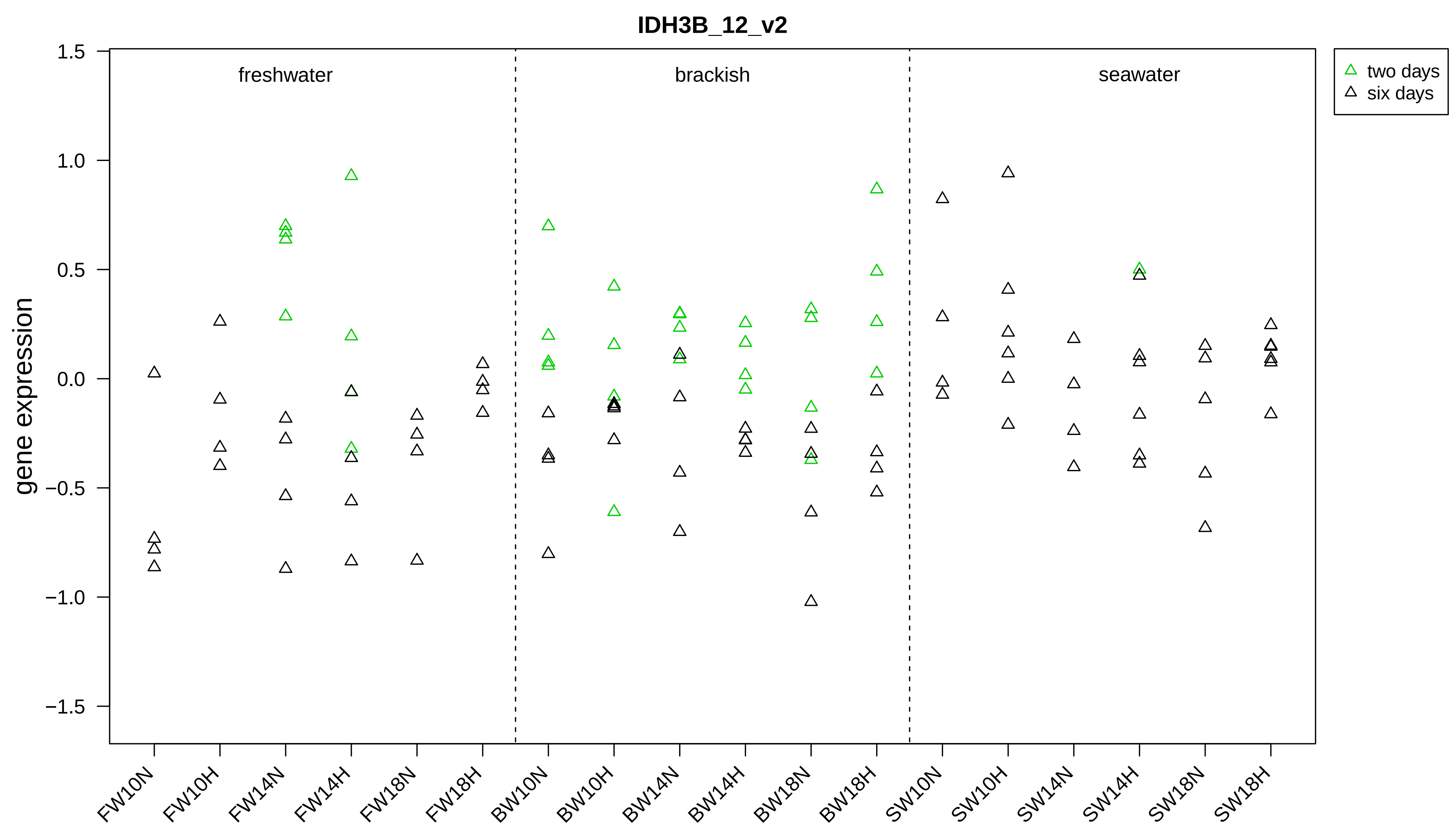

IDH3B_12_v2
gene expression
1.5
1.0
0.5
0.0
−0.5
−1.0
−1.5
freshwater
brackish
seawater
two days
six days
FW10N
FW10H
FW14N
FW14H
FW18N
FW18H
BW10N
BW10H
BW14N
BW14H
BW18N
BW18H
SW10N
SW10H
SW14N
SW14H
SW18N
SW18H

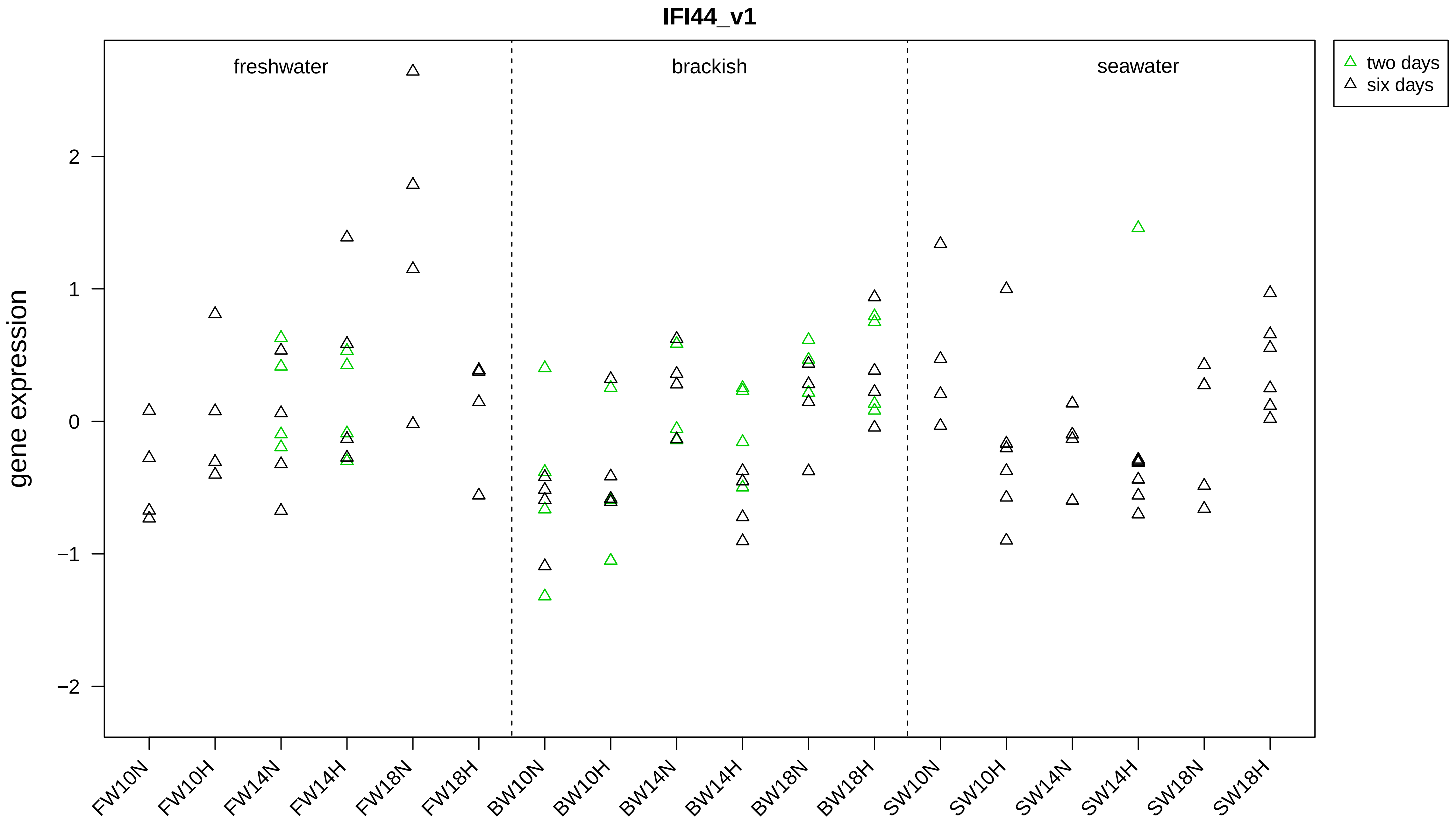

IFI44_v1
two days
six days
freshwater
brackish
seawater
gene expression
2
1
0
−1
−2
FW10N
FW10H
FW14N
FW14H
FW18N
FW18H
BW10N
BW10H
BW14N
BW14H
BW18N
BW18H
SW10N
SW10H
SW14N
SW14H
SW18N
SW18H

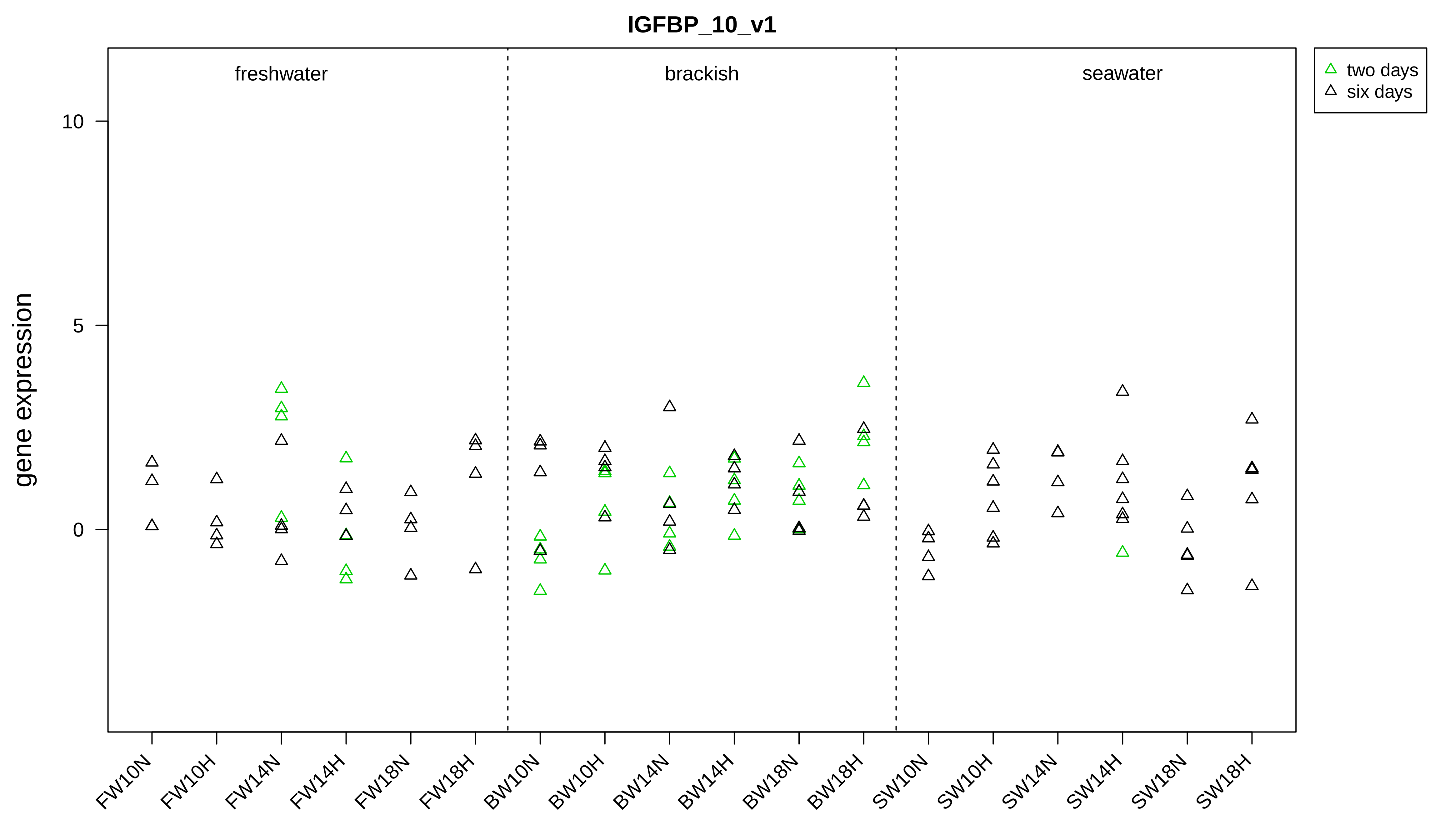
IGFBP_10_v1
gene expression
0
5
10
freshwater
brackish
seawater
two days
six days
FW10N
FW10H
FW14N
FW14H
FW18N
FW18H
BW10N
BW10H
BW14N
BW14H
BW18N
BW18H
SW10N
SW10H
SW14N
SW14H
SW18N
SW18H

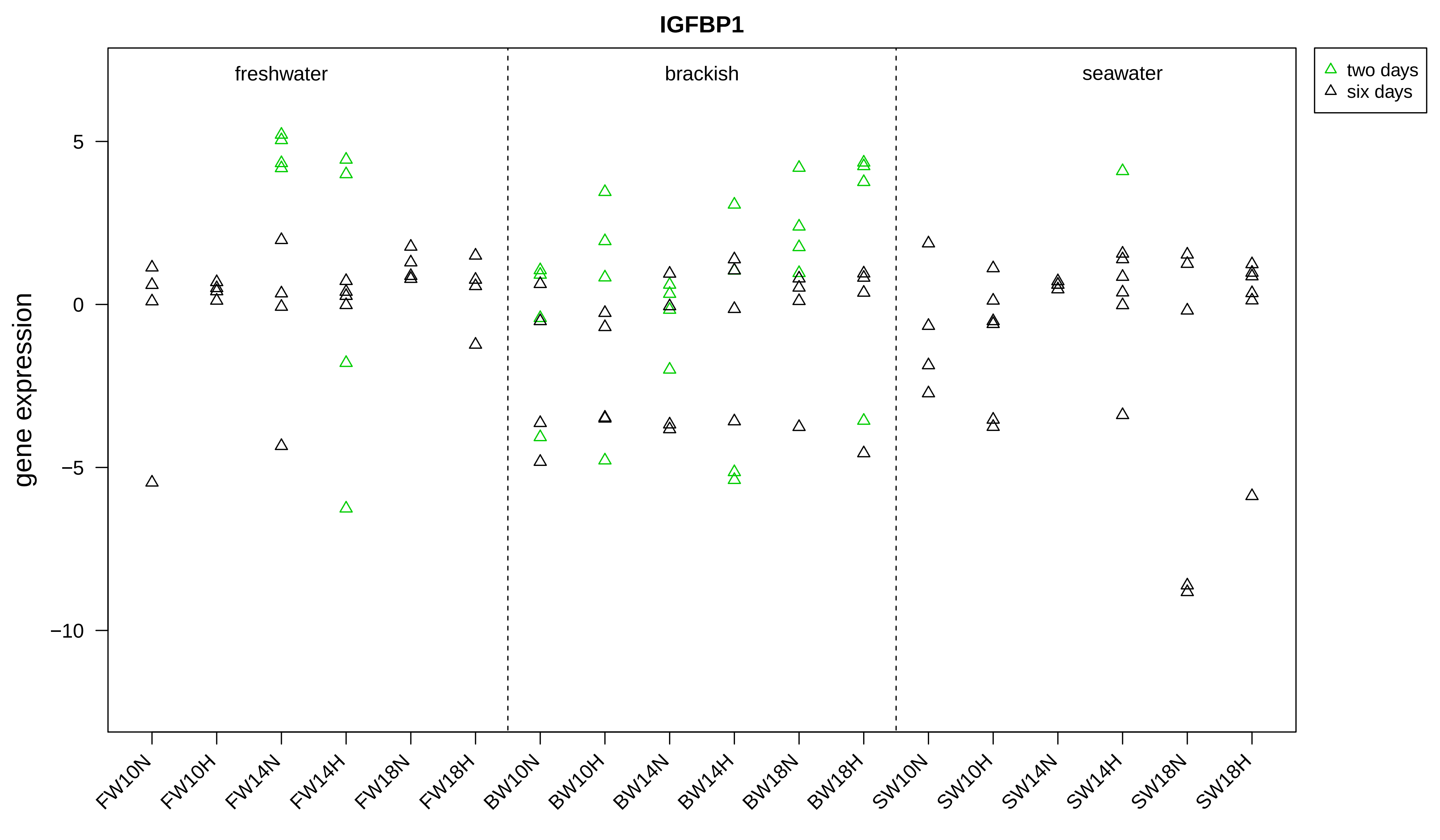
IGFBP1
gene expression
5
0
−5
−10
freshwater
brackish
seawater
two days
six days
FW10N
FW10H
FW14N
FW14H
FW18N
FW18H
BW10N
BW10H
BW14N
BW14H
BW18N
BW18H
SW10N
SW10H
SW14N
SW14H
SW18N
SW18H

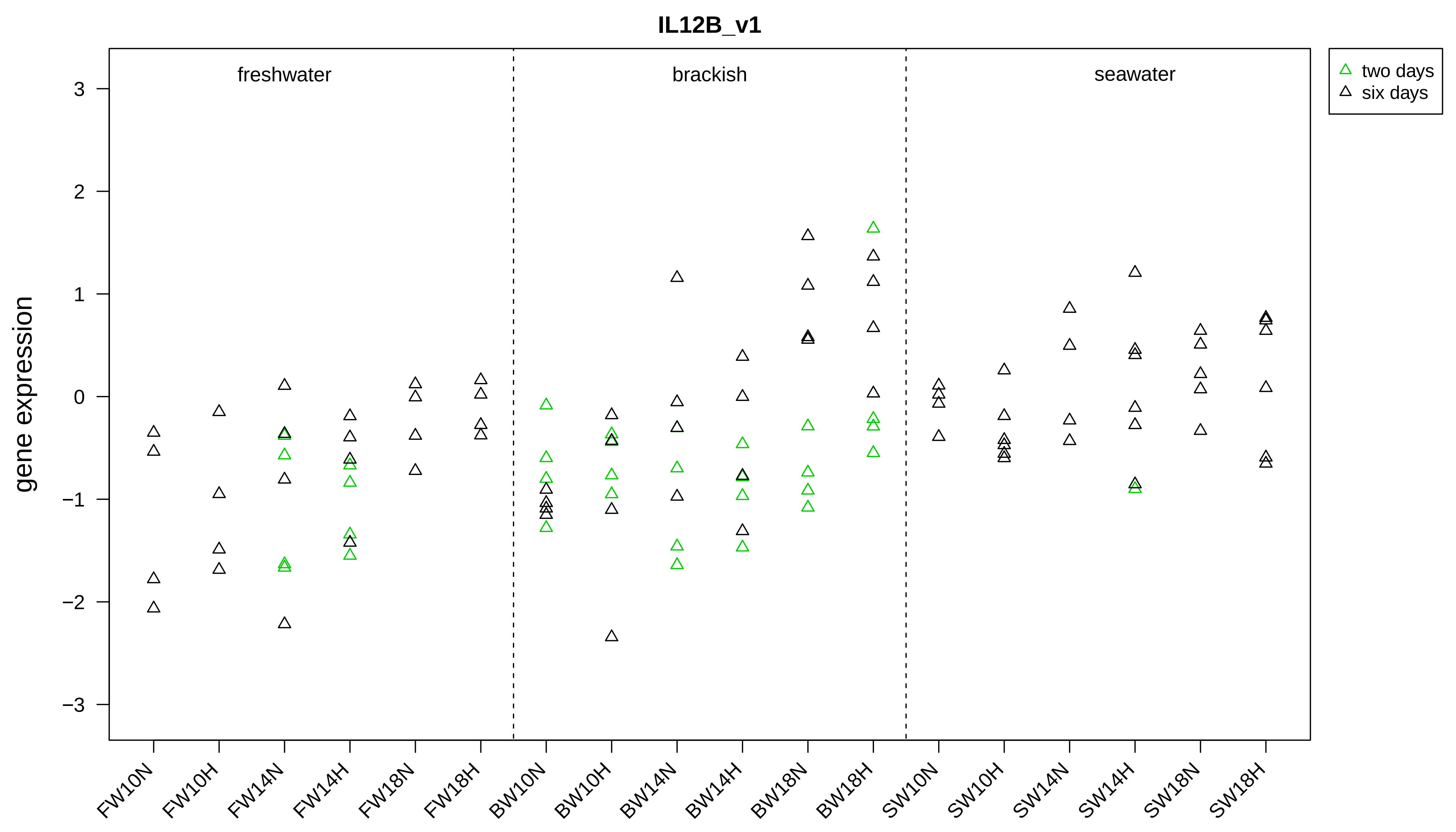
IL12B_v1
gene expression
3
2
1
0
−1
−2
−3
freshwater
brackish
seawater
two days
six days
FW10N
FW10H
FW14N
FW14H
FW18N
FW18H
BW10N
BW10H
BW14N
BW14H
BW18N
BW18H
SW10N
SW10H
SW14N
SW14H
SW18N
SW18H

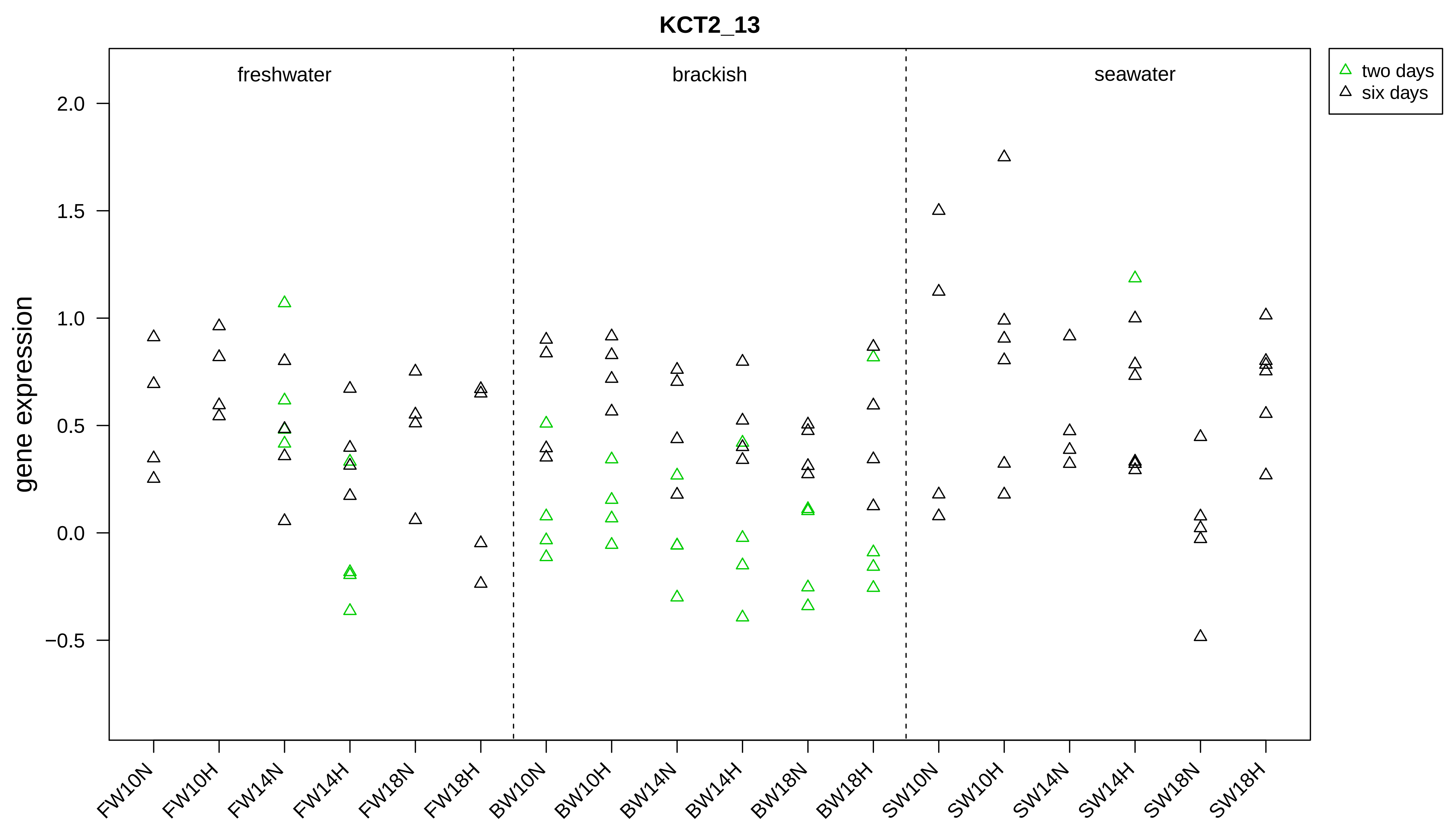

KCT2_13
freshwater
brackish
seawater
two days
six days
gene expression
2.0
1.5
1.0
0.5
0.0
−0.5
FW10N
FW10H
FW14N
FW14H
FW18N
FW18H
BW10N
BW10H
BW14N
BW14H
BW18N
BW18H
SW10N
SW10H
SW14N
SW14H
SW18N
SW18H

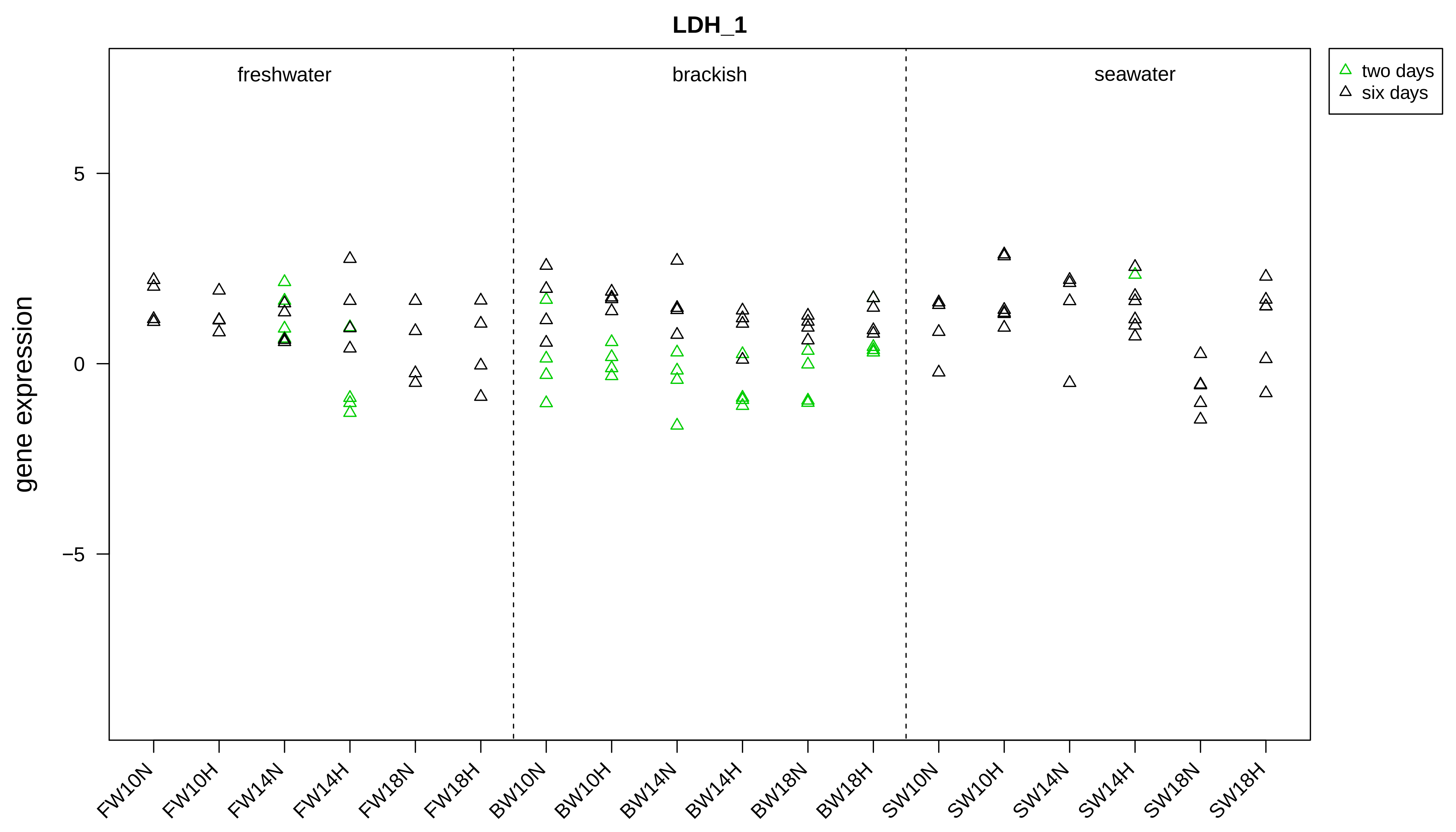

LDH_1
freshwater
brackish
seawater
two days
six days
gene expression
5
0
−5
FW10N
FW10H
FW14N
FW14H
FW18N
FW18H
BW10N
BW10H
BW14N
BW14H
BW18N
BW18H
SW10N
SW10H
SW14N
SW14H
SW18N
SW18H

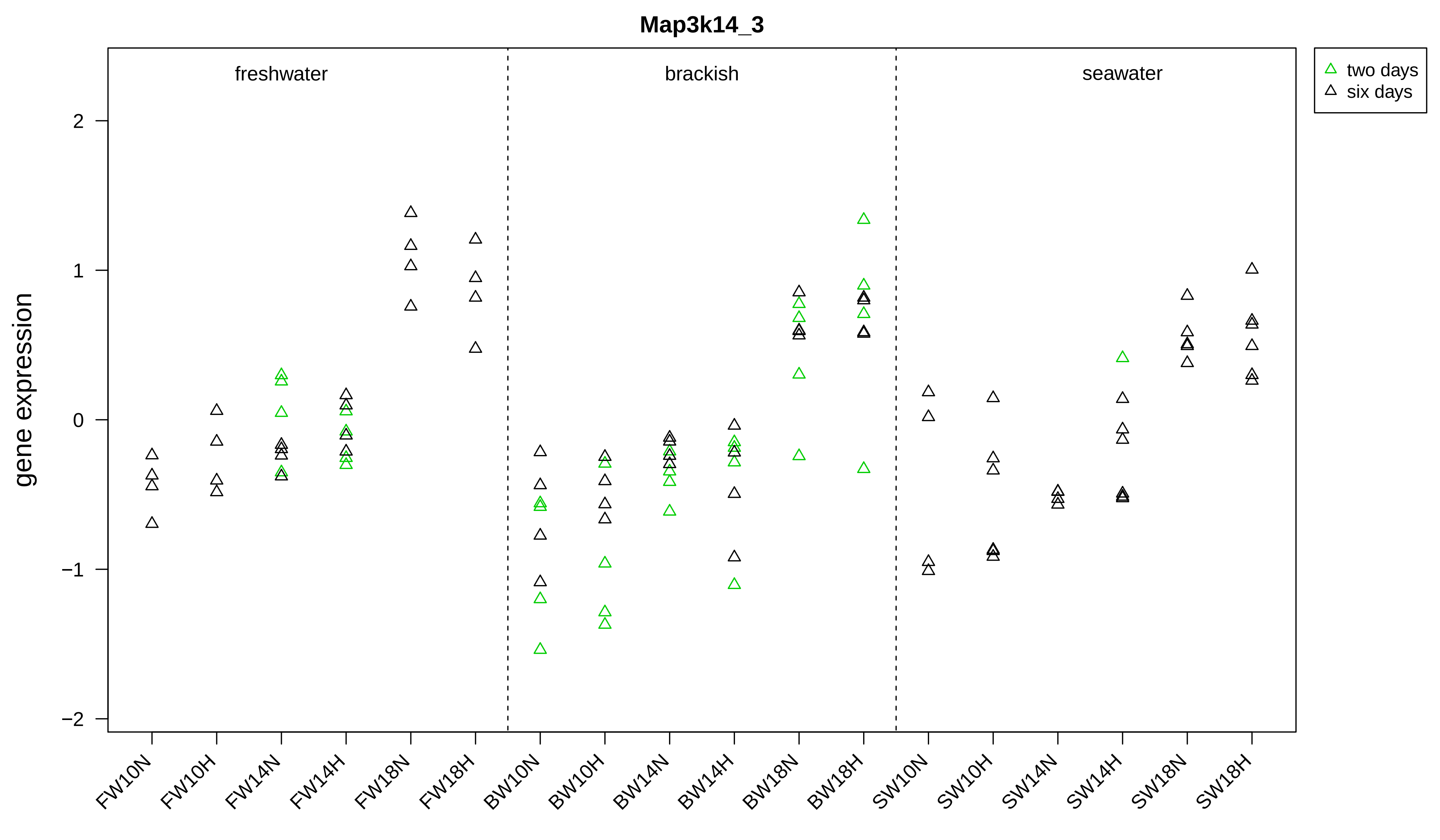
Map3k14_3
freshwater
brackish
seawater
two days
six days
gene expression
2
1
0
−1
−2
FW10N
FW10H
FW14N
FW14H
FW18N
FW18H
BW10N
BW10H
BW14N
BW14H
BW18N
BW18H
SW10N
SW10H
SW14N
SW14H
SW18N
SW18H

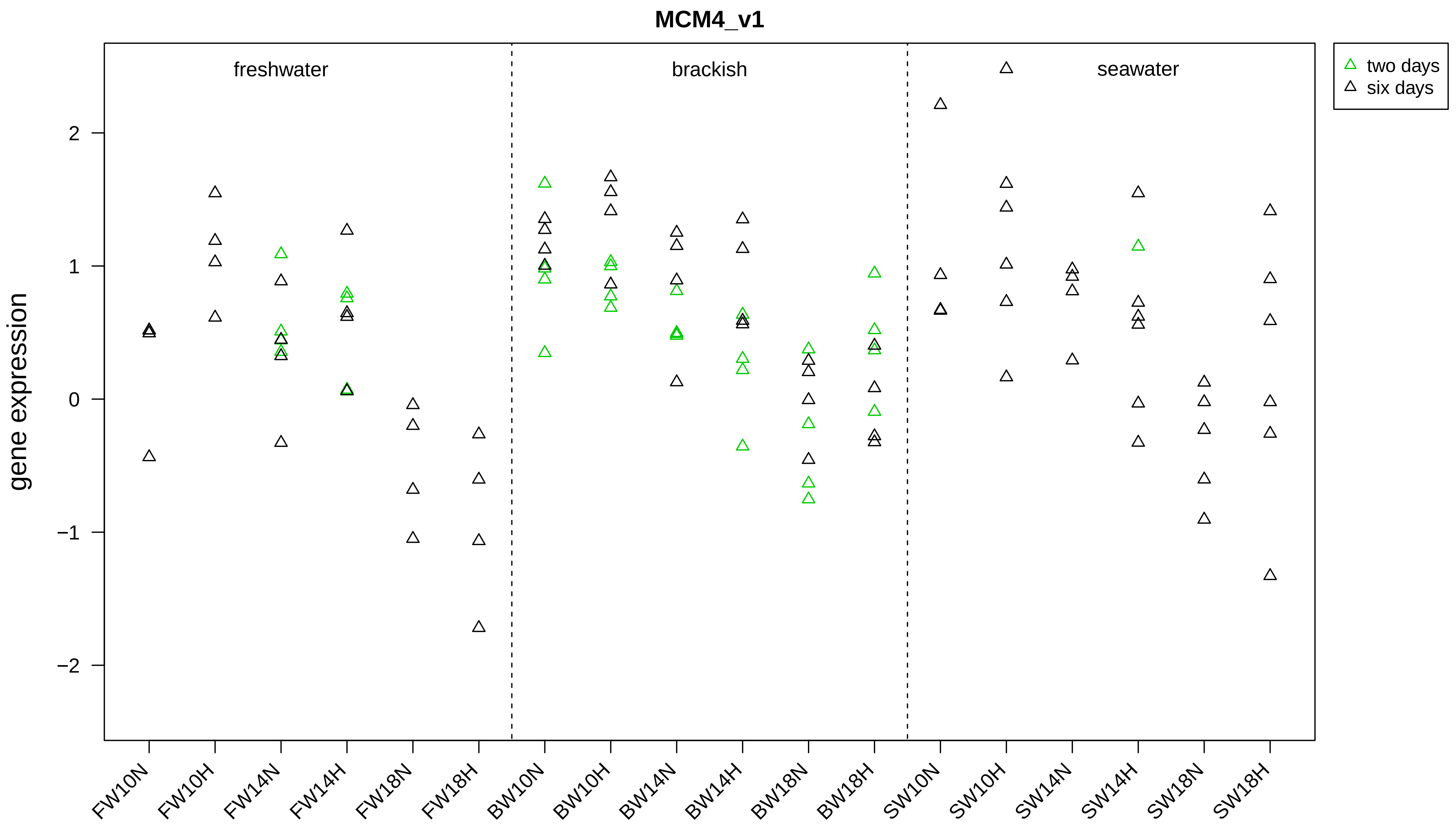

MCM4_v1
freshwater
brackish
seawater
two days
six days
gene expression
2
1
0
−1
−2
FW10N
FW10H
FW14N
FW14H
FW18N
FW18H
BW10N
BW10H
BW14N
BW14H
BW18N
BW18H
SW10N
SW10H
SW14N
SW14H
SW18N
SW18H

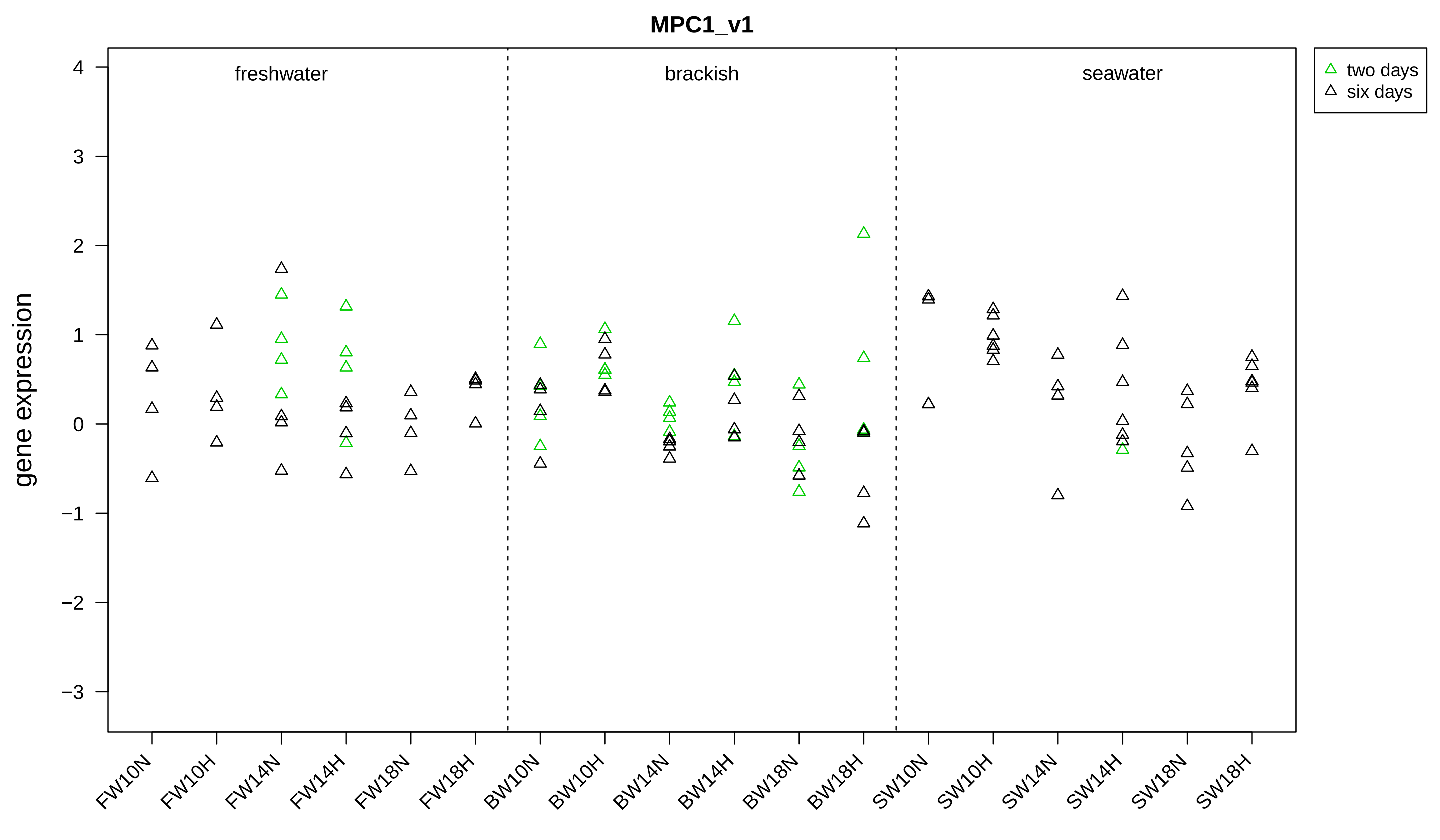
MPC1_v1
gene expression
4
3
2
1
0
−1
−2
−3
freshwater
brackish
seawater
two days
six days
FW10N
FW10H
FW14N
FW14H
FW18N
FW18H
BW10N
BW10H
BW14N
BW14H
BW18N
BW18H
SW10N
SW10H
SW14N
SW14H
SW18N
SW18H

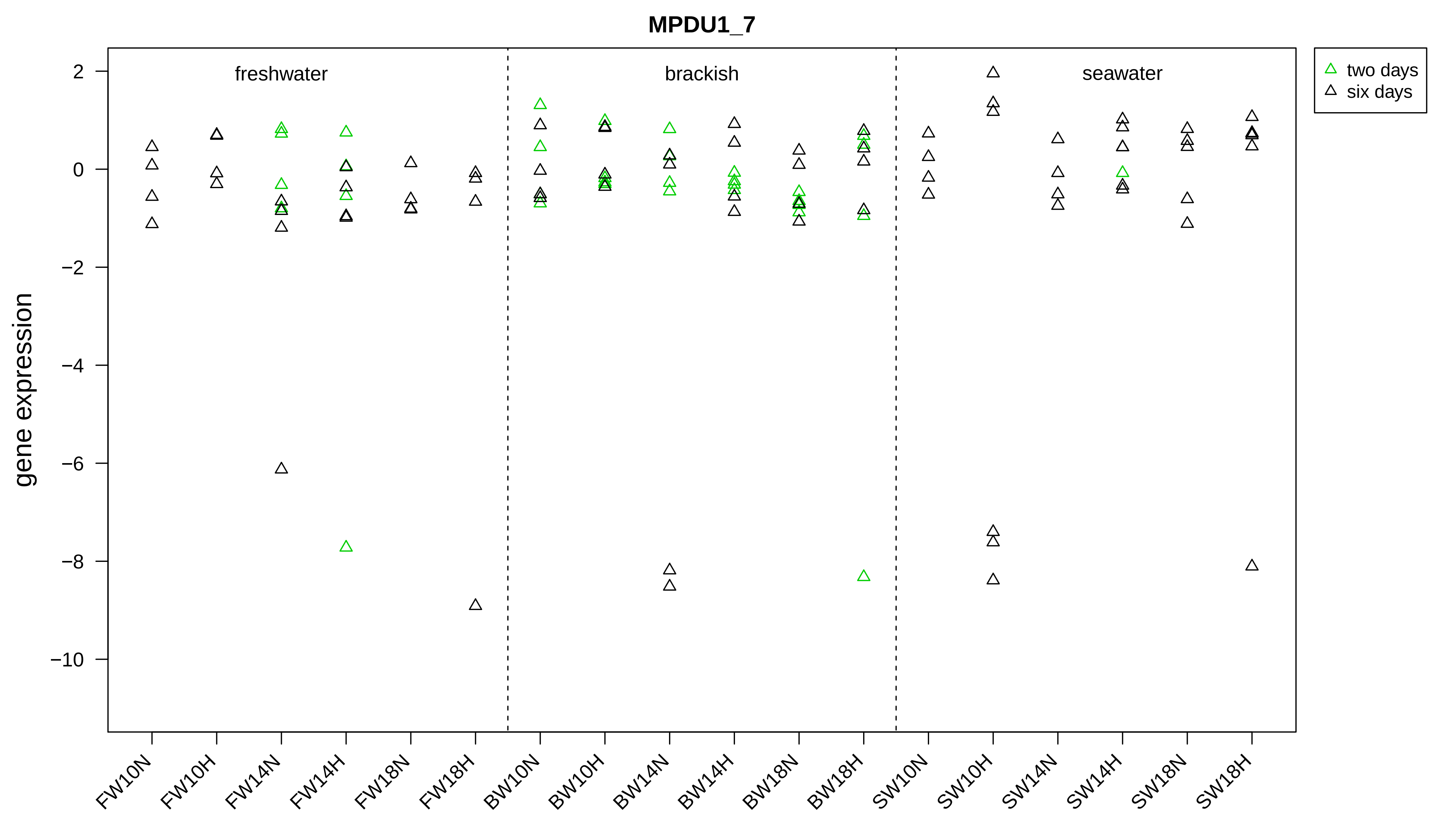

MPDU1_7
freshwater
brackish
seawater
two days
six days
gene expression
2
0
−2
−4
−6
−8
−10
FW10N
FW10H
FW14N
FW14H
FW18N
FW18H
BW10N
BW10H
BW14N
BW14H
BW18N
BW18H
SW10N
SW10H
SW14N
SW14H
SW18N
SW18H

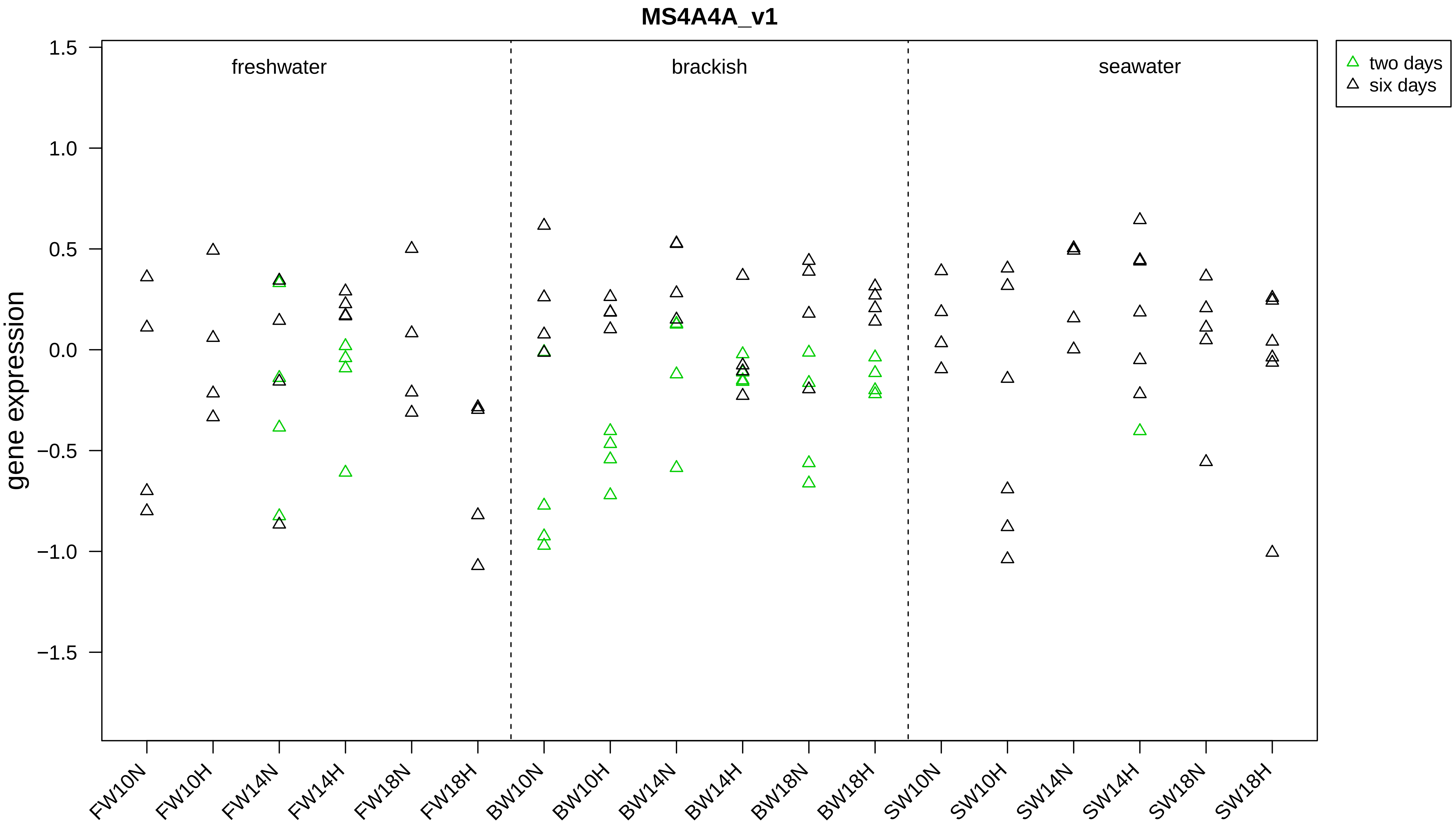
MS4A4A_v1
gene expression
1.5
1.0
0.5
0.0
−0.5
−1.0
−1.5
freshwater
brackish
seawater
two days
six days
FW10N
FW10H
FW14N
FW14H
FW18N
FW18H
BW10N
BW10H
BW14N
BW14H
BW18N
BW18H
SW10N
SW10H
SW14N
SW14H
SW18N
SW18H

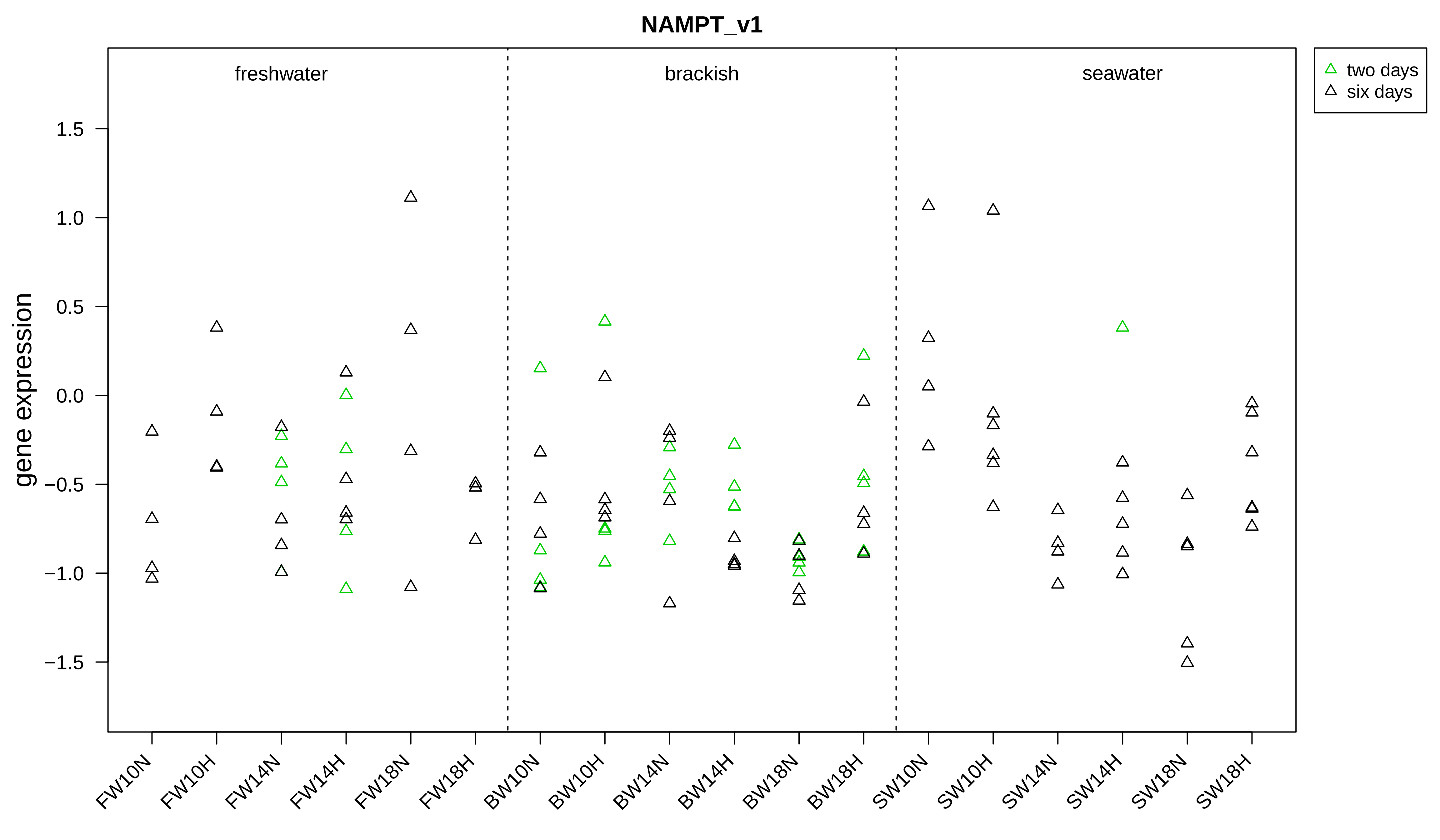
NAMPT_v1
freshwater
brackish
seawater
two days
six days
gene expression
1.5
1.0
0.5
0.0
−0.5
−1.0
−1.5
FW10N
FW10H
FW14N
FW14H
FW18N
FW18H
BW10N
BW10H
BW14N
BW14H
BW18N
BW18H
SW10N
SW10H
SW14N
SW14H
SW18N
SW18H

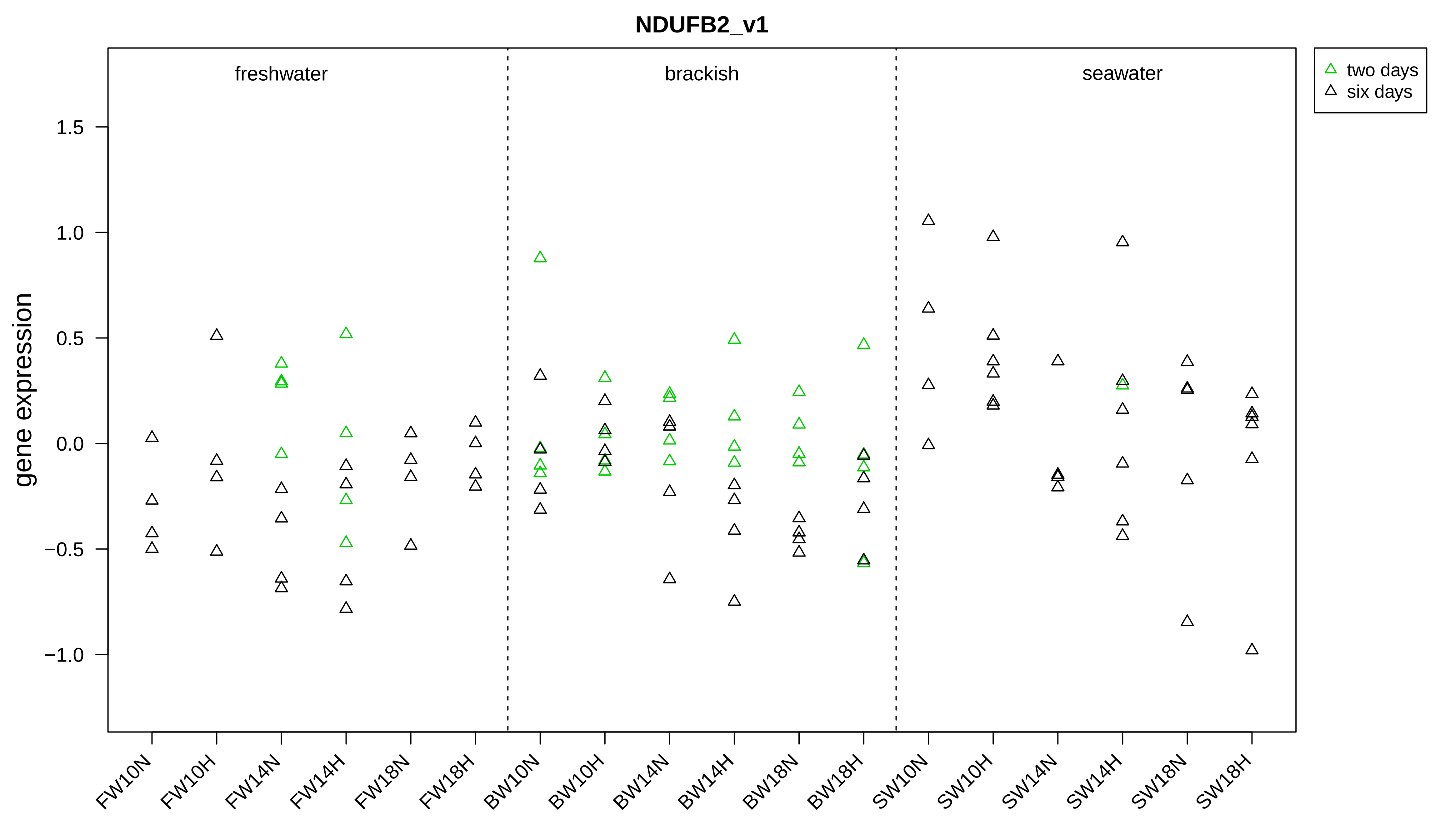

NDUFB2_v1
freshwater
brackish
seawater
two days
six days
gene expression
1.5
1.0
0.5
0.0
−0.5
−1.0
FW10N
FW10H
FW14N
FW14H
FW18N
FW18H
BW10N
BW10H
BW14N
BW14H
BW18N
BW18H
SW10N
SW10H
SW14N
SW14H
SW18N
SW18H

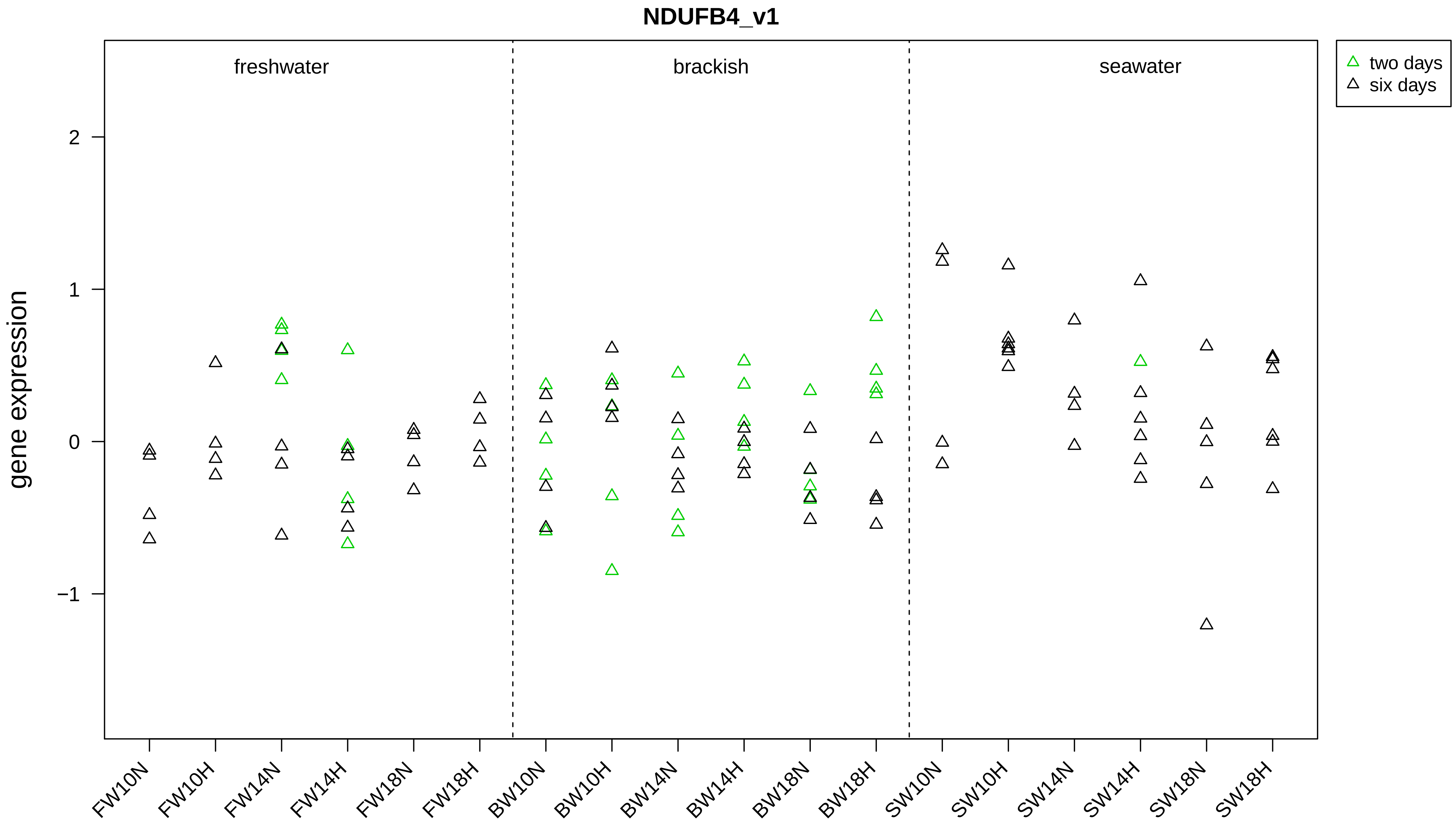

NDUFB4_v1
freshwater
brackish
seawater
two days
six days
gene expression
2
1
0
−1
FW10N
FW10H
FW14N
FW14H
FW18N
FW18H
BW10N
BW10H
BW14N
BW14H
BW18N
BW18H
SW10N
SW10H
SW14N
SW14H
SW18N
SW18H

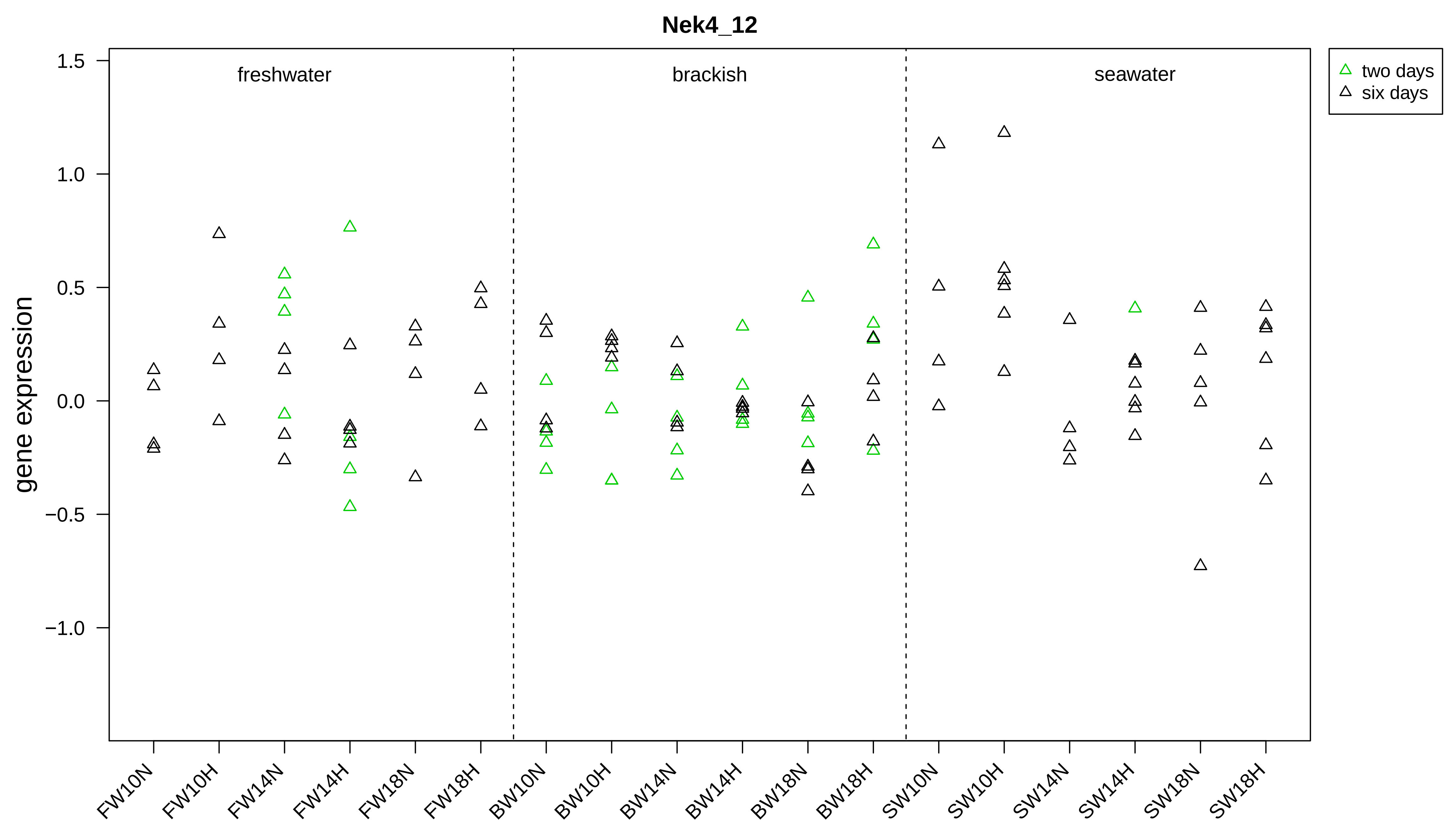
Nek4_12
gene expression
freshwater
brackish
seawater
two days
six days
1.5
1.0
0.5
0.0
−0.5
−1.0
FW10N
FW10H
FW14N
FW14H
FW18N
FW18H
BW10N
BW10H
BW14N
BW14H
BW18N
BW18H
SW10N
SW10H
SW14N
SW14H
SW18N
SW18H

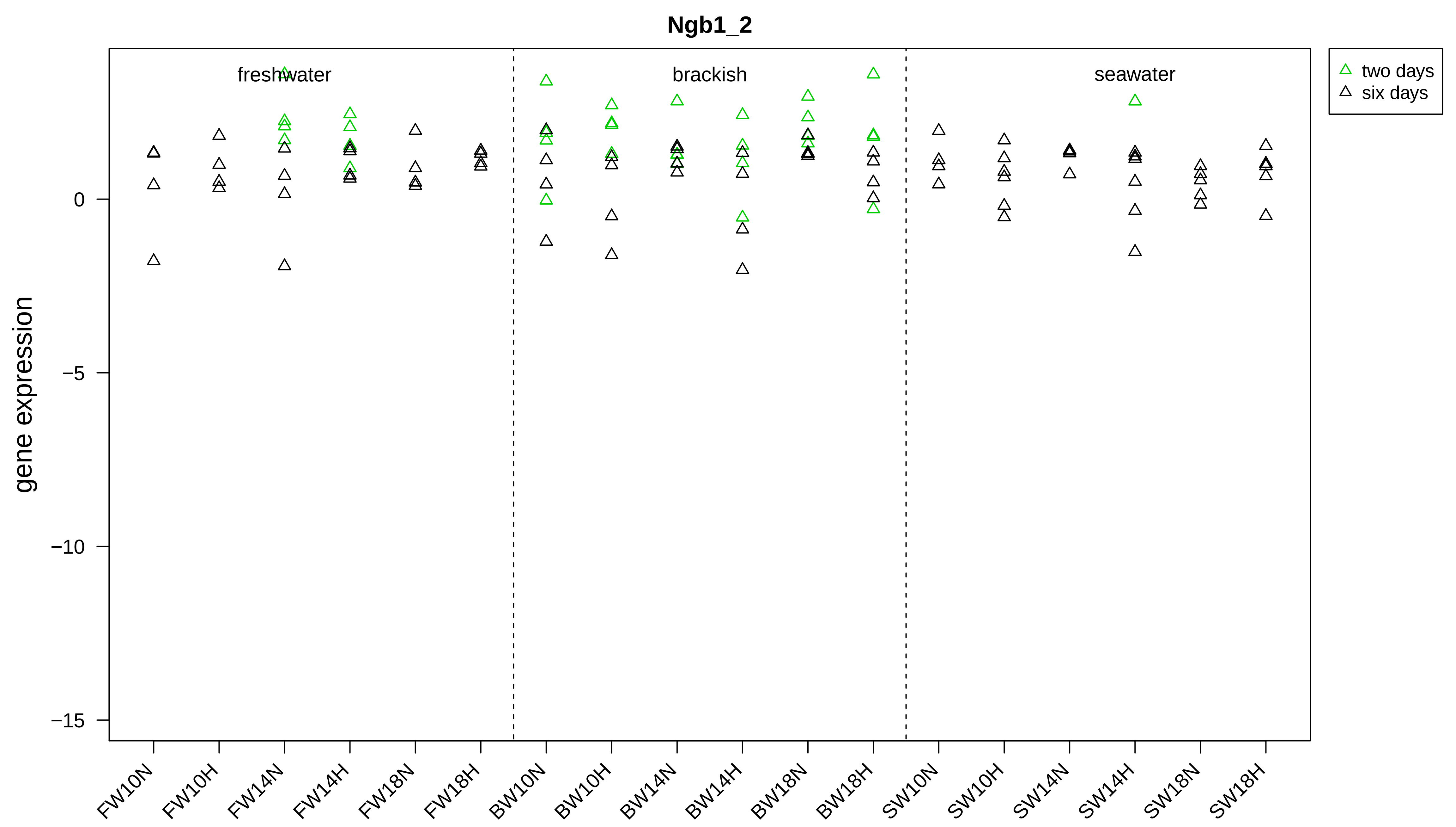
Ngb1_2
freshwater
brackish
seawater
two days
six days
gene expression
0
−5
−10
−15
FW10N
FW10H
FW14N
FW14H
FW18N
FW18H
BW10N
BW10H
BW14N
BW14H
BW18N
BW18H
SW10N
SW10H
SW14N
SW14H
SW18N
SW18H

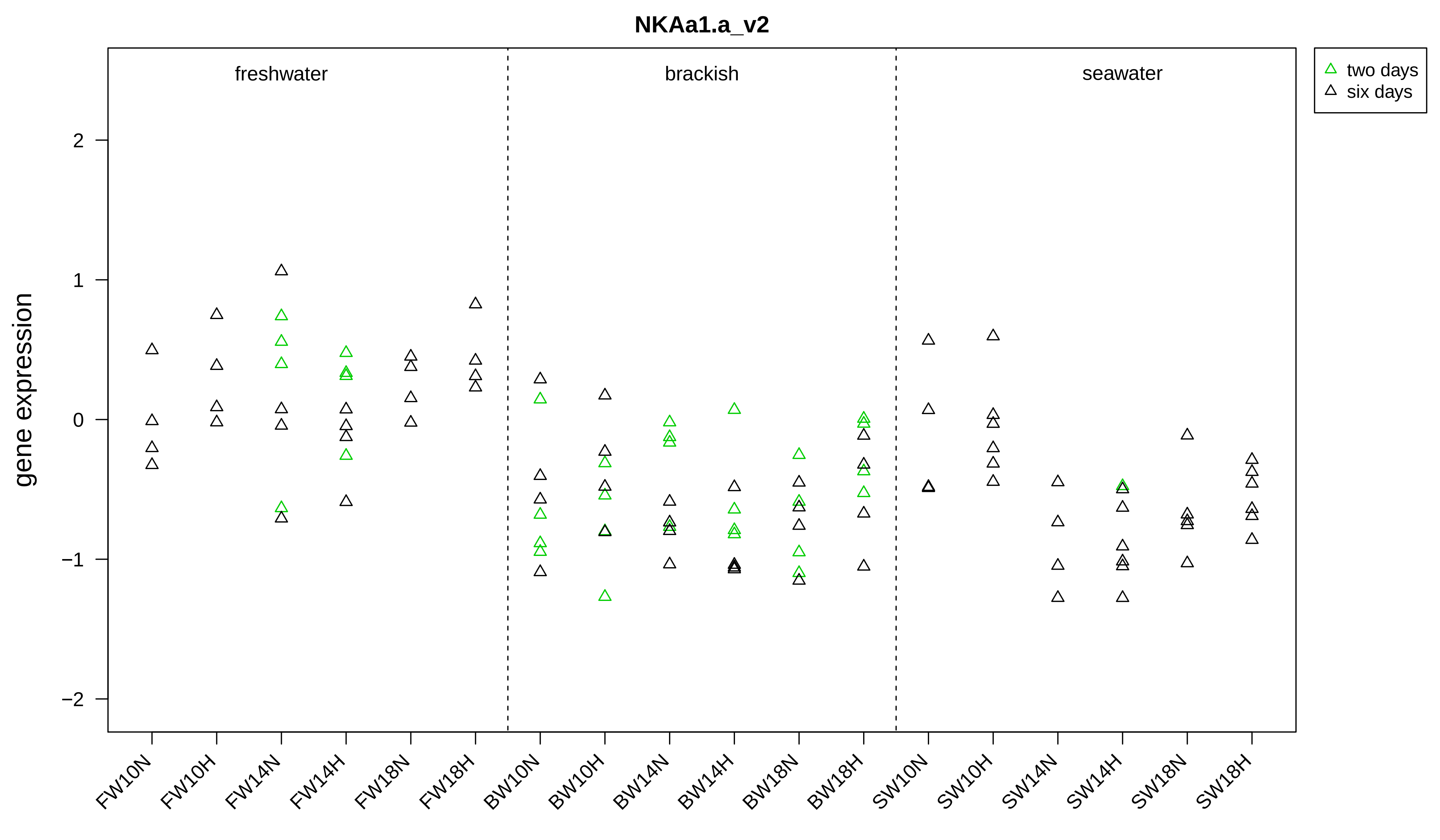
NKAa1.a_v2
freshwater
brackish
seawater
two days
six days
gene expression
2
1
0
−1
−2
FW10N
FW10H
FW14N
FW14H
FW18N
FW18H
BW10N
BW10H
BW14N
BW14H
BW18N
BW18H
SW10N
SW10H
SW14N
SW14H
SW18N
SW18H

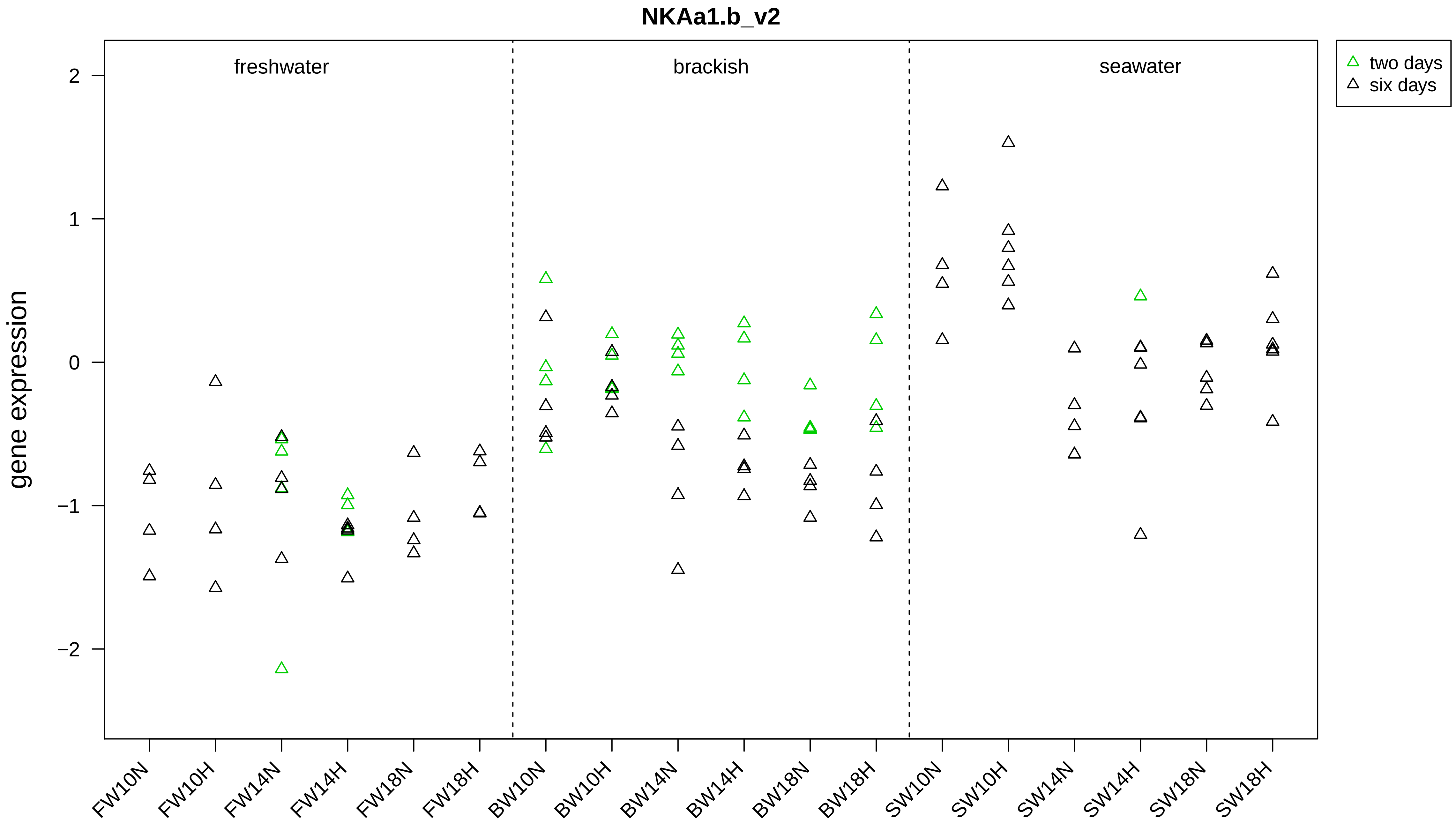
NKAa1.b_v2
freshwater
brackish
seawater
two days
six days
gene expression
2
1
0
−1
−2
FW10N
FW10H
FW14N
FW14H
FW18N
FW18H
BW10N
BW10H
BW14N
BW14H
BW18N
BW18H
SW10N
SW10H
SW14N
SW14H
SW18N
SW18H

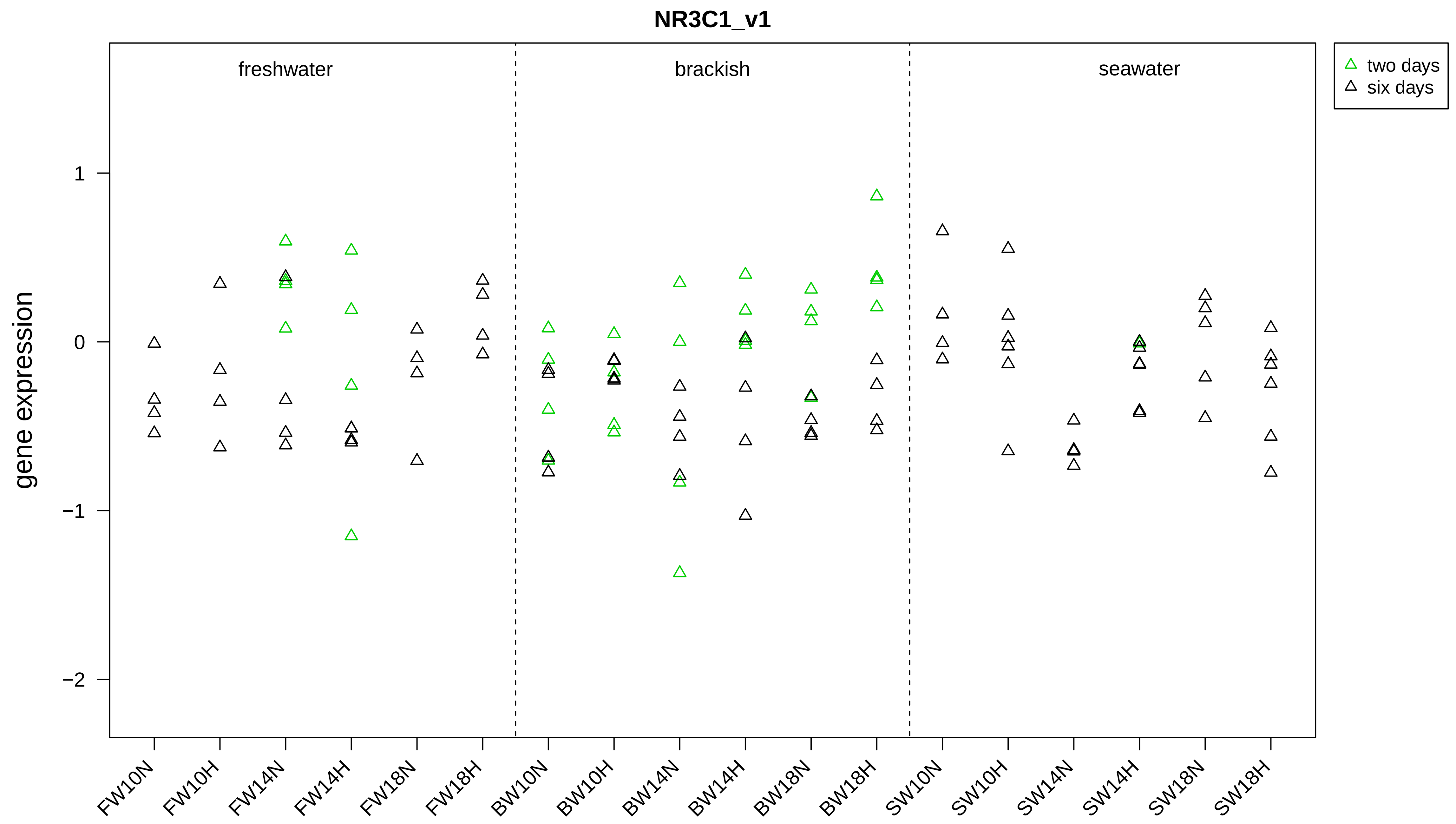
NR3C1_v1
freshwater
brackish
seawater
two days
six days
gene expression
1
0
−1
−2
FW10N
FW10H
FW14N
FW14H
FW18N
FW18H
BW10N
BW10H
BW14N
BW14H
BW18N
BW18H
SW10N
SW10H
SW14N
SW14H
SW18N
SW18H

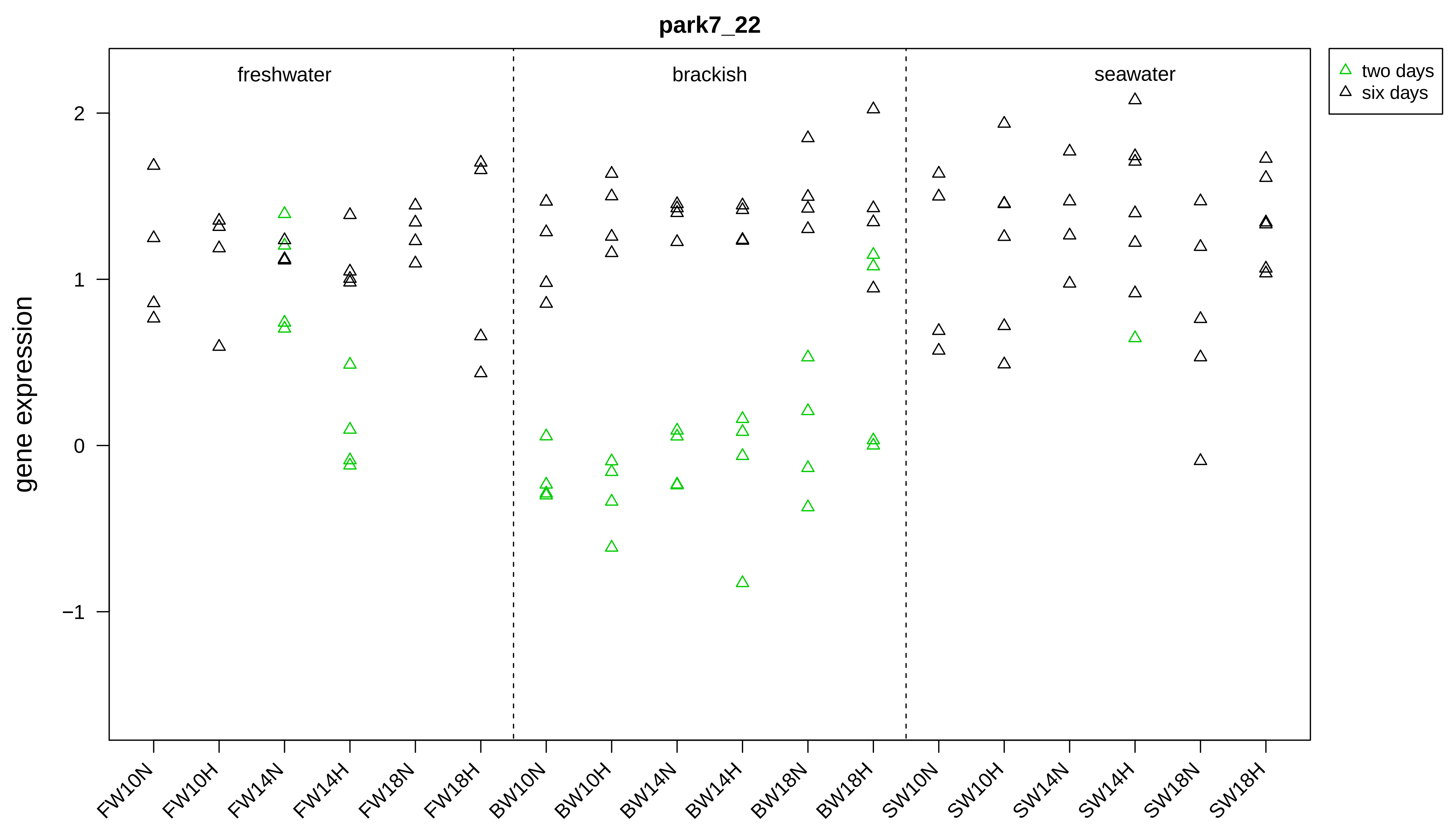

park7_22
freshwater
brackish
seawater
two days
six days
gene expression
2
1
0
−1
FW10N
FW10H
FW14N
FW14H
FW18N
FW18H
BW10N
BW10H
BW14N
BW14H
BW18N
BW18H
SW10N
SW10H
SW14N
SW14H
SW18N
SW18H

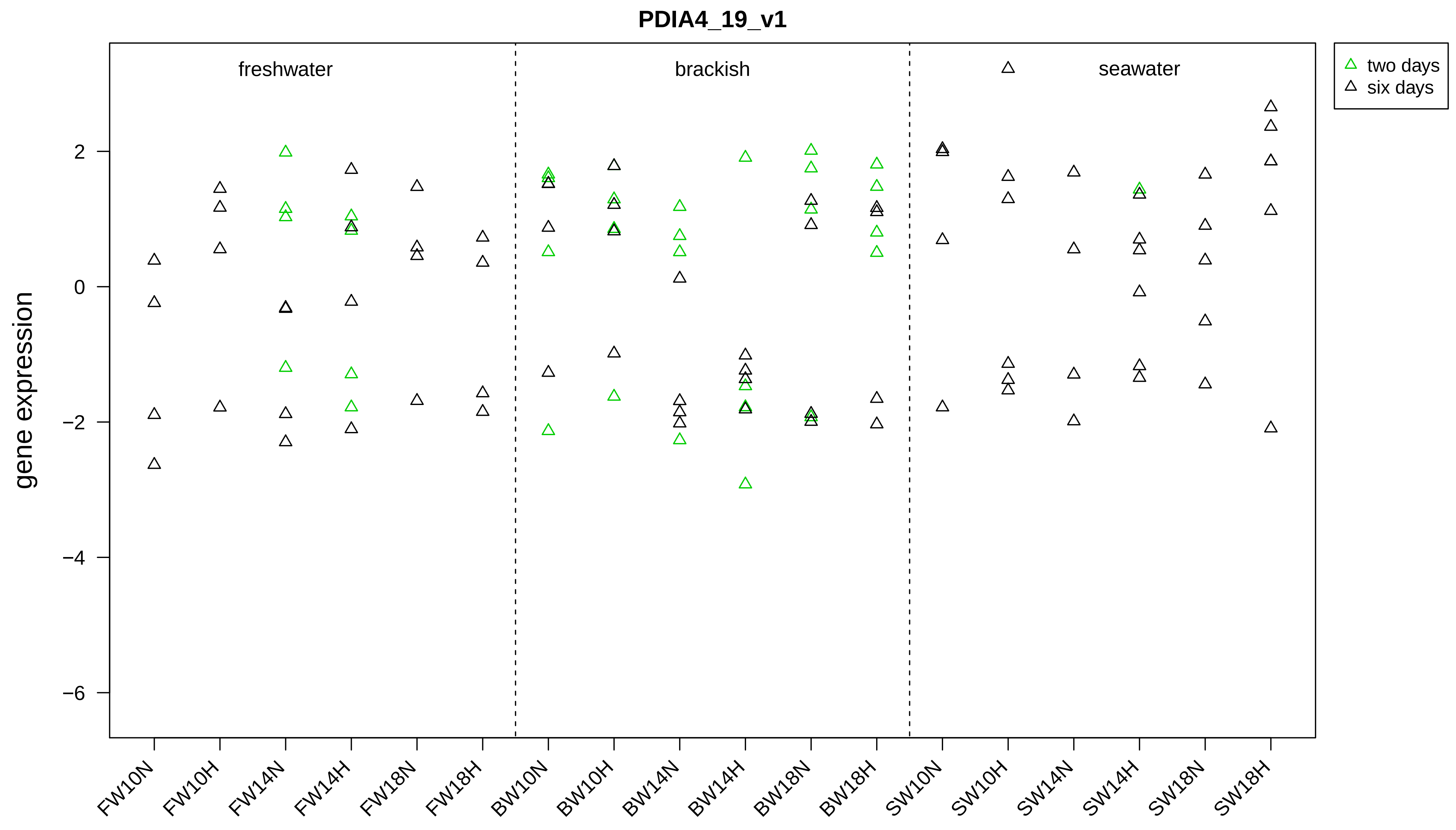

PDIA4_19_v1
freshwater
brackish
seawater
two days
six days
gene expression
2
0
−2
−4
−6
FW10N
FW10H
FW14N
FW14H
FW18N
FW18H
BW10N
BW10H
BW14N
BW14H
BW18N
BW18H
SW10N
SW10H
SW14N
SW14H
SW18N
SW18H

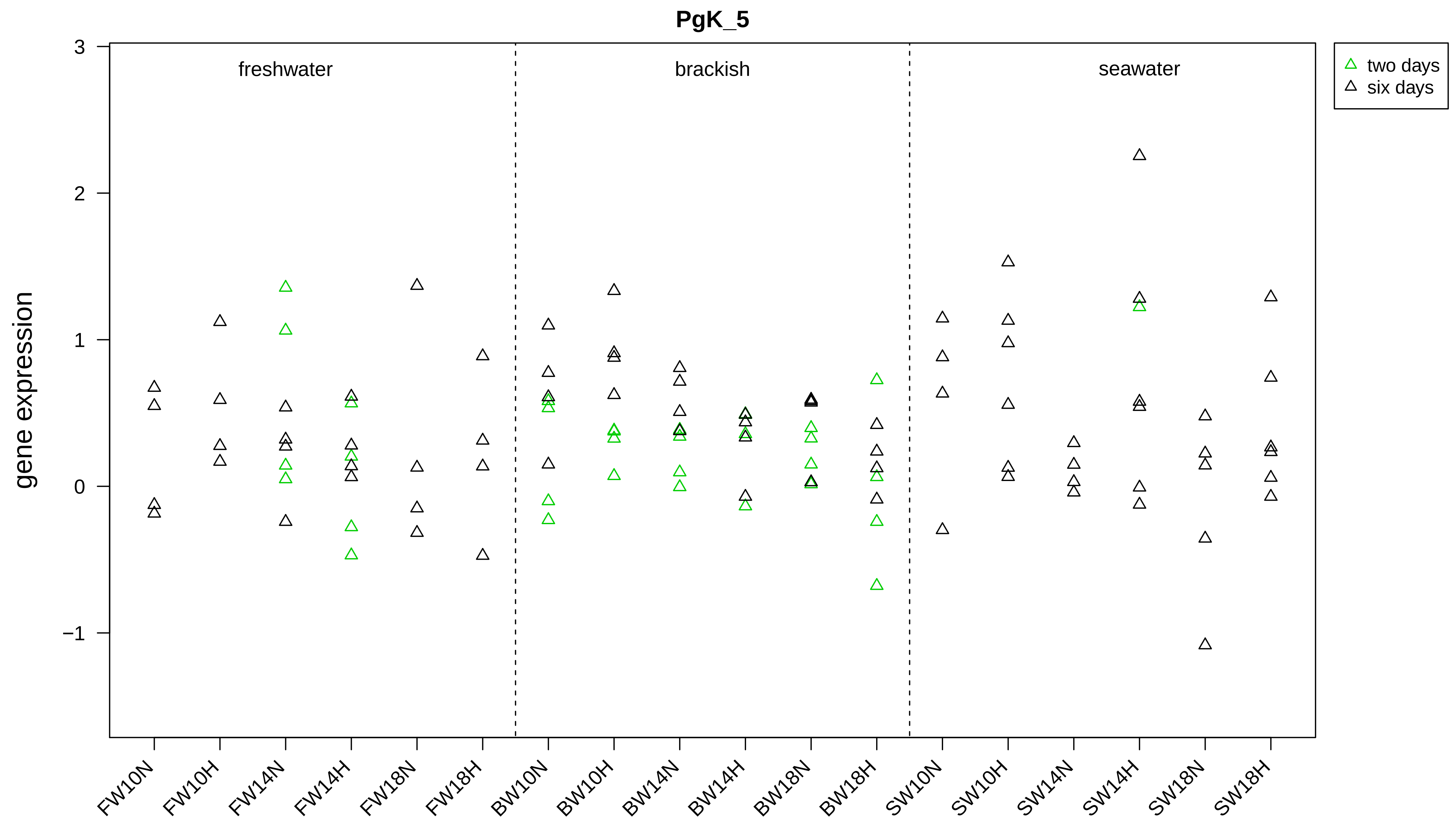

PgK_5
gene expression
3
2
1
0
−1
freshwater
brackish
seawater
two days
six days
FW10N
FW10H
FW14N
FW14H
FW18N
FW18H
BW10N
BW10H
BW14N
BW14H
BW18N
BW18H
SW10N
SW10H
SW14N
SW14H
SW18N
SW18H

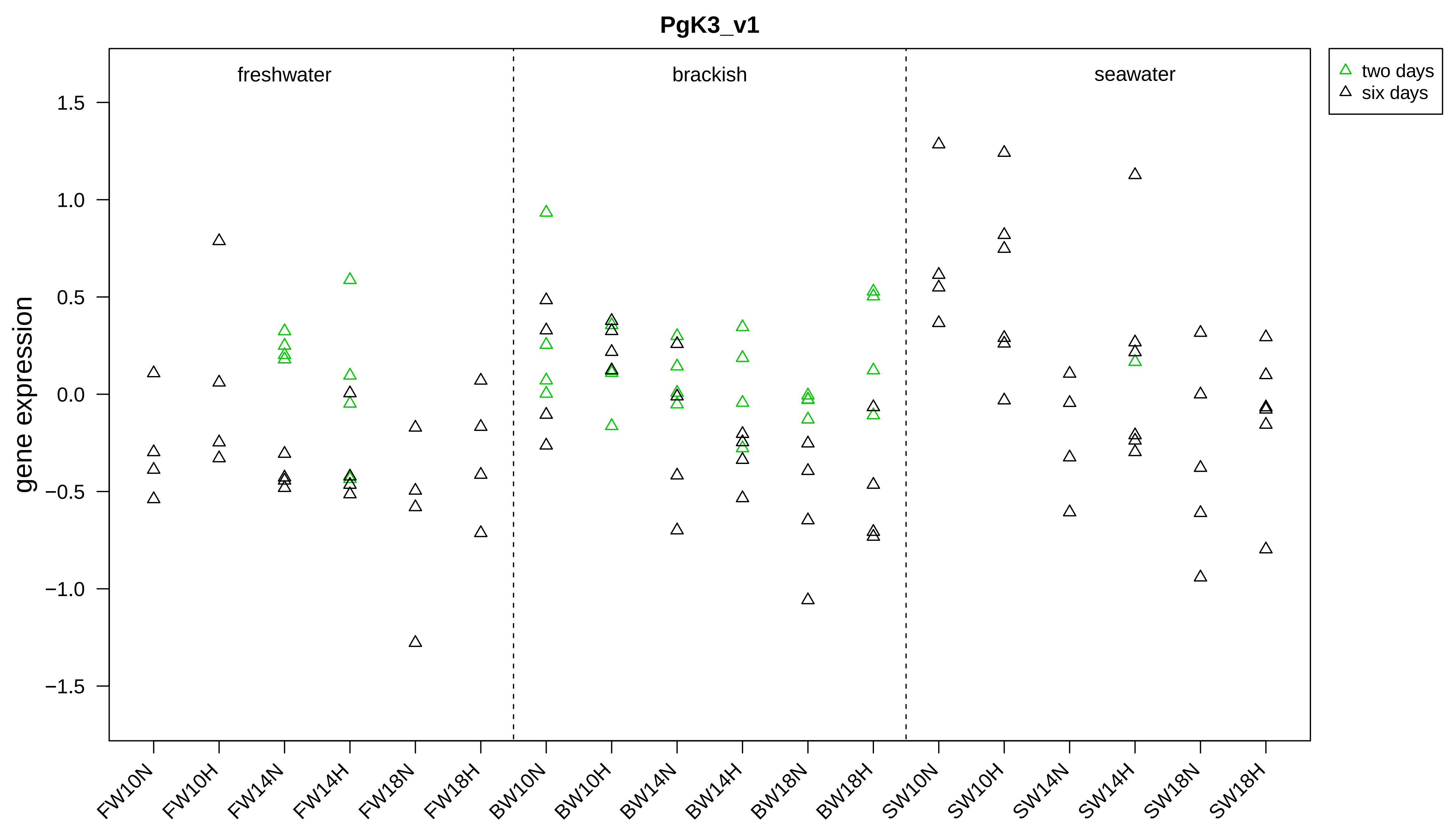

PgK3_v1
gene expression
1.5
1.0
0.5
0.0
−0.5
−1.0
−1.5
freshwater
brackish
seawater
two days
six days
FW10N
FW10H
FW14N
FW14H
FW18N
FW18H
BW10N
BW10H
BW14N
BW14H
BW18N
BW18H
SW10N
SW10H
SW14N
SW14H
SW18N
SW18H

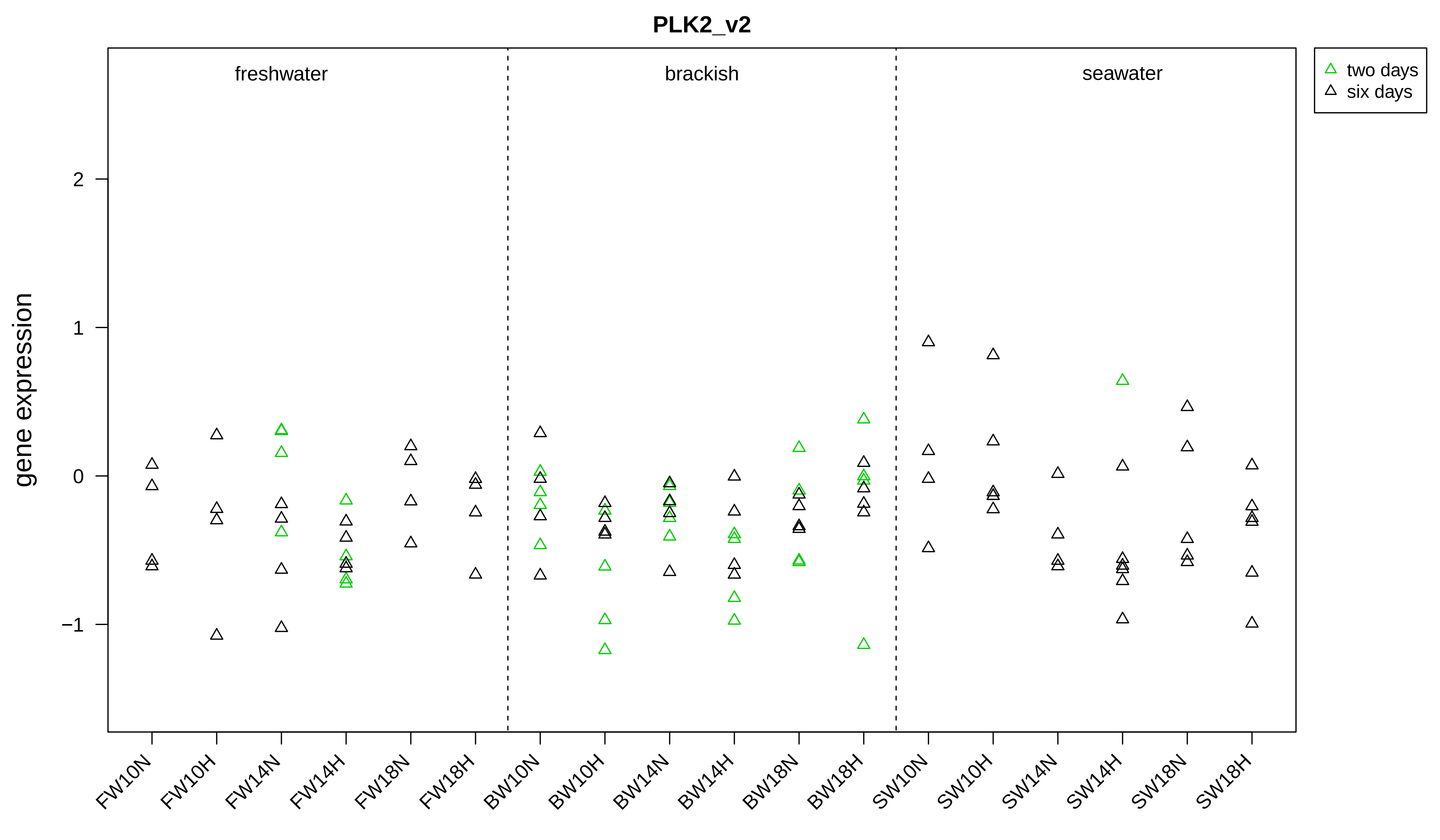

PLK2_v2
gene expression
freshwater
brackish
seawater
two days
six days
2
1
0
−1
FW10N
FW10H
FW14N
FW14H
FW18N
FW18H
BW10N
BW10H
BW14N
BW14H
BW18N
BW18H
SW10N
SW10H
SW14N
SW14H
SW18N
SW18H

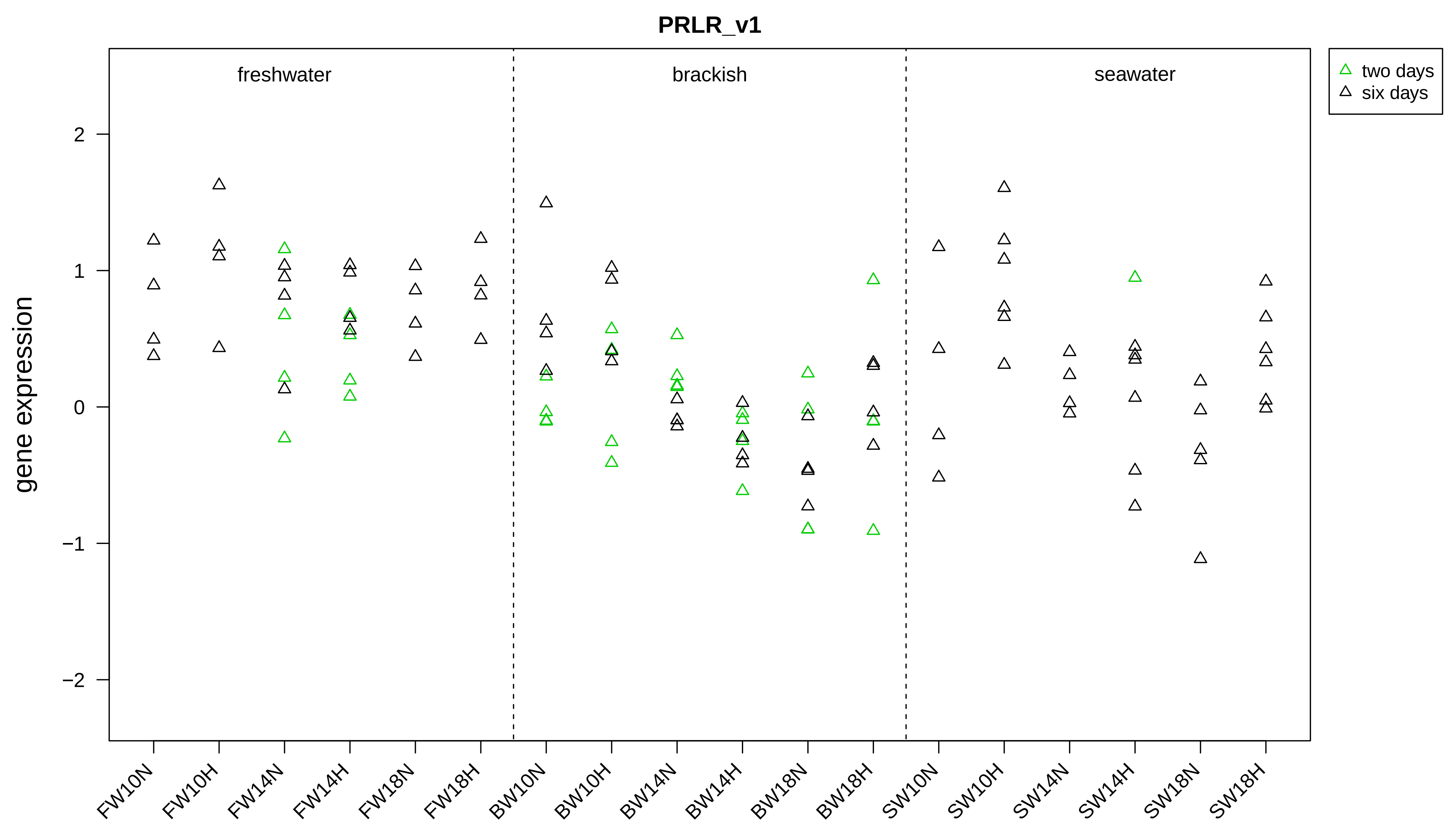

PRLR_v1
freshwater
brackish
seawater
two days
six days
gene expression
2
1
0
−1
−2
FW10N
FW10H
FW14N
FW14H
FW18N
FW18H
BW10N
BW10H
BW14N
BW14H
BW18N
BW18H
SW10N
SW10H
SW14N
SW14H
SW18N
SW18H

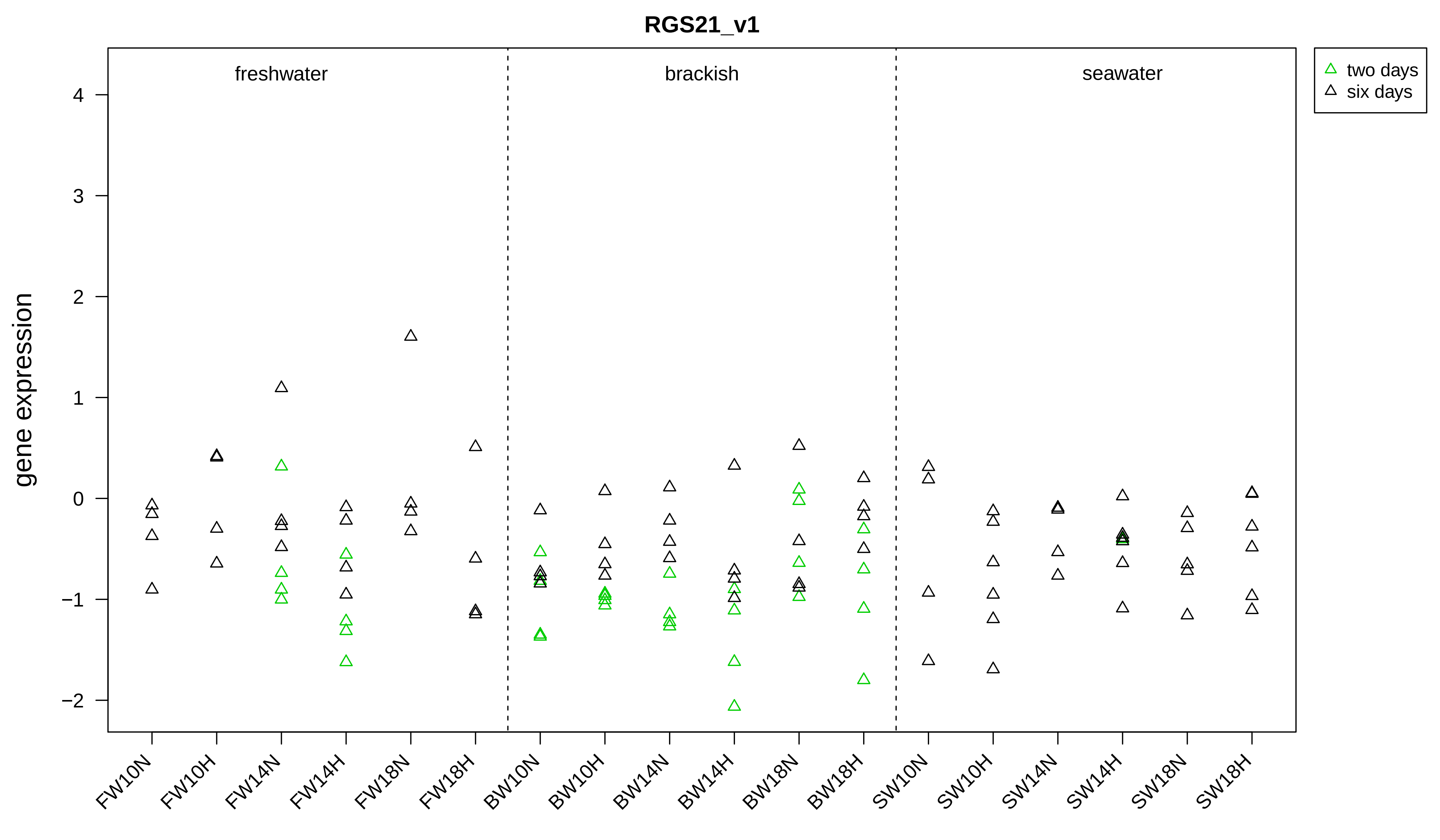

RGS21_v1
freshwater
brackish
seawater
two days
six days
gene expression
4
3
2
1
0
−1
−2
FW10N
FW10H
FW14N
FW14H
FW18N
FW18H
BW10N
BW10H
BW14N
BW14H
BW18N
BW18H
SW10N
SW10H
SW14N
SW14H
SW18N
SW18H

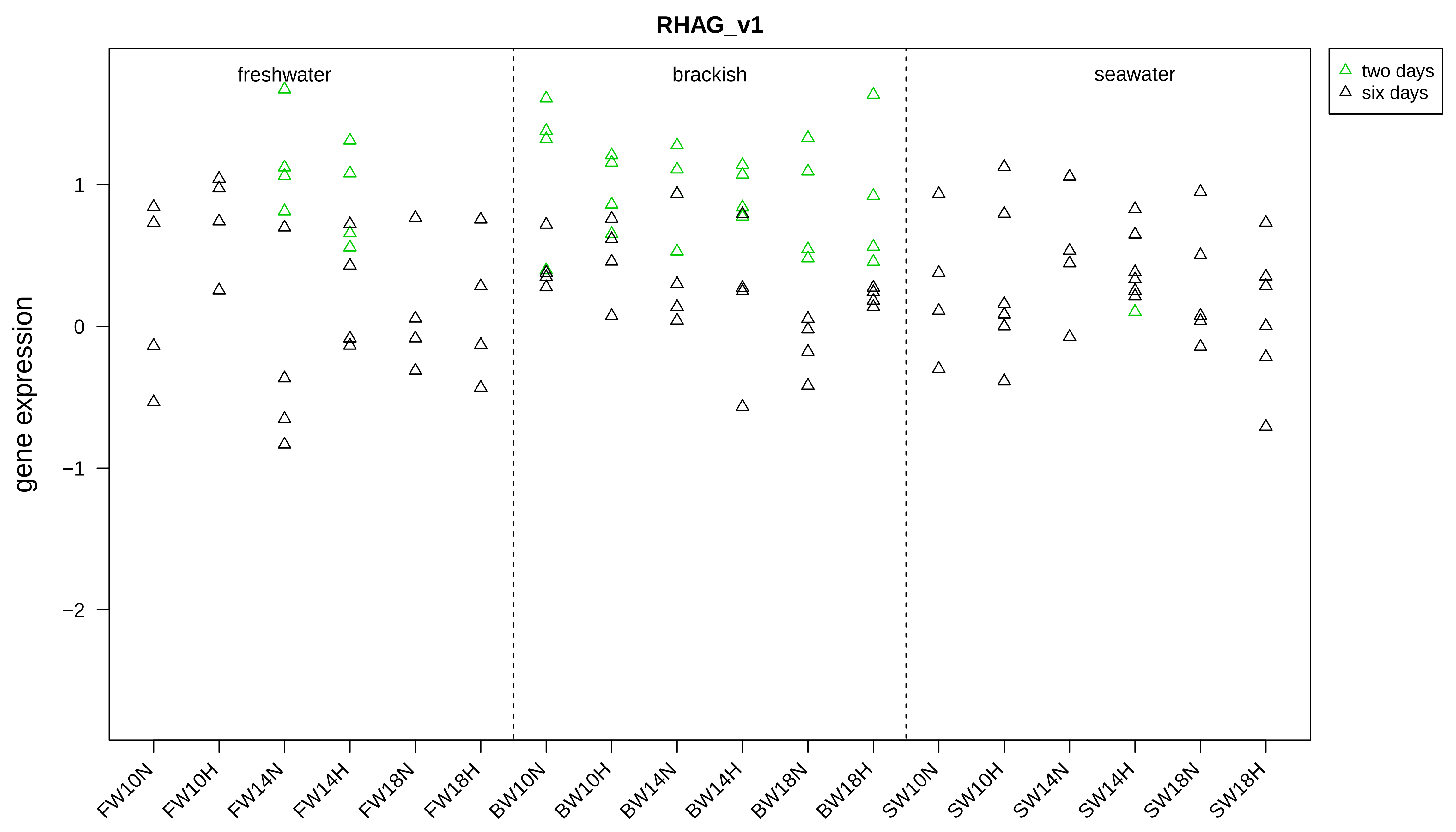
RHAG_v1
freshwater
brackish
seawater
two days
six days
gene expression
1
0
−1
−2
FW10N
FW10H
FW14N
FW14H
FW18N
FW18H
BW10N
BW10H
BW14N
BW14H
BW18N
BW18H
SW10N
SW10H
SW14N
SW14H
SW18N
SW18H

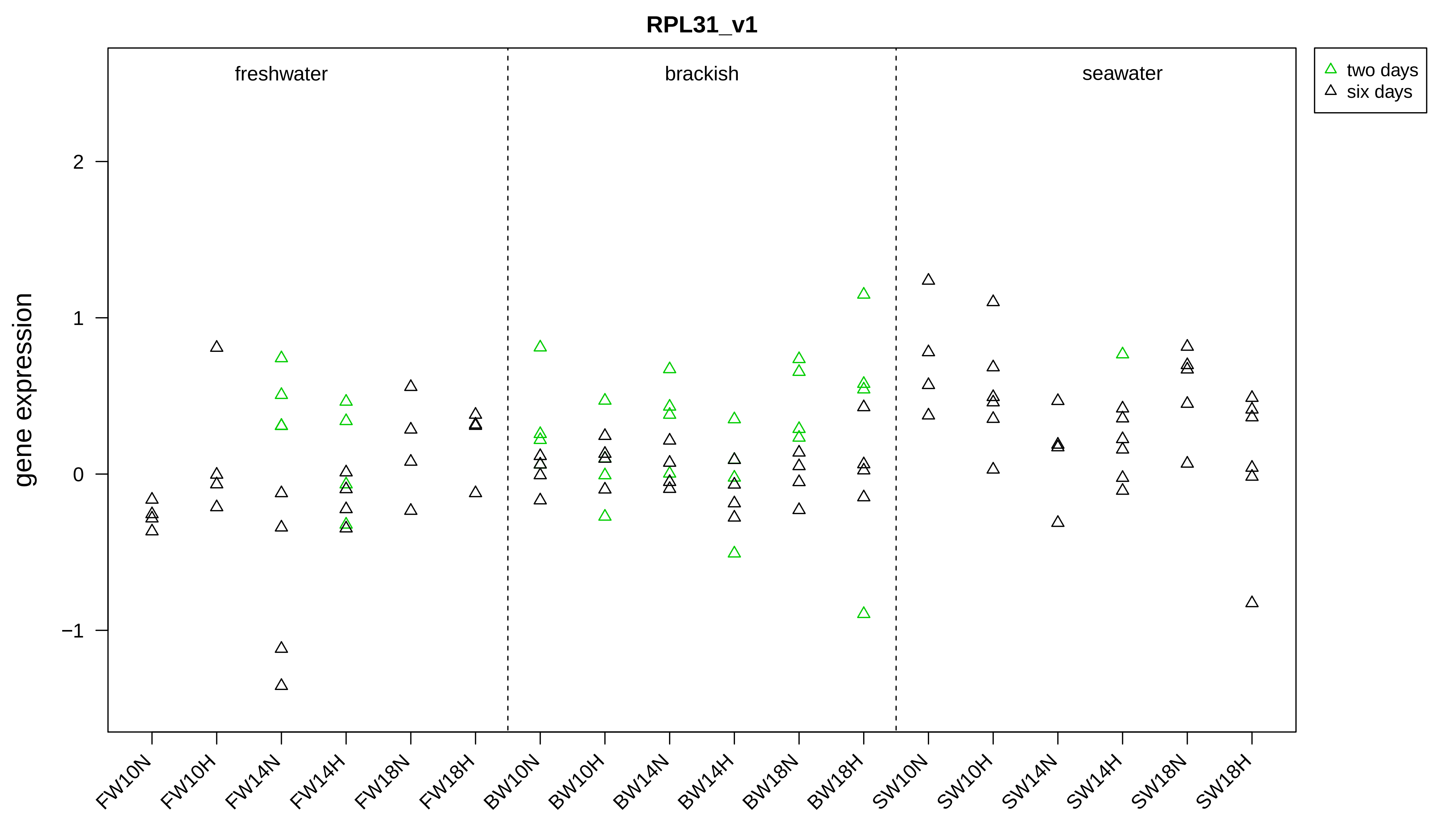

RPL31_v1
gene expression
freshwater
brackish
seawater
two days
six days
2
1
0
−1
FW10N
FW10H
FW14N
FW14H
FW18N
FW18H
BW10N
BW10H
BW14N
BW14H
BW18N
BW18H
SW10N
SW10H
SW14N
SW14H
SW18N
SW18H

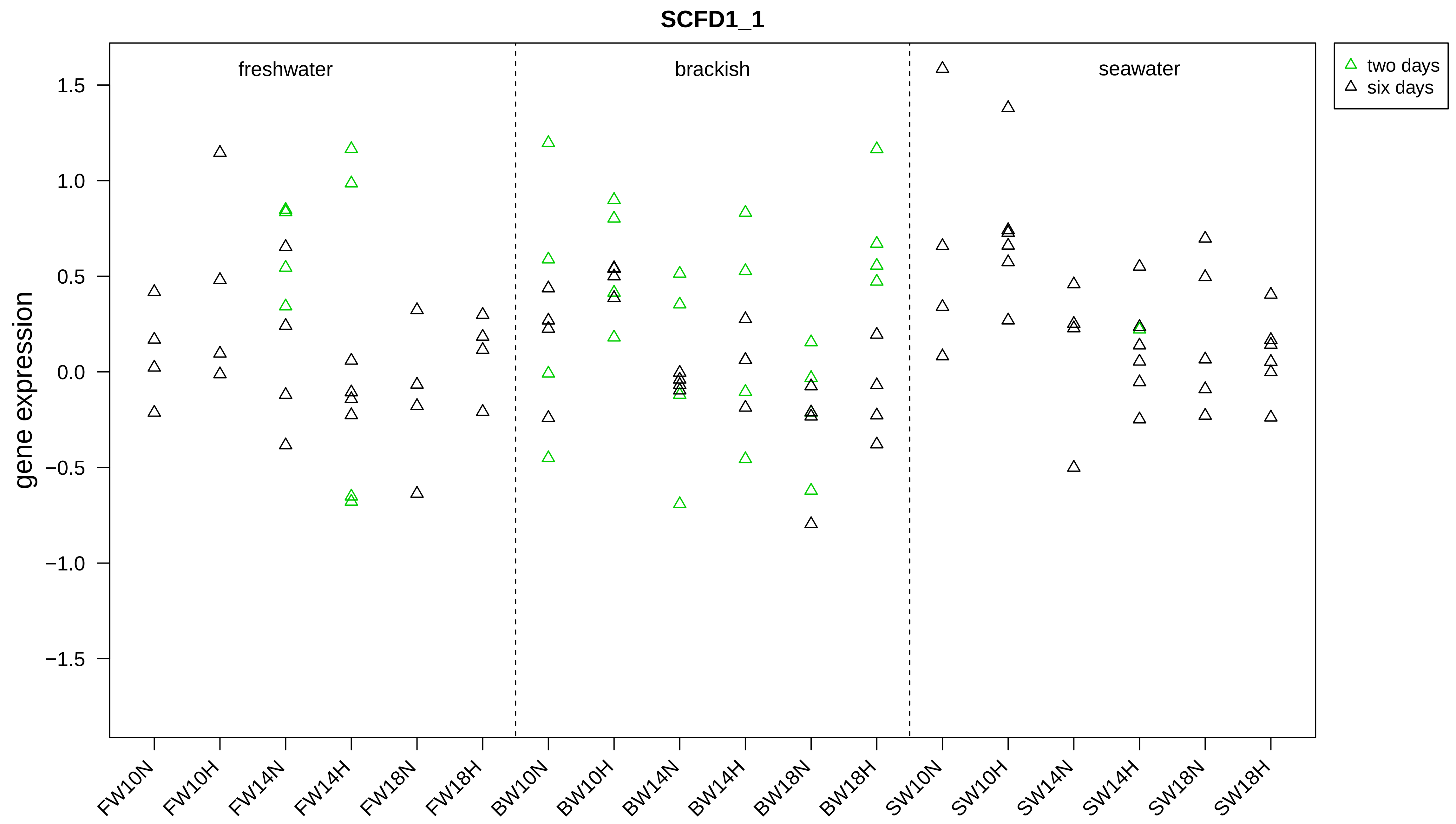
SCFD1_1
gene expression
freshwater
brackish
seawater
two days
six days
1.5
1.0
0.5
0.0
−0.5
−1.0
−1.5
FW10N
FW10H
FW14N
FW14H
FW18N
FW18H
BW10N
BW10H
BW14N
BW14H
BW18N
BW18H
SW10N
SW10H
SW14N
SW14H
SW18N
SW18H

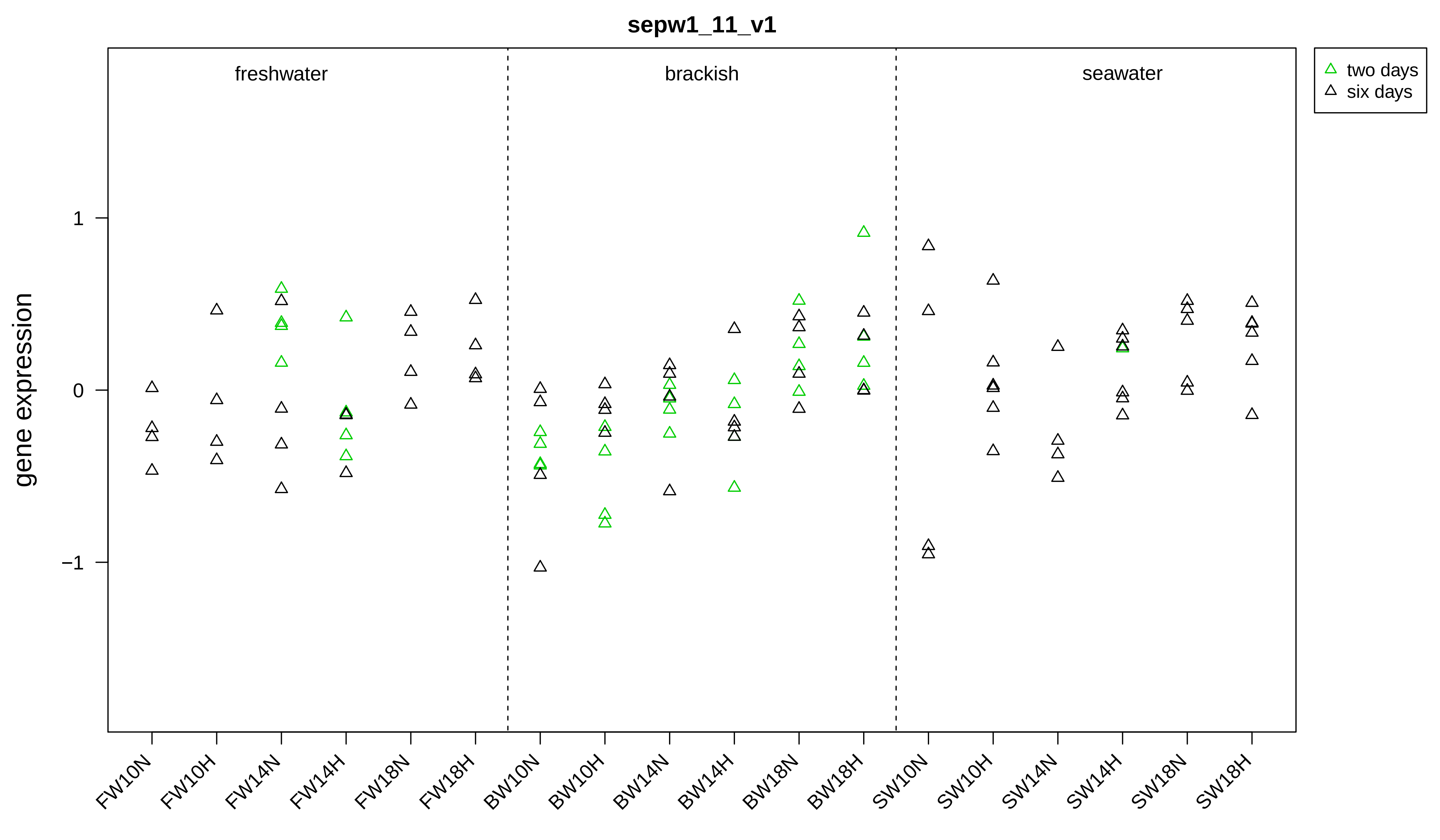
sepw1_11_v1
freshwater
brackish
seawater
two days
six days
gene expression
1
0
−1
FW10N
FW10H
FW14N
FW14H
FW18N
FW18H
BW10N
BW10H
BW14N
BW14H
BW18N
BW18H
SW10N
SW10H
SW14N
SW14H
SW18N
SW18H

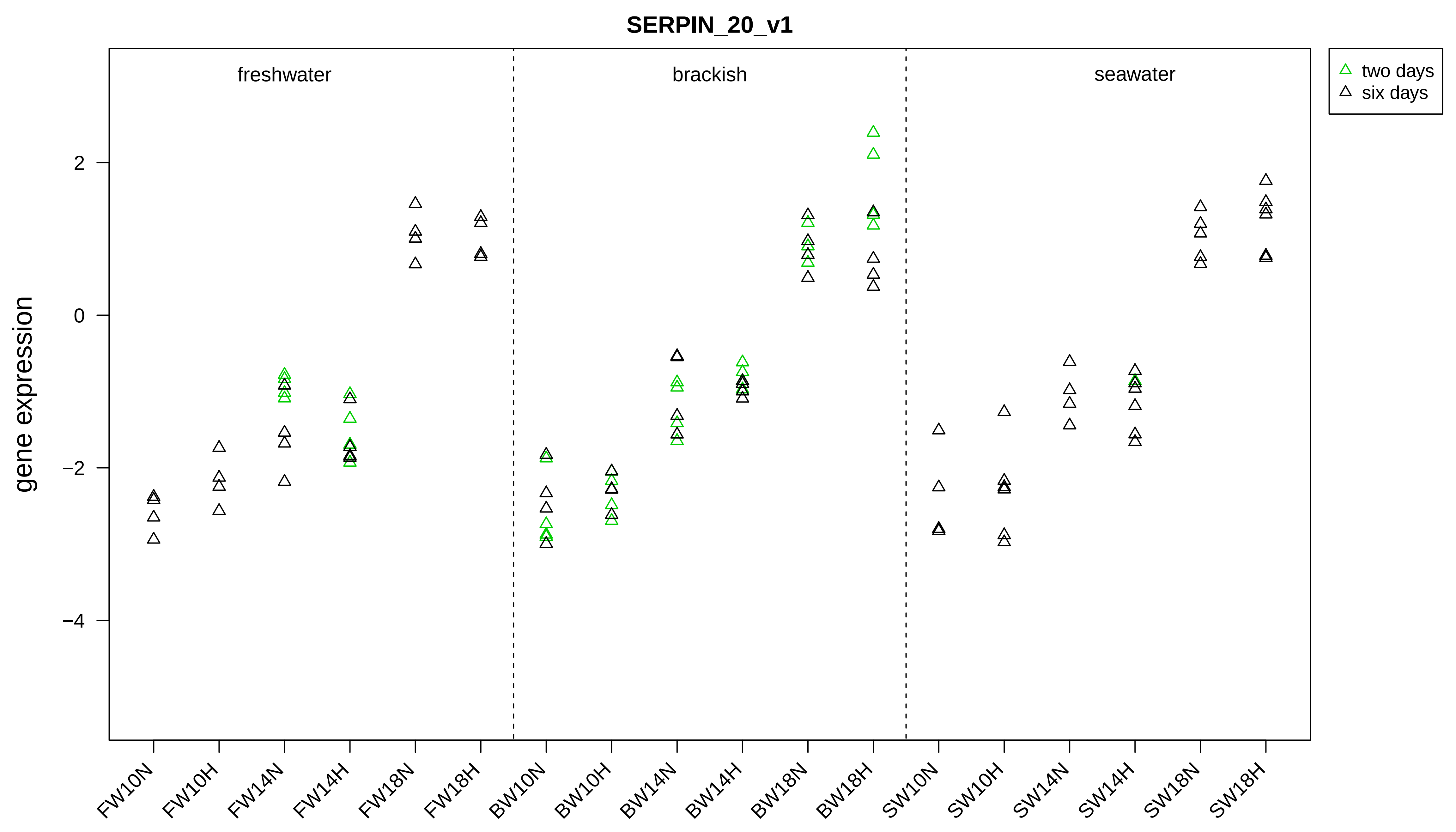

SERPIN_20_v1
freshwater
brackish
seawater
two days
six days
gene expression
2
0
−2
−4
FW10N
FW10H
FW14N
FW14H
FW18N
FW18H
BW10N
BW10H
BW14N
BW14H
BW18N
BW18H
SW10N
SW10H
SW14N
SW14H
SW18N
SW18H

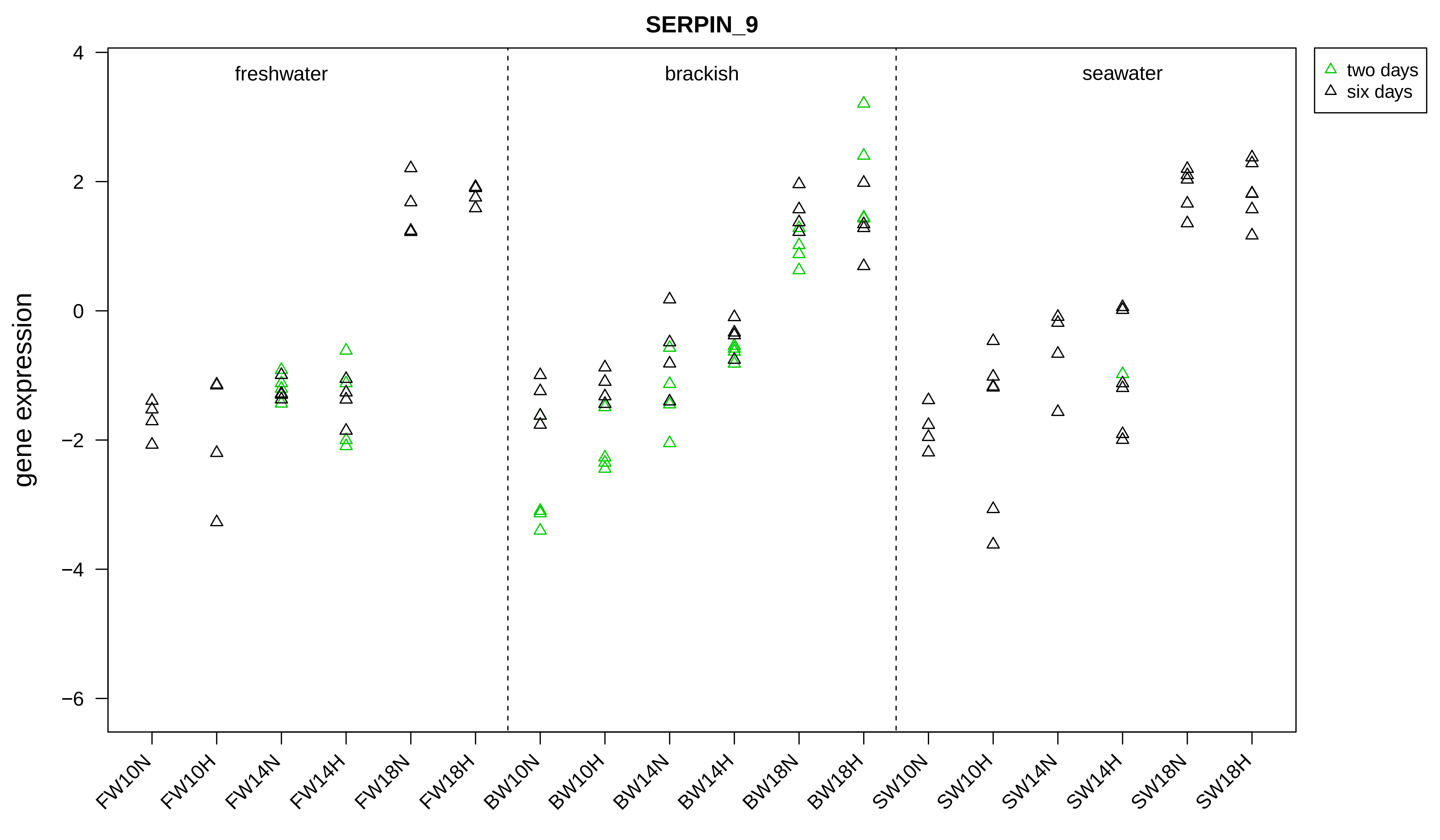
SERPIN_9
gene expression
4
2
0
−2
−4
−6
freshwater
brackish
seawater
two days
six days
FW10N
FW10H
FW14N
FW14H
FW18N
FW18H
BW10N
BW10H
BW14N
BW14H
BW18N
BW18H
SW10N
SW10H
SW14N
SW14H
SW18N
SW18H

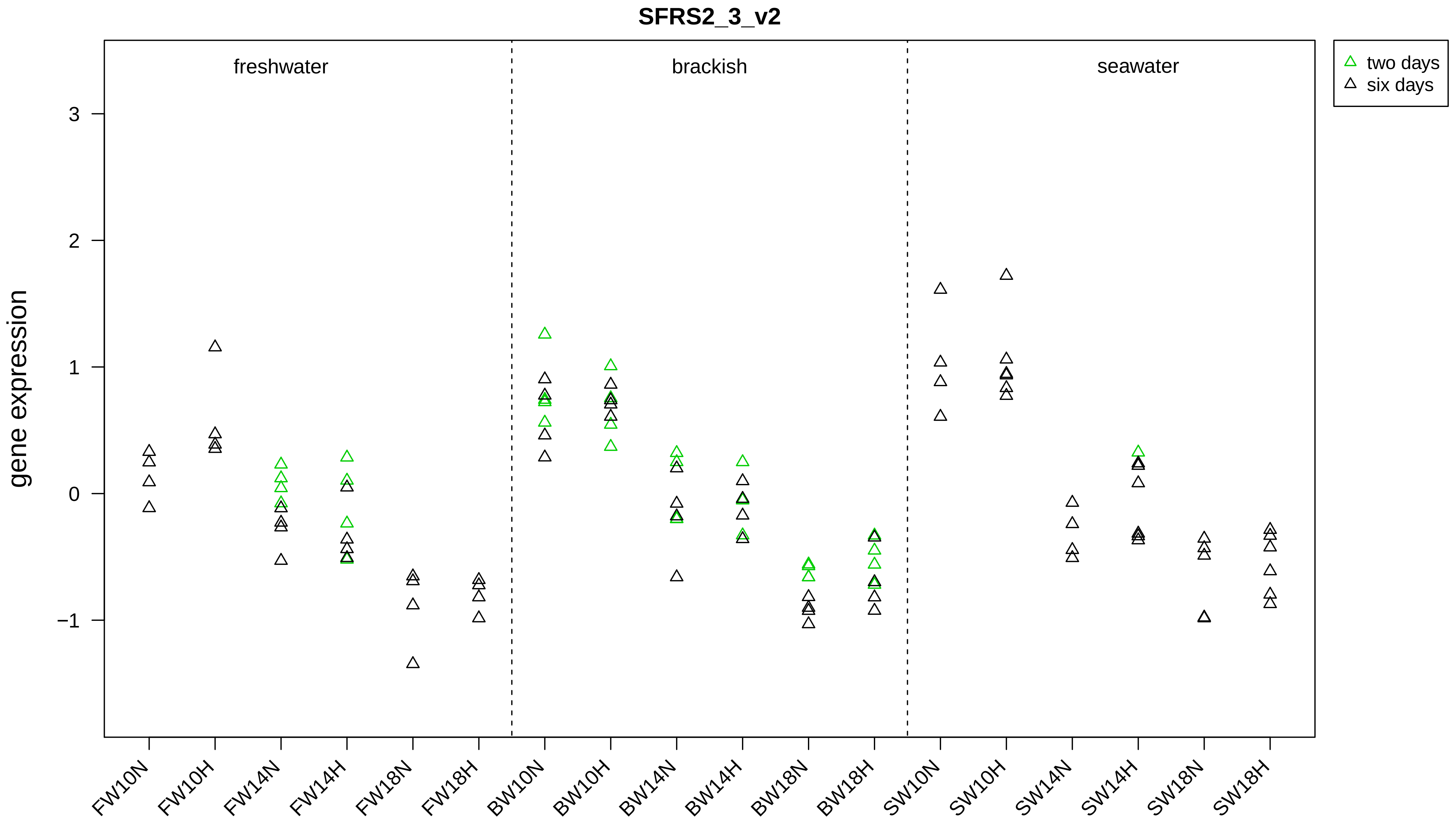

SFRS2_3_v2
gene expression
freshwater
brackish
seawater
two days
six days
3
2
1
0
−1
FW10N
FW10H
FW14N
FW14H
FW18N
FW18H
BW10N
BW10H
BW14N
BW14H
BW18N
BW18H
SW10N
SW10H
SW14N
SW14H
SW18N
SW18H

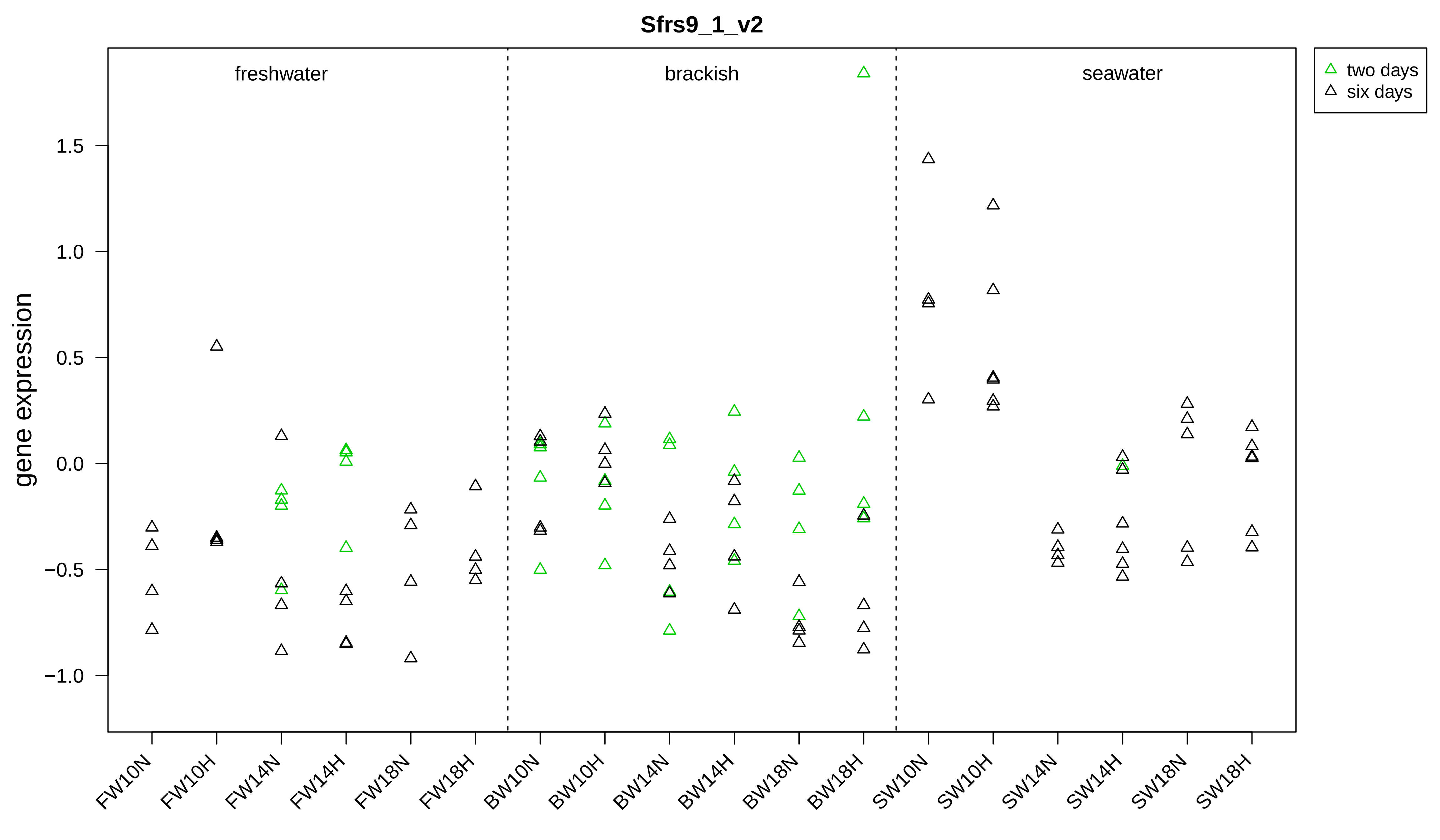
Sfrs9_1_v2
freshwater
brackish
seawater
two days
six days
gene expression
1.5
1.0
0.5
0.0
−0.5
−1.0
FW10N
FW10H
FW14N
FW14H
FW18N
FW18H
BW10N
BW10H
BW14N
BW14H
BW18N
BW18H
SW10N
SW10H
SW14N
SW14H
SW18N
SW18H

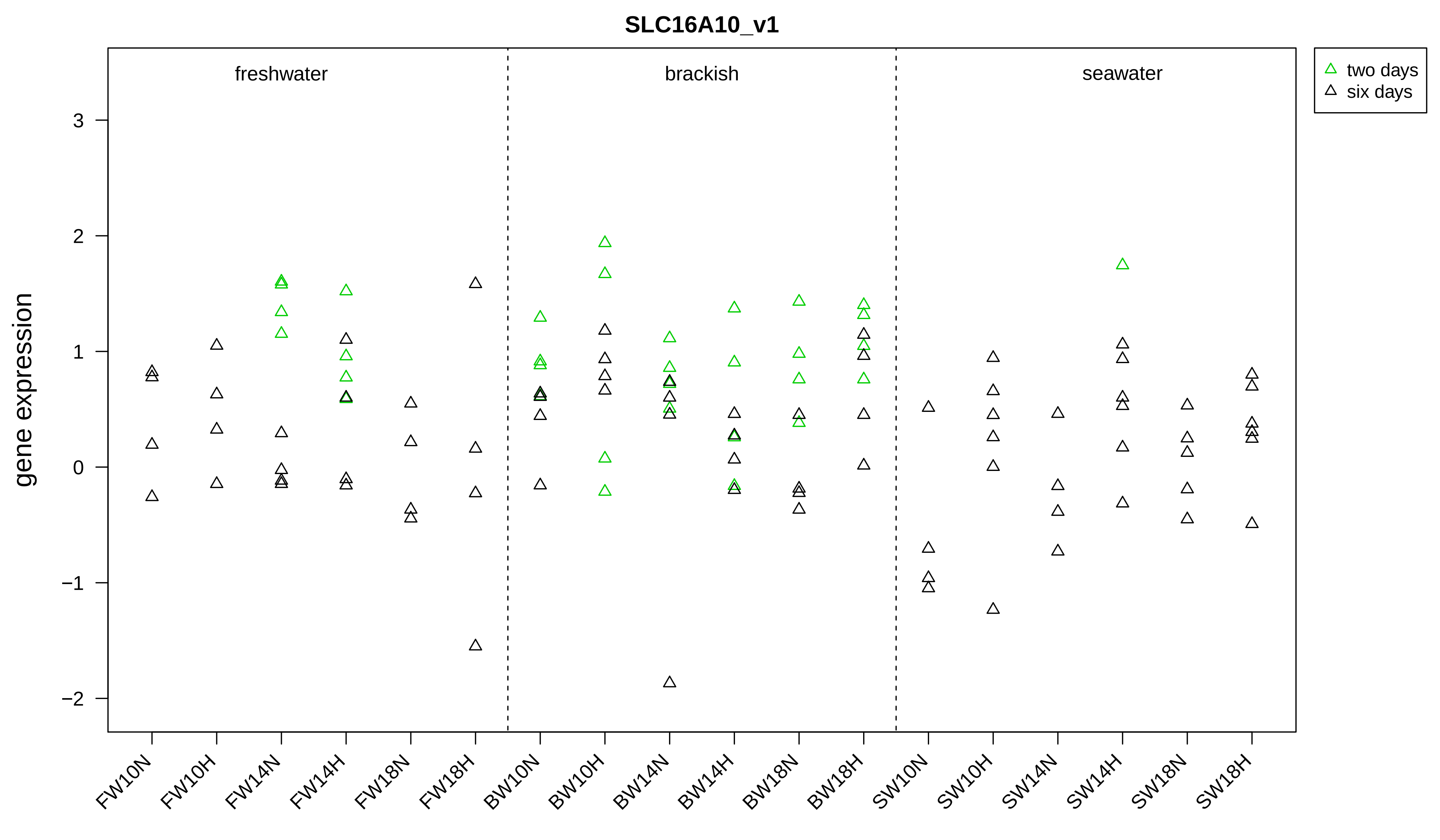

SLC16A10_v1
freshwater
brackish
seawater
two days
six days
gene expression
3
2
1
0
−1
−2
FW10N
FW10H
FW14N
FW14H
FW18N
FW18H
BW10N
BW10H
BW14N
BW14H
BW18N
BW18H
SW10N
SW10H
SW14N
SW14H
SW18N
SW18H

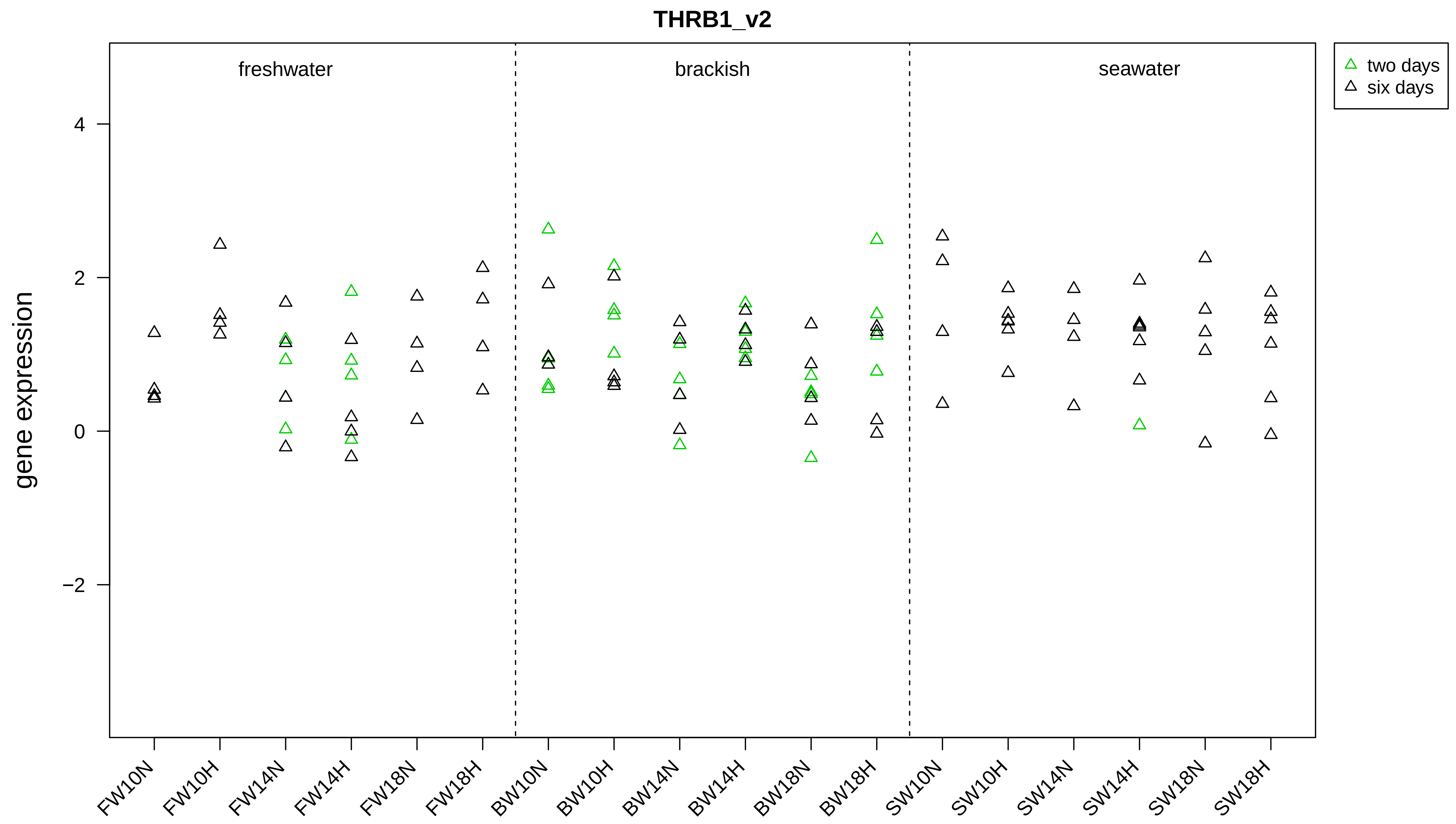
THRB1_v2
freshwater
brackish
seawater
two days
six days
gene expression
4
2
0
−2
FW10N
FW10H
FW14N
FW14H
FW18N
FW18H
BW10N
BW10H
BW14N
BW14H
BW18N
BW18H
SW10N
SW10H
SW14N
SW14H
SW18N
SW18H

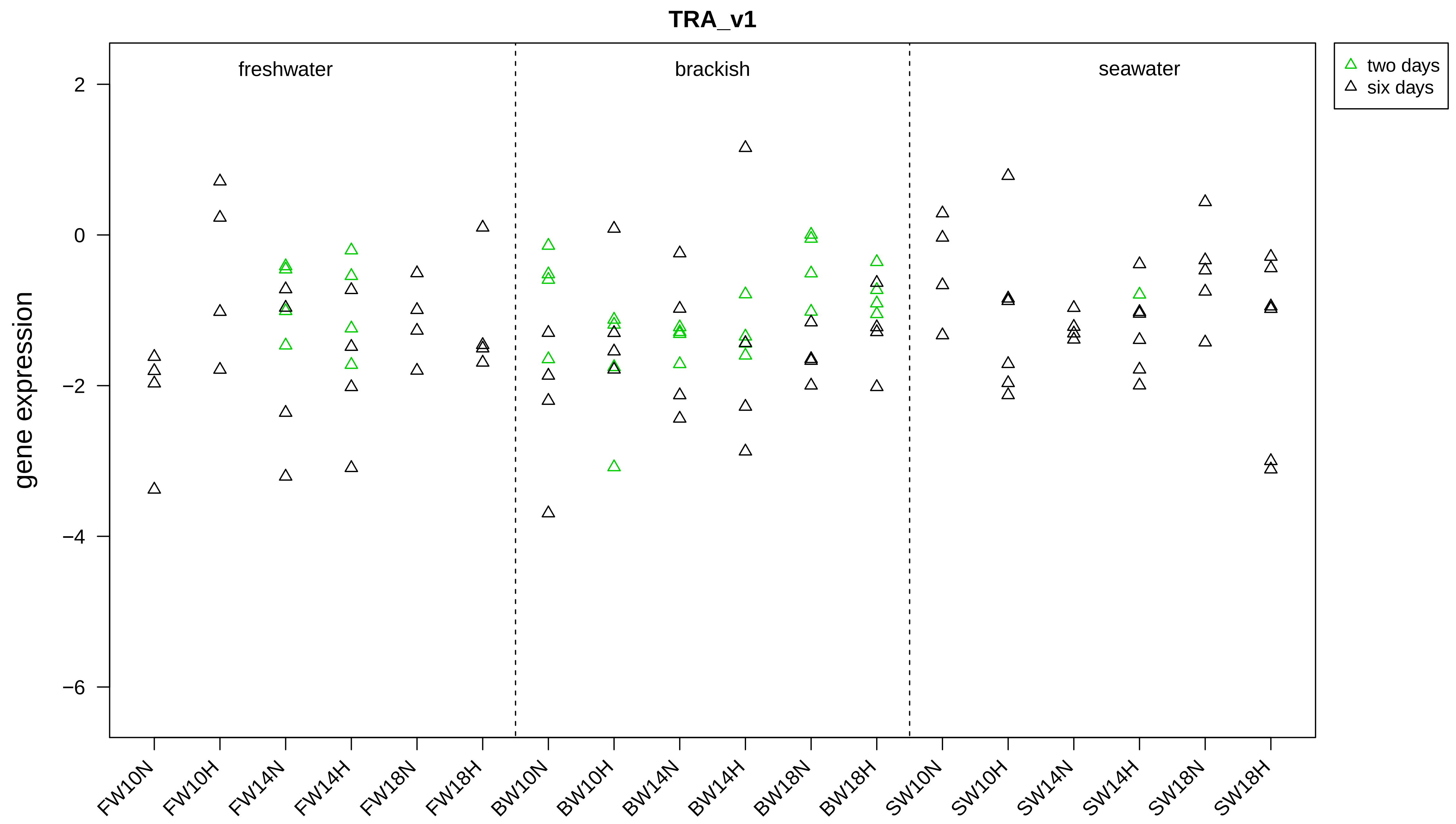
TRA_v1
freshwater
brackish
seawater
two days
six days
gene expression
2
0
−2
−4
−6
FW10N
FW10H
FW14N
FW14H
FW18N
FW18H
BW10N
BW10H
BW14N
BW14H
BW18N
BW18H
SW10N
SW10H
SW14N
SW14H
SW18N
SW18H

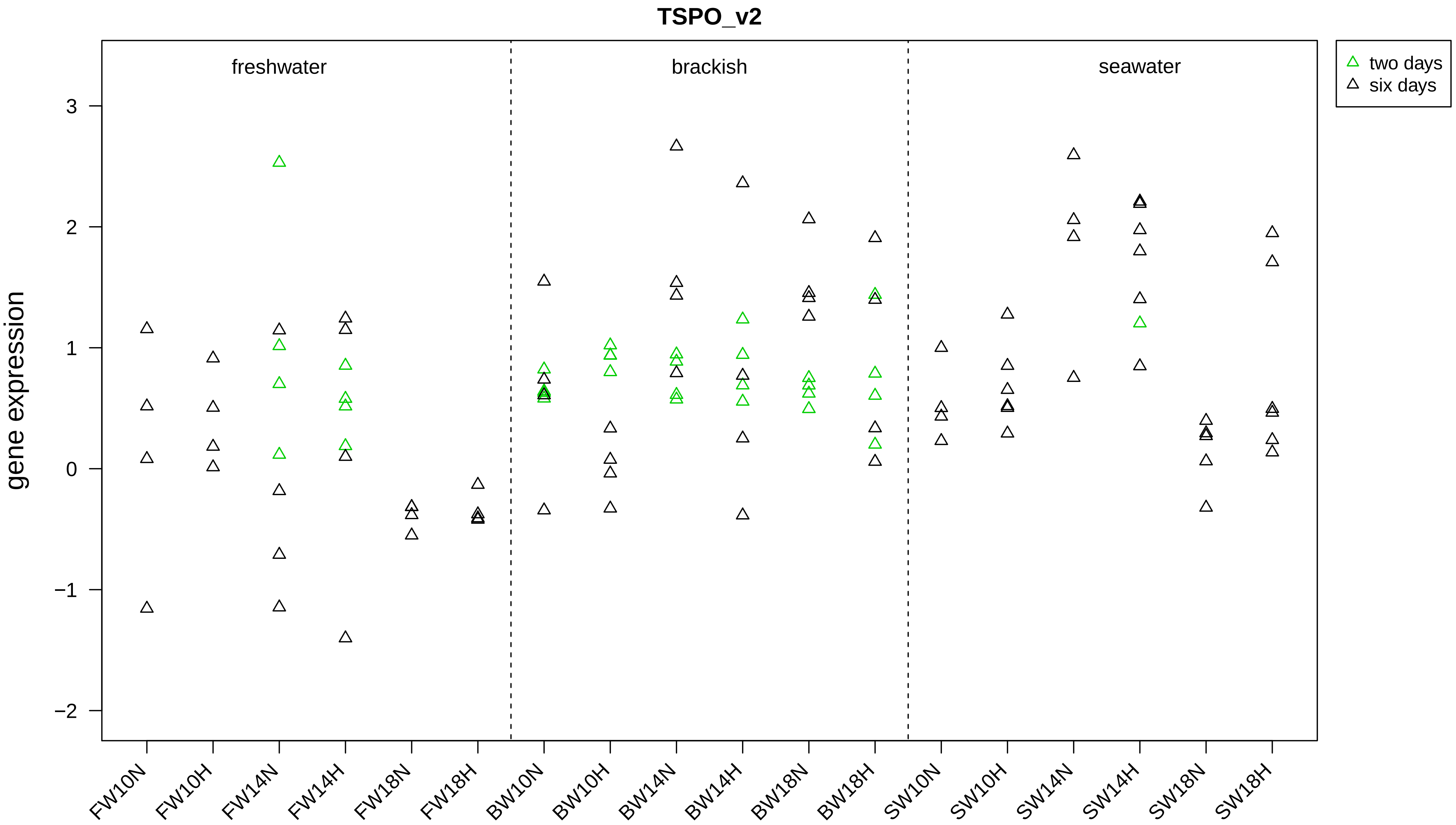

TSPO_v2
freshwater
brackish
seawater
two days
six days
gene expression
3
2
1
0
−1
−2
FW10N
FW10H
FW14N
FW14H
FW18N
FW18H
BW10N
BW10H
BW14N
BW14H
BW18N
BW18H
SW10N
SW10H
SW14N
SW14H
SW18N
SW18H

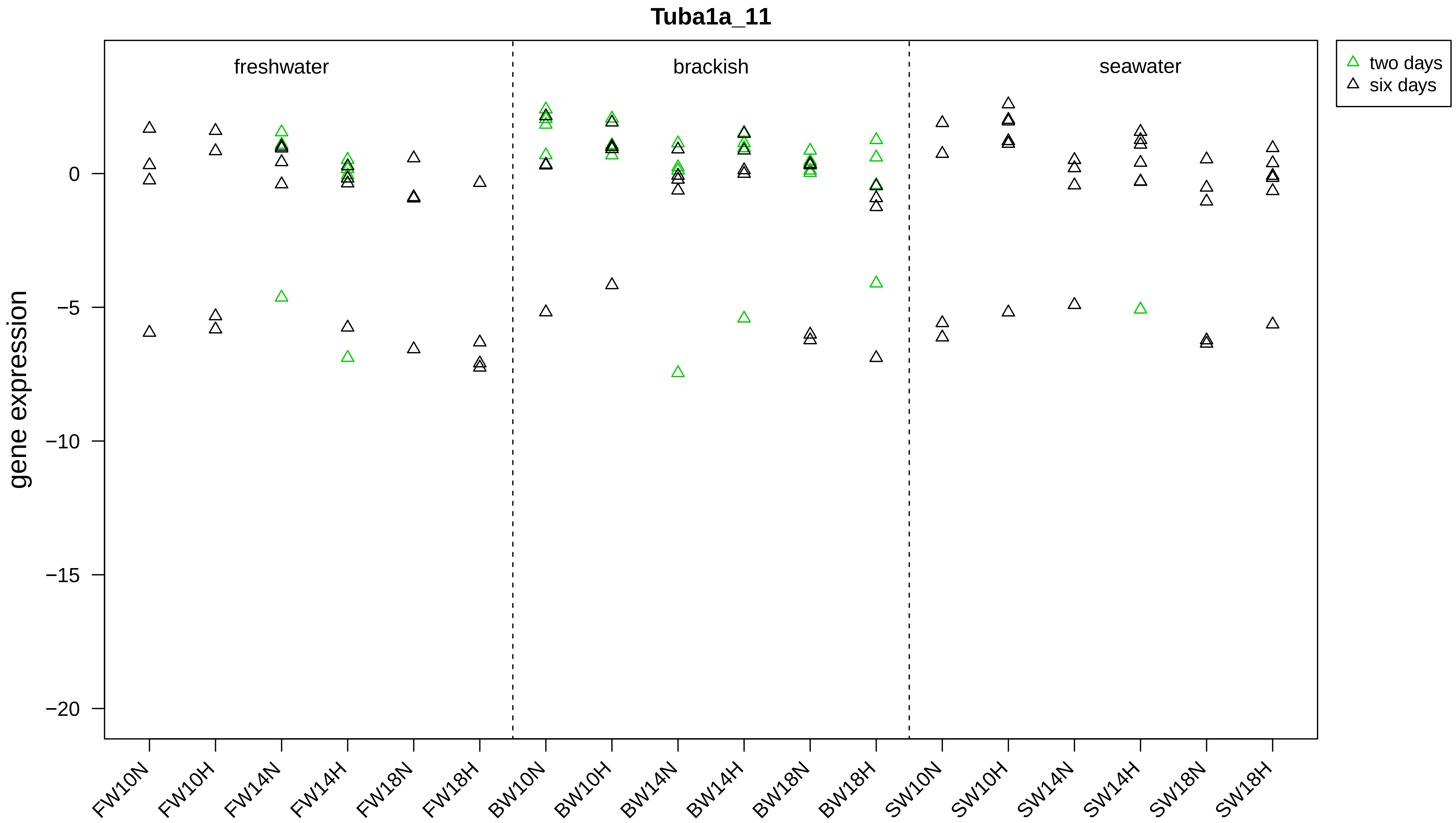

Tuba1a_11
gene expression
0
−5
−10
−15
−20
freshwater
brackish
seawater
two days
six days
FW10N
FW10H
FW14N
FW14H
FW18N
FW18H
BW10N
BW10H
BW14N
BW14H
BW18N
BW18H
SW10N
SW10H
SW14N
SW14H
SW18N
SW18H

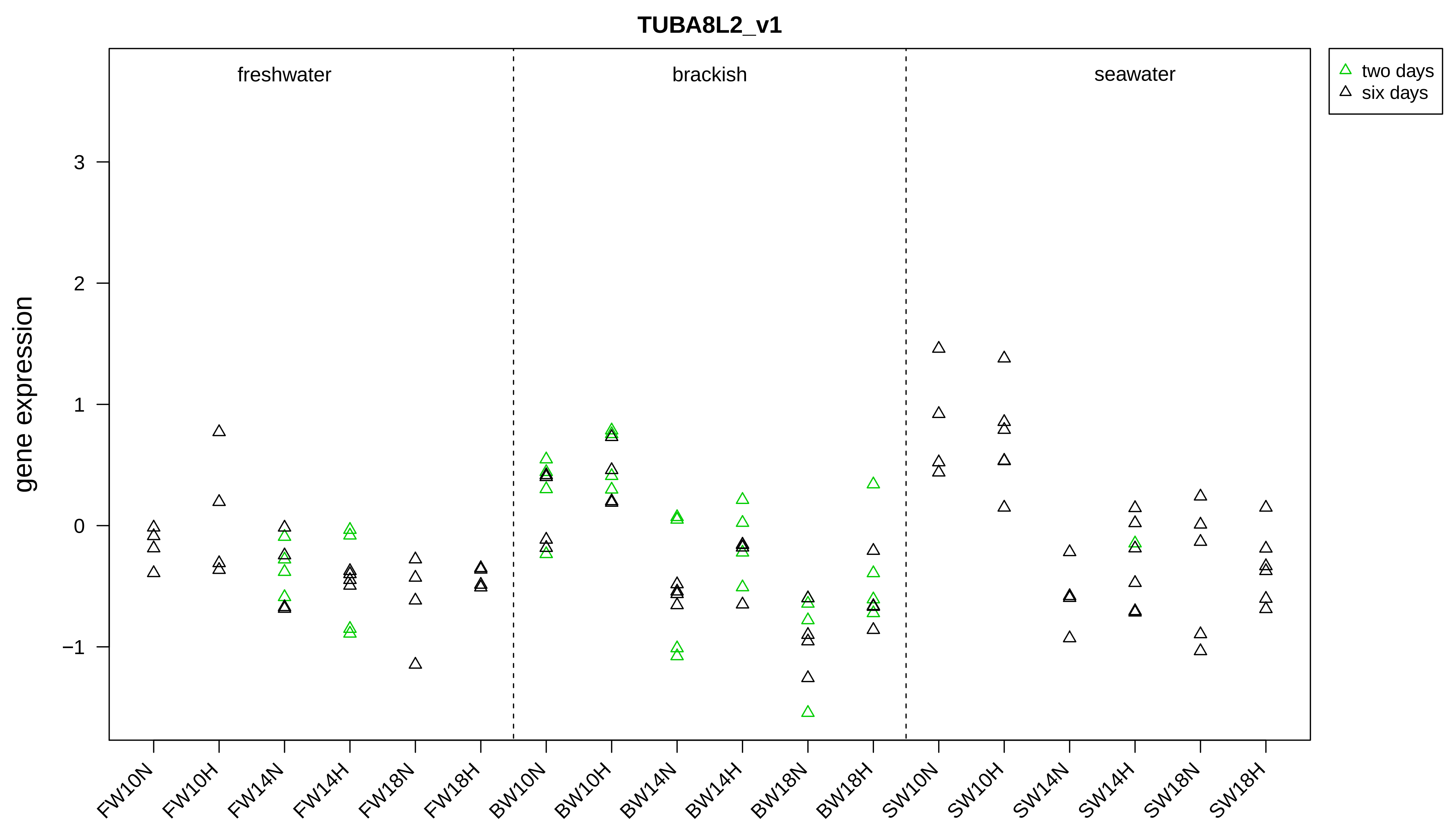

TUBA8L2_v1
freshwater
brackish
seawater
two days
six days
gene expression
3
2
1
0
−1
FW10N
FW10H
FW14N
FW14H
FW18N
FW18H
BW10N
BW10H
BW14N
BW14H
BW18N
BW18H
SW10N
SW10H
SW14N
SW14H
SW18N
SW18H

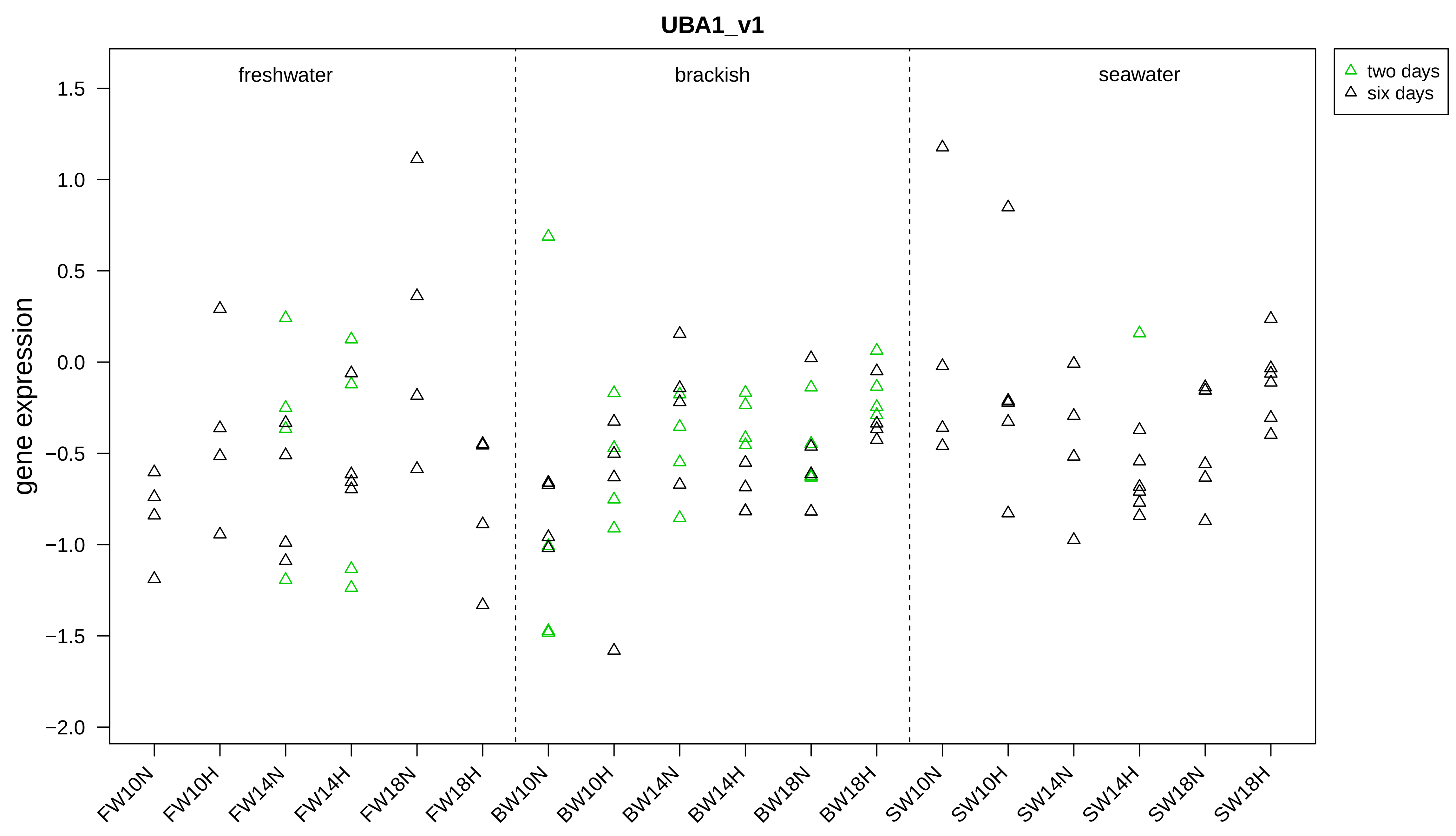

UBA1_v1
gene expression
freshwater
brackish
seawater
two days
six days
1.5
1.0
0.5
0.0
−0.5
−1.0
−1.5
−2.0
FW10N
FW10H
FW14N
FW14H
FW18N
FW18H
BW10N
BW10H
BW14N
BW14H
BW18N
BW18H
SW10N
SW10H
SW14N
SW14H
SW18N
SW18H

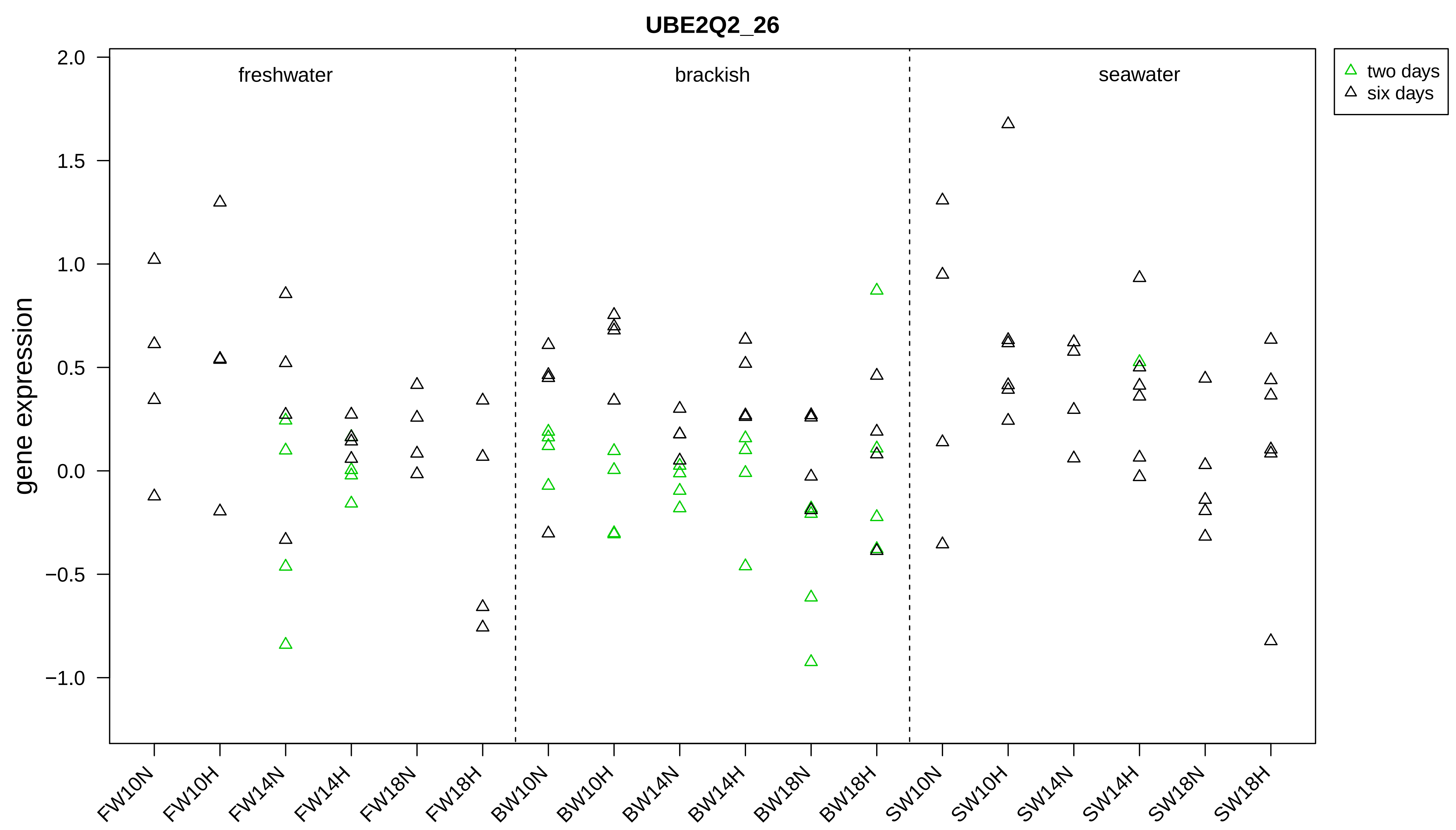

UBE2Q2_26
gene expression
freshwater
brackish
seawater
two days
six days
2.0
1.5
1.0
0.5
0.0
−0.5
−1.0
FW10N
FW10H
FW14N
FW14H
FW18N
FW18H
BW10N
BW10H
BW14N
BW14H
BW18N
BW18H
SW10N
SW10H
SW14N
SW14H
SW18N
SW18H

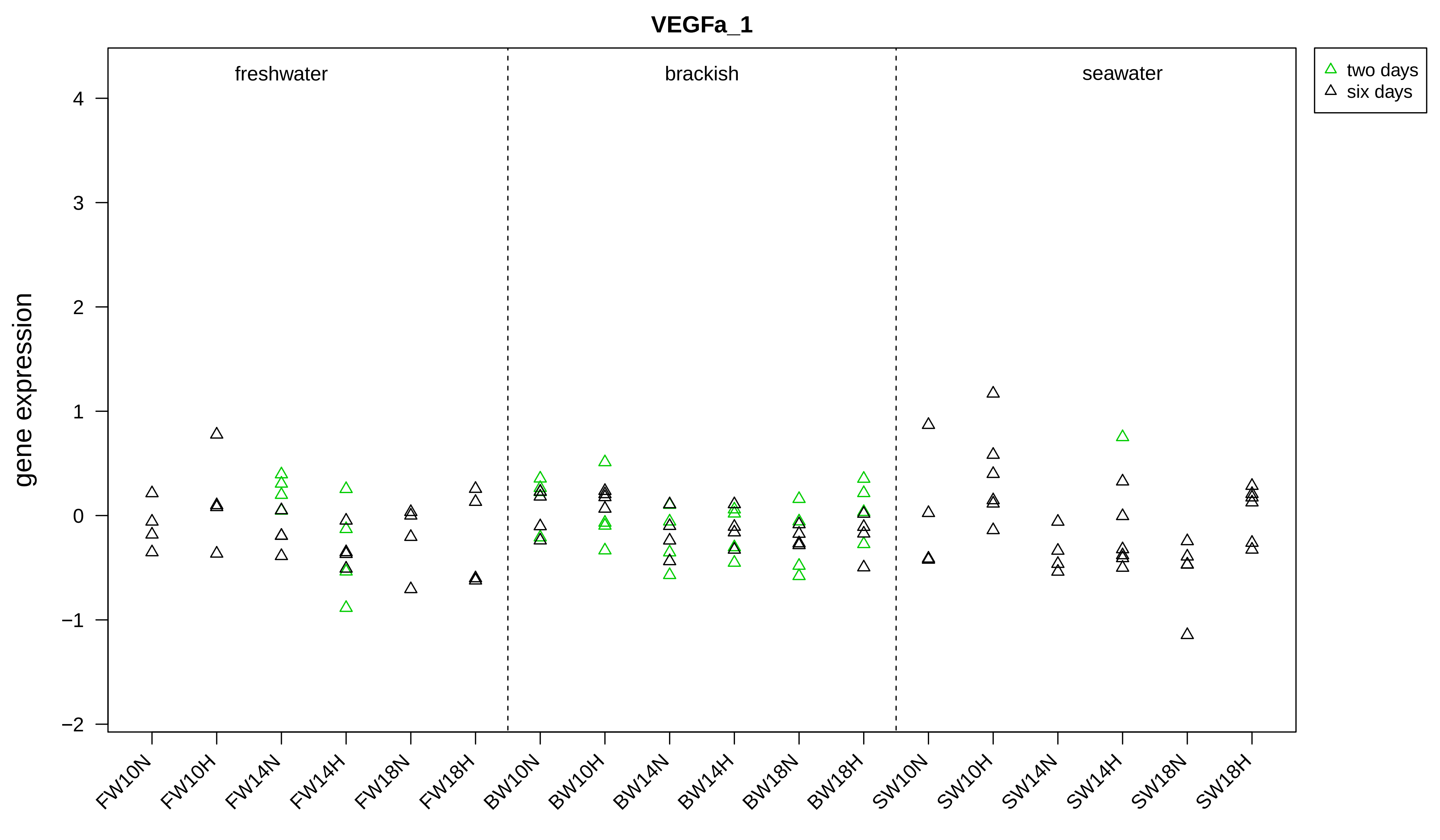

VEGFa_1
freshwater
brackish
seawater
two days
six days
gene expression
4
3
2
1
0
−1
−2
FW10N
FW10H
FW14N
FW14H
FW18N
FW18H
BW10N
BW10H
BW14N
BW14H
BW18N
BW18H
SW10N
SW10H
SW14N
SW14H
SW18N
SW18H

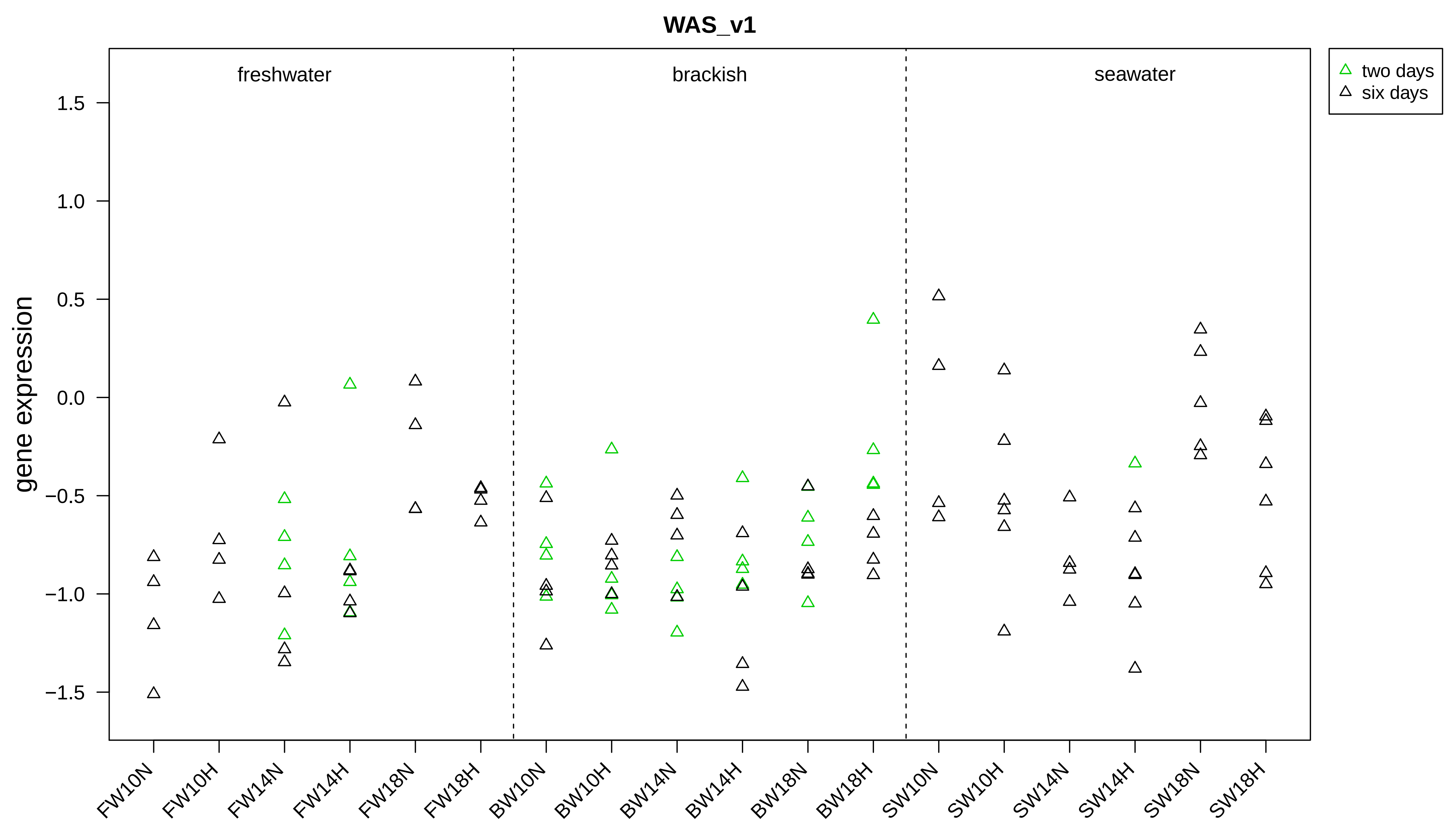

WAS_v1
freshwater
brackish
seawater
two days
six days
gene expression
1.5
1.0
0.5
0.0
−0.5
−1.0
−1.5
FW10N
FW10H
FW14N
FW14H
FW18N
FW18H
BW10N
BW10H
BW14N
BW14H
BW18N
BW18H
SW10N
SW10H
SW14N
SW14H
SW18N
SW18H

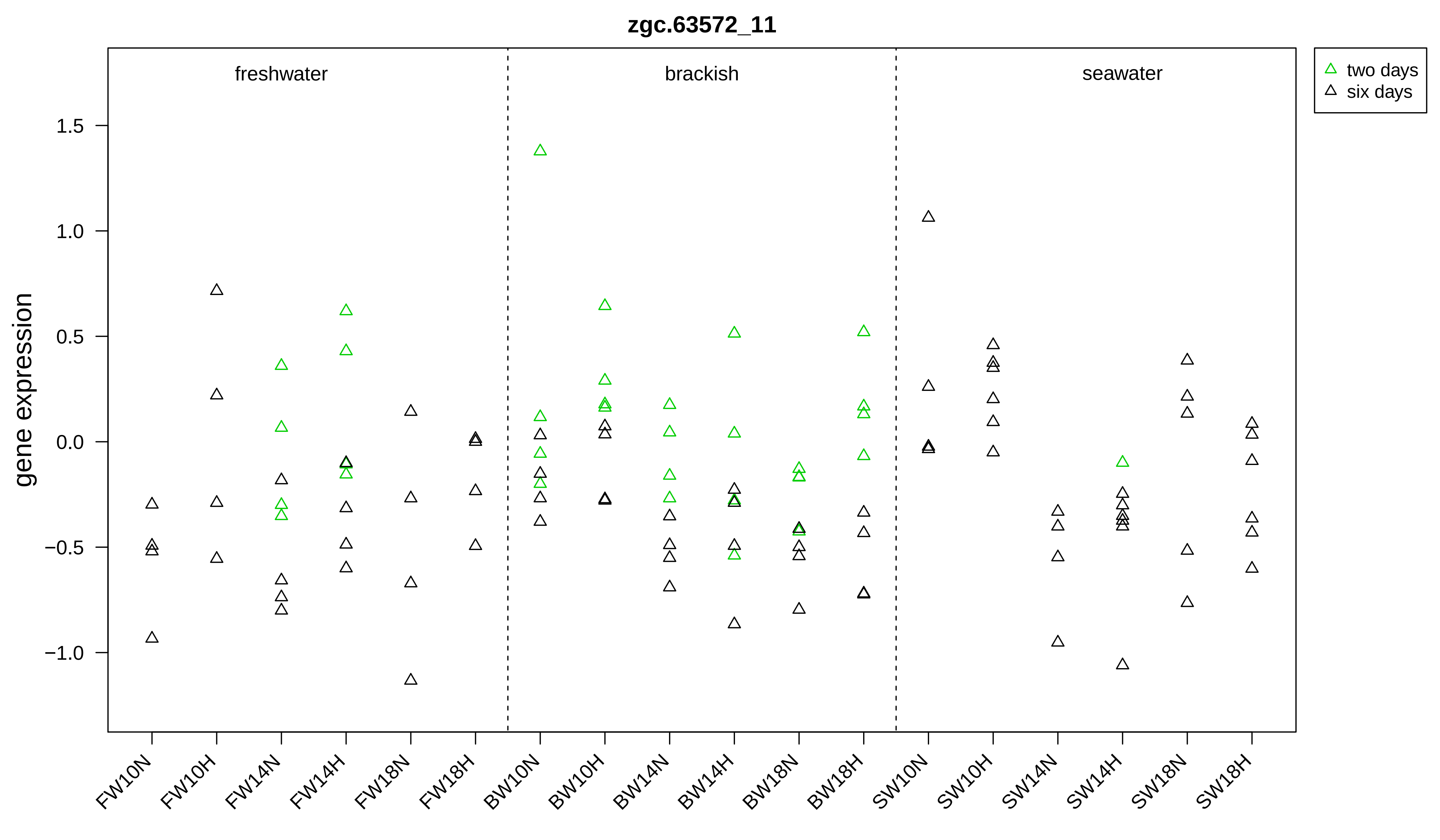

zgc.63572_11
freshwater
brackish
seawater
two days
six days
gene expression
1.5
1.0
0.5
0.0
−0.5
−1.0
FW10N
FW10H
FW14N
FW14H
FW18N
FW18H
BW10N
BW10H
BW14N
BW14H
BW18N
BW18H
SW10N
SW10H
SW14N
SW14H
SW18N
SW18H

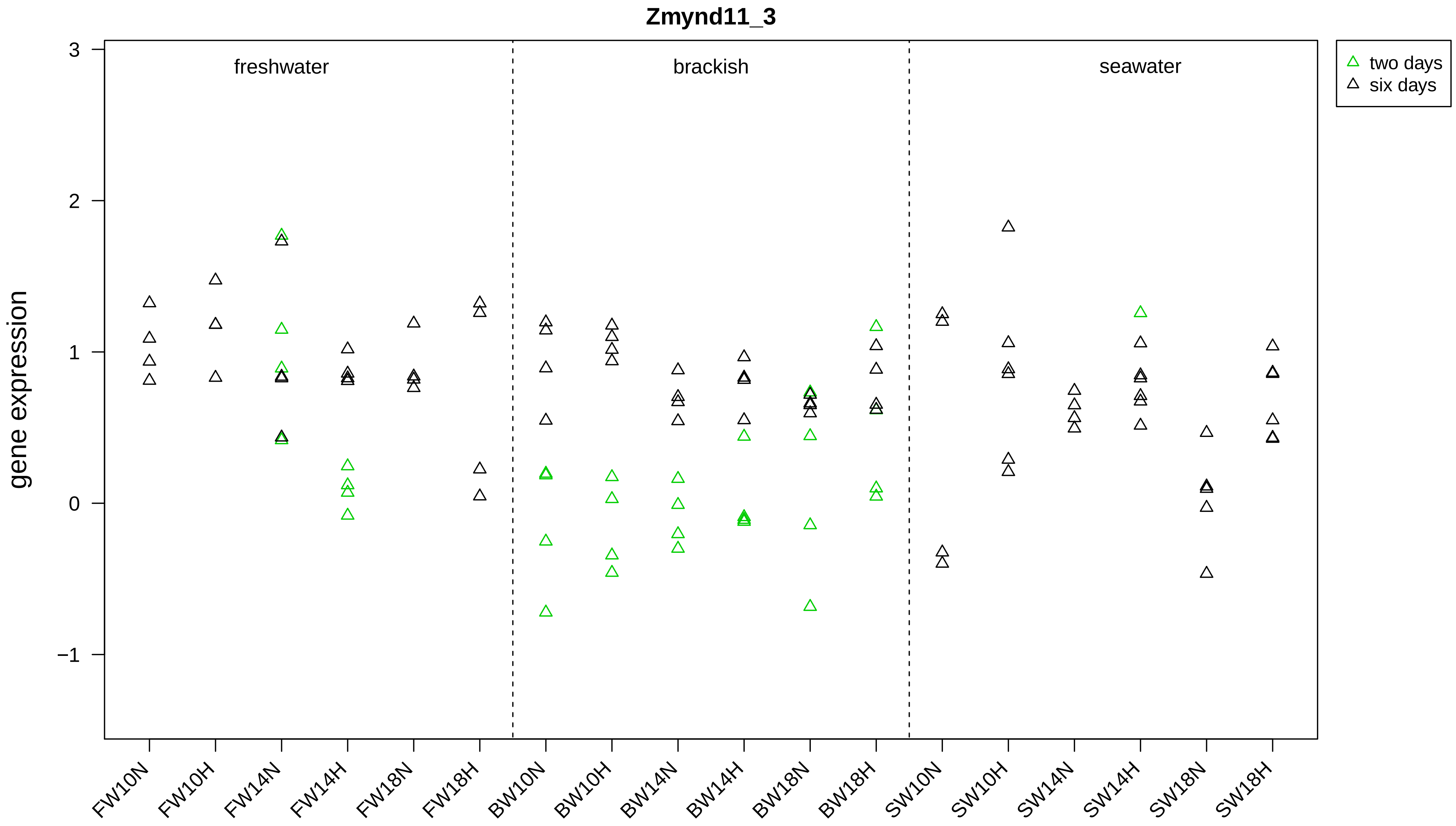

Zmynd11_3
freshwater
brackish
seawater
two days
six days
gene expression
3
2
1
0
−1
FW10N
FW10H
FW14N
FW14H
FW18N
FW18H
BW10N
BW10H
BW14N
BW14H
BW18N
BW18H
SW10N
SW10H
SW14N
SW14H
SW18N
SW18H
