## Supplementary material for "Salmonid gene expression biomarkers indicative of physiological responses to changes in salinity, temperature, but not dissolved oxygen"

Appendix 3. Treatment and mortality effects for physiological and body variables.

*Effects on behaviour*

Relative startle response was influenced by salinity for all four trials. Visually juveniles from freshwater were swimming faster after disturbance than seawater, with brackish in between. Visual gill ventilation rate scores of trial 2 were correlated with video values of the remaining trials (trial 1 Spearman's $rho = 0.70$, $p < 0.001$, trial 3 $rho = 0.65$, $p < 0.001$, and trial 4 $rho = 0.66$, $p < 0.001$). Trial date had an influence on gill ventilation rate (ANOVA, $p < 0.001$), with a higher ventilation rate for trial 3 than 1 or 4 (mean ± SE, trial 1 = 91 ± 2, trial 3 = 133 ± 3, trial 4 = 96 ± 2 gill ventilations per minute); this was likely an effect from the two different trained observers. After accounting for these differences (individual value – mean value for the trial), model selection detected primarily oxygen followed by temperature effects for gill ventilation rate ($p < 0.001$ for both). Juveniles from hypoxia had higher ventilation rate than normoxia. Juveniles from 14 and 18°C had higher ventilation rate than 10°C (Figure S1).

*Effects on $Na^+/K^+$-ATPase activity and plasma variables*

Salinity effects were detected for $Na^+/K^+$-ATPase (NKA) activity and chloride concentrations after six days for live juveniles using model selection (ANOVA, $p < 0.001$ and $p = 0.007$). Juveniles from seawater had higher NKA activity and chloride than freshwater with brackish in between (Figure S2). Temperature effects were detected for lactate and glucose concentrations ($p = 0.029$ and $p < 0.001$). Juveniles from 18°C had higher lactate and lower glucose than 10°C with 14°C in between. There was also a temperature trend for NKA activity ($p = 0.060$). Juveniles from 18°C had a trend for higher NKA activity than 10°C with 14°C in between. Within seawater, there were differences in NKA activity, chloride, and lactate between moribund or dead and live juveniles, but not glucose. NKA activity was lower and chloride and lactate were higher for moribund or dead than live juveniles (Table S3). Within brackish, there were differences in glucose and chloride, but not ATPase and lactate. Within freshwater, there was difference in glucose only. Similar to seawater, in brackish chloride was higher for moribund or dead than live juveniles. For brackish and freshwater, glucose was lower for moribund or dead than live juveniles. No oxygen effects were detected for ATPase or the plasma variables.

*Effects on body size, skin pigmentation, and body morphology*

After accounting for the differences in the initial values for the trials, across the trials treatment effects were detected by model selection for all body variables (individual values – mean of initial values for the trial), with exception of caudal fin darkness and posterior brightness  (Figure S3, S4, and S5). Temperature influenced body length, mass, condition, elongation, roundness, and caudal fin yellowness (ANOVA, $p < 0.012$ for all). Juveniles from 18°C had lower body mass, condition, and elongation, and higher back roundness than 10°C; juveniles from 14°C were generally in between. Juveniles from 14°C had lower body length than 10°C and caudal fin yellowness than 18°C. Salinity influenced caudal peduncle length ($p < 0.001$) and body thickness ($p = 0.027$). Juveniles from seawater had lower caudal peduncle length and body thickness than freshwater and brackish (thickness brackish trend $p = 0.071$). Additionally, there was a secondary temperature effect for caudal peduncle length ($p = 0.002$). Juveniles from 14 and 18°C had longer caudal peduncles than 10°C. Oxygen influenced anterior brightness ($p = 0.039$). Juveniles from hypoxia had higher anterior brightness than normoxia.

Comparing the average tank values within each trial, moribund or dead juveniles had lower body length, mass, condition, elongation, and thickness than live juveniles (Table S4). Moribund or dead juveniles also had rounder backs and more yellow caudal fins than live individuals. There were no differences in remaining skin pigmentation and body morphology variables. Albeit the moribund and dead sample sizes for freshwater and brackish were considerably smaller than for seawater.

Table S3. Summary of gill $Na^+/K^+$-ATPase activity and plasma variables for mortality and salinity.

Presented are the mean ± SD. Bold values represent significant differences (Student's t-tests, $p <$ 0.05) between live and distressed (moribund or dead) juveniles. $Na^+/K^+$-ATPase activity units are µmol ADP (mg protein)$^{-1}$ h$^{-1}$.

| Variable | Live | Distressed |
|---|---|---|
| $Na^+/K^+$-ATPase activity | | |
| freshwater | 4.5 ± 2.3 | 5.3 ± 3.6 |
| brackish | 5.7 ± 2.6 | 6.3 ± 3.7 |
| seawater | **8.3 ± 3.4** | **5.7 ± 3.7** |
| Chloride (mmol L$^{-1}$) | | |
| freshwater | 115.7 ± 14.5 | 73.8 ± 75.1 |
| brackish | **124.9 ± 24.7** | **206.1 ± 21.8** |
| seawater | **138.6 ± 25.8** | **205.8 ± 58.8** |
| Lactate (mmol L$^{-1}$) | | |
| freshwater | 3.2 ± 2.1 | 5.9 ± 6.1 |
| brackish | 2.4 ± 1.0 | 4.0 ± 2.6 |
| seawater | **2.4 ± 1.2** | **9.1 ± 6.4** |
| Glucose (mmol L$^{-1}$) | | |
| freshwater | **3.9 ± 2.7** | **1.6 ± 1.3** |
| brackish | **3.5 ± 1.6** | **1.5 ± 1.2** |
| seawater | 3.4 ± 1.5 | 2.6 ± 4.4 |

Table S4. Summary of body variables for mortality.

Presented are the mean ± SD. Bold values represent significant differences (paired Student's t-tests, $p < 0.05$) between live and distressed (moribund or dead) juveniles of the same tank for each trial.

| Body variable | live | distressed |
| --- | --- | --- |
| Size | | |
| length (cm) | **7.7 ± 1.9** | **7.3 ± 1.9** |
| mass (g) | **5.53 ± 3.35** | **4.41 ± 2.92** |
| condition (100 × mass ÷ length$^3$) | **1.05 ± 0.05** | **0.95 ± 0.10** |
| Skin pigmentation | | |
| anterior brightness (PC1) | -0.8 ± 7.5 | -1.3 ± 9.3 |
| caudal fin darkness (PC2) | -3.5 ± 3.8 | -1.4 ± 7.9 |
| posterior brightness (PC3) | 0.6 ± 3.5 | 1.5 ± 6.3 |
| caudal fin yellowness (PC4) | **-2.5 ± 2.2** | **-1.0 ± 5.2** |
| Morphology | | |
| elongation (RW1) | **-0.0013 ± 0.0105** | **-0.0045 ± 0.0109** |
| back roundness (RW2) | **-0.0001 ± 0.0039** | **0.0058 ± 0.0076** |
| caudal peduncle length (RW4) | 0.0015 ± 0.0033 | 0.0030 ± 0.0060 |
| thickness (RW5) | **-0.0012 ± 0.0033** | **-0.0084 ± 0.0070** |

**Figure legends**

Figure S1. Beanplot of the gill ventilation rate after four days.

Because of the difference average rates among trials, the data are corrected by taking the values and subtracting the mean value of the trial. Bold horizontal line is the mean and smaller horizontal lines are for three individuals per tank and trial. Salinity symbols are SW, BW, and FW for seawater, brackish, and freshwater for groups. Temperature treatment symbols are 10, 14, and 18 for °C groups. Dissolved oxygen treatment symbols are N for normoxia and H for hypoxia groups. For trial 2 (24 June), data from one FW10N and BW14N replicate tank are missing because of low video quality. Different letters represent Tukey's test significant contrasts ($p < 0.05$): capital letters represent dissolved oxygen contrasts and miniscule letters represent temperature contrasts. Note that the video was not set-up for trial 2 (late May, smolt), so no data are available.

Figure S2. Beanplots of gill $Na^+/K^+$-ATPase activity and plasma variables and after six days of treatment. Bold horizontal line is the mean and smaller horizontal lines are for the individuals from the trials combined. Salinity treatment symbols are SW, BW, and FW for seawater, brackish, and freshwater groups. Temperature treatment symbols are 10, 14, and 18 for °C groups. Different letters represent Tukey's test significant contrasts among salinities or temperatures defined in the x-axis ($p < 0.05$).

Figure S3. Beanplots of body length, mass, and condition.

Displayed are the data by initial trial date (left panel) and any detected salinity, temperature, or dissolved oxygen effects from model selection (right panel). Because of the influence of trial date on body variables, right panel data are corrected by taking the individual values and subtracting the mean of the initial values of the trial. Bold horizontal line is the mean. Different letters represent Tukey's test significant contrasts for groups ($p < 0.05$).

Figure S4. Beanplots of four skin pigmentation variables.

See Figure S3 legend.

Figure S5. Beanplots of four body morphology variables.

See Figure S3 legend.

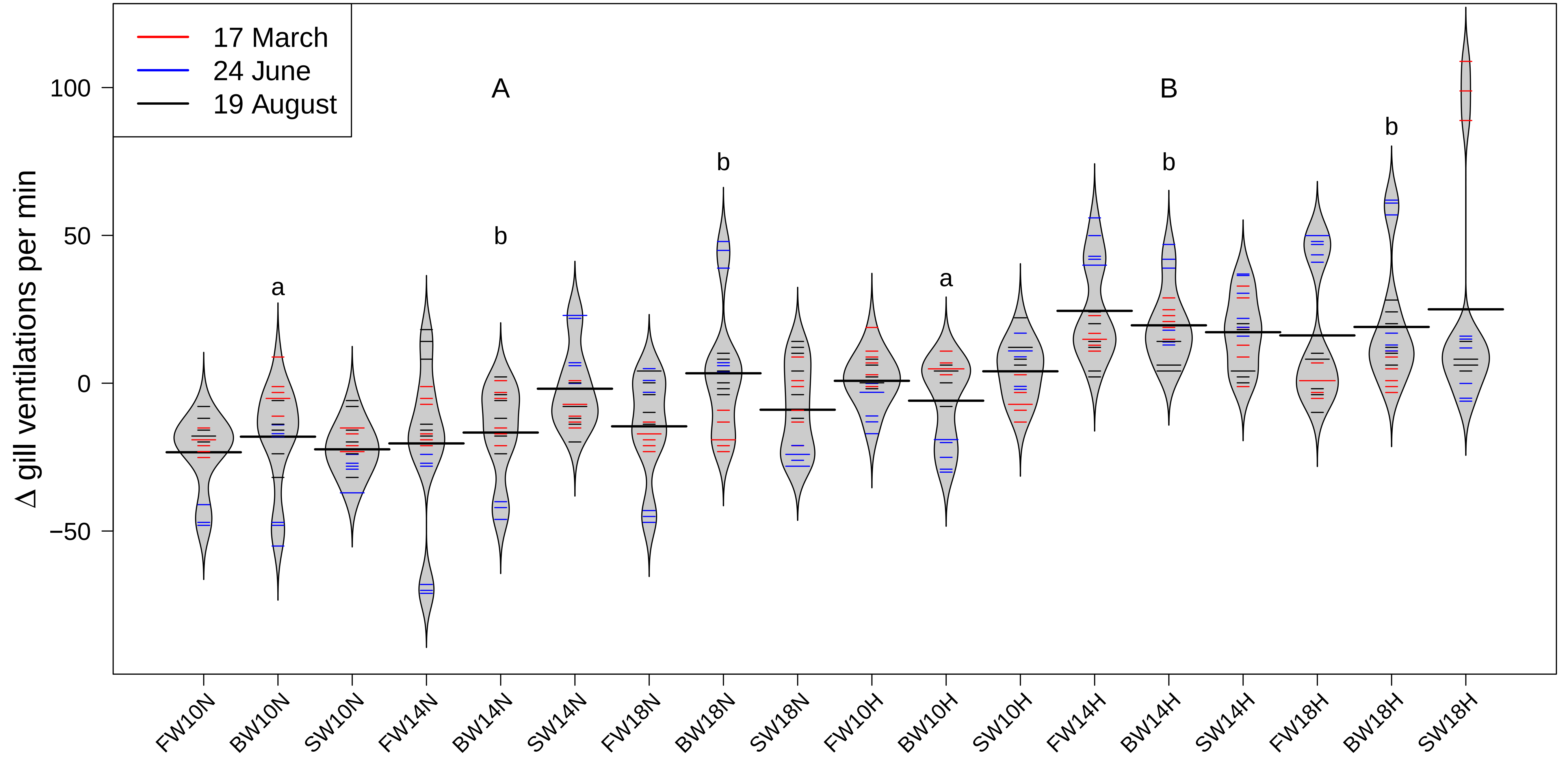

17 March
24 June
19 August
A
B
a
b
b
b
a
b
b
Δ gill ventilations per min
100
50
0
−50
FW10N
BW10N
SW10N
FW14N
BW14N
SW14N
FW18N
BW18N
SW18N
FW10H
BW10H
SW10H
FW14H
BW14H
SW14H
FW18H
BW18H
SW18H

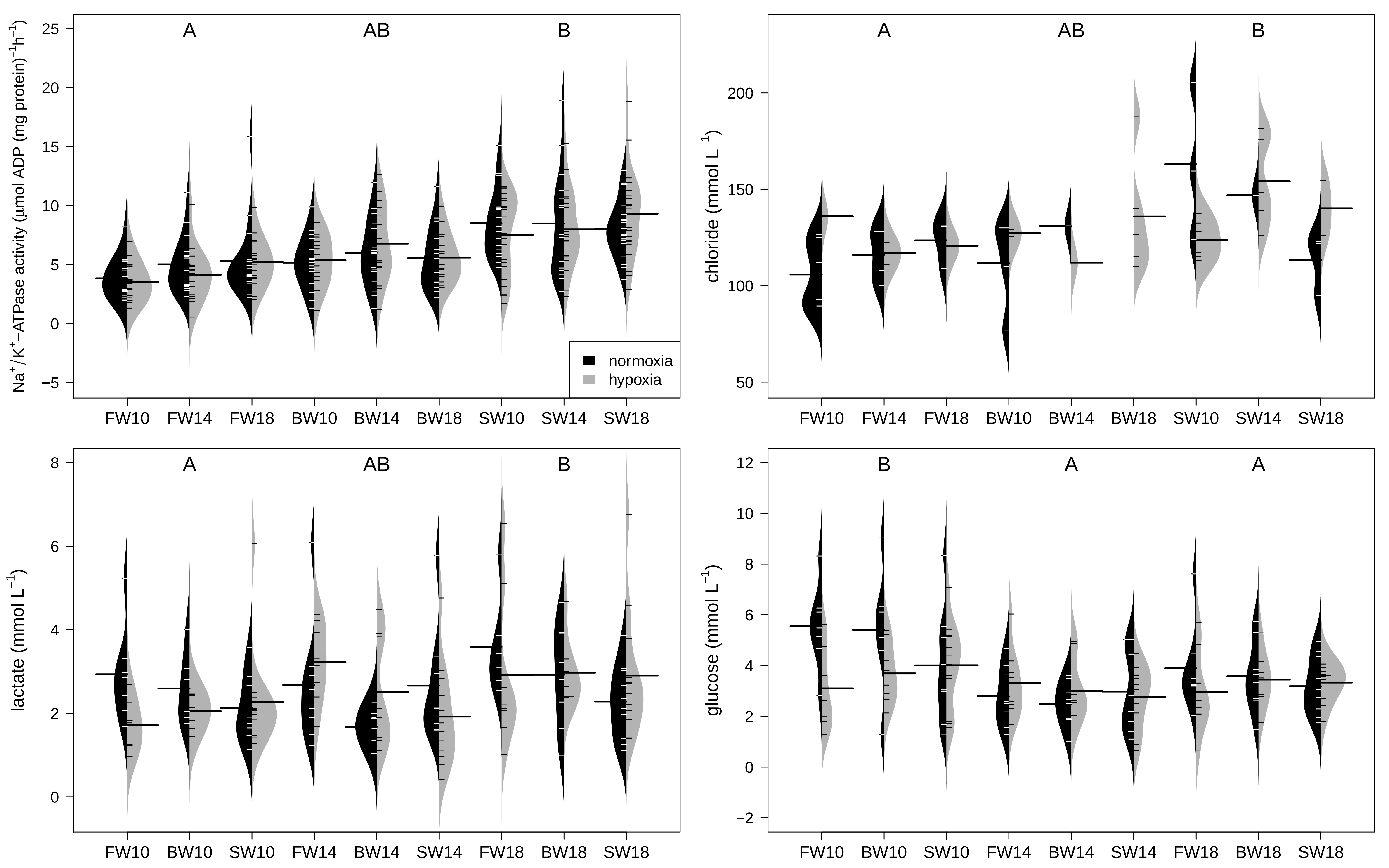

Na+/K+−ATPase activity (μmol ADP (mg protein)−1h−1)
A
AB
B
normoxia
hypoxia
FW10
FW14
FW18
BW10
BW14
BW18
SW10
SW14
SW18
chloride (mmol L−1)
lactate (mmol L−1)
glucose (mmol L−1)

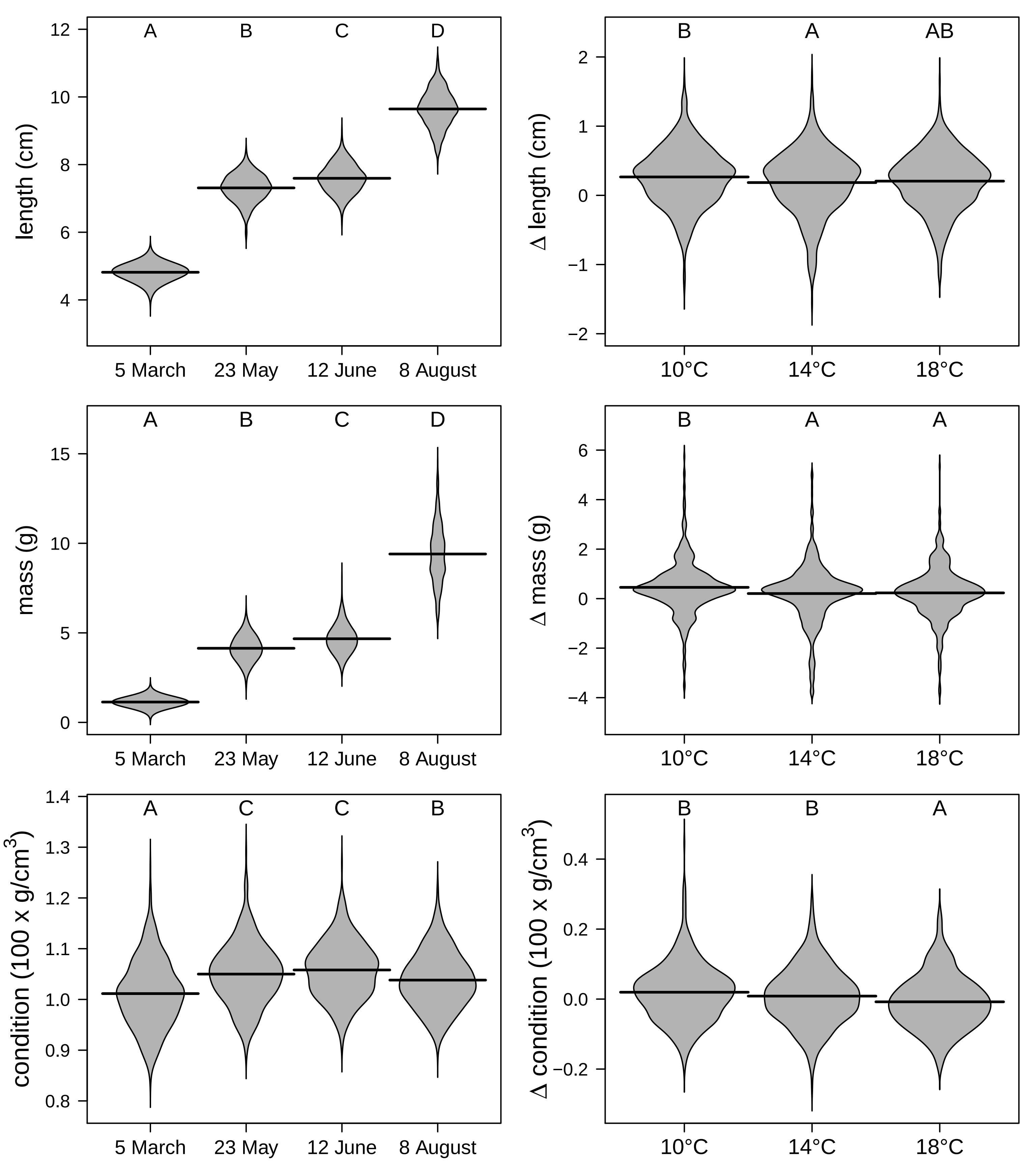

A
B
C
D
length (cm)
5 March
23 May
12 June
8 August
B
A
AB
Δ length (cm)
10°C
14°C
18°C
A
B
C
D
mass (g)
5 March
23 May
12 June
8 August
B
A
A
Δ mass (g)
10°C
14°C
18°C
A
C
C
B
condition (100 x g/cm3)
5 March
23 May
12 June
8 August
B
B
A
Δ condition (100 x g/cm3)
10°C
14°C
18°C

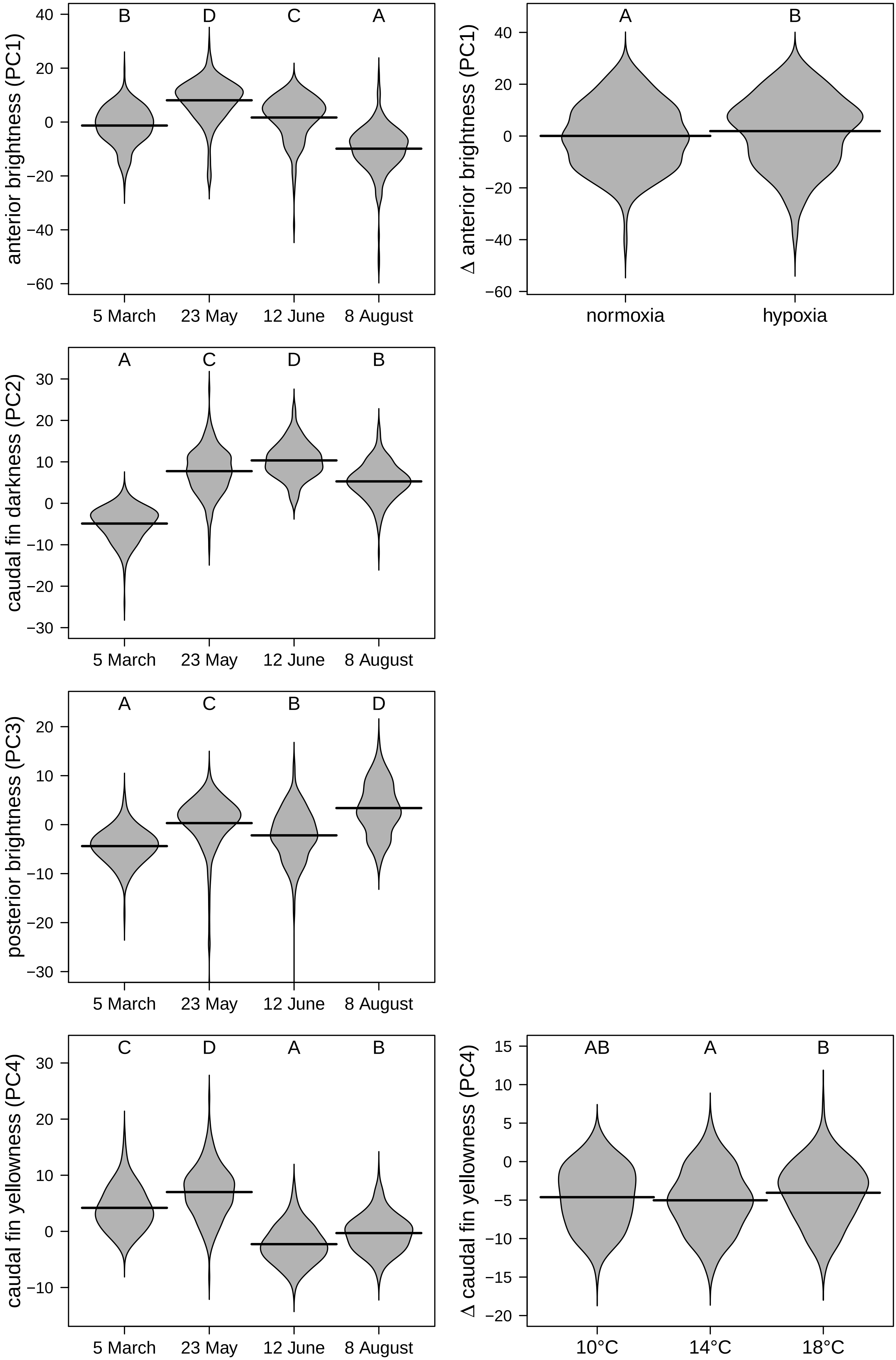

anterior brightness (PC1)
B
D
C
A
5 March
23 May
12 June
8 August
Δ anterior brightness (PC1)
A
B
normoxia
hypoxia
caudal fin darkness (PC2)
A
C
D
B
5 March
23 May
12 June
8 August
posterior brightness (PC3)
A
C
B
D
5 March
23 May
12 June
8 August
caudal fin yellowness (PC4)
C
D
A
B
5 March
23 May
12 June
8 August
Δ caudal fin yellowness (PC4)
AB
A
B
10°C
14°C
18°C

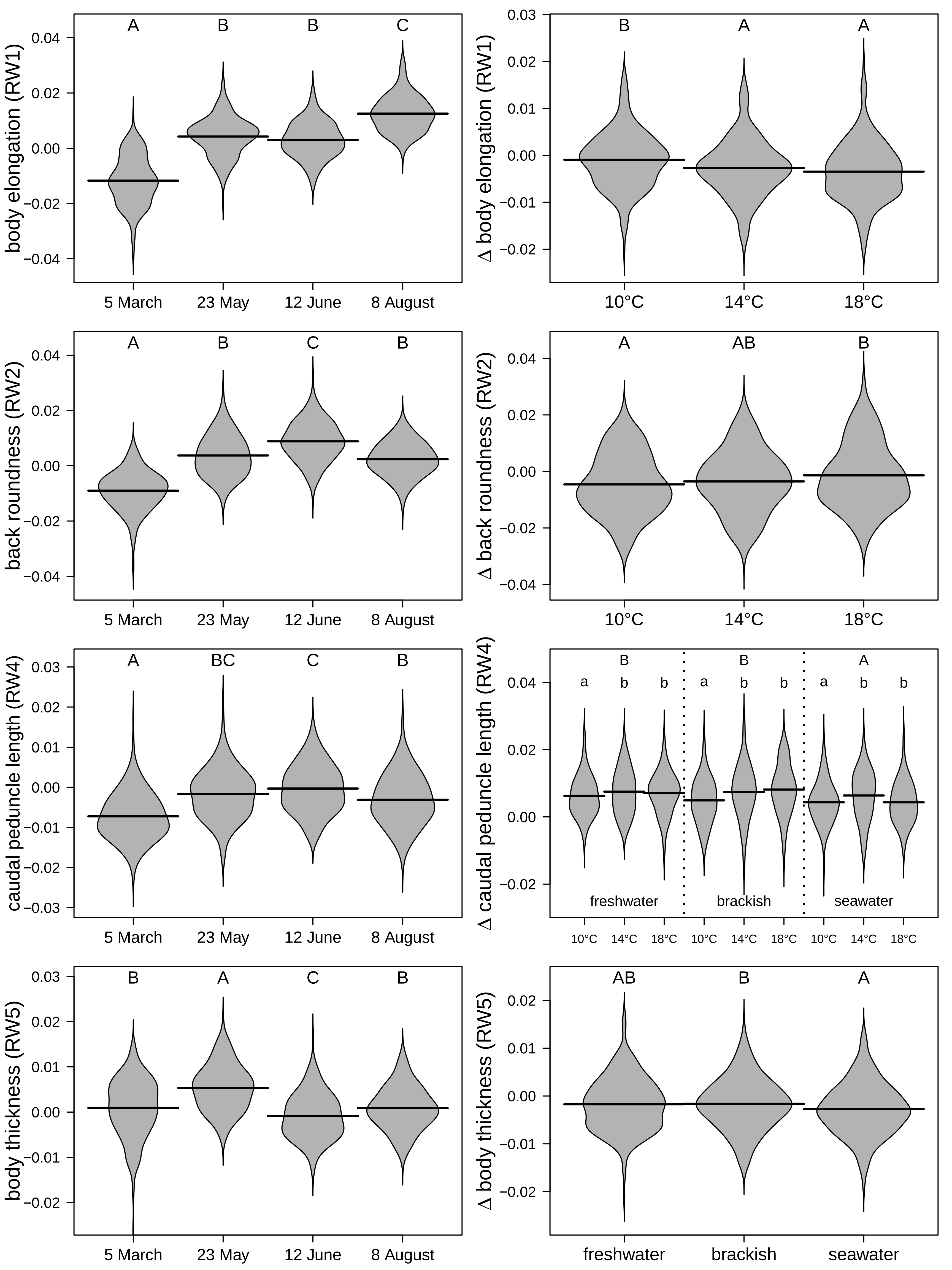
body elongation (RW1)
Δ body elongation (RW1)
back roundness (RW2)
Δ back roundness (RW2)
caudal peduncle length (RW4)
Δ caudal peduncle length (RW4)
body thickness (RW5)
Δ body thickness (RW5)
5 March
23 May
12 June
8 August
10°C
14°C
18°C
freshwater
brackish
seawater
A
B
C
AB
BC
a
b
