## Supplementary material for "Salmonid gene expression biomarkers indicative of physiological responses to changes in salinity, temperature, but not dissolved oxygen"

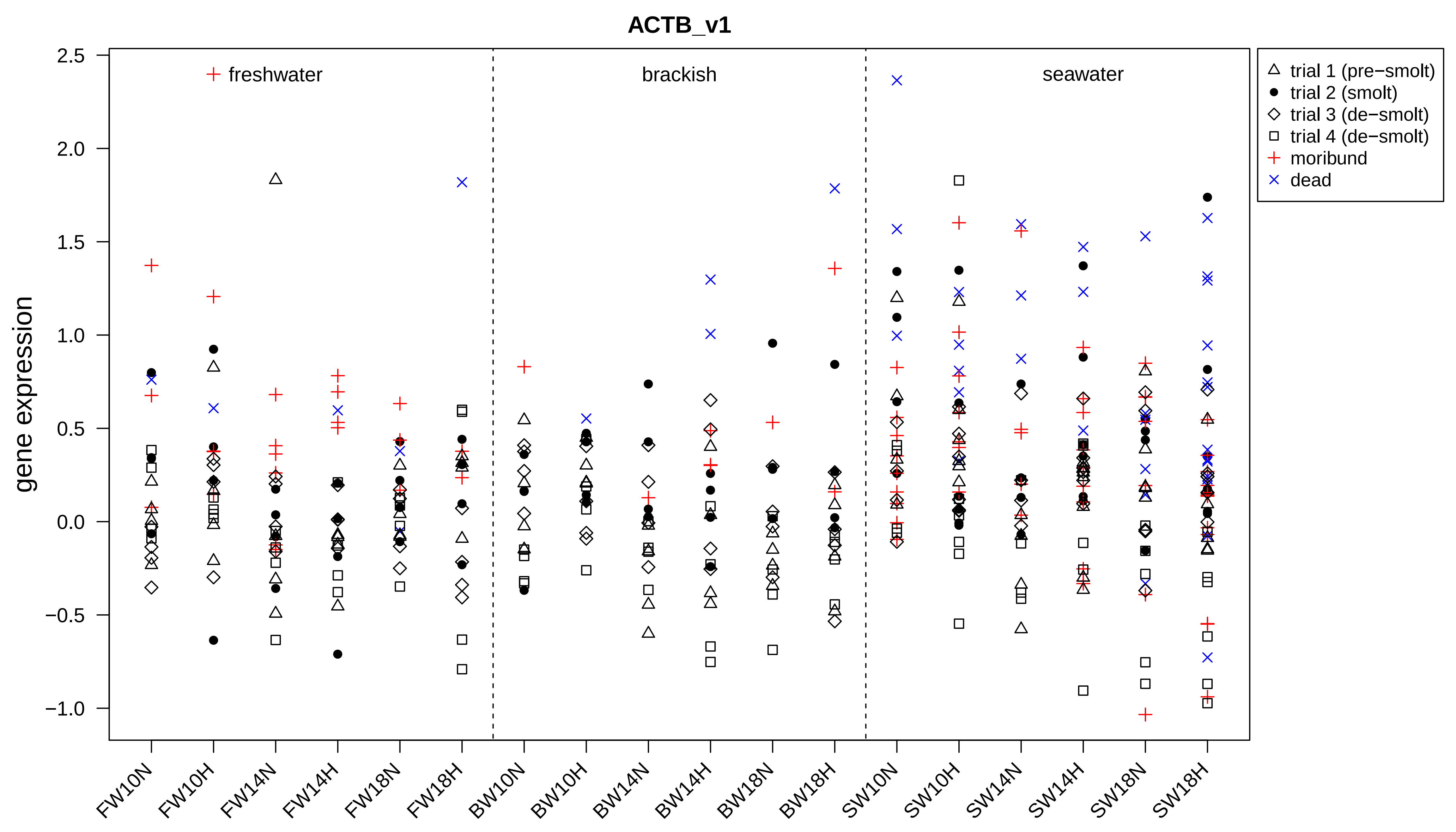

ACTB_v1
gene expression
2.5
2.0
1.5
1.0
0.5
0.0
−0.5
−1.0
freshwater
brackish
seawater
trial 1 (pre−smolt)
trial 2 (smolt)
trial 3 (de−smolt)
trial 4 (de−smolt)
moribund
dead
FW10N
FW10H
FW14N
FW14H
FW18N
FW18H
BW10N
BW10H
BW14N
BW14H
BW18N
BW18H
SW10N
SW10H
SW14N
SW14H
SW18N
SW18H

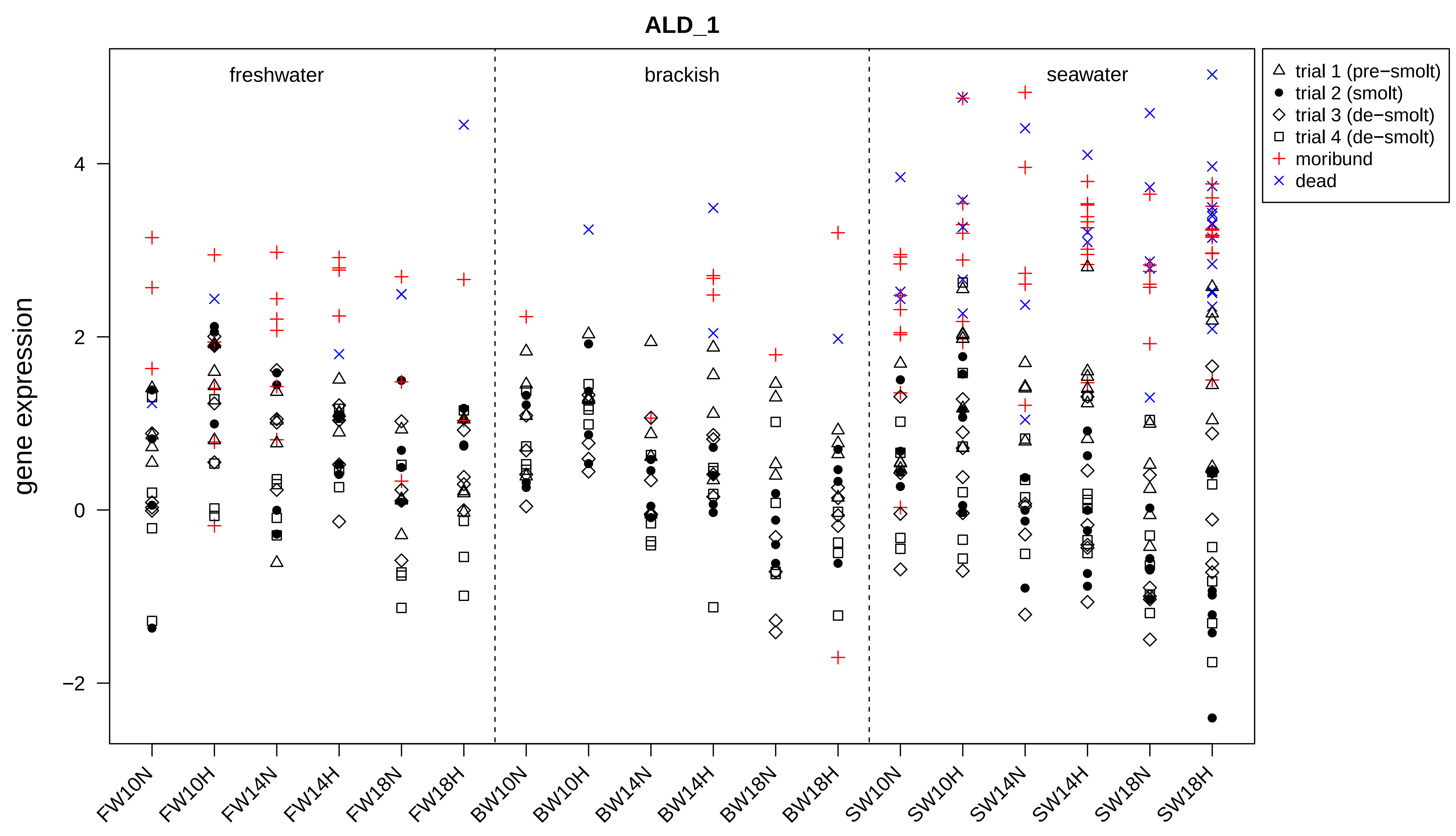
ALD_1
freshwater
brackish
seawater
trial 1 (pre−smolt)
trial 2 (smolt)
trial 3 (de−smolt)
trial 4 (de−smolt)
moribund
dead
gene expression
4
2
0
−2
FW10N
FW10H
FW14N
FW14H
FW18N
FW18H
BW10N
BW10H
BW14N
BW14H
BW18N
BW18H
SW10N
SW10H
SW14N
SW14H
SW18N
SW18H

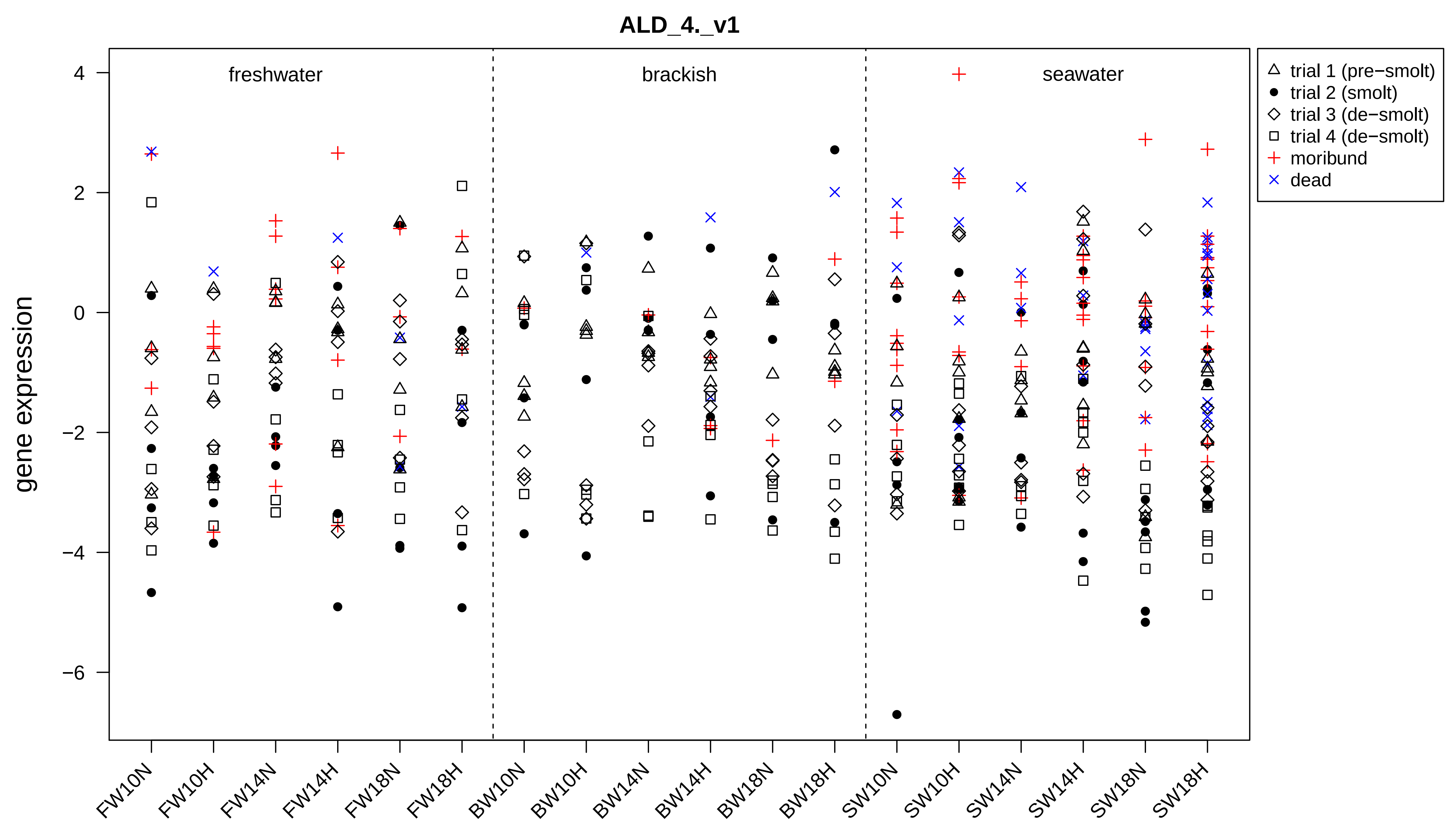
ALD_4._v1
gene expression
freshwater
brackish
seawater
trial 1 (pre−smolt)
trial 2 (smolt)
trial 3 (de−smolt)
trial 4 (de−smolt)
moribund
dead
4
2
0
−2
−4
−6
FW10N
FW10H
FW14N
FW14H
FW18N
FW18H
BW10N
BW10H
BW14N
BW14H
BW18N
BW18H
SW10N
SW10H
SW14N
SW14H
SW18N
SW18H

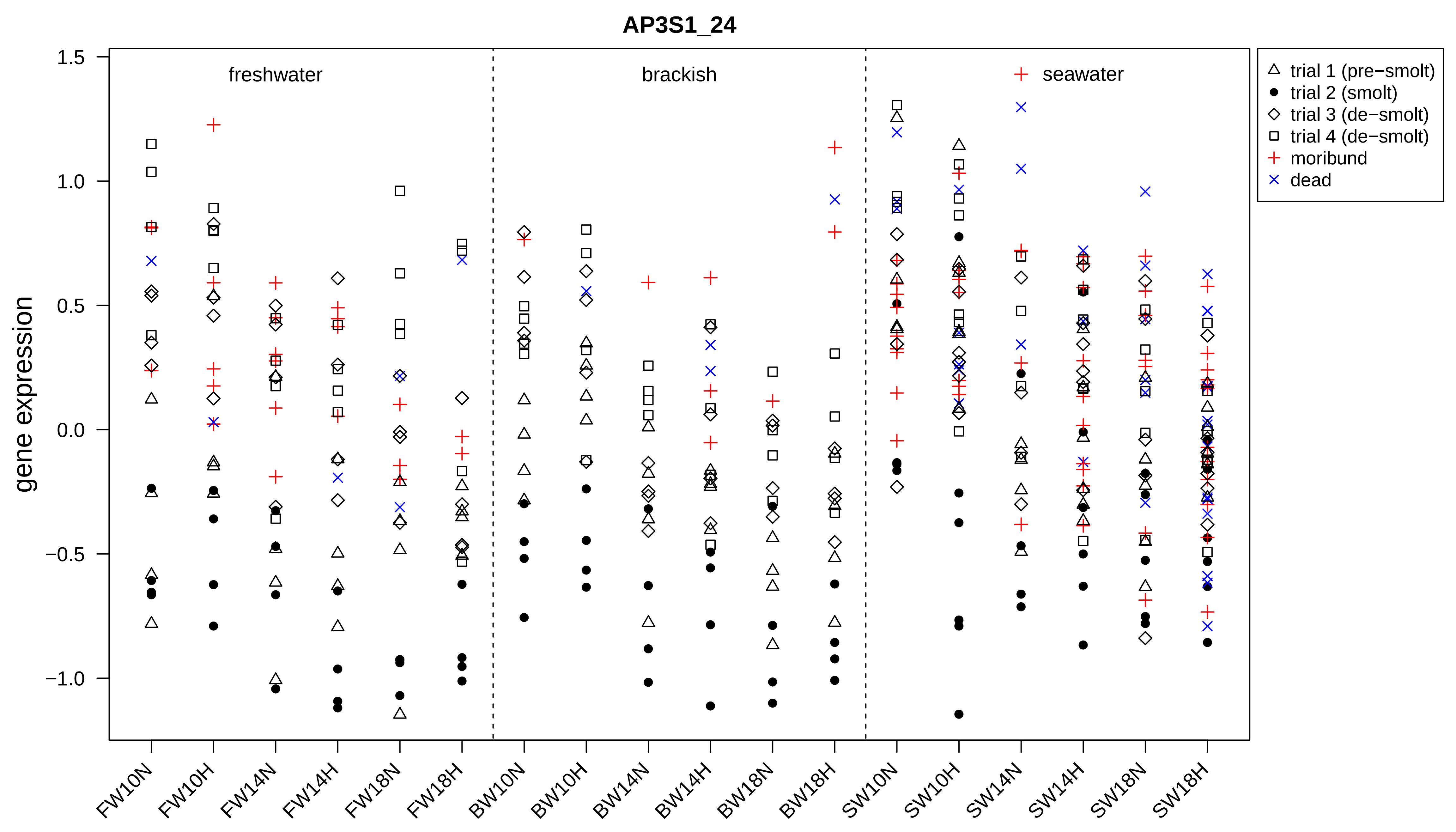

AP3S1_24
gene expression
freshwater
brackish
seawater
trial 1 (pre−smolt)
trial 2 (smolt)
trial 3 (de−smolt)
trial 4 (de−smolt)
moribund
dead
1.5
1.0
0.5
0.0
−0.5
−1.0
FW10N
FW10H
FW14N
FW14H
FW18N
FW18H
BW10N
BW10H
BW14N
BW14H
BW18N
BW18H
SW10N
SW10H
SW14N
SW14H
SW18N
SW18H

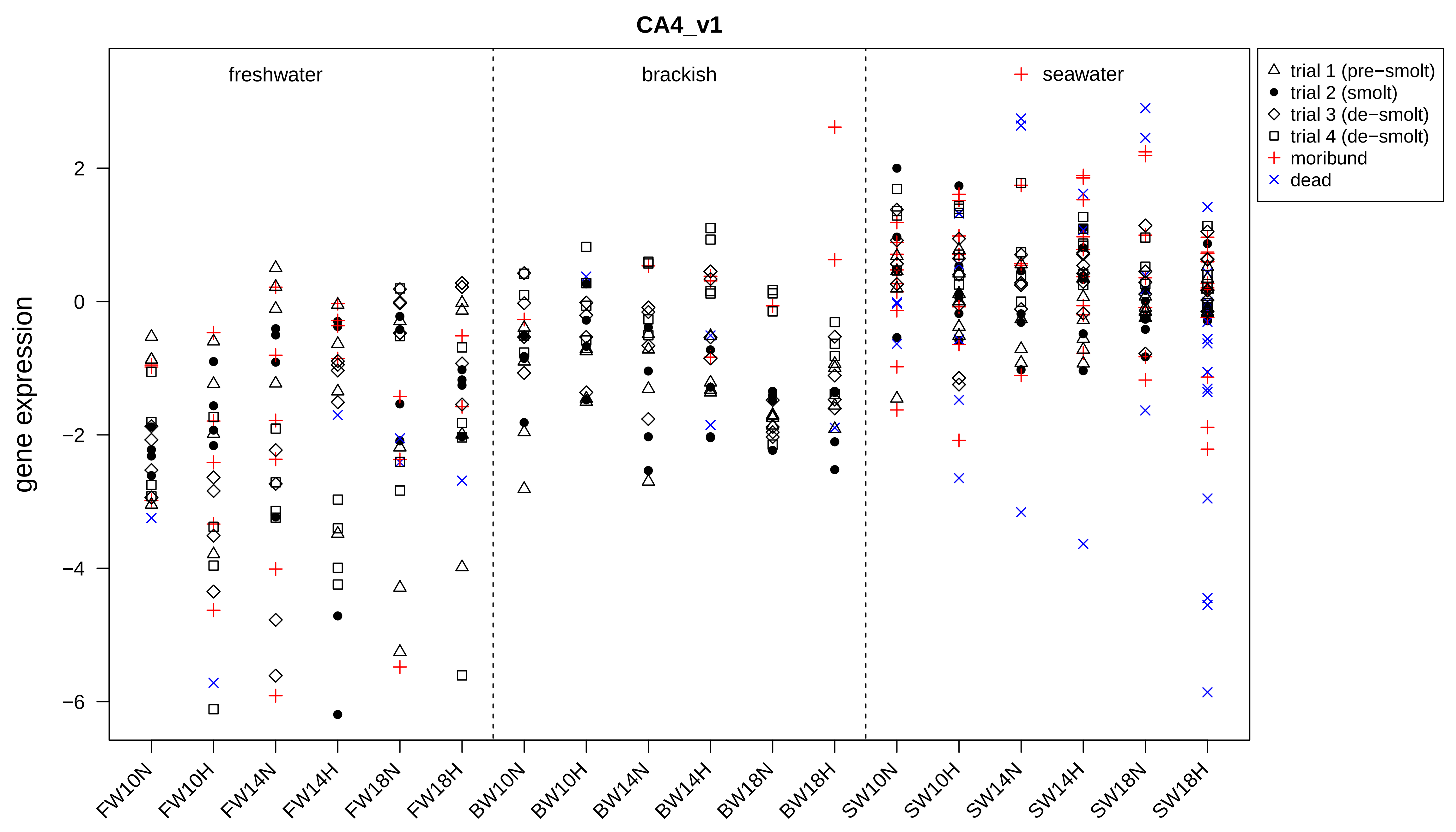

CA4_v1
freshwater
brackish
seawater
gene expression
2
0
−2
−4
−6
FW10N
FW10H
FW14N
FW14H
FW18N
FW18H
BW10N
BW10H
BW14N
BW14H
BW18N
BW18H
SW10N
SW10H
SW14N
SW14H
SW18N
SW18H
trial 1 (pre−smolt)
trial 2 (smolt)
trial 3 (de−smolt)
trial 4 (de−smolt)
moribund
dead

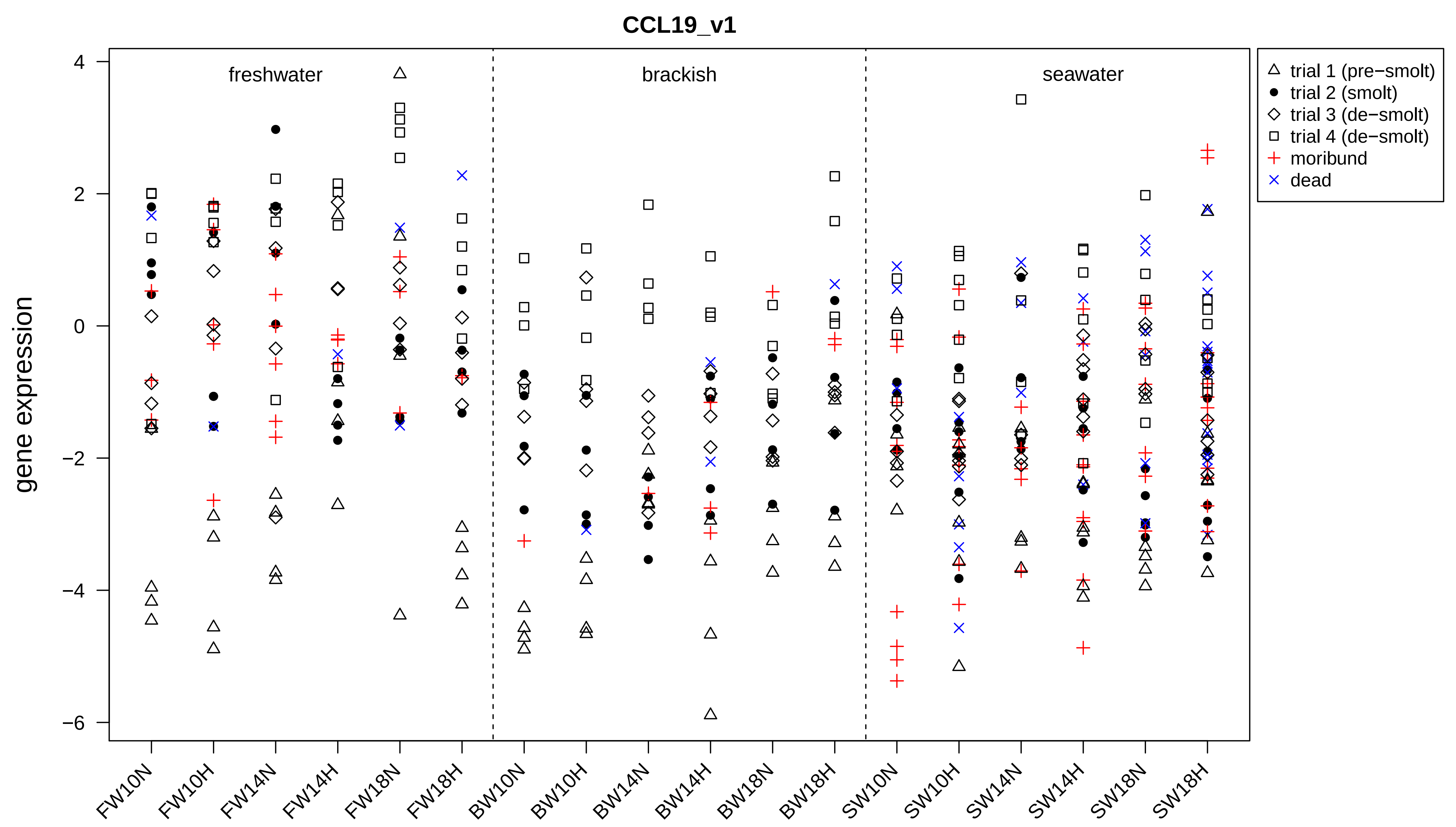

CCL19_v1
gene expression
freshwater
brackish
seawater
trial 1 (pre−smolt)
trial 2 (smolt)
trial 3 (de−smolt)
trial 4 (de−smolt)
moribund
dead
4
2
0
−2
−4
−6
FW10N
FW10H
FW14N
FW14H
FW18N
FW18H
BW10N
BW10H
BW14N
BW14H
BW18N
BW18H
SW10N
SW10H
SW14N
SW14H
SW18N
SW18H

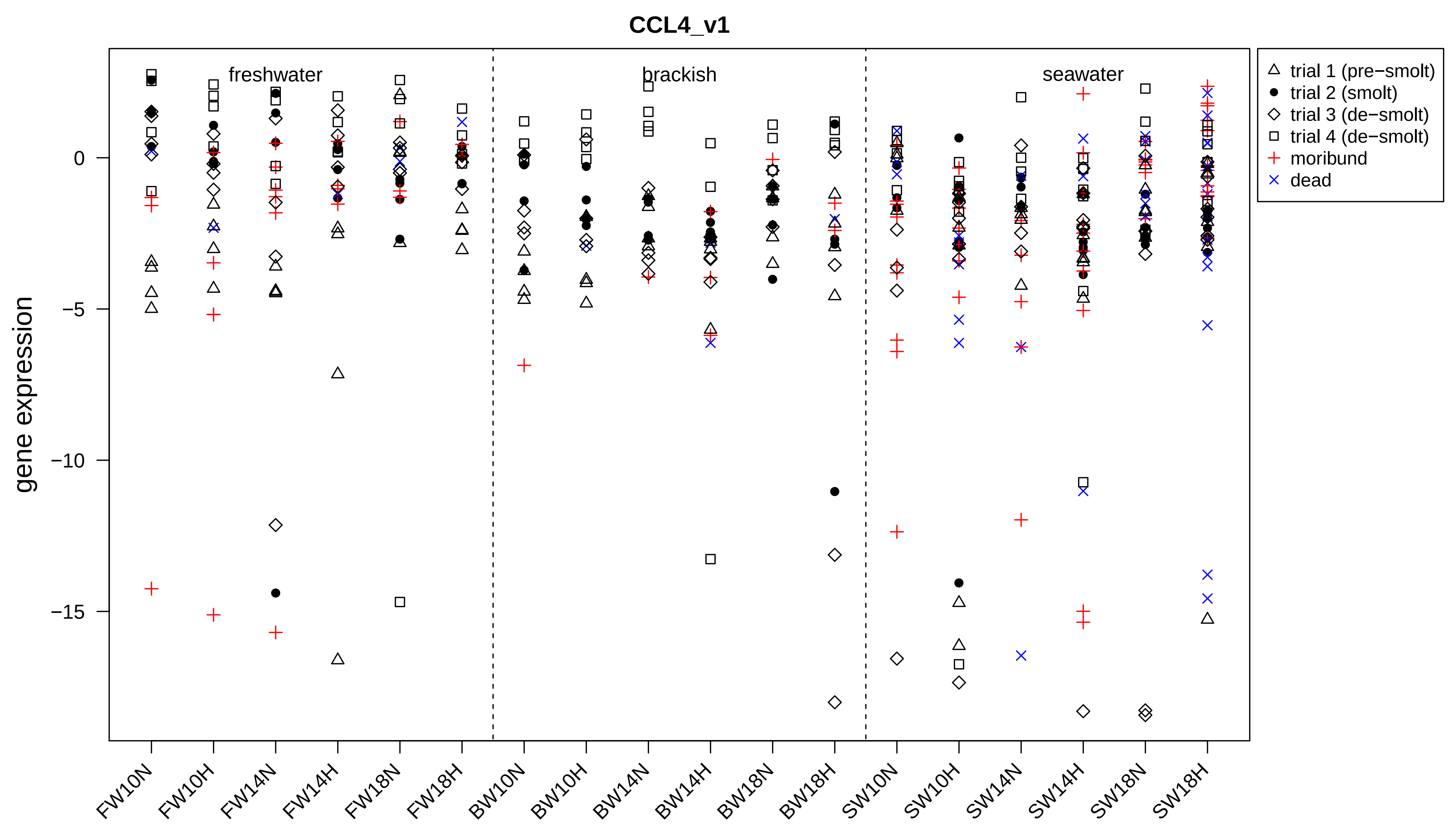
CCL4_v1
gene expression
0
−5
−10
−15
freshwater
brackish
seawater
trial 1 (pre−smolt)
trial 2 (smolt)
trial 3 (de−smolt)
trial 4 (de−smolt)
moribund
dead
FW10N
FW10H
FW14N
FW14H
FW18N
FW18H
BW10N
BW10H
BW14N
BW14H
BW18N
BW18H
SW10N
SW10H
SW14N
SW14H
SW18N
SW18H

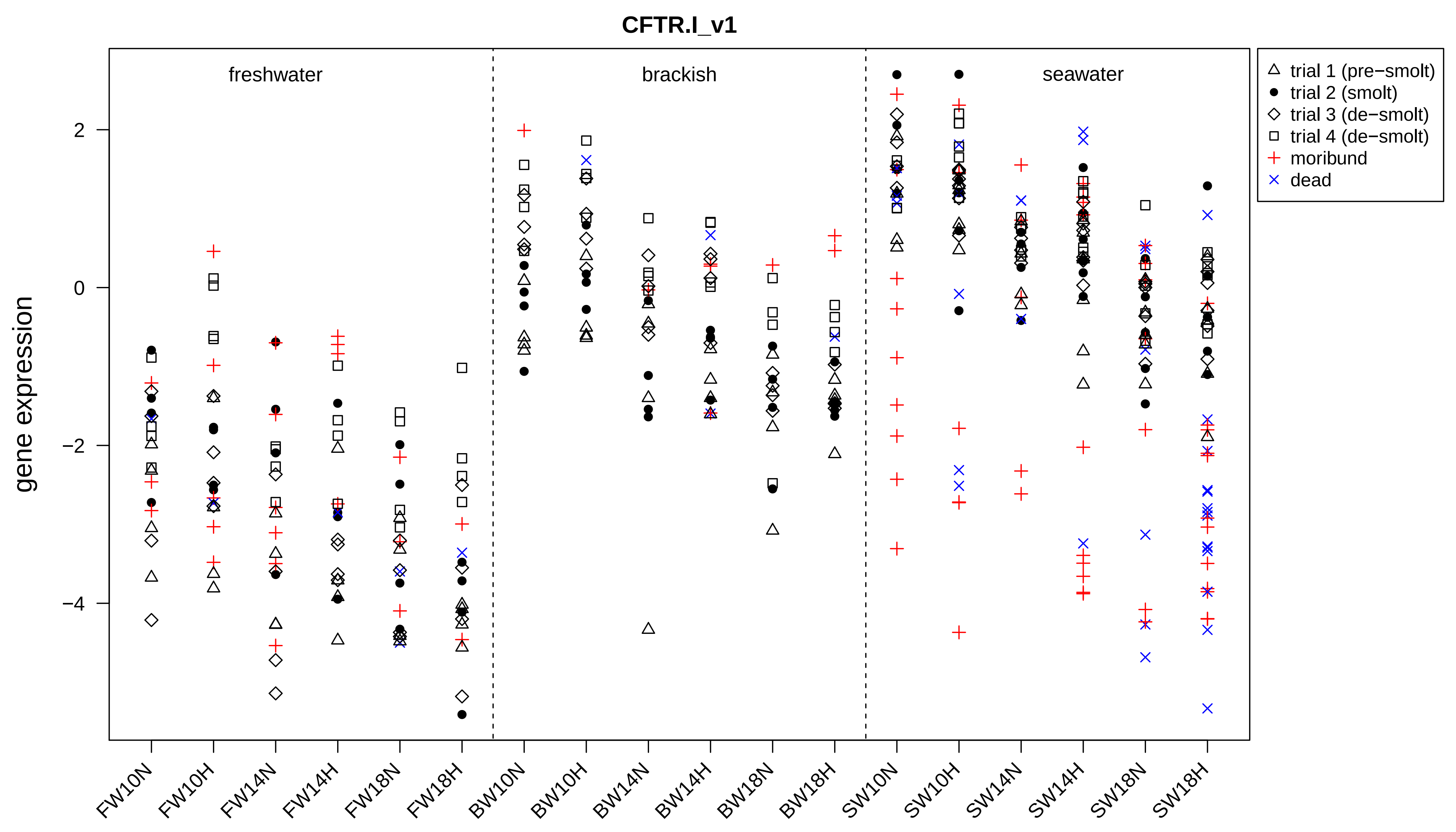

CFTR.I_v1
freshwater
brackish
seawater
gene expression
2
0
−2
−4
FW10N
FW10H
FW14N
FW14H
FW18N
FW18H
BW10N
BW10H
BW14N
BW14H
BW18N
BW18H
SW10N
SW10H
SW14N
SW14H
SW18N
SW18H
trial 1 (pre−smolt)
trial 2 (smolt)
trial 3 (de−smolt)
trial 4 (de−smolt)
moribund
dead

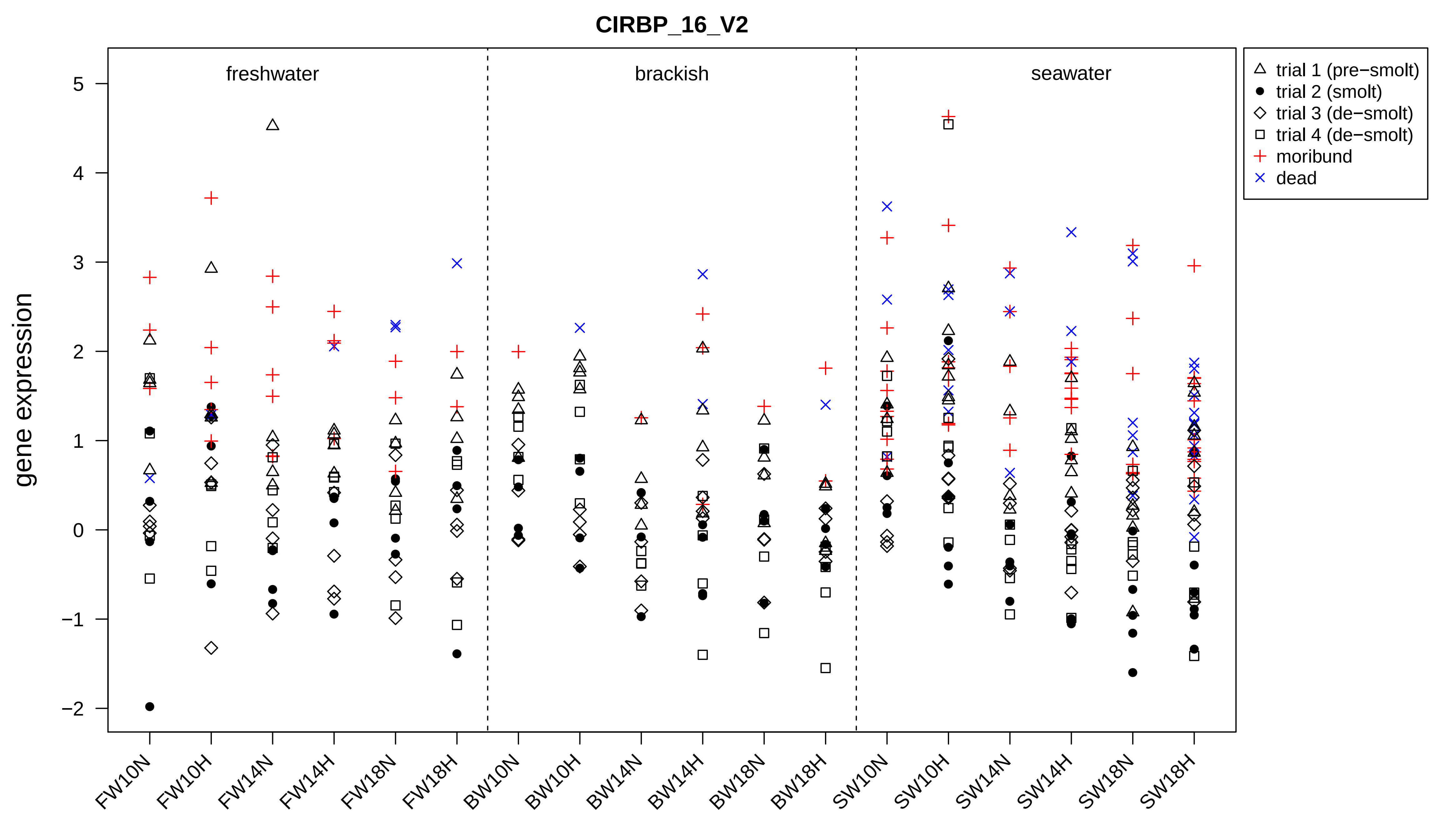

CIRBP_16_V2
freshwater
brackish
seawater
trial 1 (pre−smolt)
trial 2 (smolt)
trial 3 (de−smolt)
trial 4 (de−smolt)
moribund
dead
gene expression
5
4
3
2
1
0
−1
−2
FW10N
FW10H
FW14N
FW14H
FW18N
FW18H
BW10N
BW10H
BW14N
BW14H
BW18N
BW18H
SW10N
SW10H
SW14N
SW14H
SW18N
SW18H

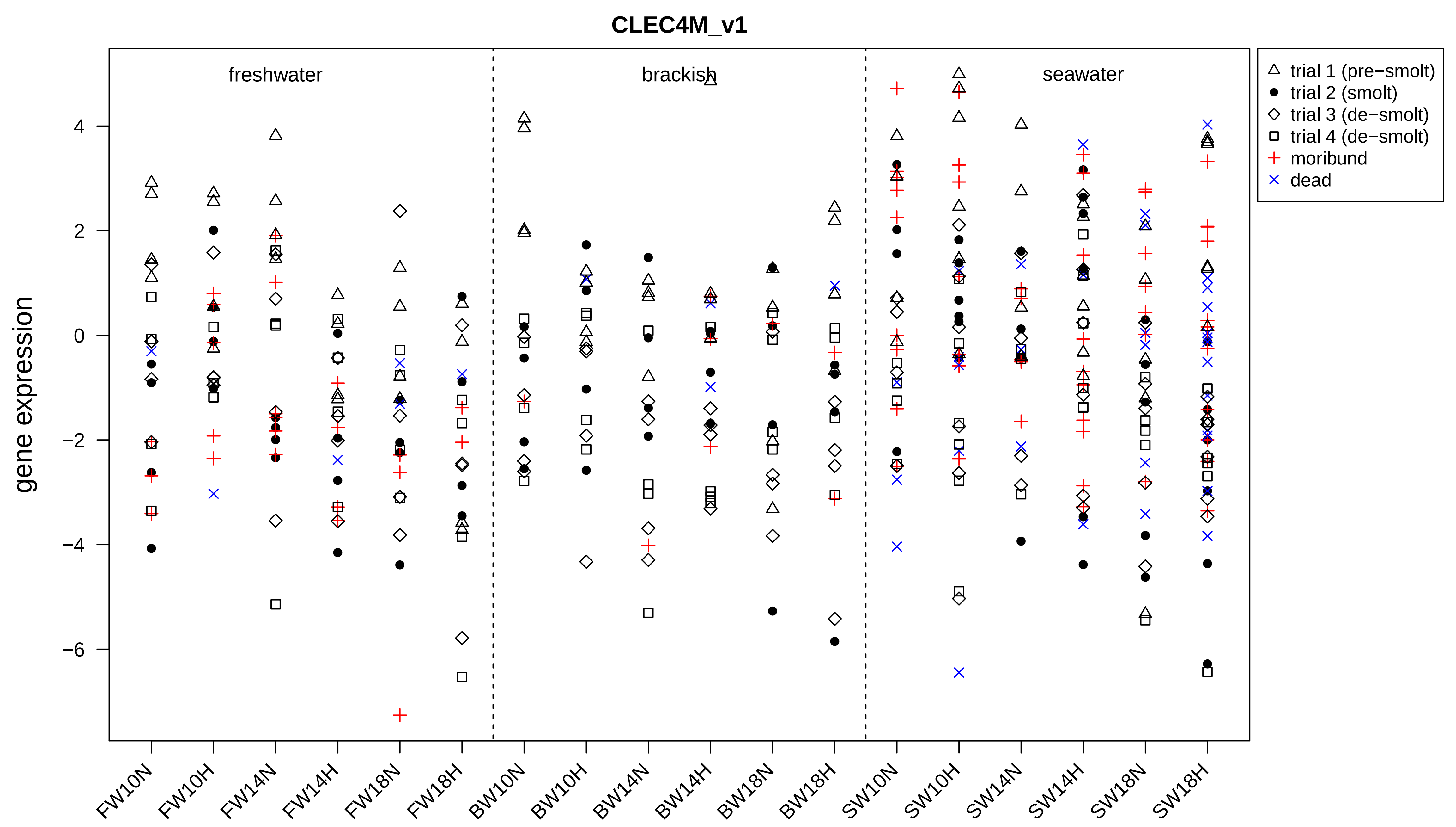
CLEC4M_v1
freshwater
brackish
seawater
gene expression
4
2
0
−2
−4
−6
FW10N
FW10H
FW14N
FW14H
FW18N
FW18H
BW10N
BW10H
BW14N
BW14H
BW18N
BW18H
SW10N
SW10H
SW14N
SW14H
SW18N
SW18H
trial 1 (pre−smolt)
trial 2 (smolt)
trial 3 (de−smolt)
trial 4 (de−smolt)
moribund
dead

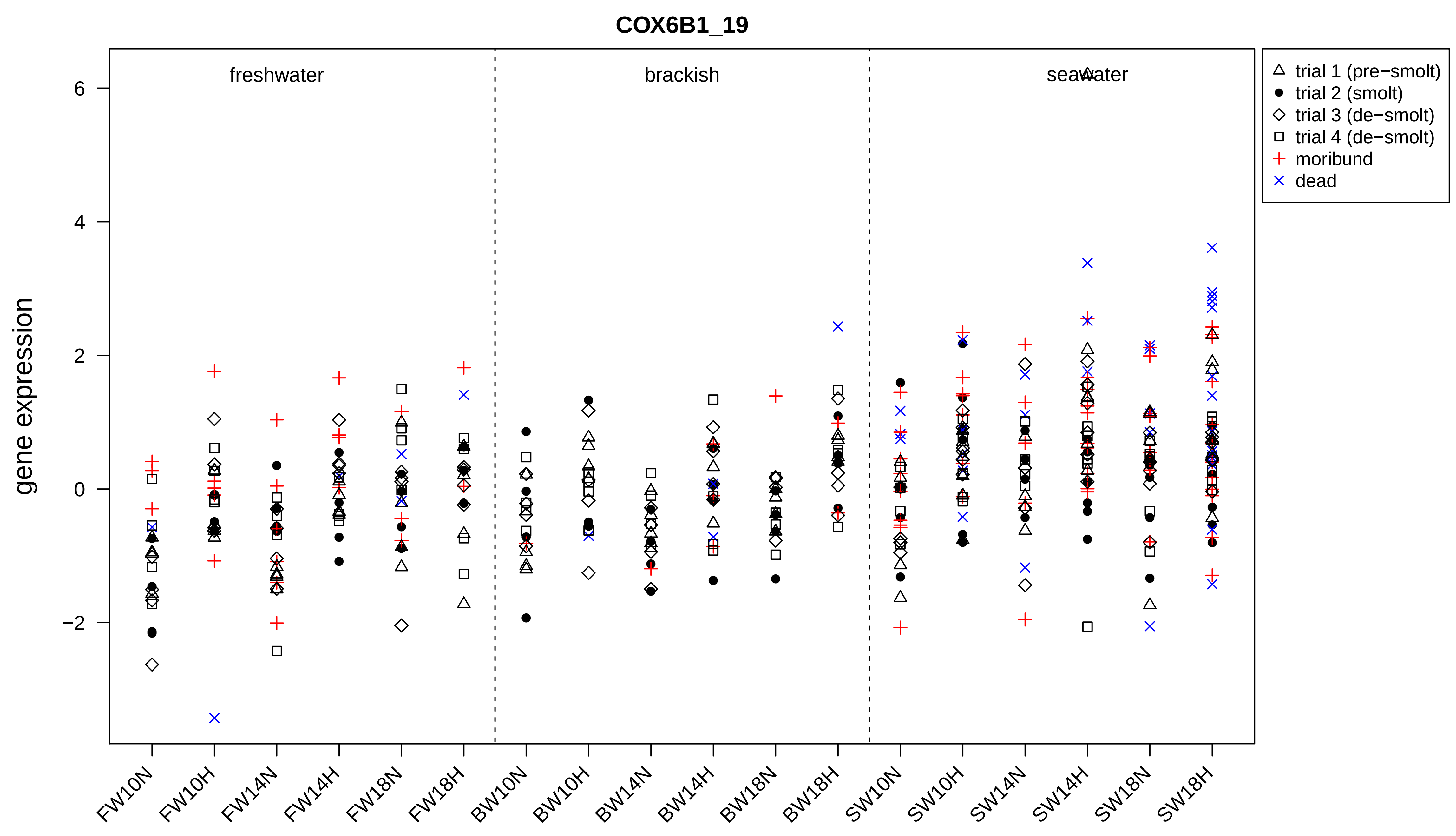
COX6B1_19
freshwater
brackish
seawater
trial 1 (pre−smolt)
trial 2 (smolt)
trial 3 (de−smolt)
trial 4 (de−smolt)
moribund
dead
gene expression
6
4
2
0
−2
FW10N
FW10H
FW14N
FW14H
FW18N
FW18H
BW10N
BW10H
BW14N
BW14H
BW18N
BW18H
SW10N
SW10H
SW14N
SW14H
SW18N
SW18H

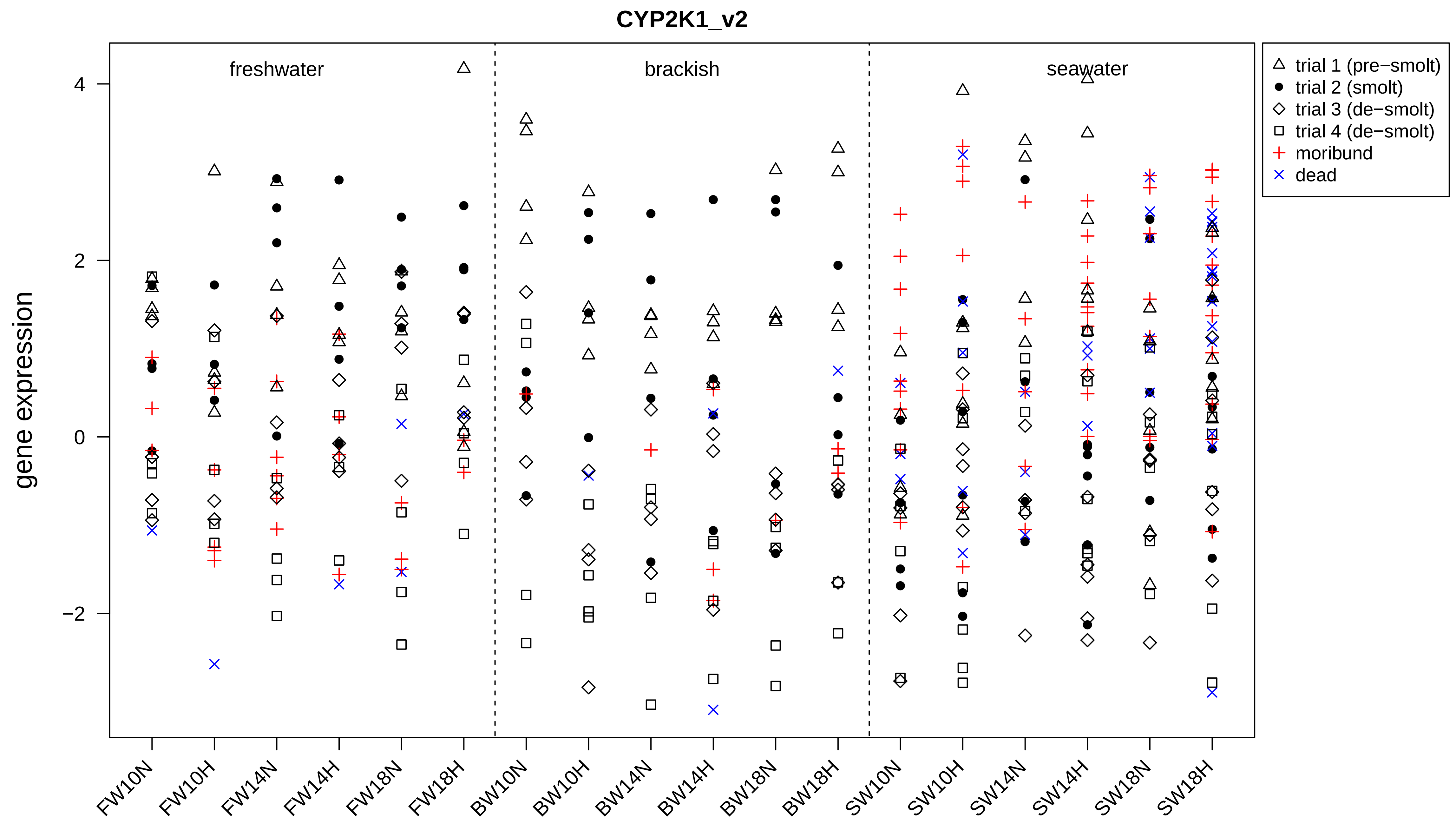
CYP2K1_v2
freshwater
brackish
seawater
gene expression
4
2
0
−2
trial 1 (pre−smolt)
trial 2 (smolt)
trial 3 (de−smolt)
trial 4 (de−smolt)
moribund
dead
FW10N
FW10H
FW14N
FW14H
FW18N
FW18H
BW10N
BW10H
BW14N
BW14H
BW18N
BW18H
SW10N
SW10H
SW14N
SW14H
SW18N
SW18H

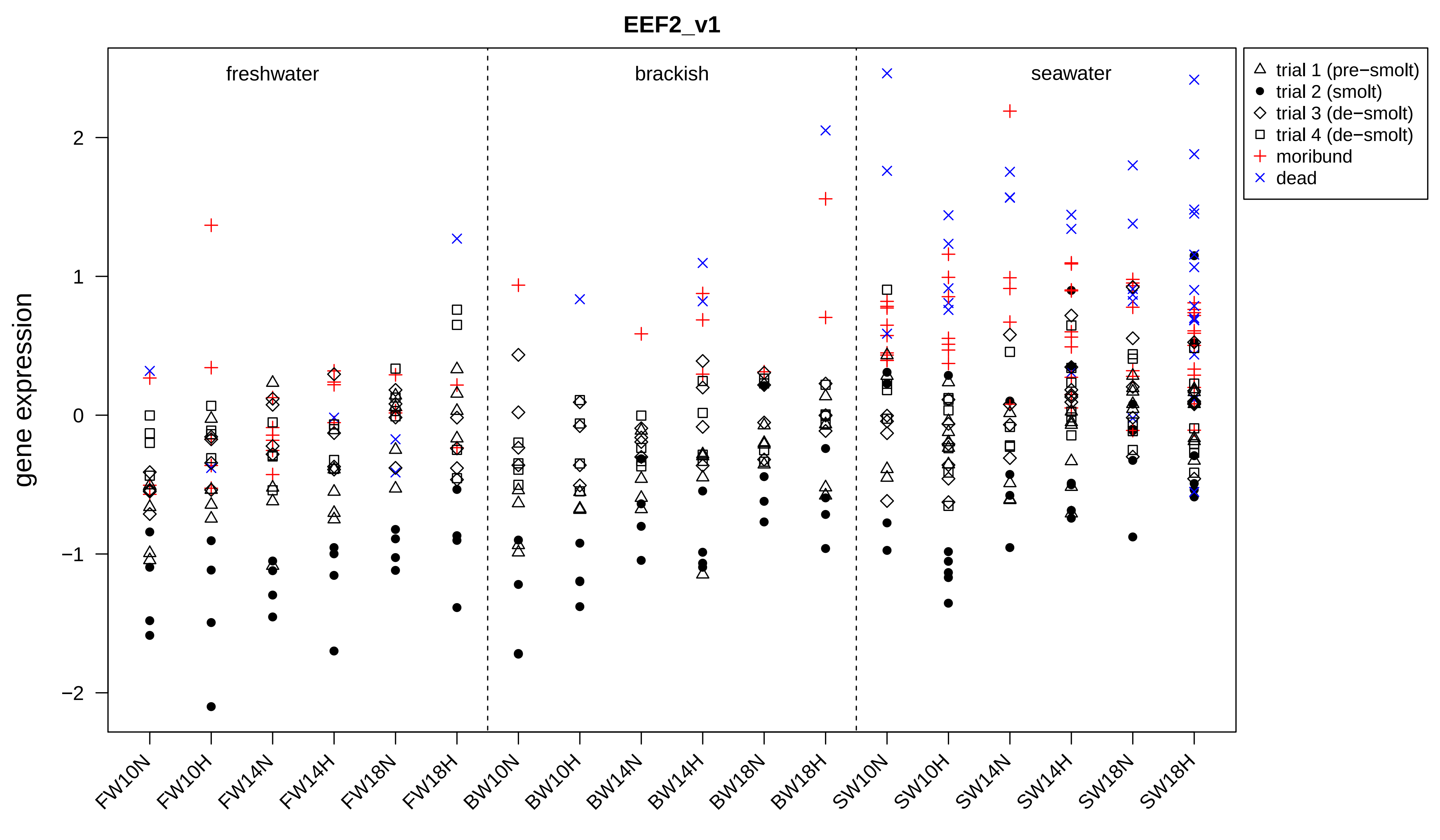
EEF2_v1
freshwater
brackish
seawater
trial 1 (pre−smolt)
trial 2 (smolt)
trial 3 (de−smolt)
trial 4 (de−smolt)
moribund
dead
gene expression
2
1
0
−1
−2
FW10N
FW10H
FW14N
FW14H
FW18N
FW18H
BW10N
BW10H
BW14N
BW14H
BW18N
BW18H
SW10N
SW10H
SW14N
SW14H
SW18N
SW18H

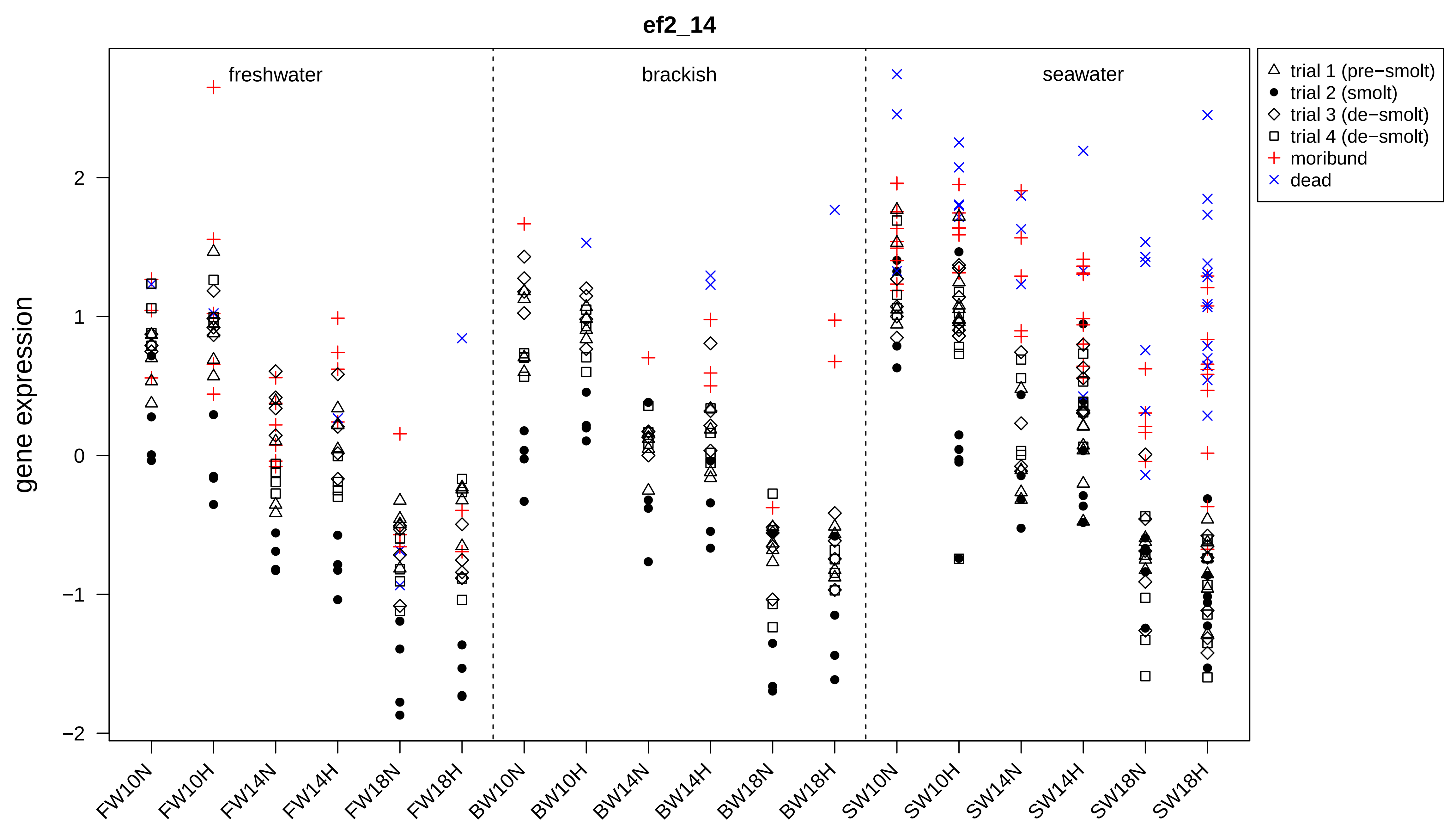
ef2_14
freshwater
brackish
seawater
trial 1 (pre−smolt)
trial 2 (smolt)
trial 3 (de−smolt)
trial 4 (de−smolt)
moribund
dead
gene expression
2
1
0
−1
−2
FW10N
FW10H
FW14N
FW14H
FW18N
FW18H
BW10N
BW10H
BW14N
BW14H
BW18N
BW18H
SW10N
SW10H
SW14N
SW14H
SW18N
SW18H

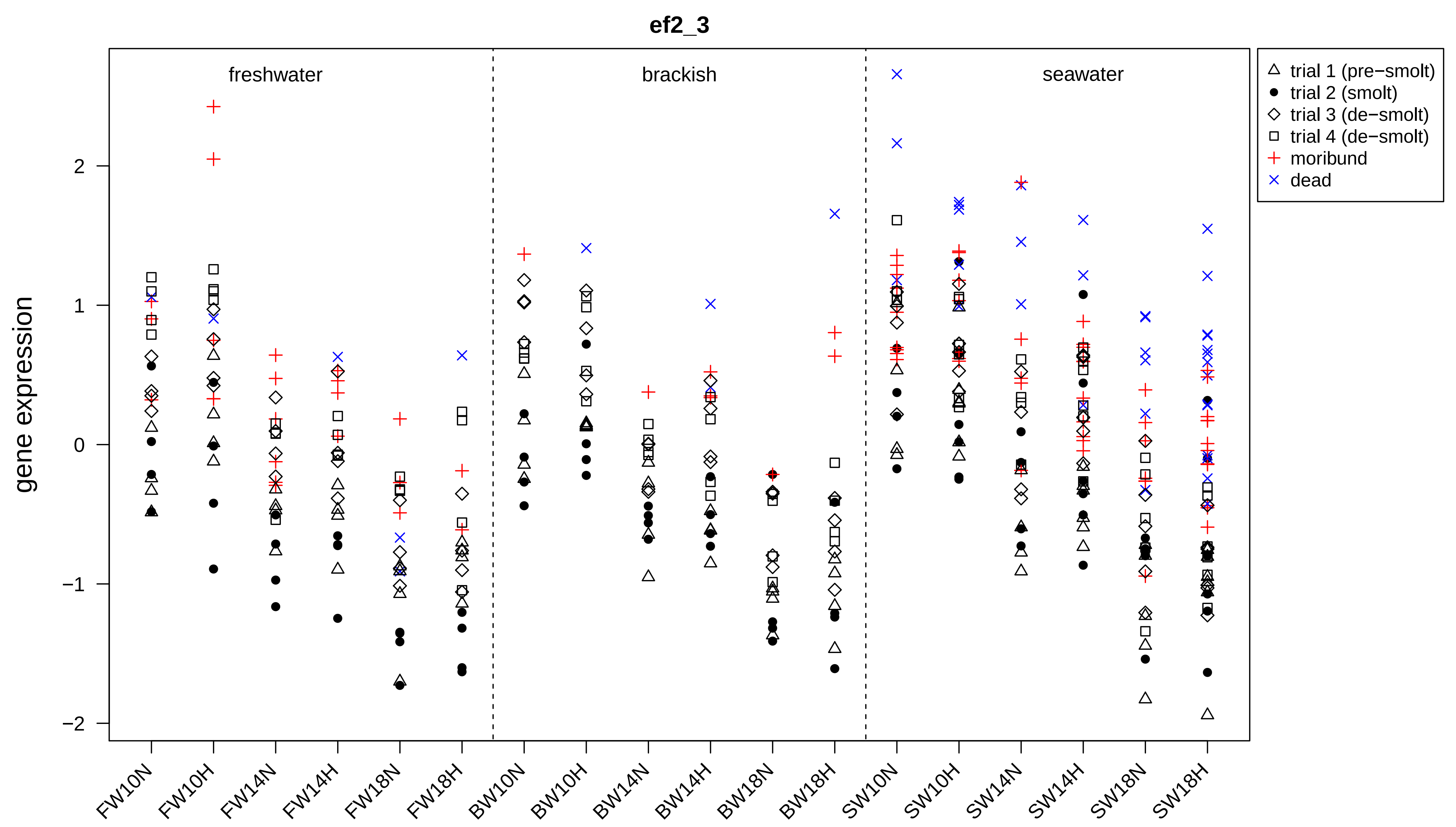
ef2_3
freshwater
brackish
seawater
trial 1 (pre−smolt)
trial 2 (smolt)
trial 3 (de−smolt)
trial 4 (de−smolt)
moribund
dead
gene expression
2
1
0
−1
−2
FW10N
FW10H
FW14N
FW14H
FW18N
FW18H
BW10N
BW10H
BW14N
BW14H
BW18N
BW18H
SW10N
SW10H
SW14N
SW14H
SW18N
SW18H

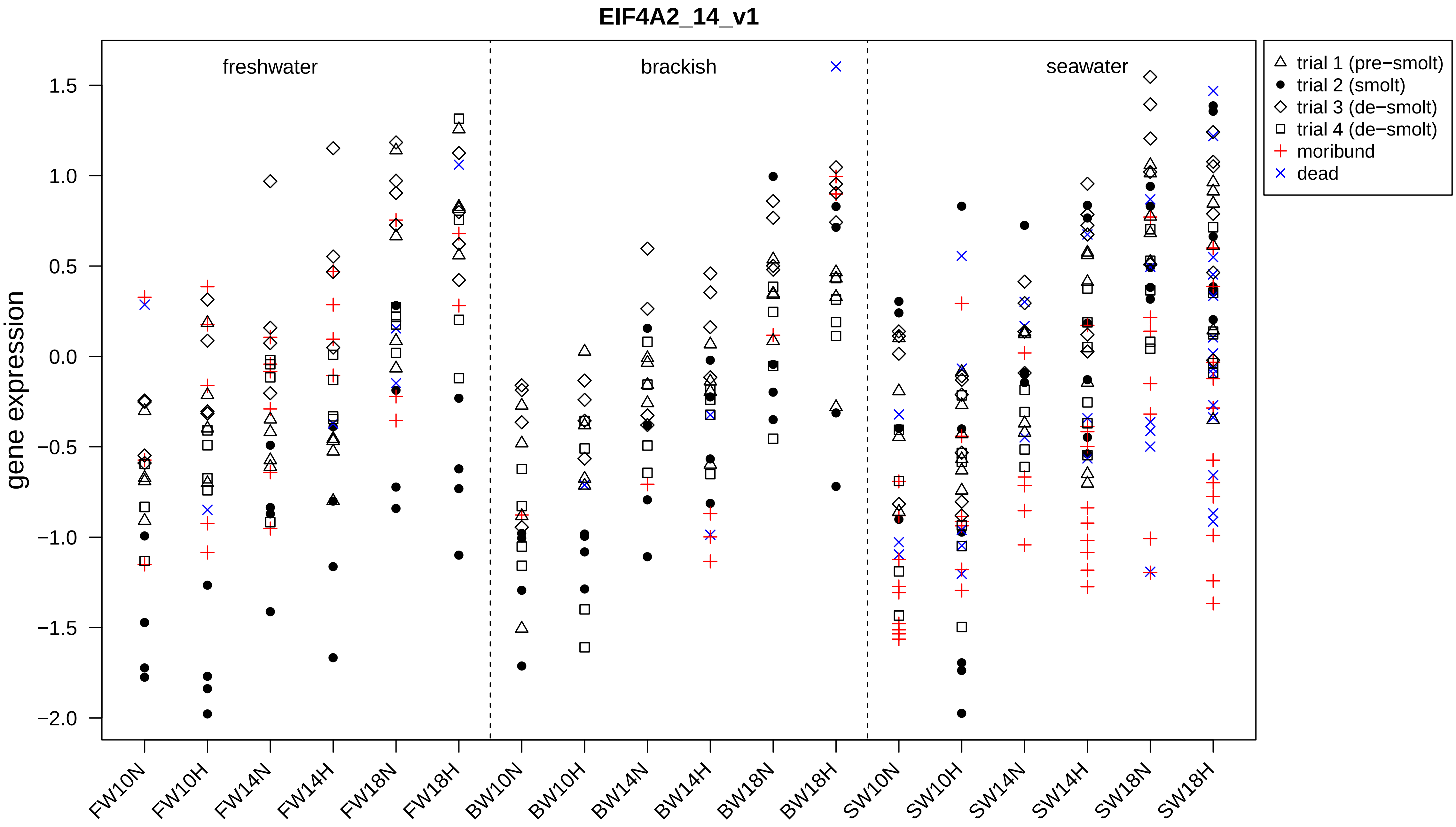

EIF4A2_14_v1
freshwater
brackish
seawater
gene expression
1.5
1.0
0.5
0.0
−0.5
−1.0
−1.5
−2.0
FW10N
FW10H
FW14N
FW14H
FW18N
FW18H
BW10N
BW10H
BW14N
BW14H
BW18N
BW18H
SW10N
SW10H
SW14N
SW14H
SW18N
SW18H
trial 1 (pre−smolt)
trial 2 (smolt)
trial 3 (de−smolt)
trial 4 (de−smolt)
moribund
dead

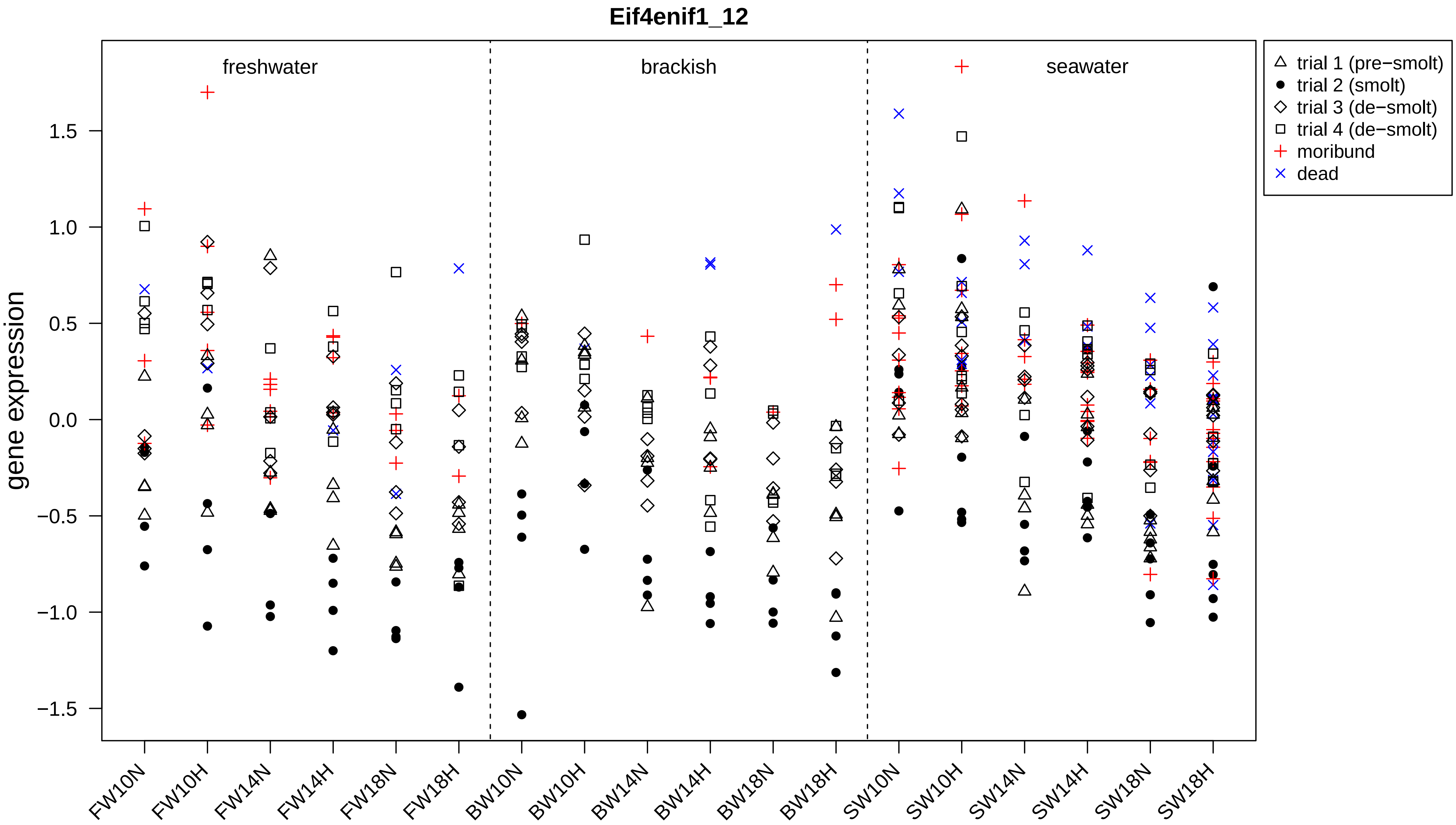

Eif4enif1_12
freshwater
brackish
seawater
trial 1 (pre−smolt)
trial 2 (smolt)
trial 3 (de−smolt)
trial 4 (de−smolt)
moribund
dead
gene expression
1.5
1.0
0.5
0.0
−0.5
−1.0
−1.5
FW10N
FW10H
FW14N
FW14H
FW18N
FW18H
BW10N
BW10H
BW14N
BW14H
BW18N
BW18H
SW10N
SW10H
SW14N
SW14H
SW18N
SW18H

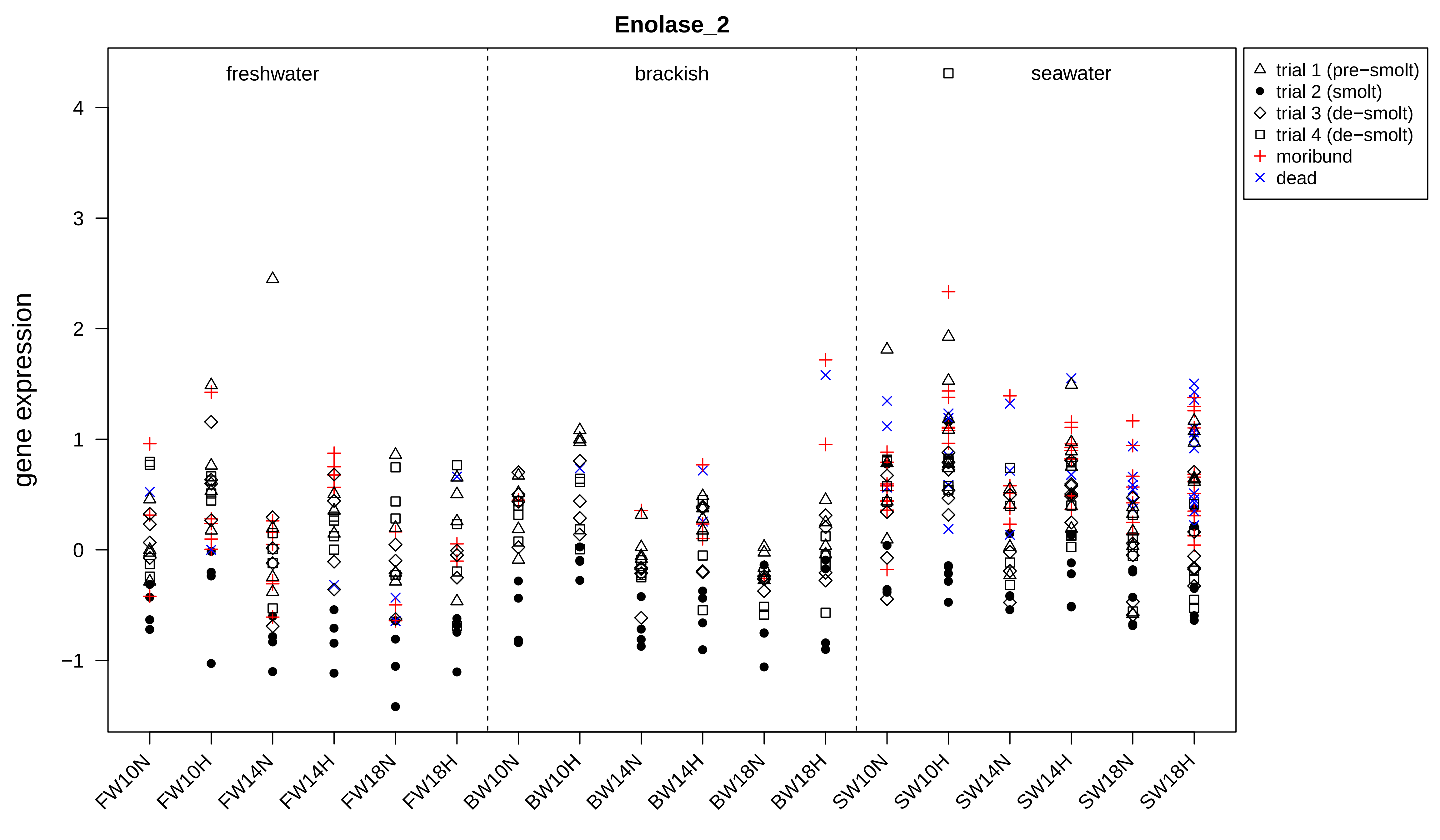
Enolase_2
freshwater
brackish
seawater
trial 1 (pre−smolt)
trial 2 (smolt)
trial 3 (de−smolt)
trial 4 (de−smolt)
moribund
dead
gene expression
4
3
2
1
0
−1
FW10N
FW10H
FW14N
FW14H
FW18N
FW18H
BW10N
BW10H
BW14N
BW14H
BW18N
BW18H
SW10N
SW10H
SW14N
SW14H
SW18N
SW18H

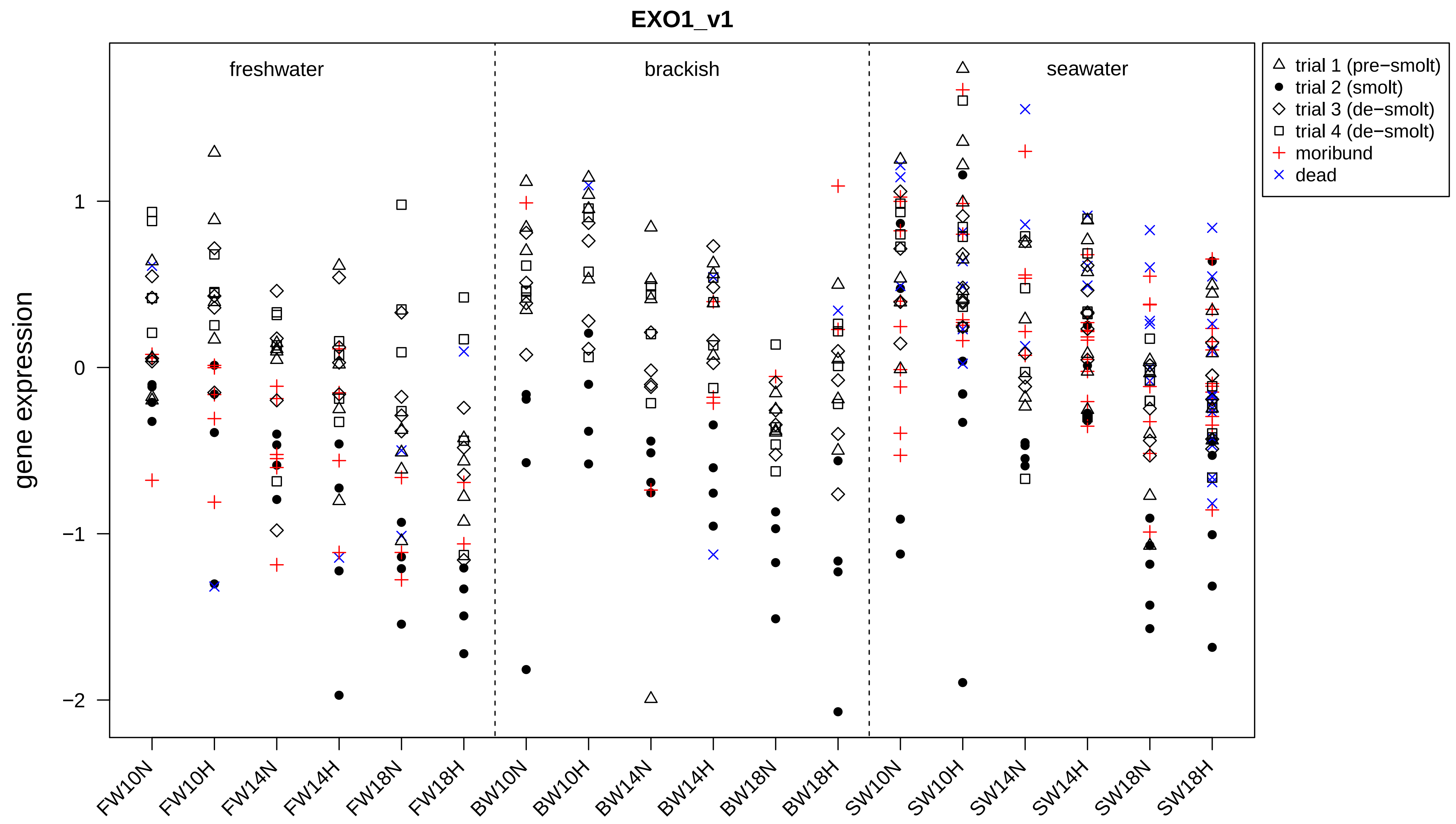
EXO1_v1
freshwater
brackish
seawater
gene expression
1
0
−1
−2
FW10N
FW10H
FW14N
FW14H
FW18N
FW18H
BW10N
BW10H
BW14N
BW14H
BW18N
BW18H
SW10N
SW10H
SW14N
SW14H
SW18N
SW18H
trial 1 (pre−smolt)
trial 2 (smolt)
trial 3 (de−smolt)
trial 4 (de−smolt)
moribund
dead

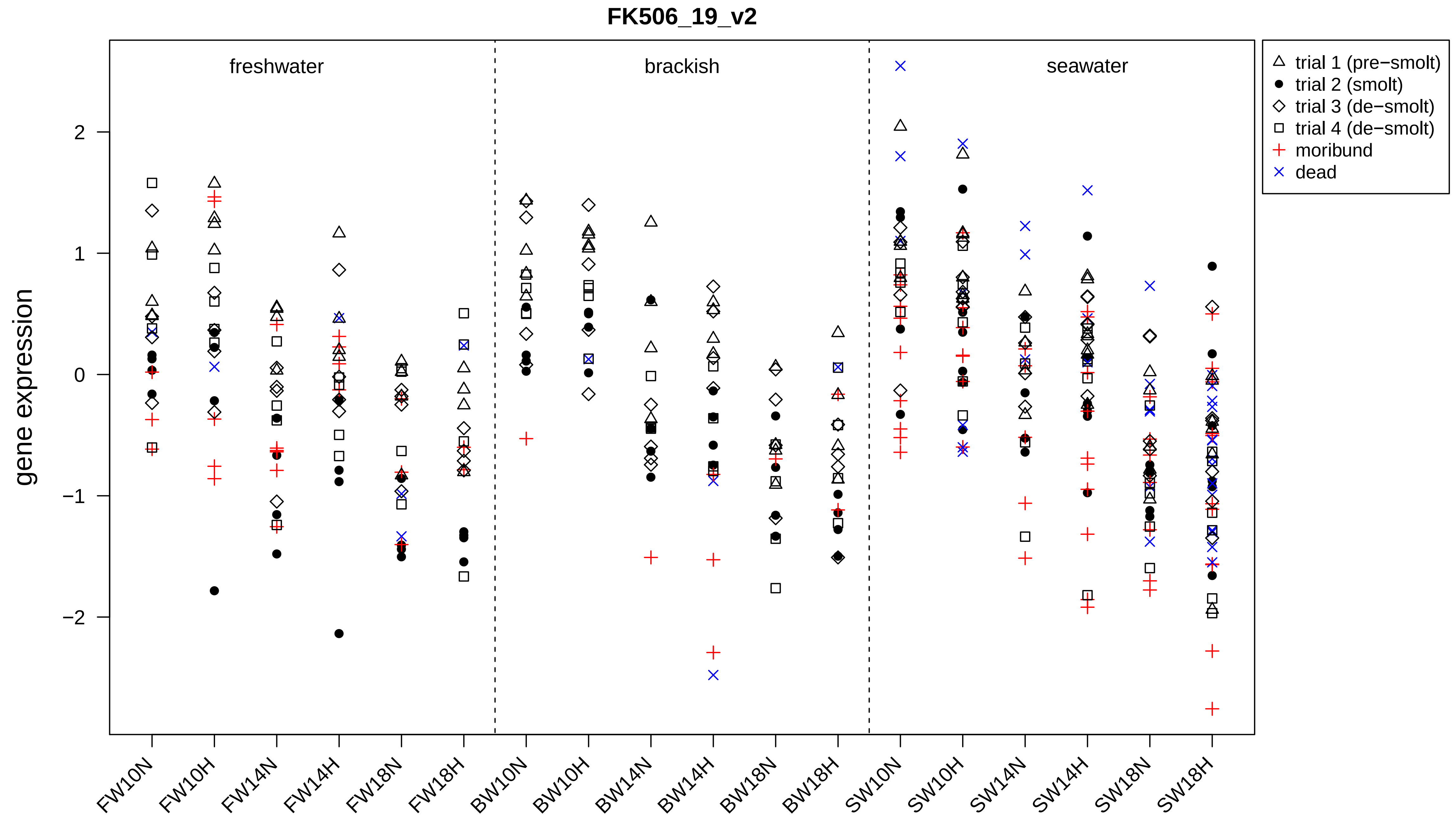

FK506_19_v2
freshwater
brackish
seawater
gene expression
2
1
0
−1
−2
trial 1 (pre−smolt)
trial 2 (smolt)
trial 3 (de−smolt)
trial 4 (de−smolt)
moribund
dead
FW10N
FW10H
FW14N
FW14H
FW18N
FW18H
BW10N
BW10H
BW14N
BW14H
BW18N
BW18H
SW10N
SW10H
SW14N
SW14H
SW18N
SW18H

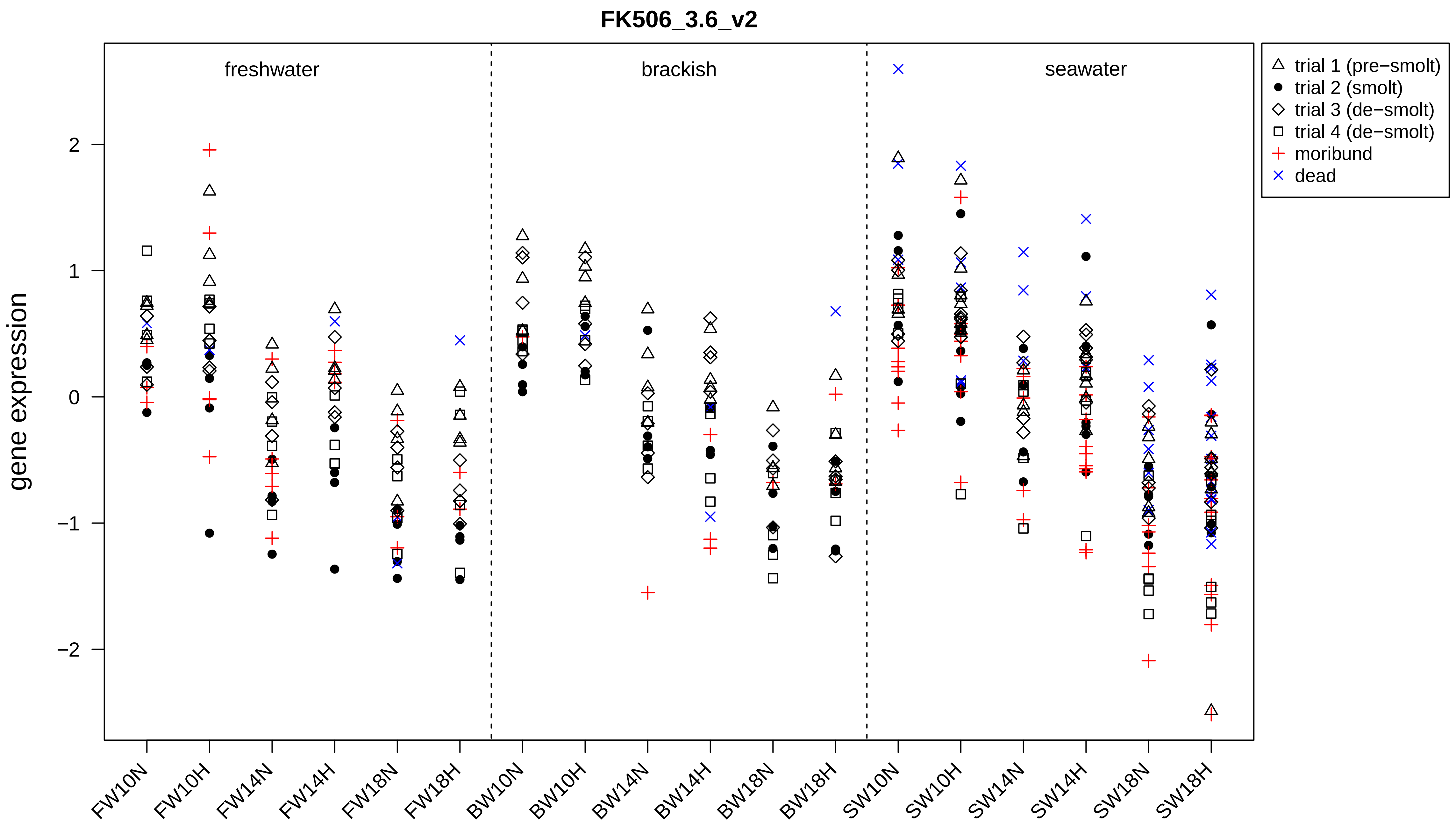

FK506_3.6_v2
freshwater
brackish
seawater
gene expression
2
1
0
−1
−2
trial 1 (pre−smolt)
trial 2 (smolt)
trial 3 (de−smolt)
trial 4 (de−smolt)
moribund
dead
FW10N
FW10H
FW14N
FW14H
FW18N
FW18H
BW10N
BW10H
BW14N
BW14H
BW18N
BW18H
SW10N
SW10H
SW14N
SW14H
SW18N
SW18H

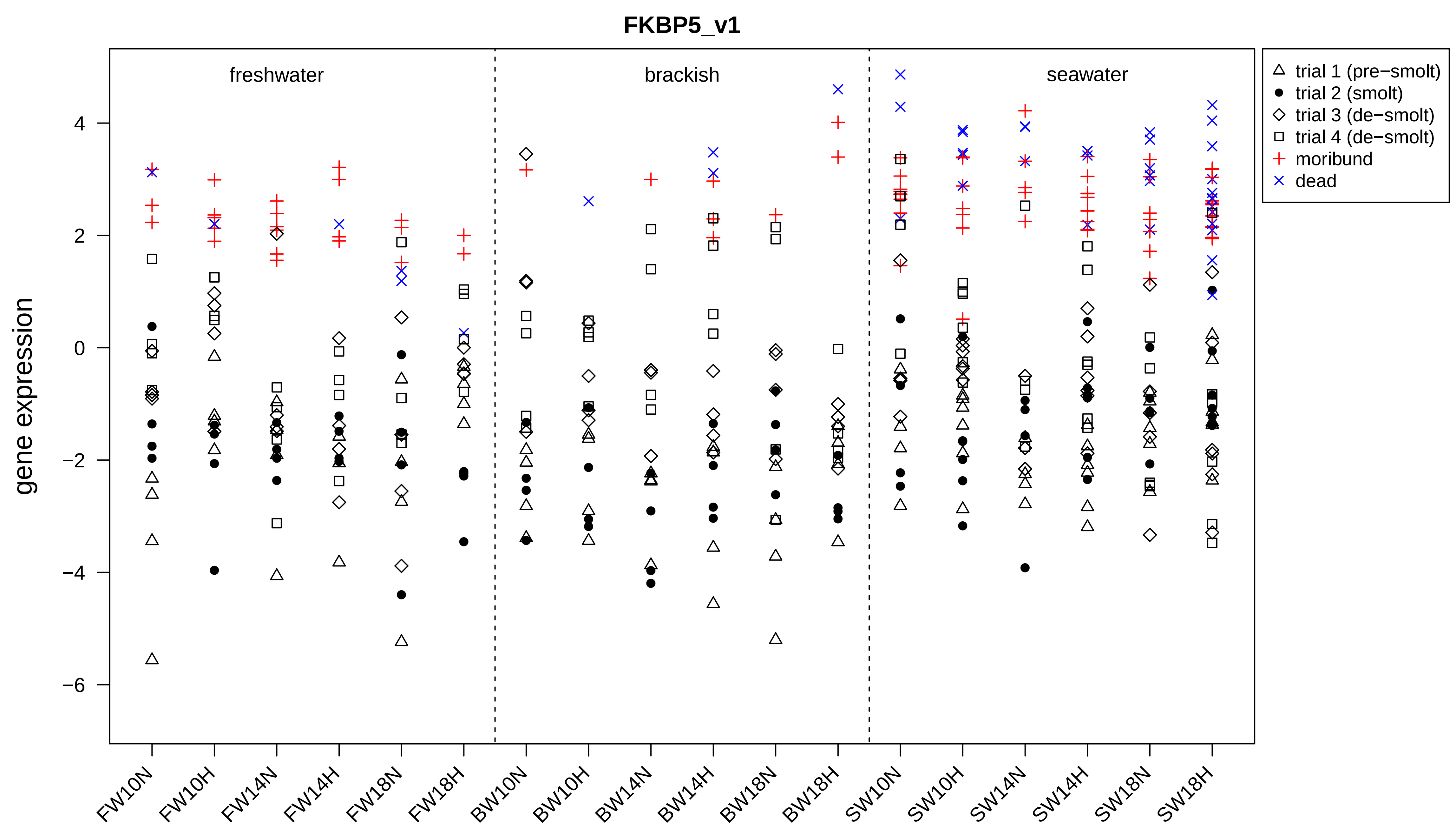
FKBP5_v1
freshwater
brackish
seawater
gene expression
4
2
0
−2
−4
−6
FW10N
FW10H
FW14N
FW14H
FW18N
FW18H
BW10N
BW10H
BW14N
BW14H
BW18N
BW18H
SW10N
SW10H
SW14N
SW14H
SW18N
SW18H
trial 1 (pre−smolt)
trial 2 (smolt)
trial 3 (de−smolt)
trial 4 (de−smolt)
moribund
dead

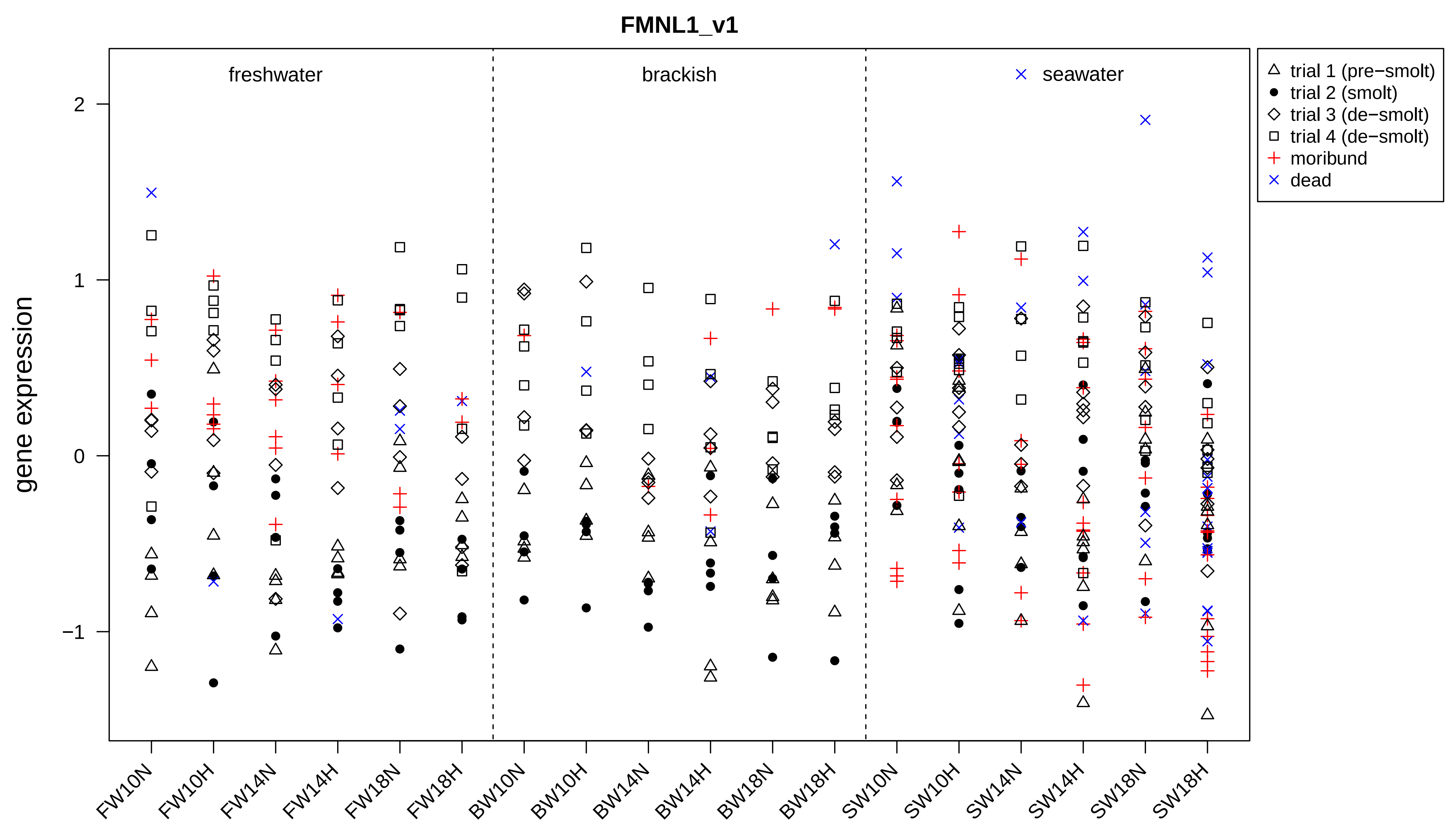

FMNL1_v1
freshwater
brackish
seawater
gene expression
2
1
0
−1
trial 1 (pre−smolt)
trial 2 (smolt)
trial 3 (de−smolt)
trial 4 (de−smolt)
moribund
dead
FW10N
FW10H
FW14N
FW14H
FW18N
FW18H
BW10N
BW10H
BW14N
BW14H
BW18N
BW18H
SW10N
SW10H
SW14N
SW14H
SW18N
SW18H

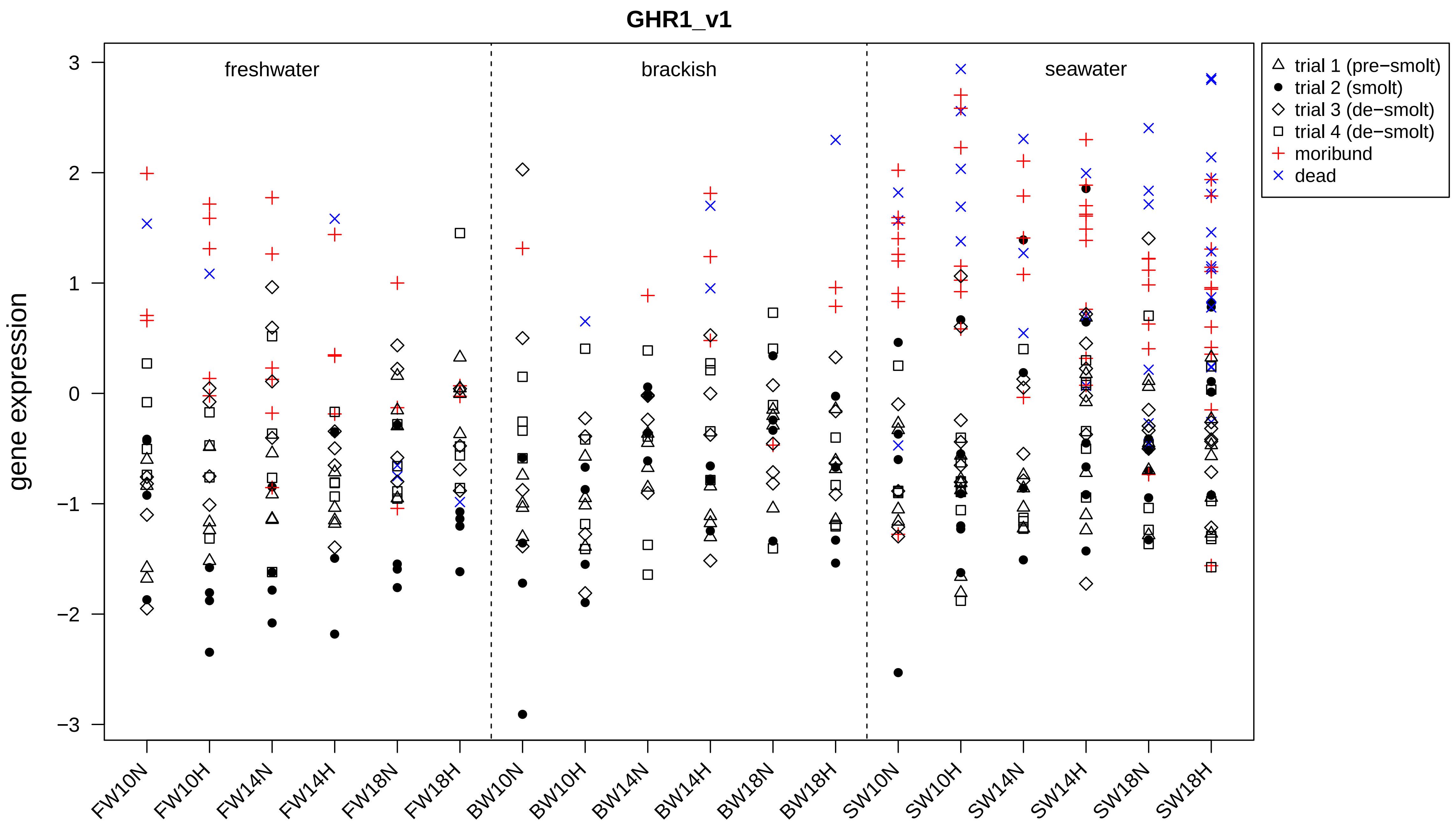
GHR1_v1
gene expression
3
2
1
0
−1
−2
−3
freshwater
brackish
seawater
FW10N
FW10H
FW14N
FW14H
FW18N
FW18H
BW10N
BW10H
BW14N
BW14H
BW18N
BW18H
SW10N
SW10H
SW14N
SW14H
SW18N
SW18H
trial 1 (pre−smolt)
trial 2 (smolt)
trial 3 (de−smolt)
trial 4 (de−smolt)
moribund
dead

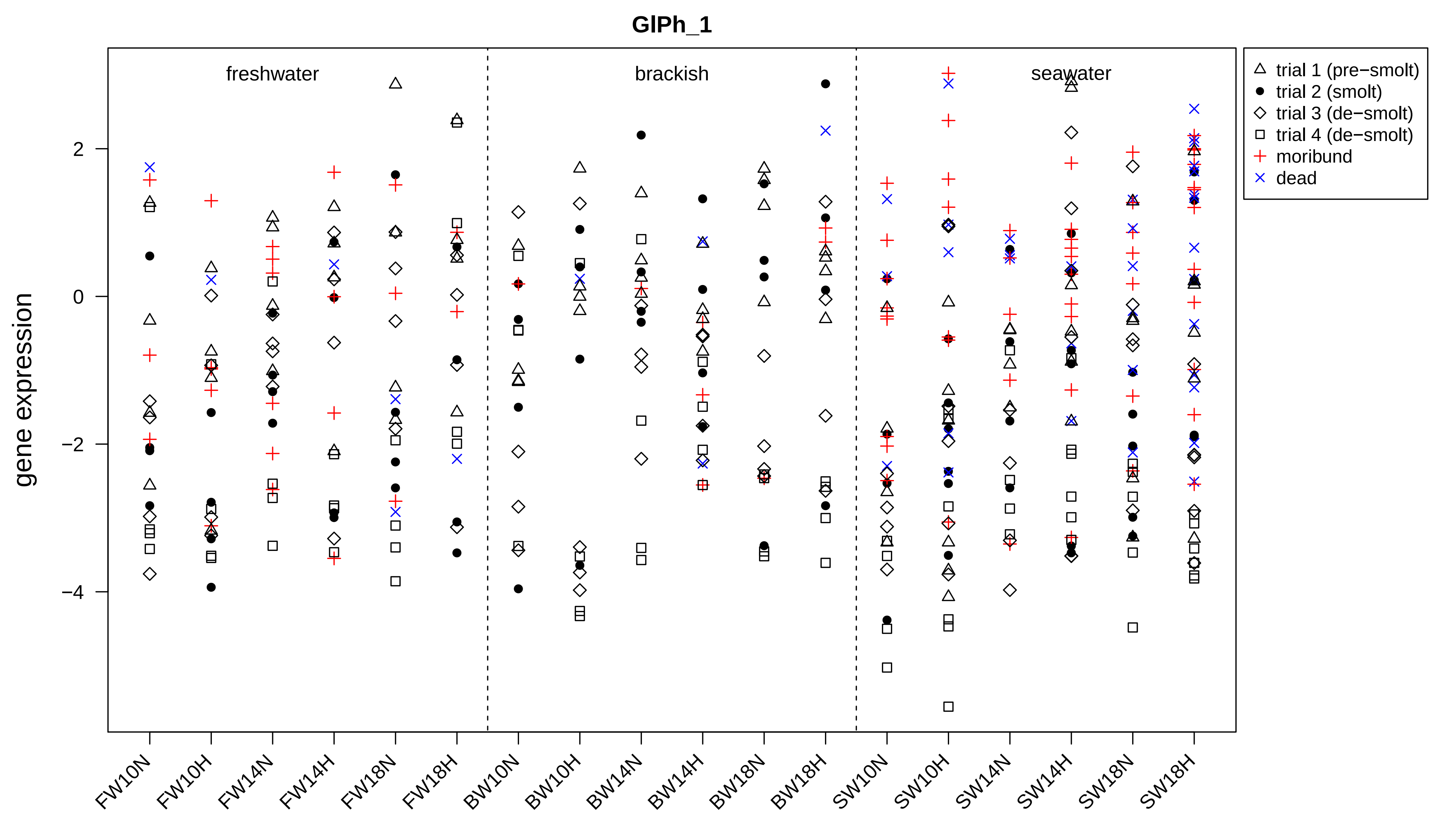

GlPh_1
freshwater
brackish
seawater
gene expression
2
0
−2
−4
trial 1 (pre−smolt)
trial 2 (smolt)
trial 3 (de−smolt)
trial 4 (de−smolt)
moribund
dead
FW10N
FW10H
FW14N
FW14H
FW18N
FW18H
BW10N
BW10H
BW14N
BW14H
BW18N
BW18H
SW10N
SW10H
SW14N
SW14H
SW18N
SW18H

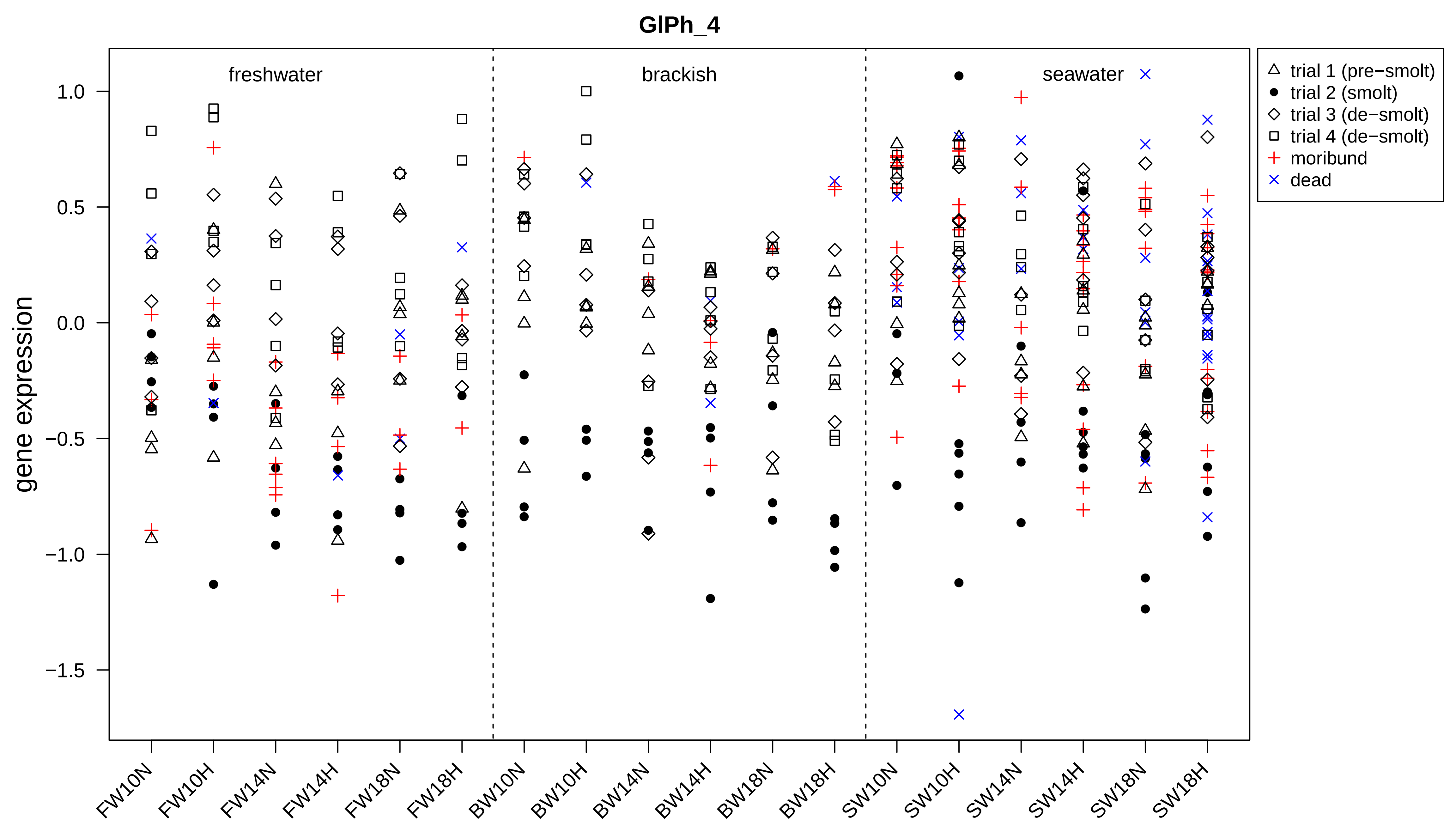

GlPh_4
freshwater
brackish
seawater
gene expression
1.0
0.5
0.0
−0.5
−1.0
−1.5
FW10N
FW10H
FW14N
FW14H
FW18N
FW18H
BW10N
BW10H
BW14N
BW14H
BW18N
BW18H
SW10N
SW10H
SW14N
SW14H
SW18N
SW18H
trial 1 (pre−smolt)
trial 2 (smolt)
trial 3 (de−smolt)
trial 4 (de−smolt)
moribund
dead

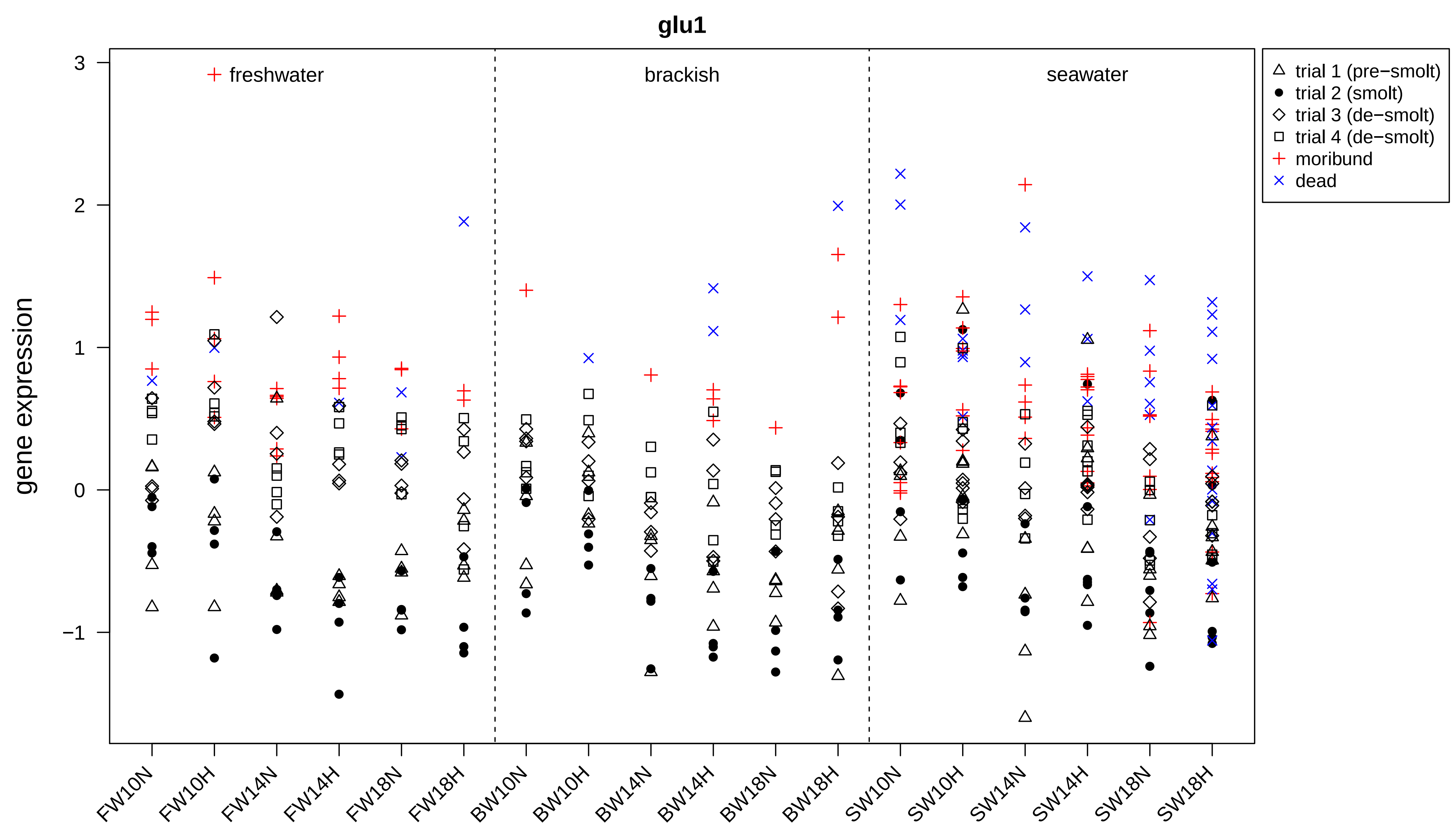

glu1
gene expression
3
2
1
0
−1
freshwater
brackish
seawater
trial 1 (pre−smolt)
trial 2 (smolt)
trial 3 (de−smolt)
trial 4 (de−smolt)
moribund
dead
FW10N
FW10H
FW14N
FW14H
FW18N
FW18H
BW10N
BW10H
BW14N
BW14H
BW18N
BW18H
SW10N
SW10H
SW14N
SW14H
SW18N
SW18H

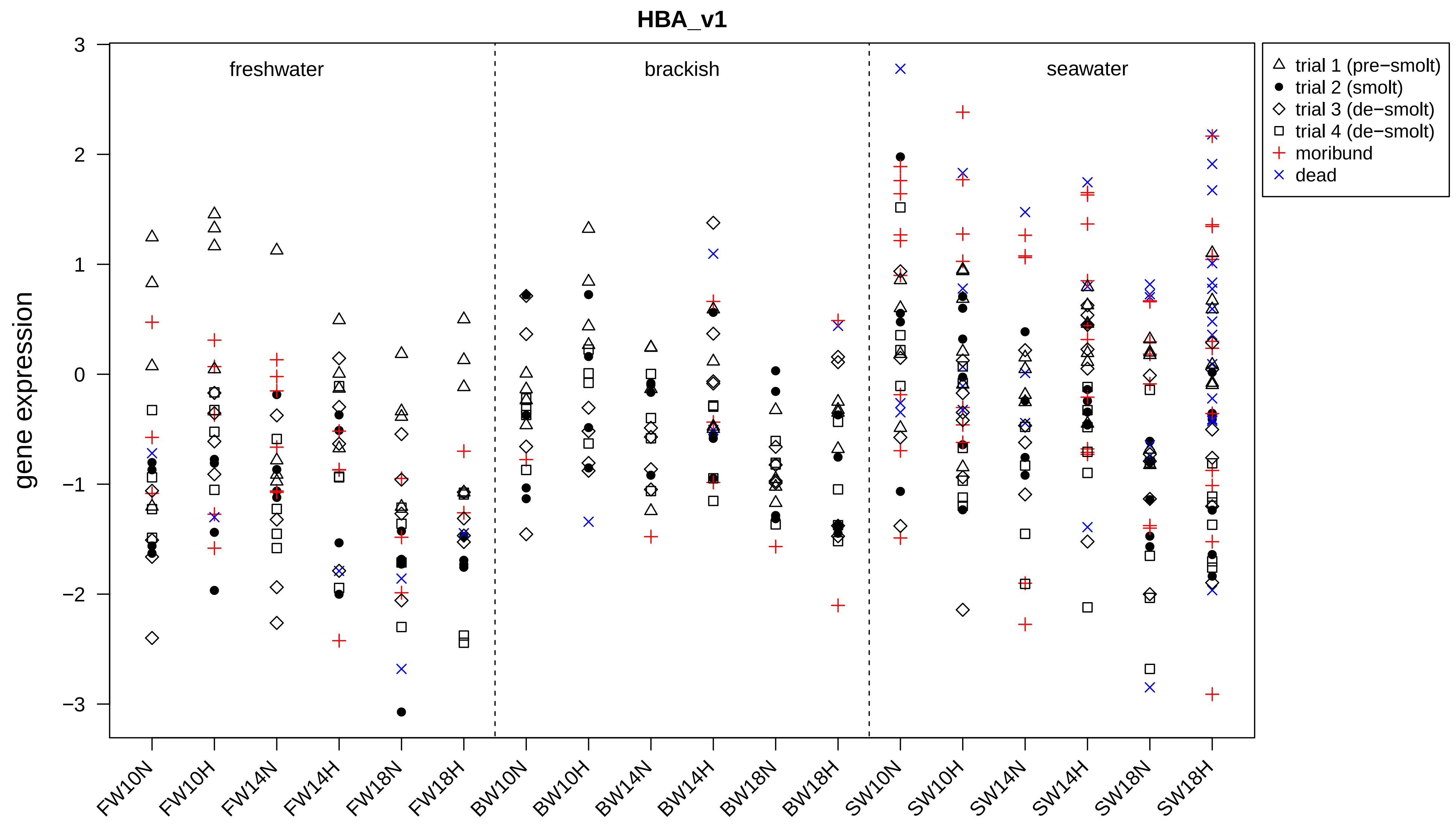
HBA_v1
gene expression
freshwater
brackish
seawater
trial 1 (pre−smolt)
trial 2 (smolt)
trial 3 (de−smolt)
trial 4 (de−smolt)
moribund
dead
3
2
1
0
−1
−2
−3
FW10N
FW10H
FW14N
FW14H
FW18N
FW18H
BW10N
BW10H
BW14N
BW14H
BW18N
BW18H
SW10N
SW10H
SW14N
SW14H
SW18N
SW18H

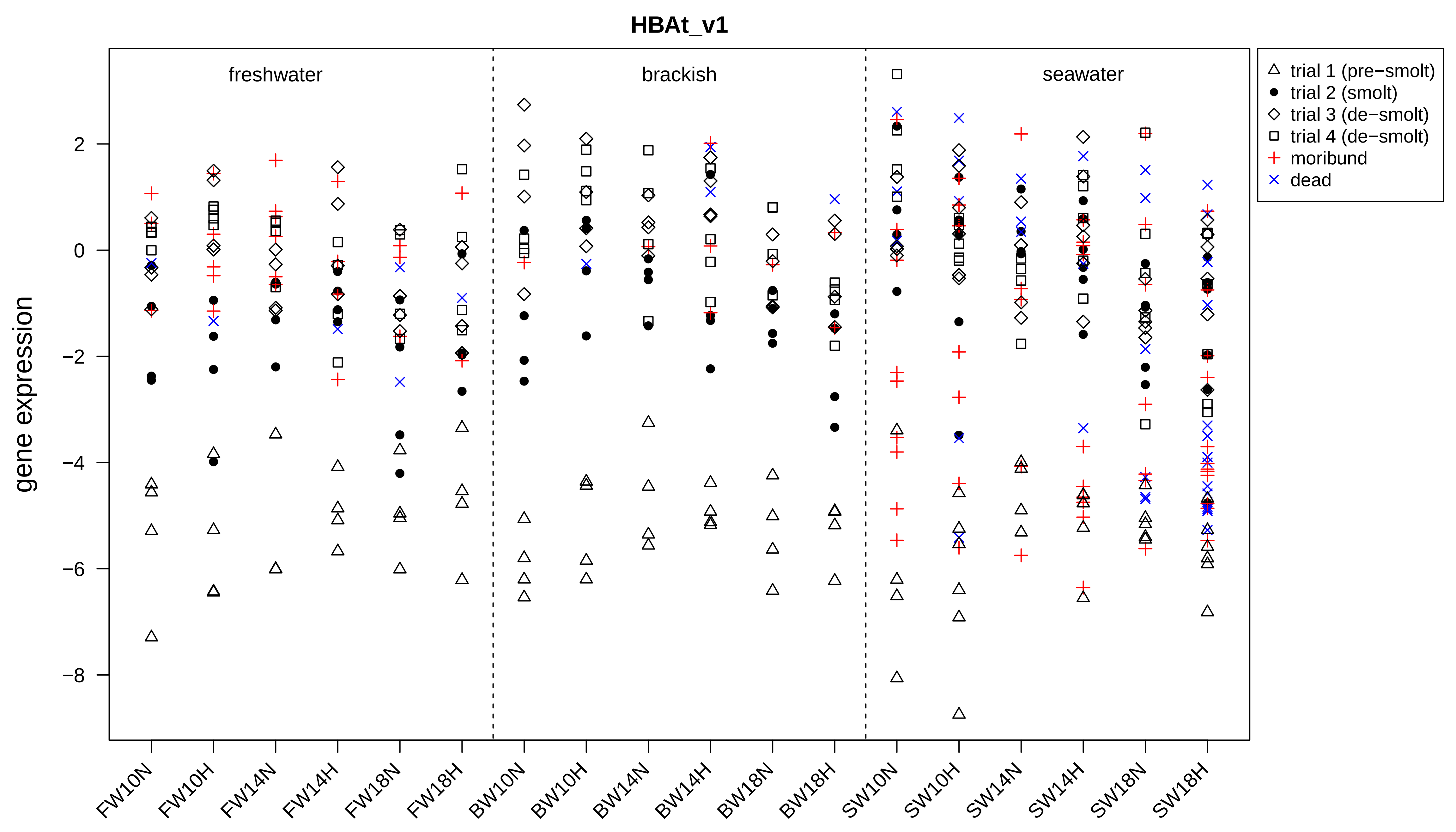
HBAt_v1
freshwater
brackish
seawater
trial 1 (pre−smolt)
trial 2 (smolt)
trial 3 (de−smolt)
trial 4 (de−smolt)
moribund
dead
gene expression
2
0
−2
−4
−6
−8
FW10N
FW10H
FW14N
FW14H
FW18N
FW18H
BW10N
BW10H
BW14N
BW14H
BW18N
BW18H
SW10N
SW10H
SW14N
SW14H
SW18N
SW18H

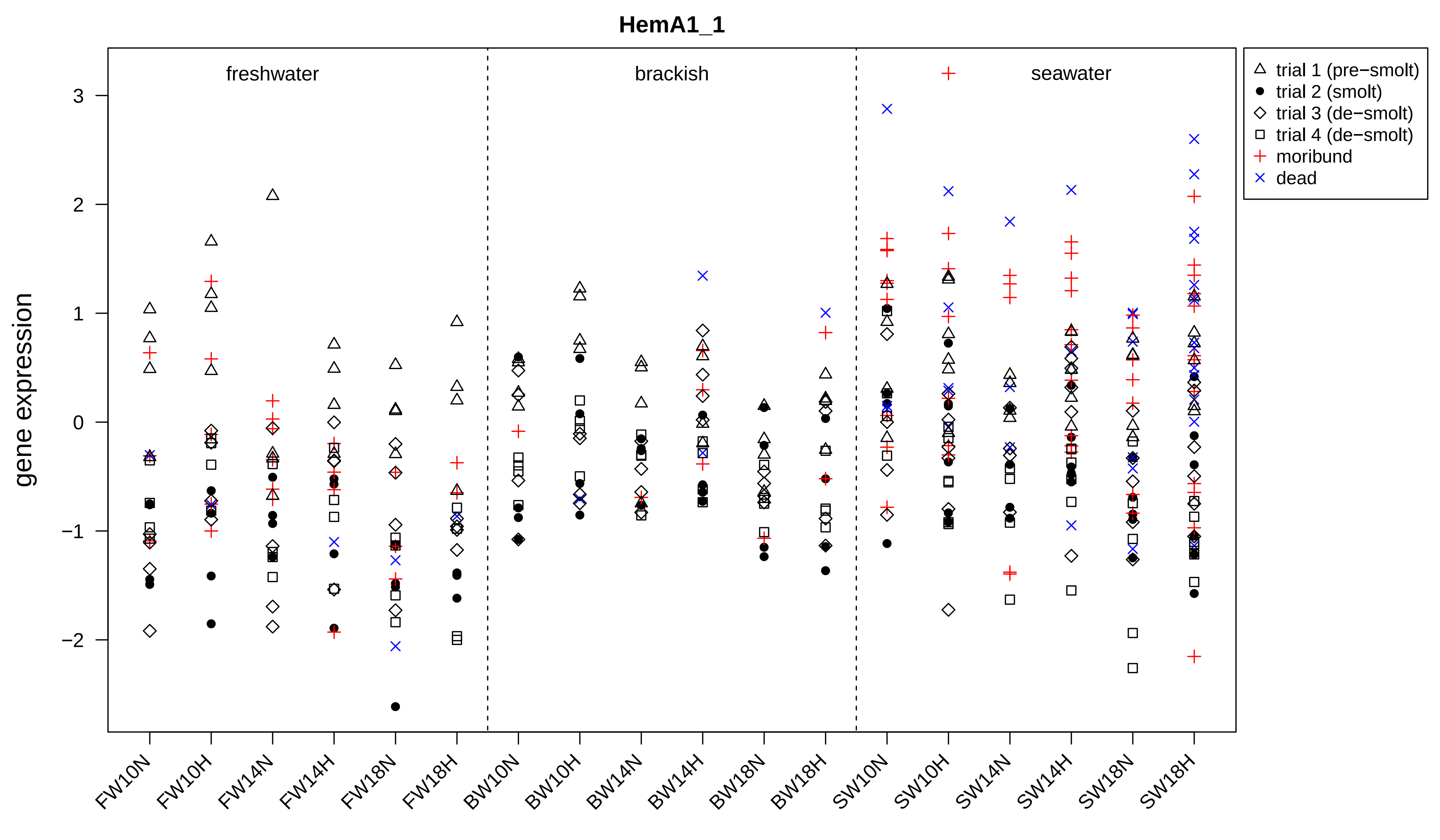

HemA1_1
freshwater
brackish
seawater
gene expression
3
2
1
0
−1
−2
trial 1 (pre−smolt)
trial 2 (smolt)
trial 3 (de−smolt)
trial 4 (de−smolt)
moribund
dead
FW10N
FW10H
FW14N
FW14H
FW18N
FW18H
BW10N
BW10H
BW14N
BW14H
BW18N
BW18H
SW10N
SW10H
SW14N
SW14H
SW18N
SW18H

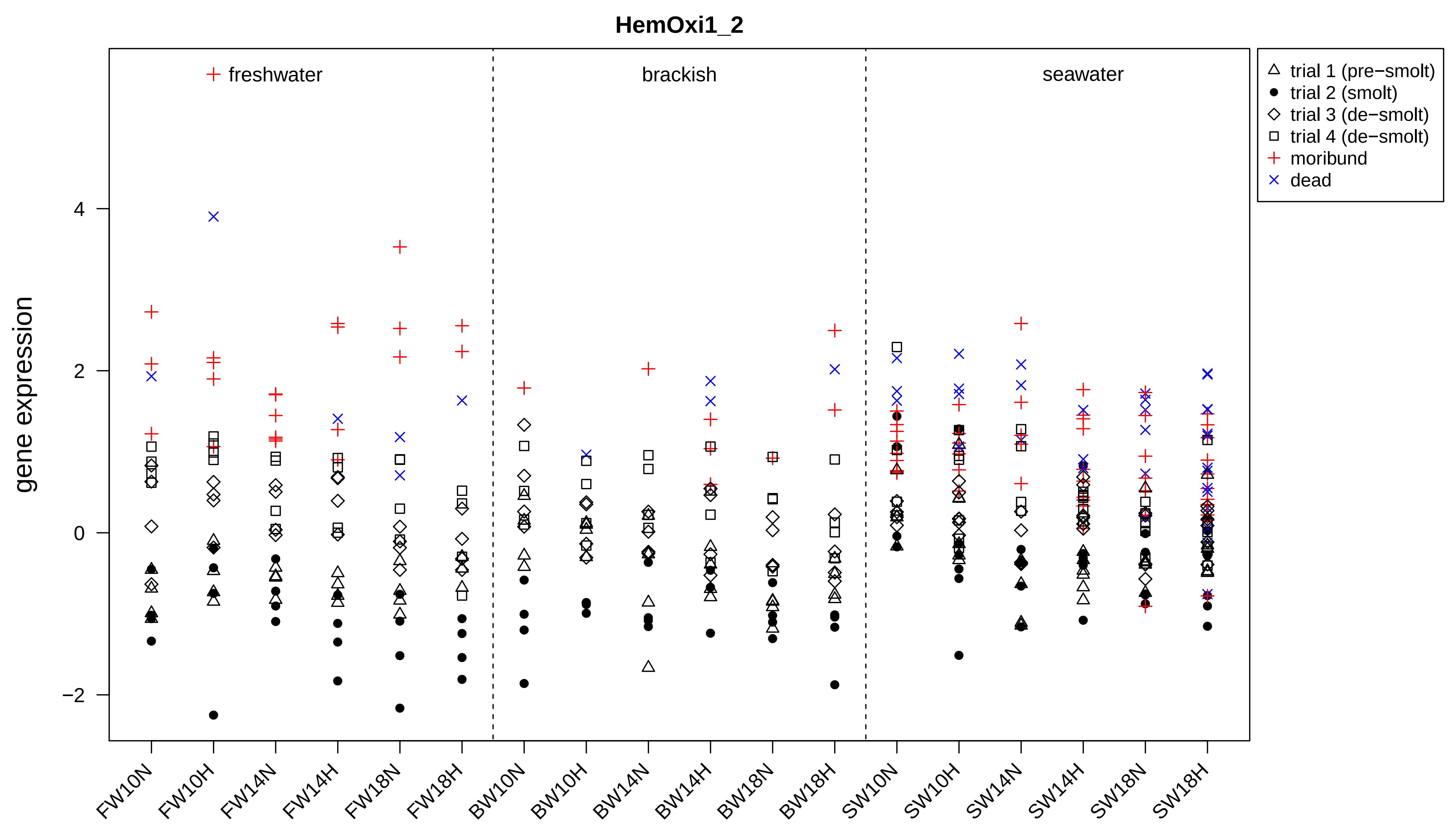

HemOxi1_2
gene expression
freshwater
brackish
seawater
trial 1 (pre−smolt)
trial 2 (smolt)
trial 3 (de−smolt)
trial 4 (de−smolt)
moribund
dead
4
2
0
−2
FW10N
FW10H
FW14N
FW14H
FW18N
FW18H
BW10N
BW10H
BW14N
BW14H
BW18N
BW18H
SW10N
SW10H
SW14N
SW14H
SW18N
SW18H

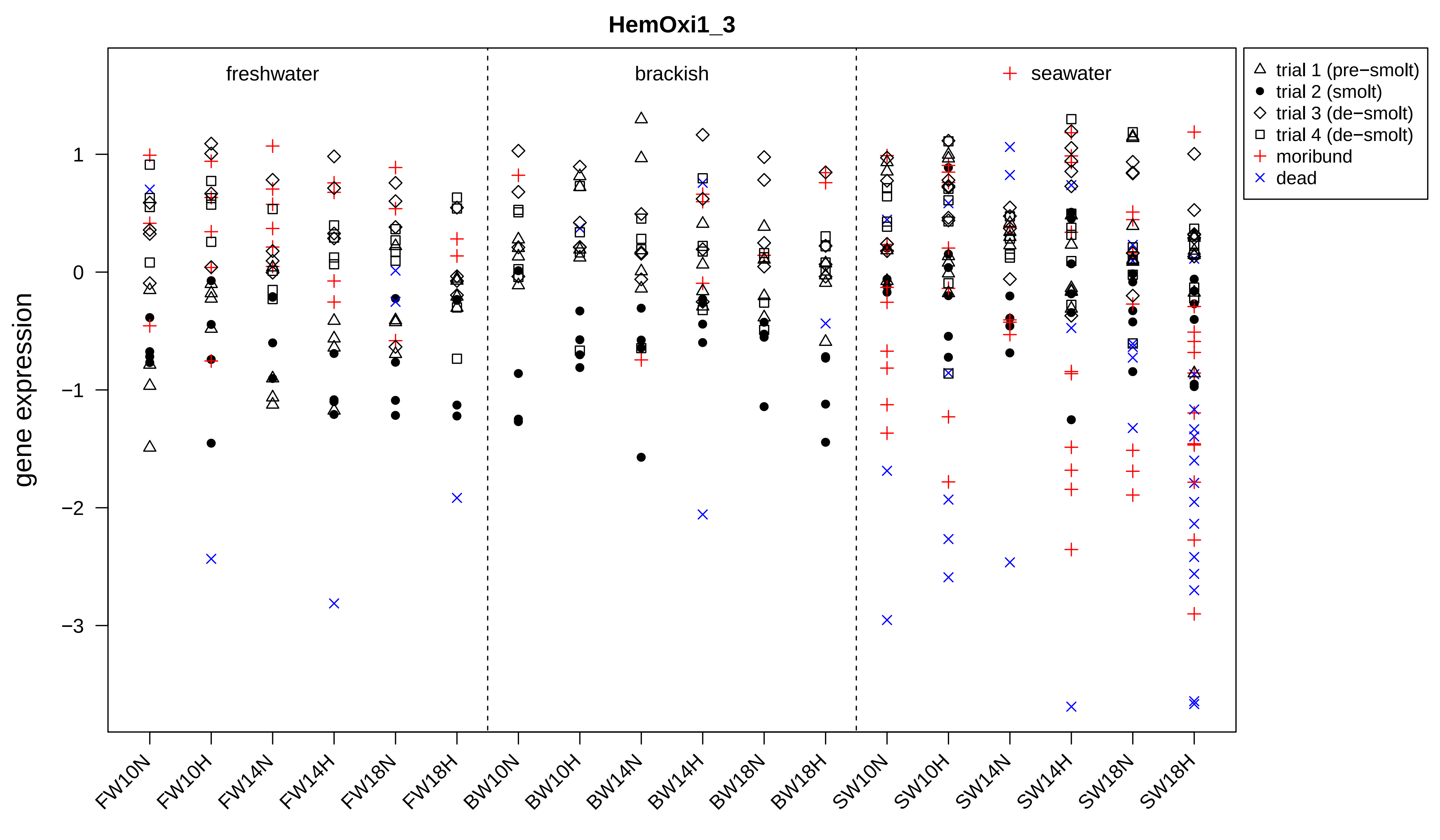

HemOxi1_3
freshwater
brackish
seawater
trial 1 (pre−smolt)
trial 2 (smolt)
trial 3 (de−smolt)
trial 4 (de−smolt)
moribund
dead
gene expression
1
0
−1
−2
−3
FW10N
FW10H
FW14N
FW14H
FW18N
FW18H
BW10N
BW10H
BW14N
BW14H
BW18N
BW18H
SW10N
SW10H
SW14N
SW14H
SW18N
SW18H

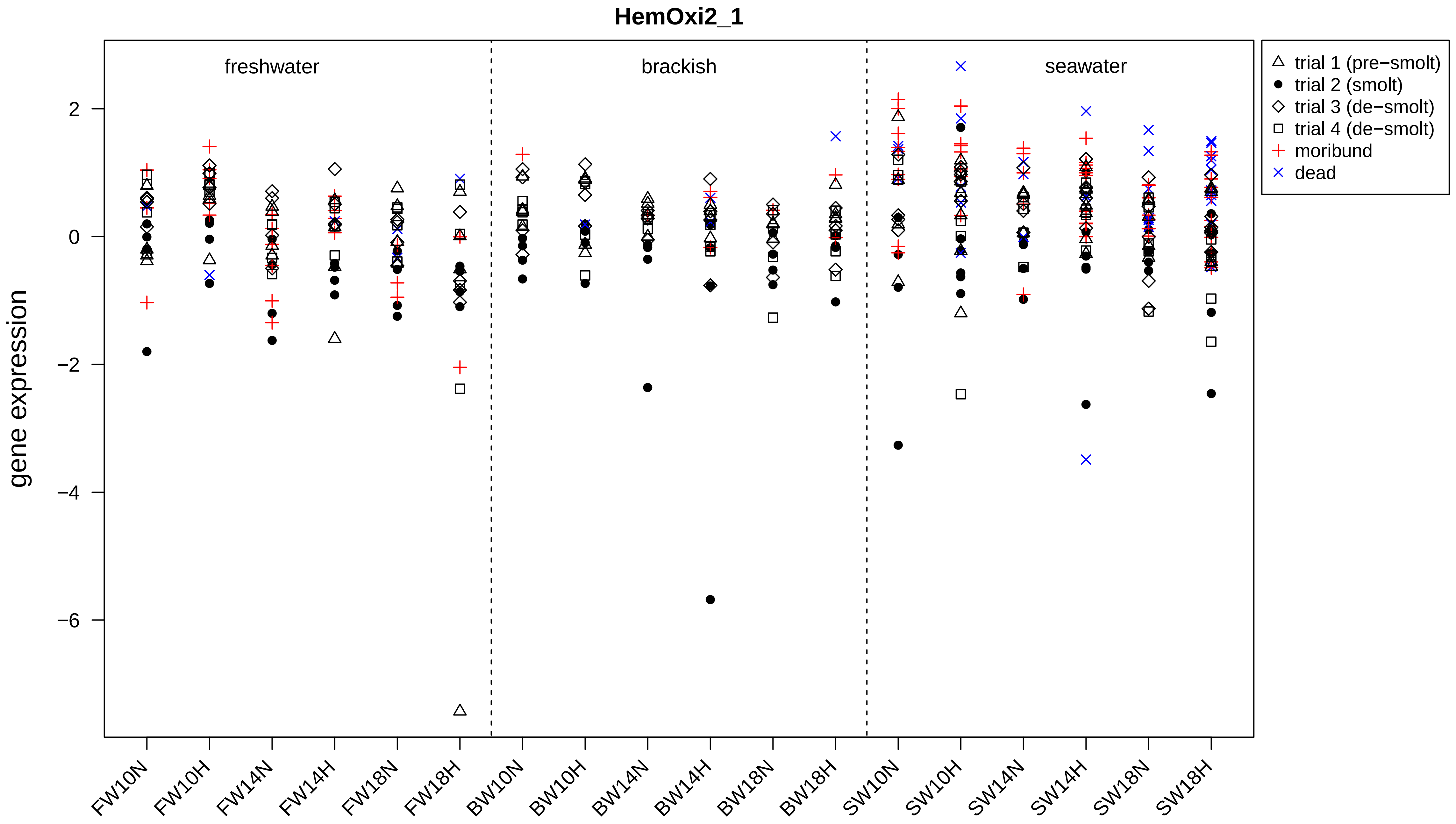

HemOxi2_1
gene expression
freshwater
brackish
seawater
2
0
−2
−4
−6
FW10N
FW10H
FW14N
FW14H
FW18N
FW18H
BW10N
BW10H
BW14N
BW14H
BW18N
BW18H
SW10N
SW10H
SW14N
SW14H
SW18N
SW18H
trial 1 (pre−smolt)
trial 2 (smolt)
trial 3 (de−smolt)
trial 4 (de−smolt)
moribund
dead

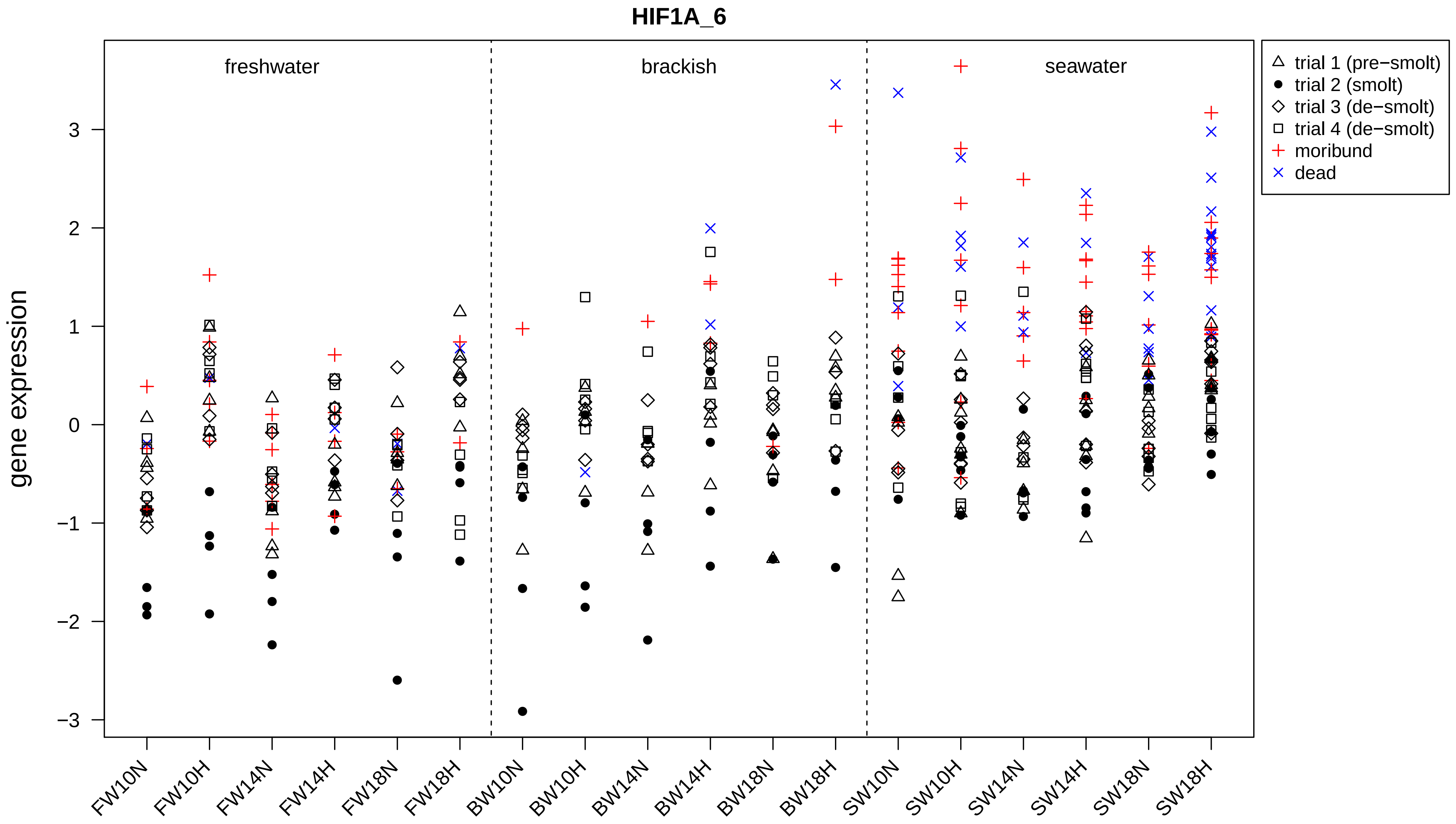

HIF1A_6
freshwater
brackish
seawater
gene expression
3
2
1
0
−1
−2
−3
FW10N
FW10H
FW14N
FW14H
FW18N
FW18H
BW10N
BW10H
BW14N
BW14H
BW18N
BW18H
SW10N
SW10H
SW14N
SW14H
SW18N
SW18H
trial 1 (pre−smolt)
trial 2 (smolt)
trial 3 (de−smolt)
trial 4 (de−smolt)
moribund
dead

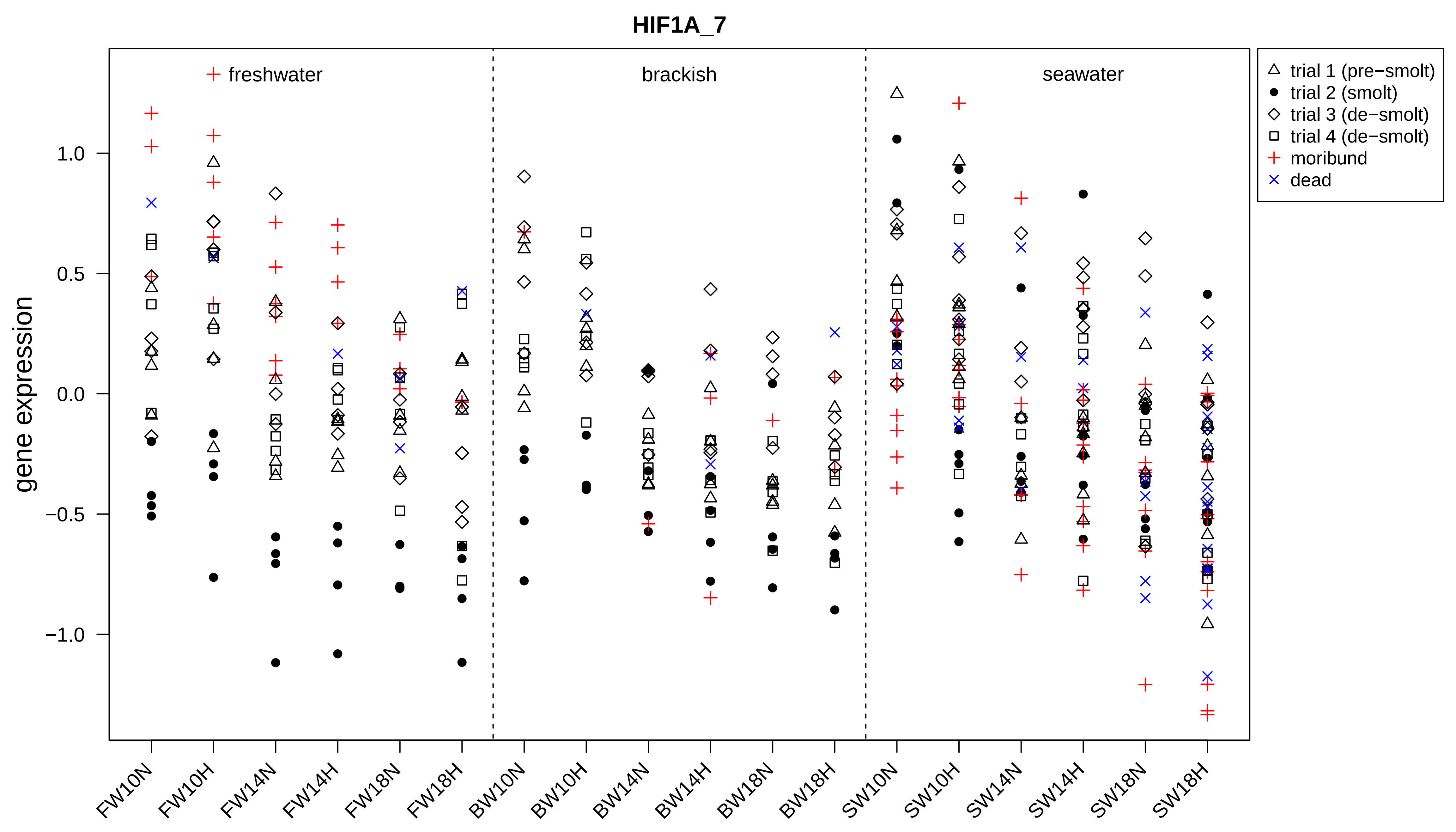

HIF1A_7
freshwater
brackish
seawater
trial 1 (pre−smolt)
trial 2 (smolt)
trial 3 (de−smolt)
trial 4 (de−smolt)
moribund
dead
gene expression
1.0
0.5
0.0
−0.5
−1.0
FW10N
FW10H
FW14N
FW14H
FW18N
FW18H
BW10N
BW10H
BW14N
BW14H
BW18N
BW18H
SW10N
SW10H
SW14N
SW14H
SW18N
SW18H

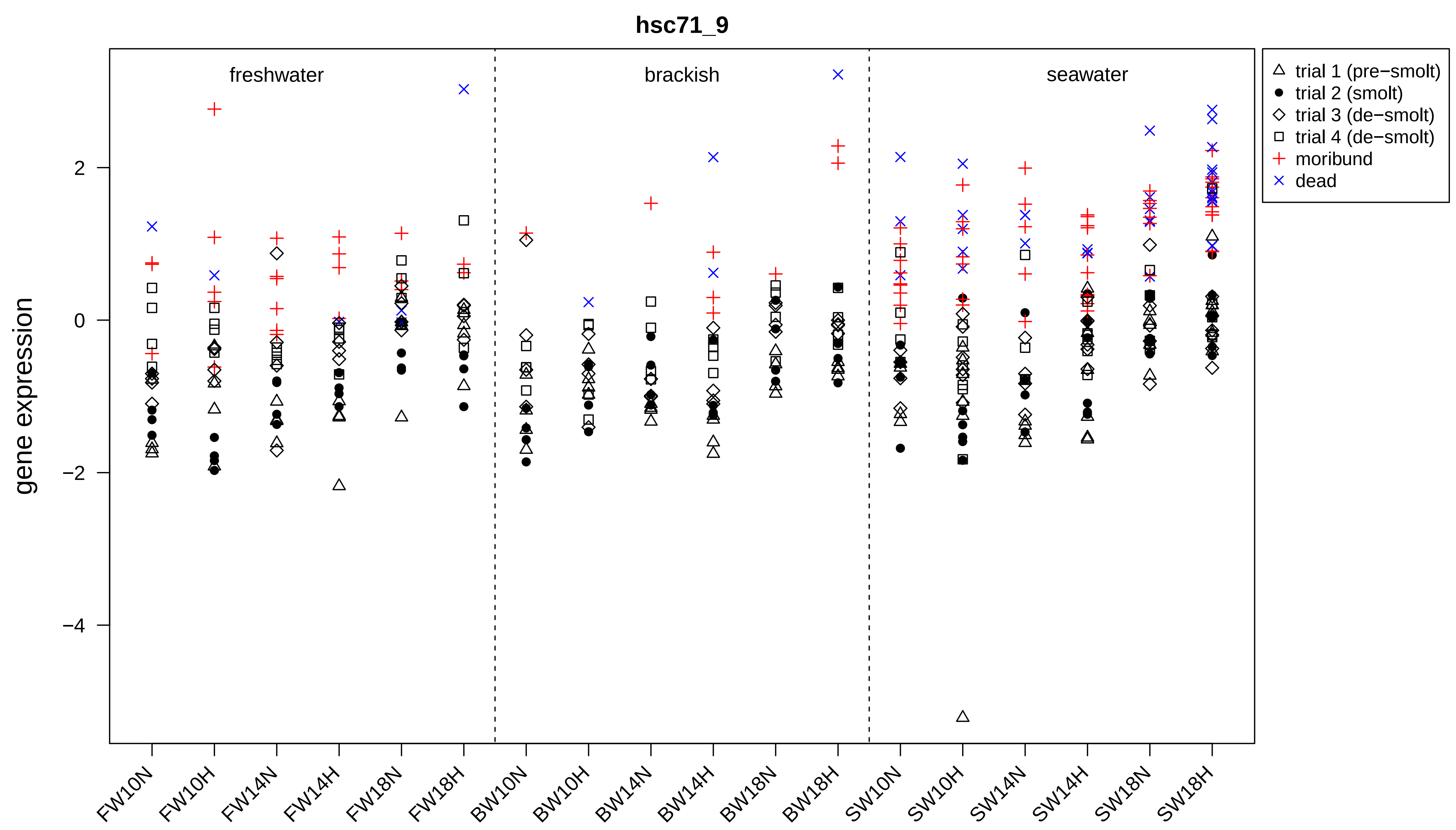
hsc71_9
freshwater
brackish
seawater
gene expression
2
0
−2
−4
FW10N
FW10H
FW14N
FW14H
FW18N
FW18H
BW10N
BW10H
BW14N
BW14H
BW18N
BW18H
SW10N
SW10H
SW14N
SW14H
SW18N
SW18H
trial 1 (pre−smolt)
trial 2 (smolt)
trial 3 (de−smolt)
trial 4 (de−smolt)
moribund
dead

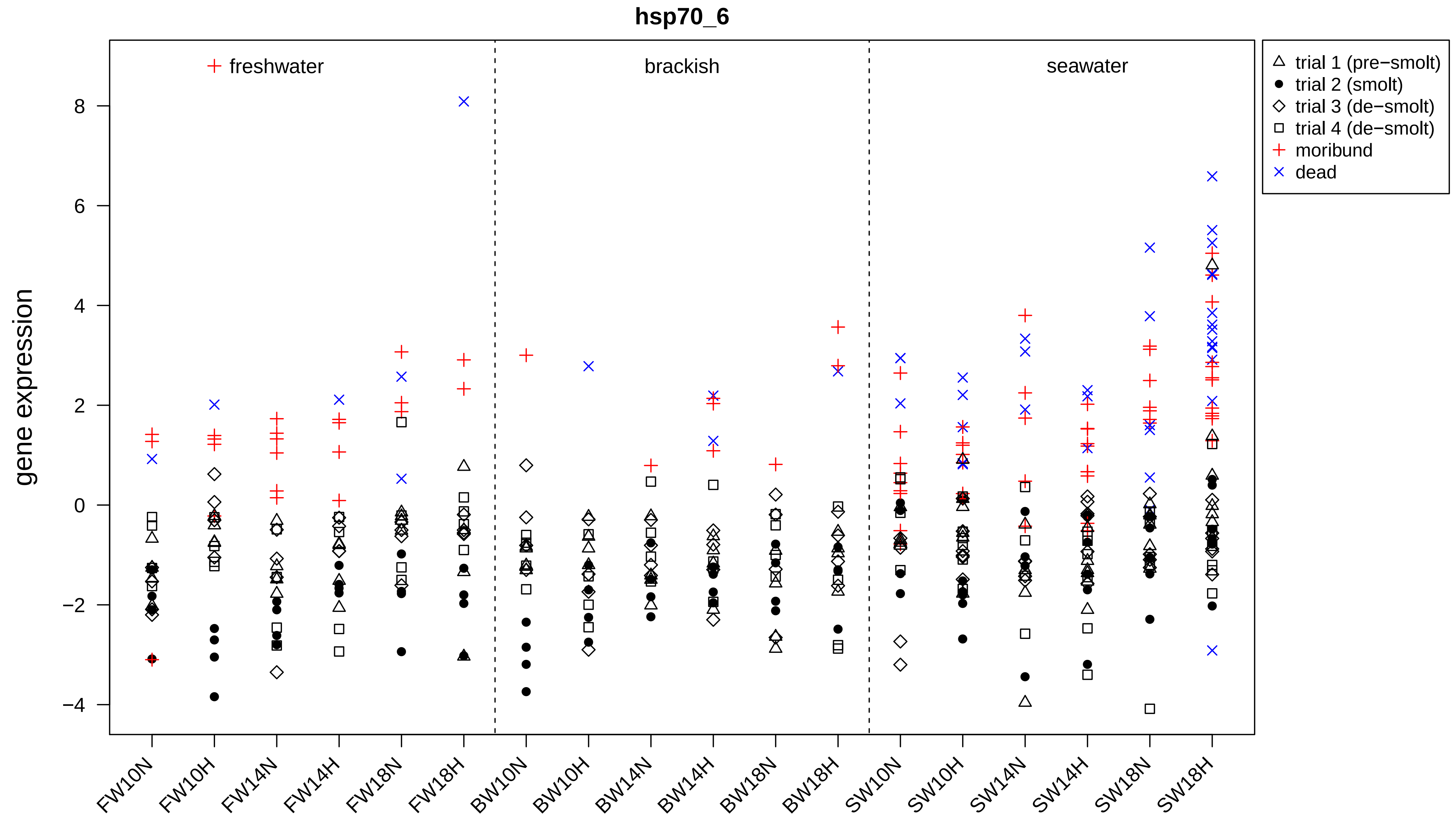

hsp70_6
freshwater
brackish
seawater
trial 1 (pre−smolt)
trial 2 (smolt)
trial 3 (de−smolt)
trial 4 (de−smolt)
moribund
dead
gene expression
8
6
4
2
0
−2
−4
FW10N
FW10H
FW14N
FW14H
FW18N
FW18H
BW10N
BW10H
BW14N
BW14H
BW18N
BW18H
SW10N
SW10H
SW14N
SW14H
SW18N
SW18H

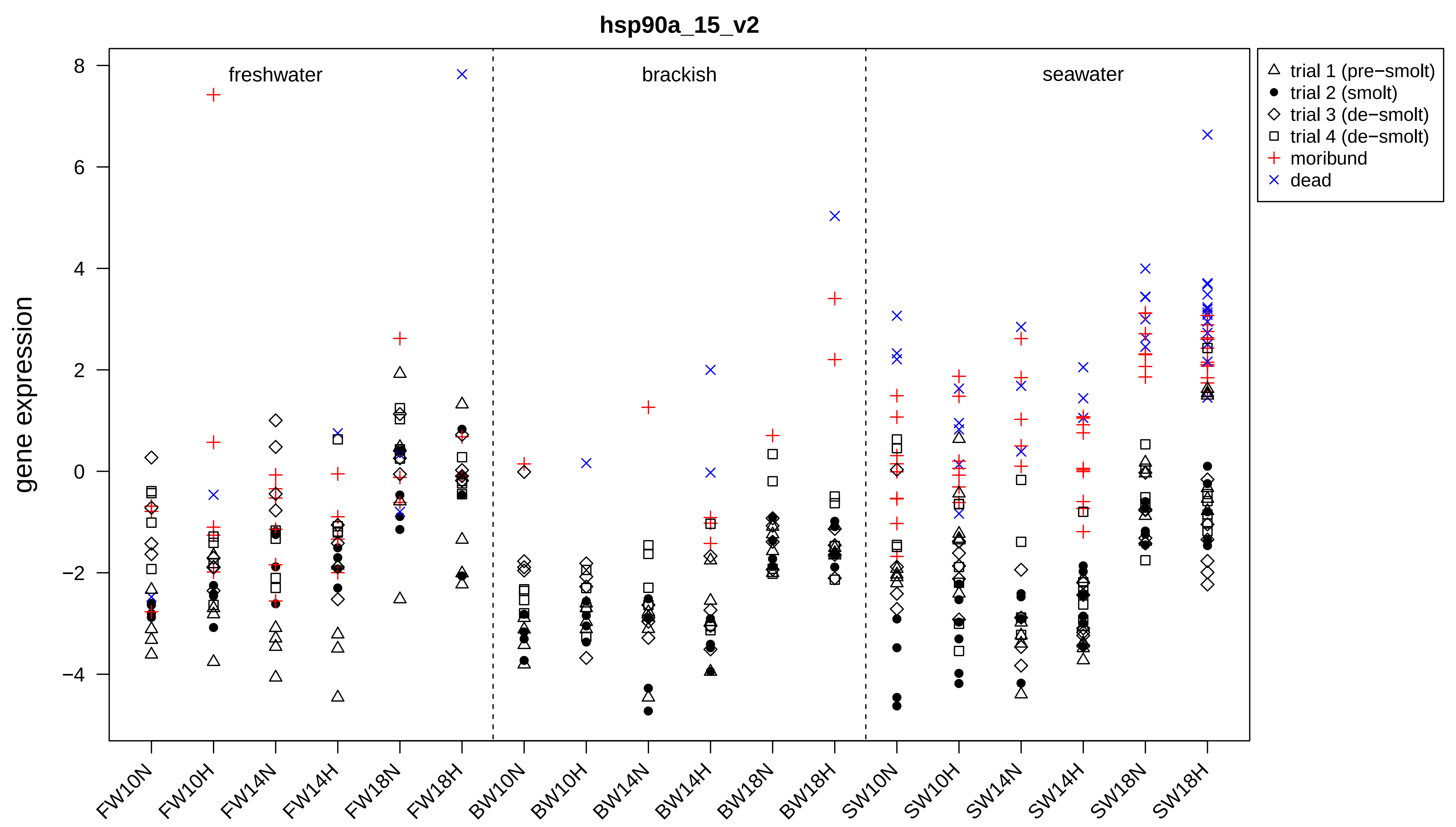
hsp90a_15_v2
freshwater
brackish
seawater
gene expression
8
6
4
2
0
−2
−4
FW10N
FW10H
FW14N
FW14H
FW18N
FW18H
BW10N
BW10H
BW14N
BW14H
BW18N
BW18H
SW10N
SW10H
SW14N
SW14H
SW18N
SW18H
trial 1 (pre−smolt)
trial 2 (smolt)
trial 3 (de−smolt)
trial 4 (de−smolt)
moribund
dead

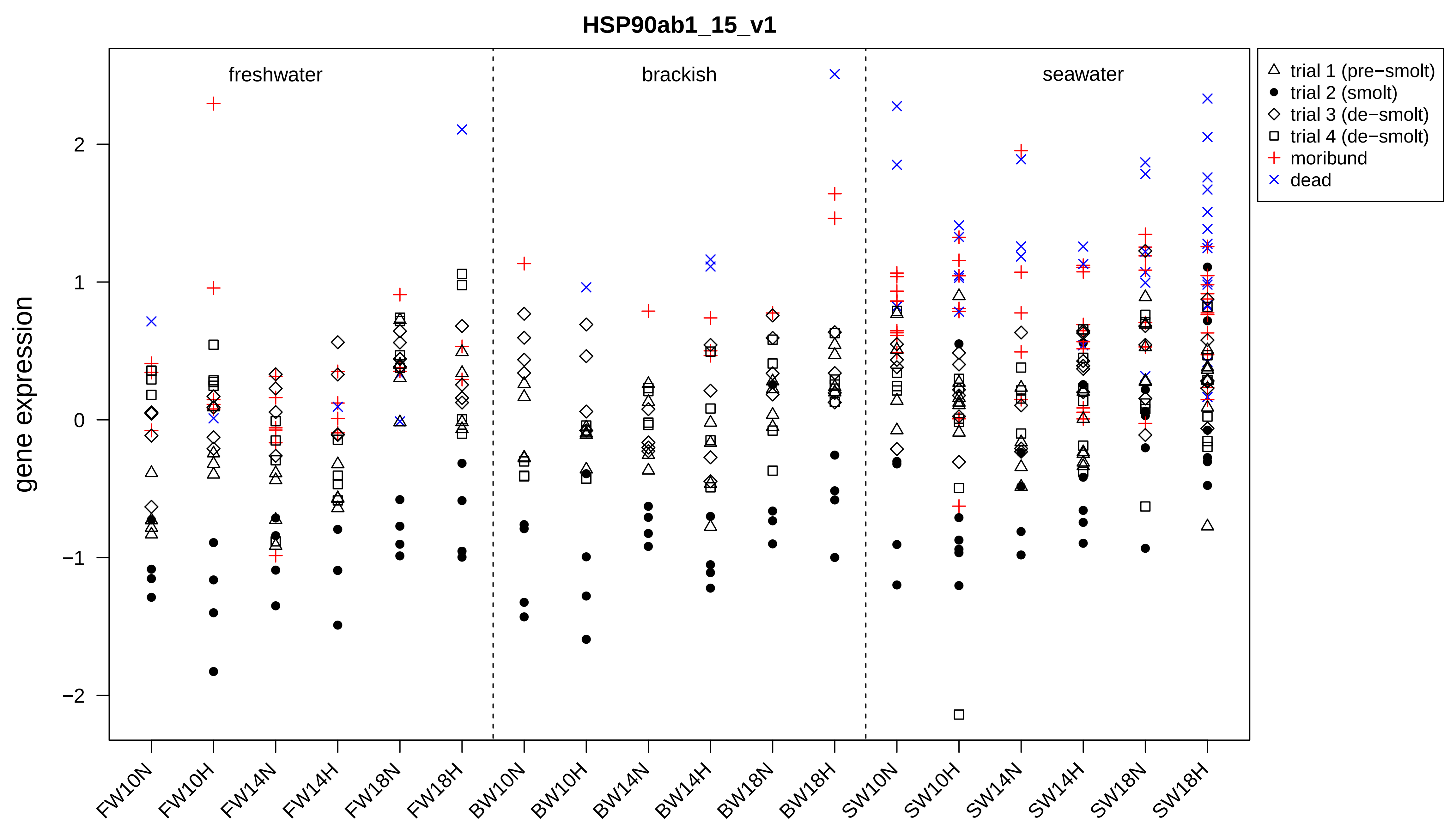

HSP90ab1_15_v1
freshwater
brackish
seawater
gene expression
2
1
0
−1
−2
FW10N
FW10H
FW14N
FW14H
FW18N
FW18H
BW10N
BW10H
BW14N
BW14H
BW18N
BW18H
SW10N
SW10H
SW14N
SW14H
SW18N
SW18H
trial 1 (pre−smolt)
trial 2 (smolt)
trial 3 (de−smolt)
trial 4 (de−smolt)
moribund
dead

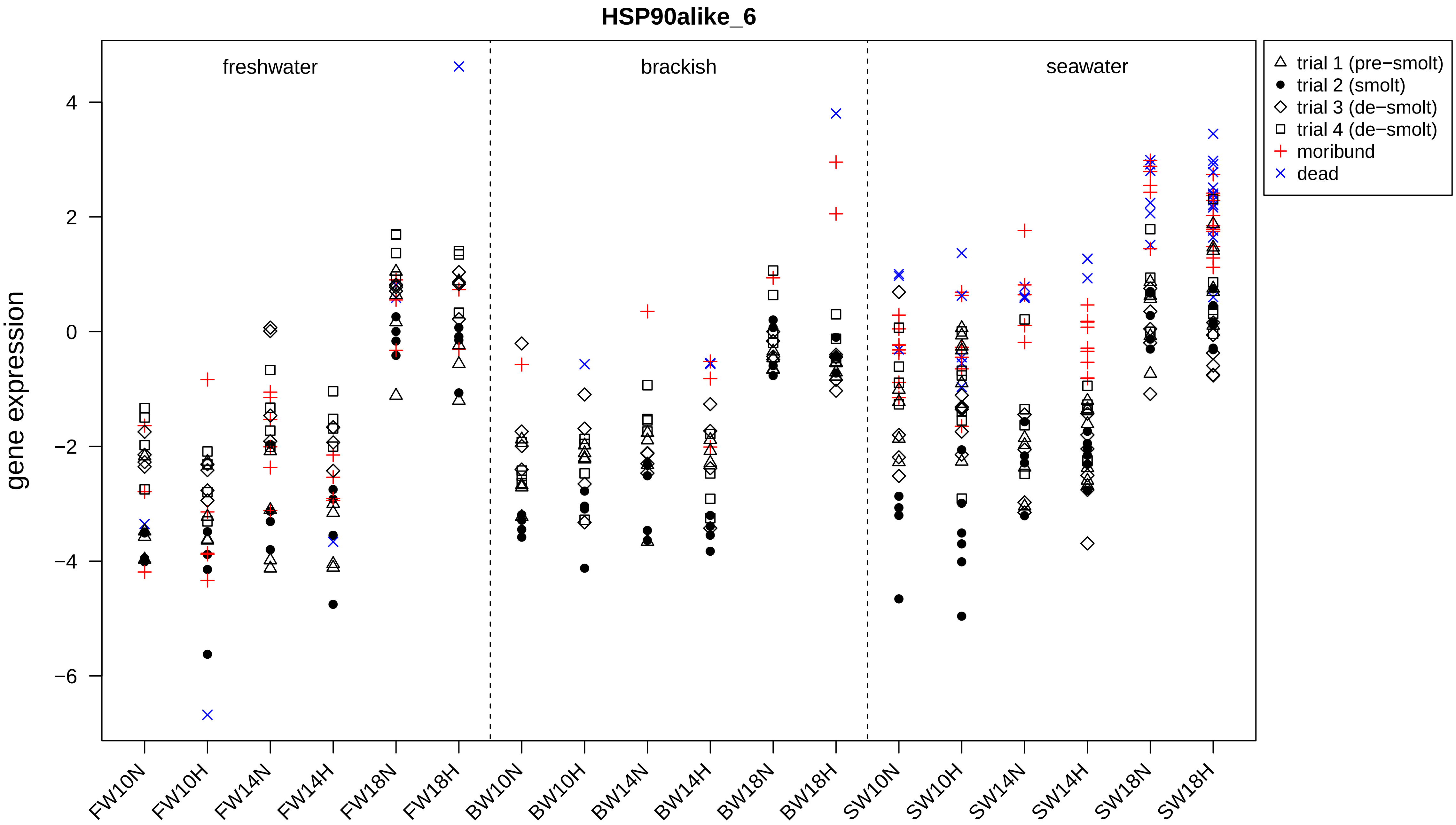
HSP90alike_6
gene expression
freshwater
brackish
seawater
4
2
0
−2
−4
−6
FW10N
FW10H
FW14N
FW14H
FW18N
FW18H
BW10N
BW10H
BW14N
BW14H
BW18N
BW18H
SW10N
SW10H
SW14N
SW14H
SW18N
SW18H
trial 1 (pre−smolt)
trial 2 (smolt)
trial 3 (de−smolt)
trial 4 (de−smolt)
moribund
dead

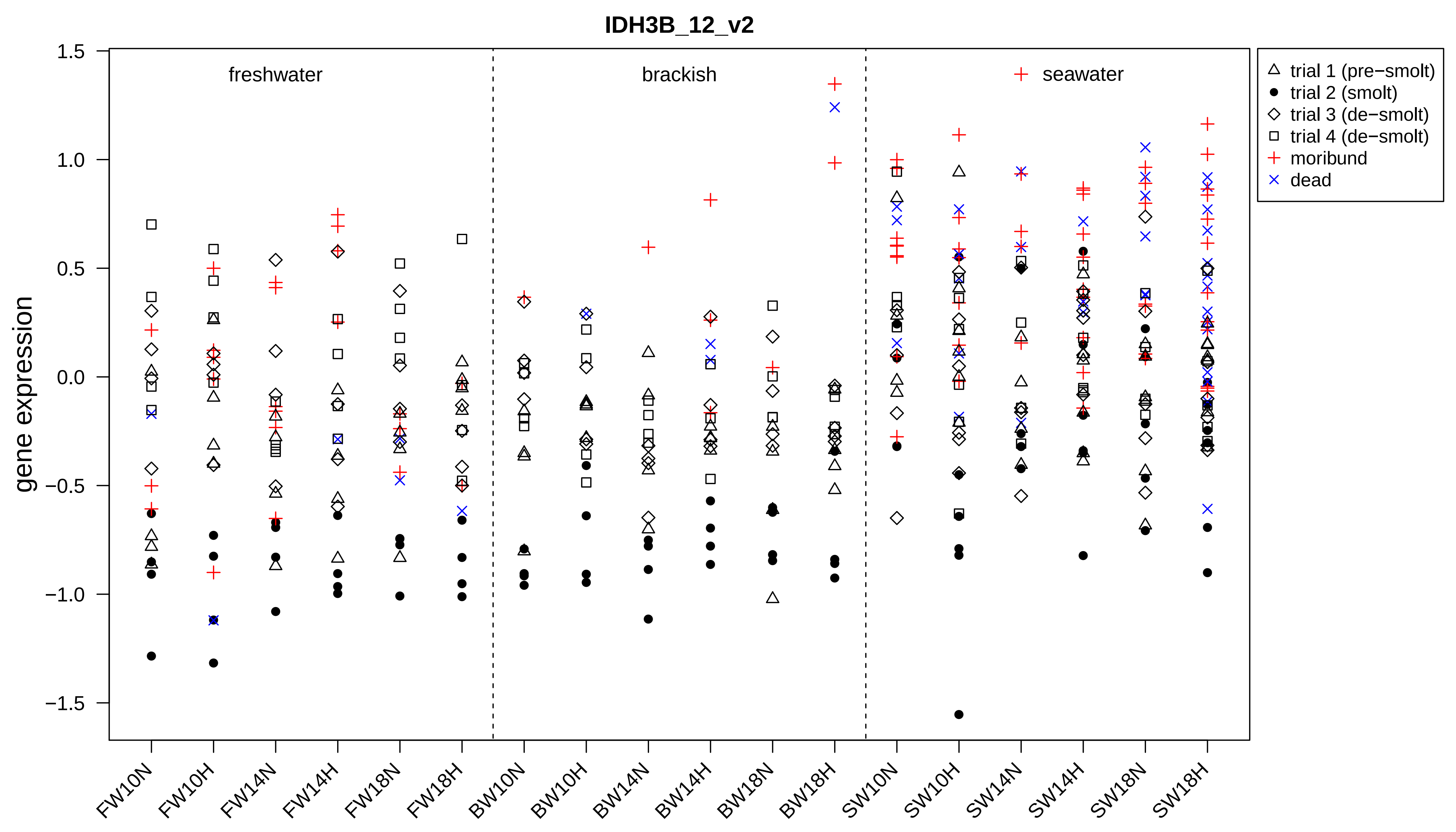

IDH3B_12_v2
gene expression
freshwater
brackish
seawater
1.5
1.0
0.5
0.0
−0.5
−1.0
−1.5
FW10N
FW10H
FW14N
FW14H
FW18N
FW18H
BW10N
BW10H
BW14N
BW14H
BW18N
BW18H
SW10N
SW10H
SW14N
SW14H
SW18N
SW18H
trial 1 (pre−smolt)
trial 2 (smolt)
trial 3 (de−smolt)
trial 4 (de−smolt)
moribund
dead

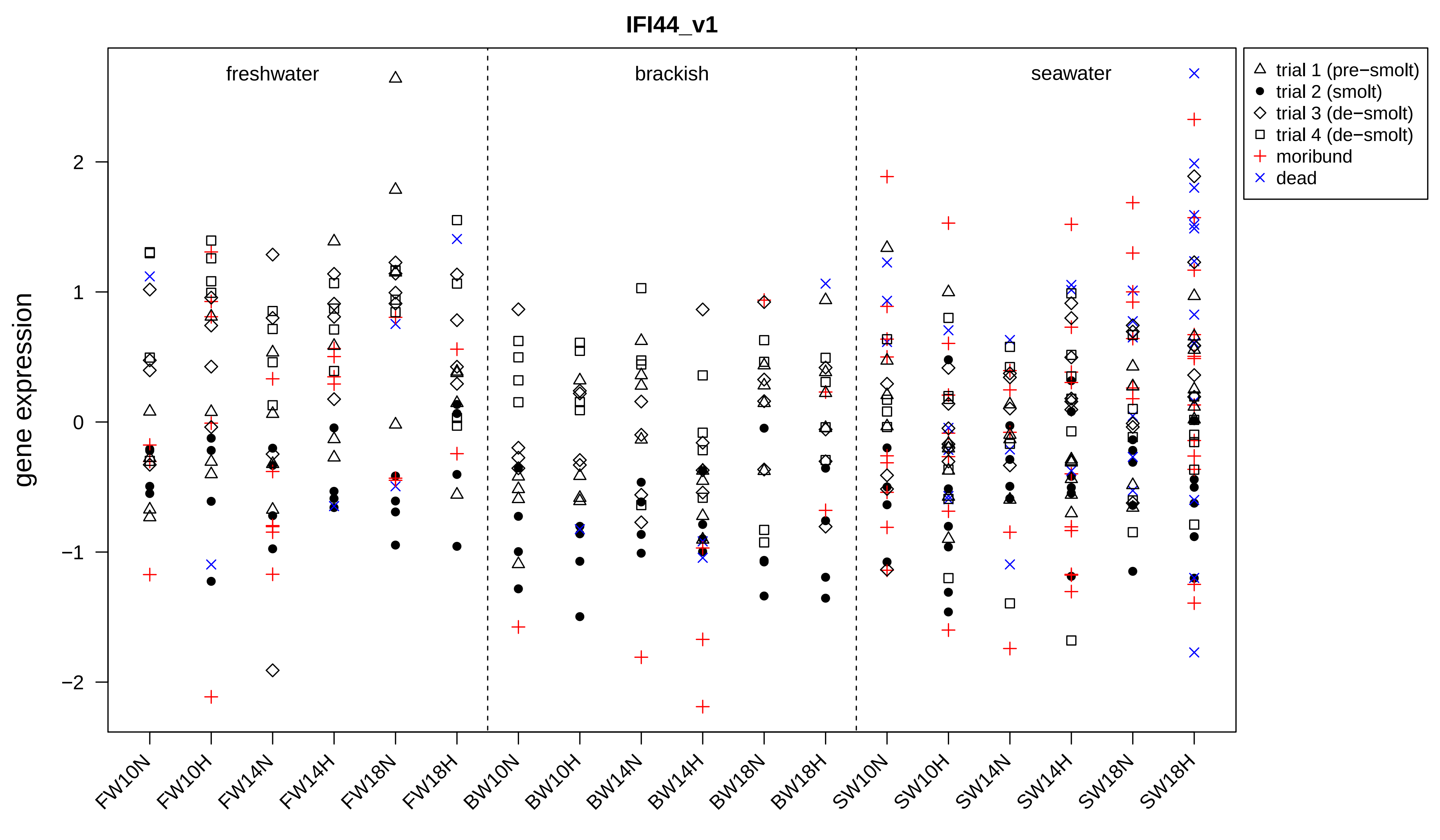

IFI44_v1
gene expression
freshwater
brackish
seawater
trial 1 (pre−smolt)
trial 2 (smolt)
trial 3 (de−smolt)
trial 4 (de−smolt)
moribund
dead
2
1
0
−1
−2
FW10N
FW10H
FW14N
FW14H
FW18N
FW18H
BW10N
BW10H
BW14N
BW14H
BW18N
BW18H
SW10N
SW10H
SW14N
SW14H
SW18N
SW18H

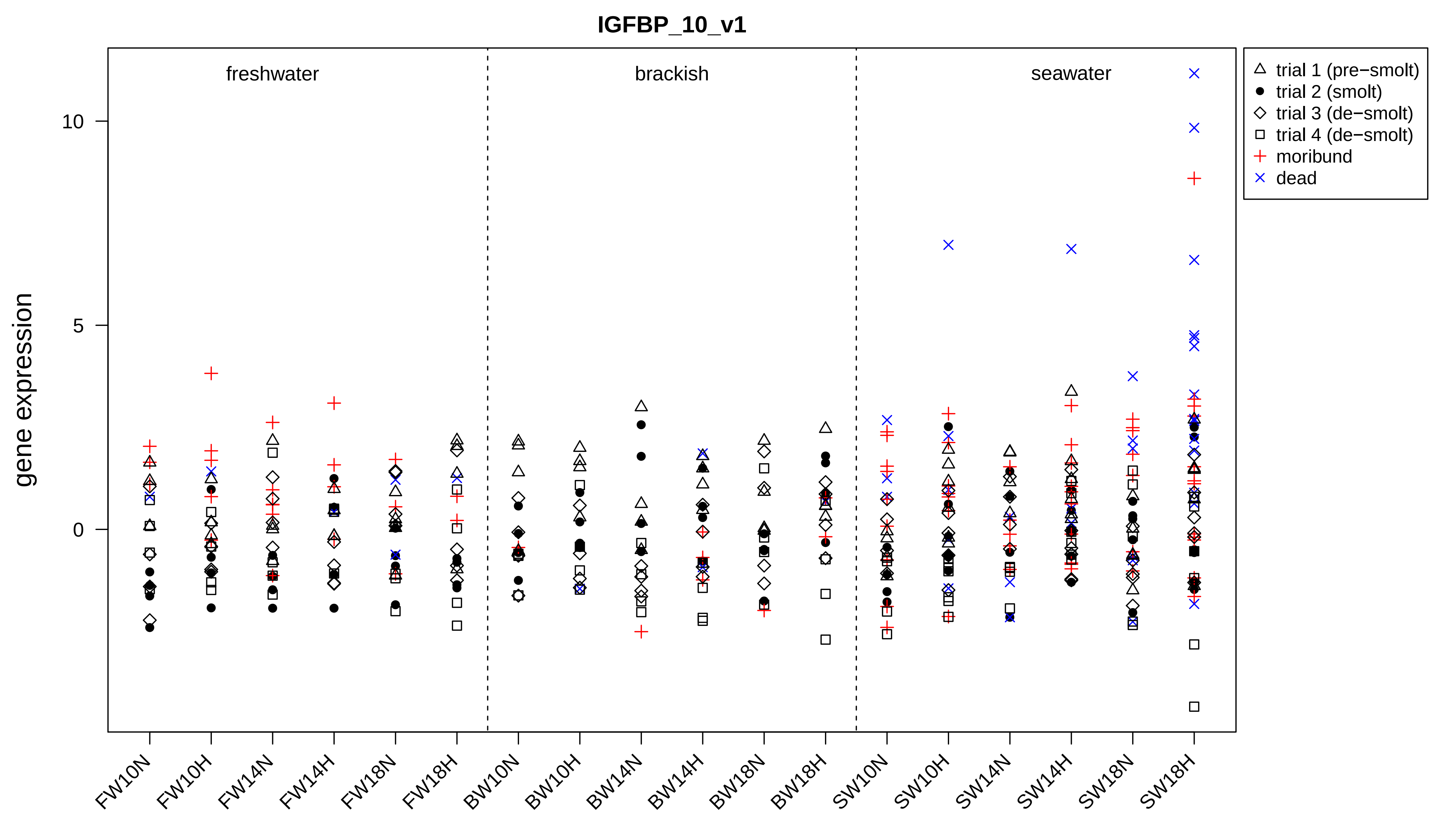

IGFBP_10_v1
freshwater
brackish
seawater
gene expression
10
5
0
trial 1 (pre−smolt)
trial 2 (smolt)
trial 3 (de−smolt)
trial 4 (de−smolt)
moribund
dead
FW10N
FW10H
FW14N
FW14H
FW18N
FW18H
BW10N
BW10H
BW14N
BW14H
BW18N
BW18H
SW10N
SW10H
SW14N
SW14H
SW18N
SW18H

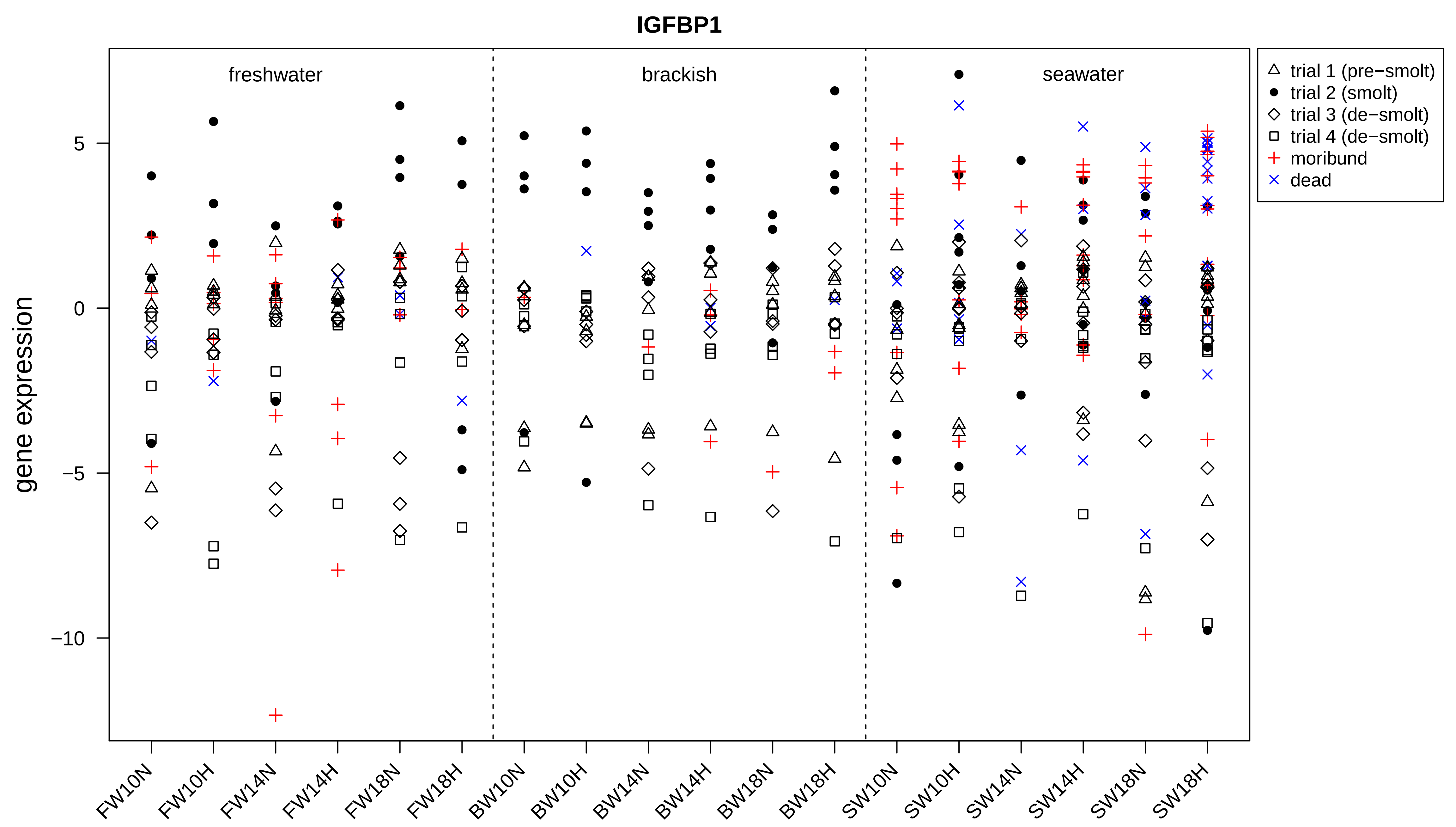

IGFBP1
gene expression
freshwater
brackish
seawater
5
0
−5
−10
FW10N
FW10H
FW14N
FW14H
FW18N
FW18H
BW10N
BW10H
BW14N
BW14H
BW18N
BW18H
SW10N
SW10H
SW14N
SW14H
SW18N
SW18H
trial 1 (pre−smolt)
trial 2 (smolt)
trial 3 (de−smolt)
trial 4 (de−smolt)
moribund
dead

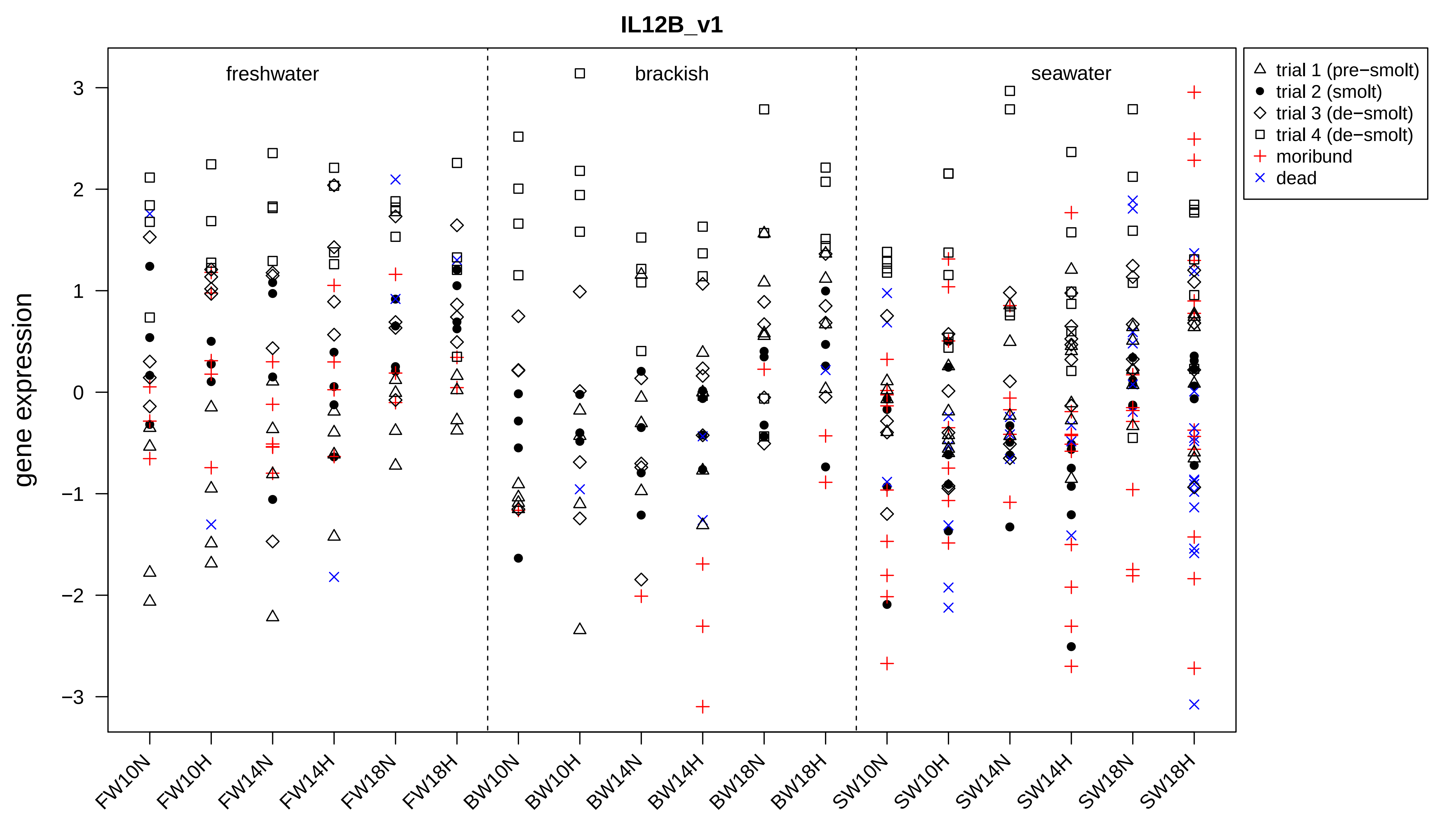
IL12B_v1
freshwater
brackish
seawater
gene expression
3
2
1
0
−1
−2
−3
trial 1 (pre−smolt)
trial 2 (smolt)
trial 3 (de−smolt)
trial 4 (de−smolt)
moribund
dead
FW10N
FW10H
FW14N
FW14H
FW18N
FW18H
BW10N
BW10H
BW14N
BW14H
BW18N
BW18H
SW10N
SW10H
SW14N
SW14H
SW18N
SW18H

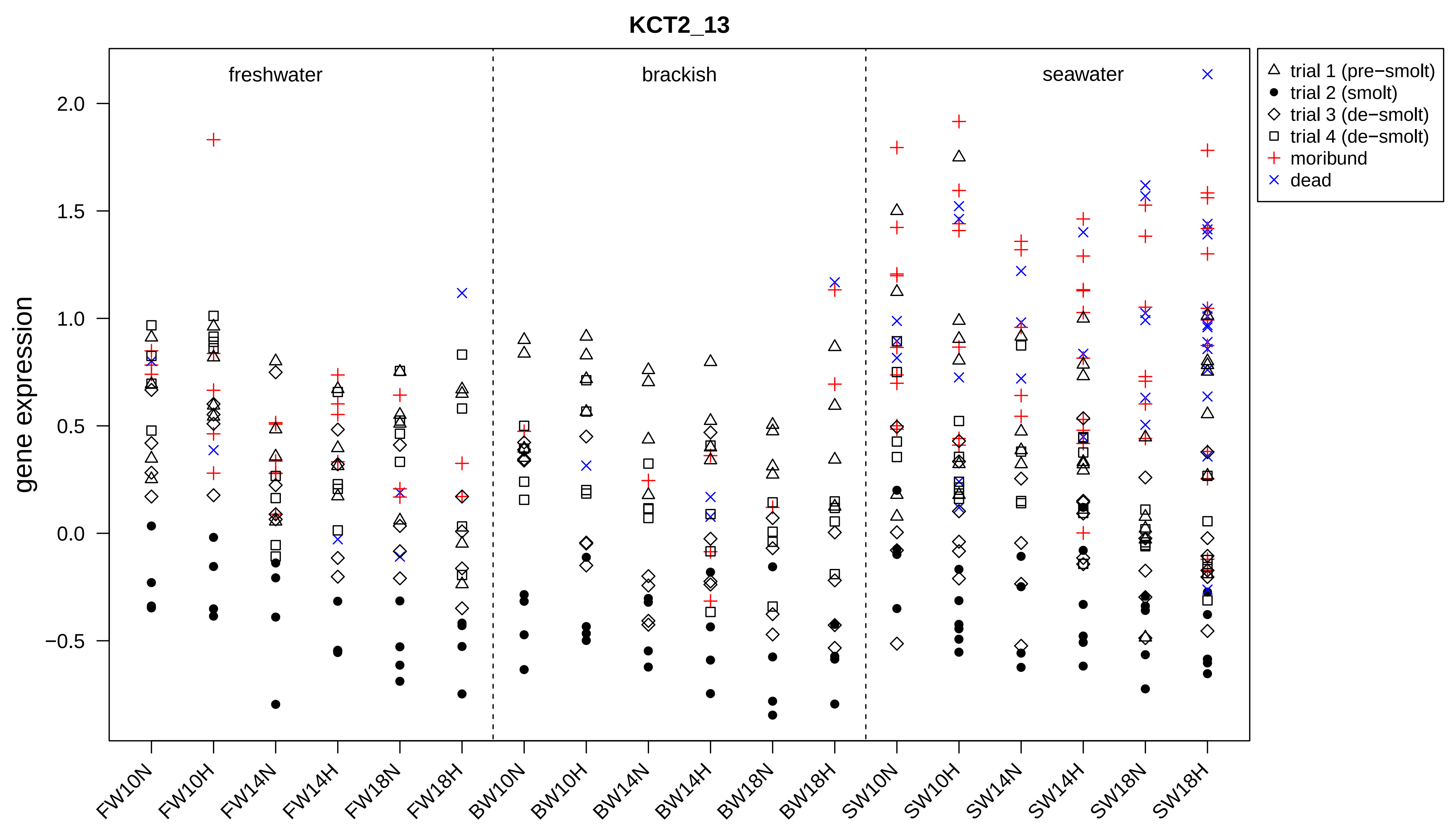

KCT2_13
gene expression
freshwater
brackish
seawater
trial 1 (pre−smolt)
trial 2 (smolt)
trial 3 (de−smolt)
trial 4 (de−smolt)
moribund
dead
2.0
1.5
1.0
0.5
0.0
−0.5
FW10N
FW10H
FW14N
FW14H
FW18N
FW18H
BW10N
BW10H
BW14N
BW14H
BW18N
BW18H
SW10N
SW10H
SW14N
SW14H
SW18N
SW18H

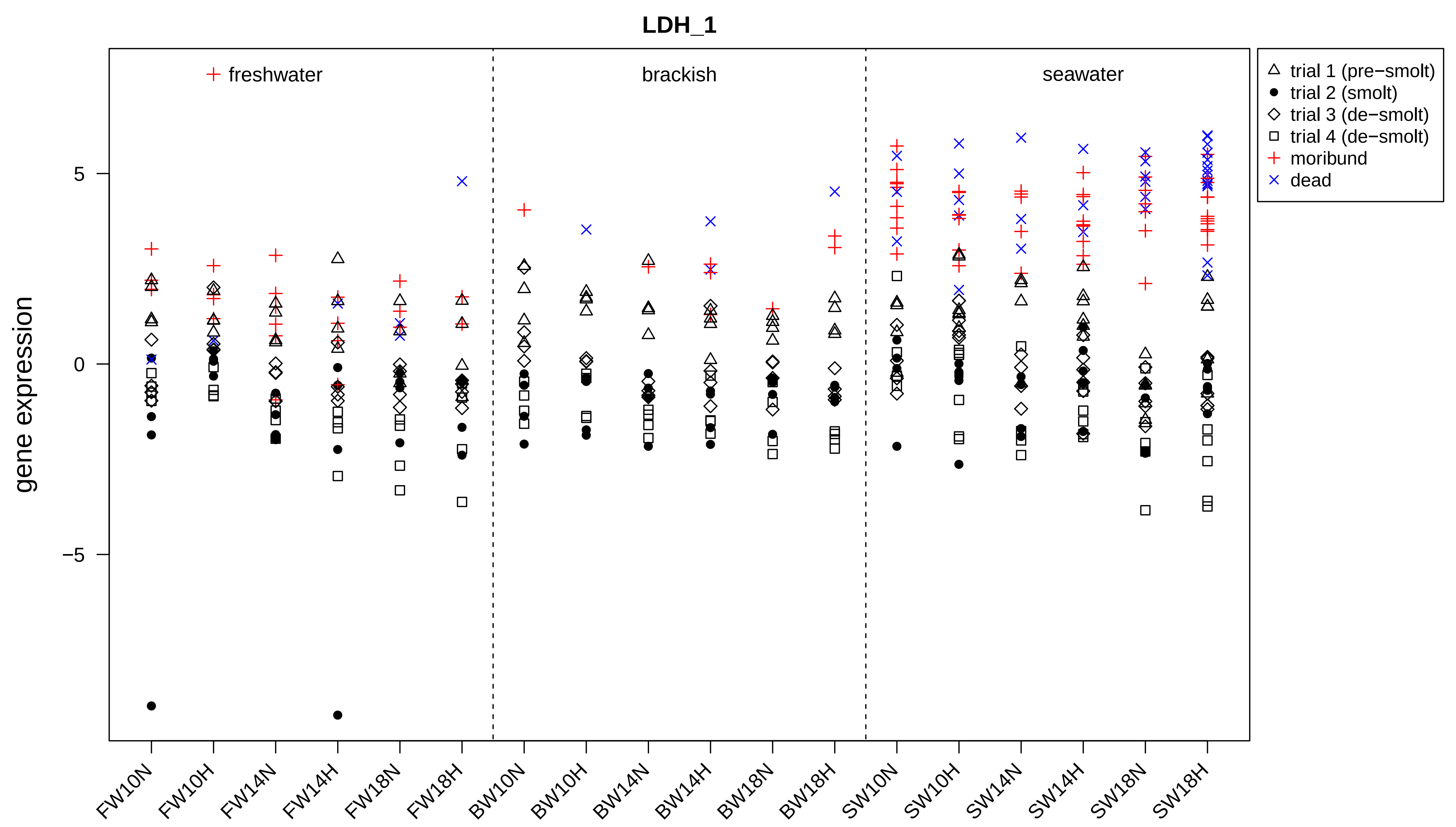

LDH_1
freshwater
brackish
seawater
trial 1 (pre−smolt)
trial 2 (smolt)
trial 3 (de−smolt)
trial 4 (de−smolt)
moribund
dead
gene expression
5
0
−5
FW10N
FW10H
FW14N
FW14H
FW18N
FW18H
BW10N
BW10H
BW14N
BW14H
BW18N
BW18H
SW10N
SW10H
SW14N
SW14H
SW18N
SW18H

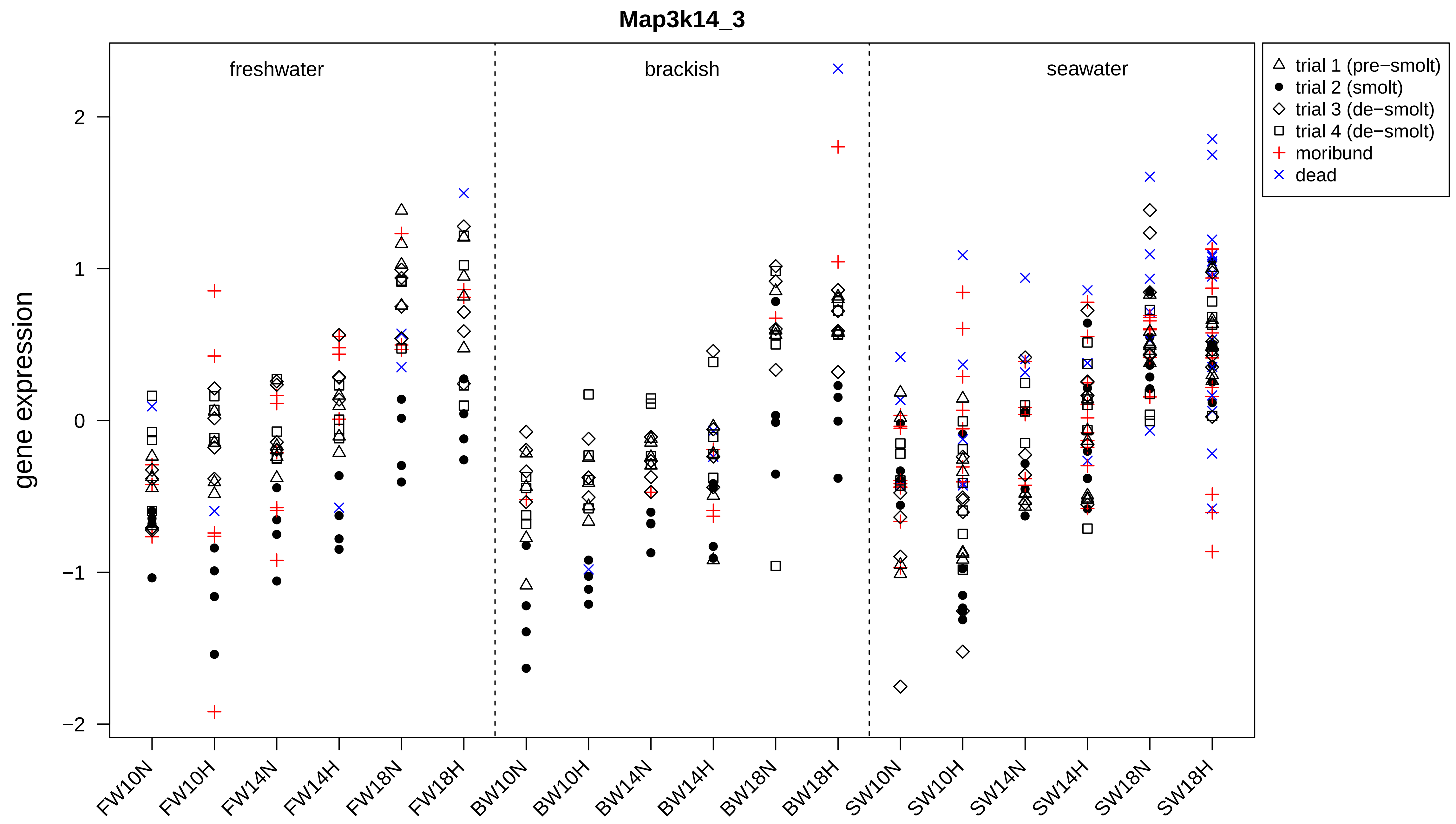
Map3k14_3
freshwater
brackish
seawater
trial 1 (pre−smolt)
trial 2 (smolt)
trial 3 (de−smolt)
trial 4 (de−smolt)
moribund
dead
gene expression
2
1
0
−1
−2
FW10N
FW10H
FW14N
FW14H
FW18N
FW18H
BW10N
BW10H
BW14N
BW14H
BW18N
BW18H
SW10N
SW10H
SW14N
SW14H
SW18N
SW18H

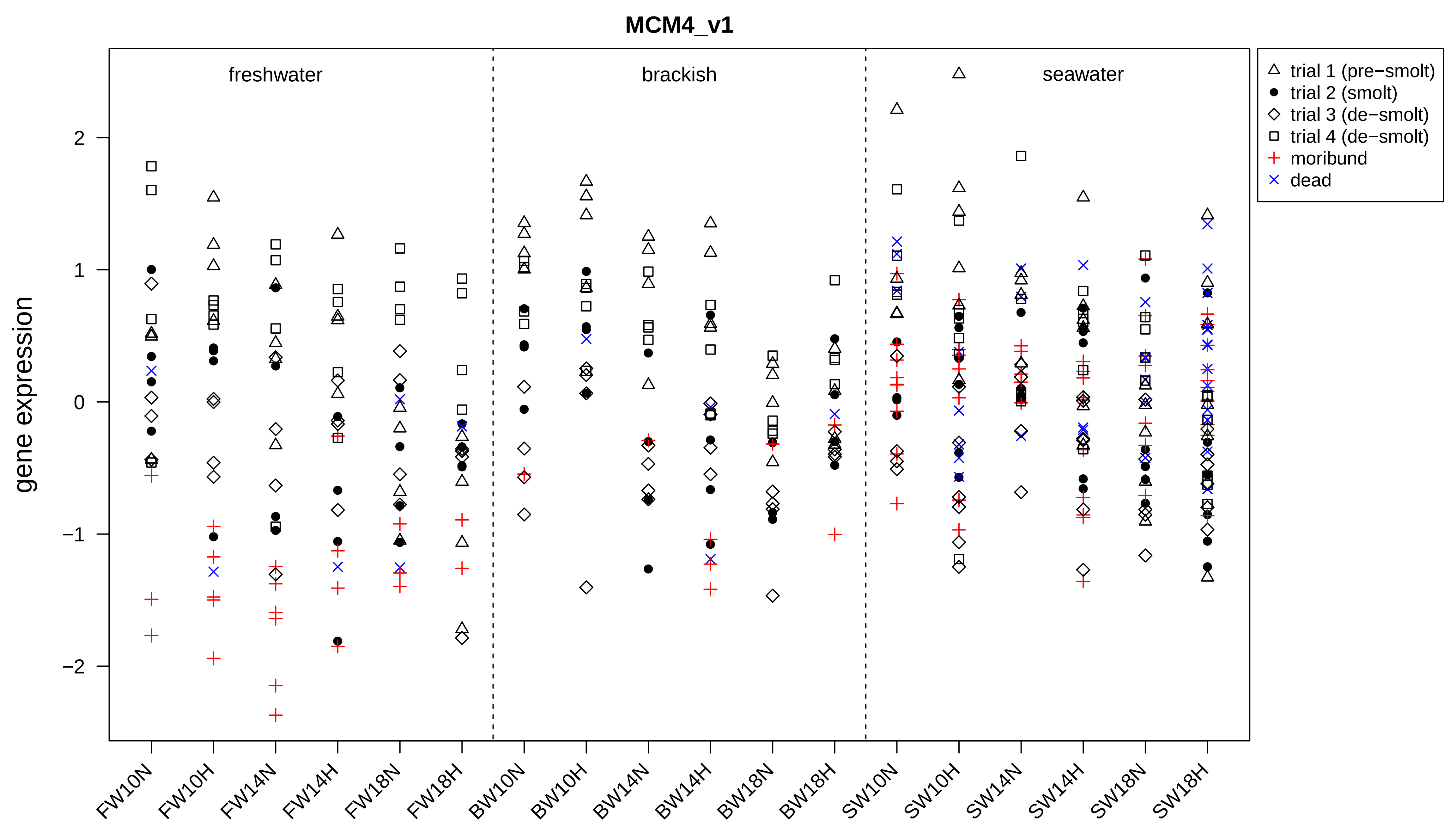
MCM4_v1
gene expression
freshwater
brackish
seawater
trial 1 (pre−smolt)
trial 2 (smolt)
trial 3 (de−smolt)
trial 4 (de−smolt)
moribund
dead
2
1
0
−1
−2
FW10N
FW10H
FW14N
FW14H
FW18N
FW18H
BW10N
BW10H
BW14N
BW14H
BW18N
BW18H
SW10N
SW10H
SW14N
SW14H
SW18N
SW18H

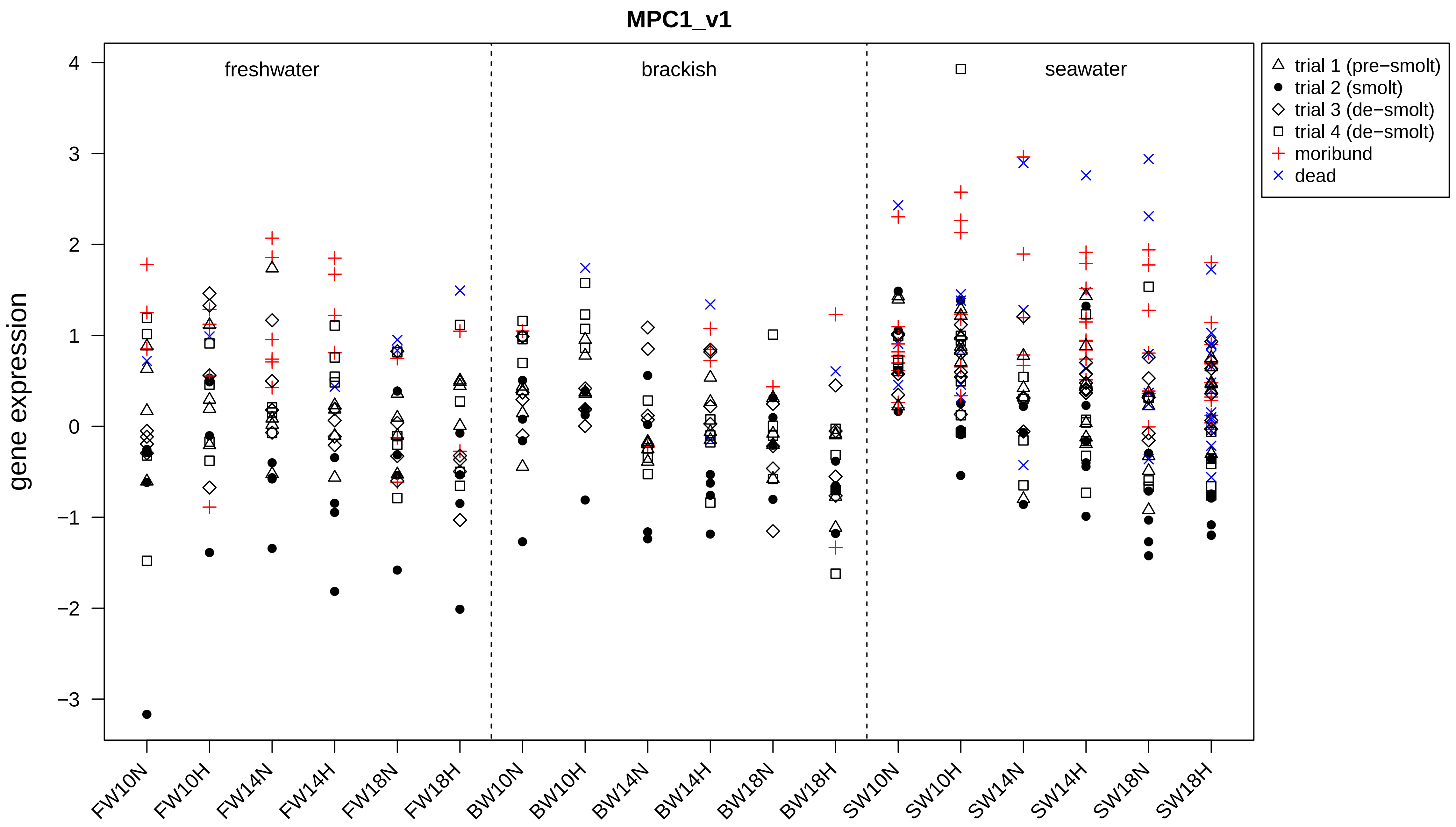
MPC1_v1
freshwater
brackish
seawater
gene expression
4
3
2
1
0
−1
−2
−3
FW10N
FW10H
FW14N
FW14H
FW18N
FW18H
BW10N
BW10H
BW14N
BW14H
BW18N
BW18H
SW10N
SW10H
SW14N
SW14H
SW18N
SW18H
trial 1 (pre−smolt)
trial 2 (smolt)
trial 3 (de−smolt)
trial 4 (de−smolt)
moribund
dead

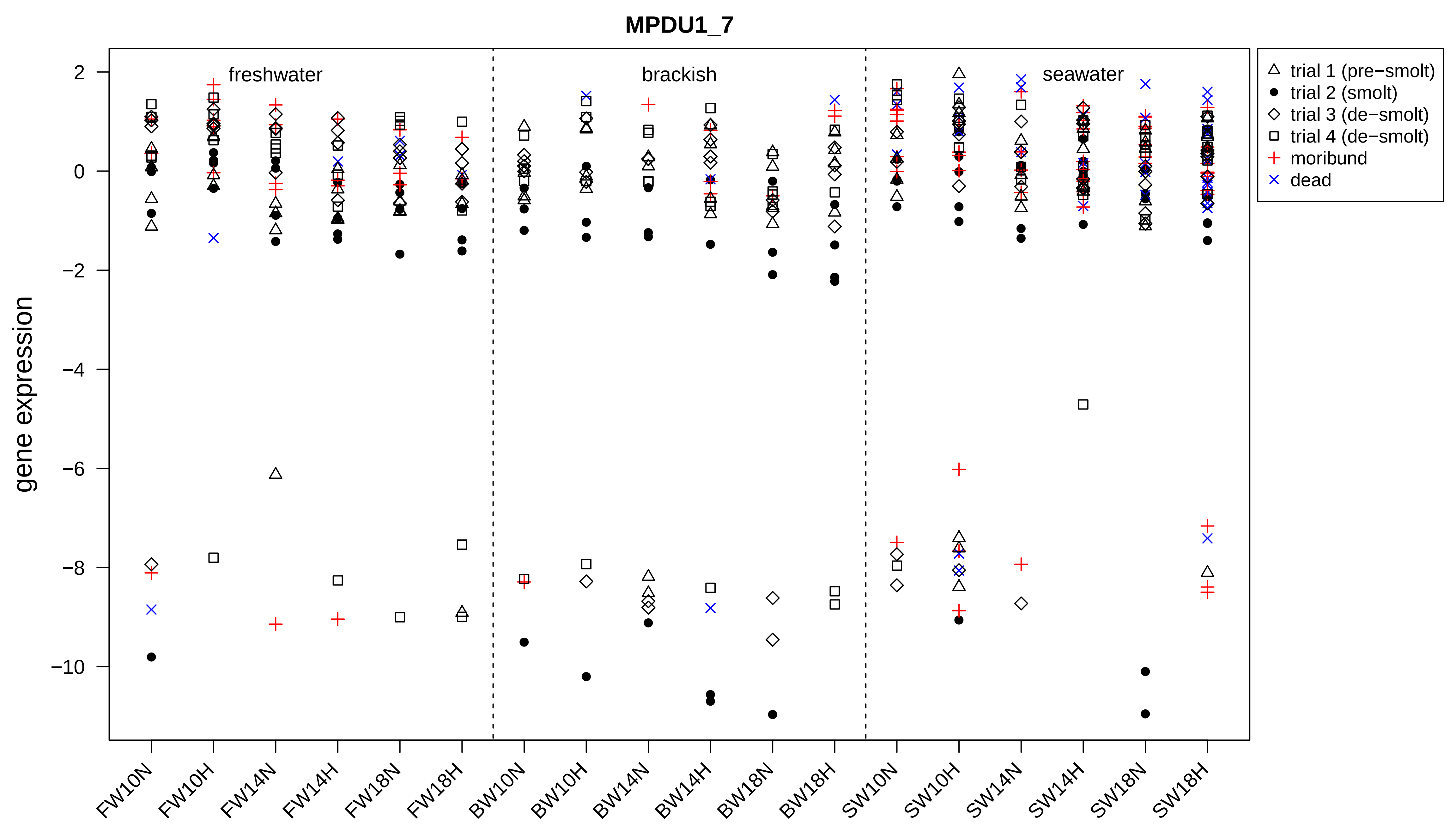

MPDU1_7
gene expression
2
0
−2
−4
−6
−8
−10
freshwater
brackish
seawater
FW10N
FW10H
FW14N
FW14H
FW18N
FW18H
BW10N
BW10H
BW14N
BW14H
BW18N
BW18H
SW10N
SW10H
SW14N
SW14H
SW18N
SW18H
trial 1 (pre−smolt)
trial 2 (smolt)
trial 3 (de−smolt)
trial 4 (de−smolt)
moribund
dead

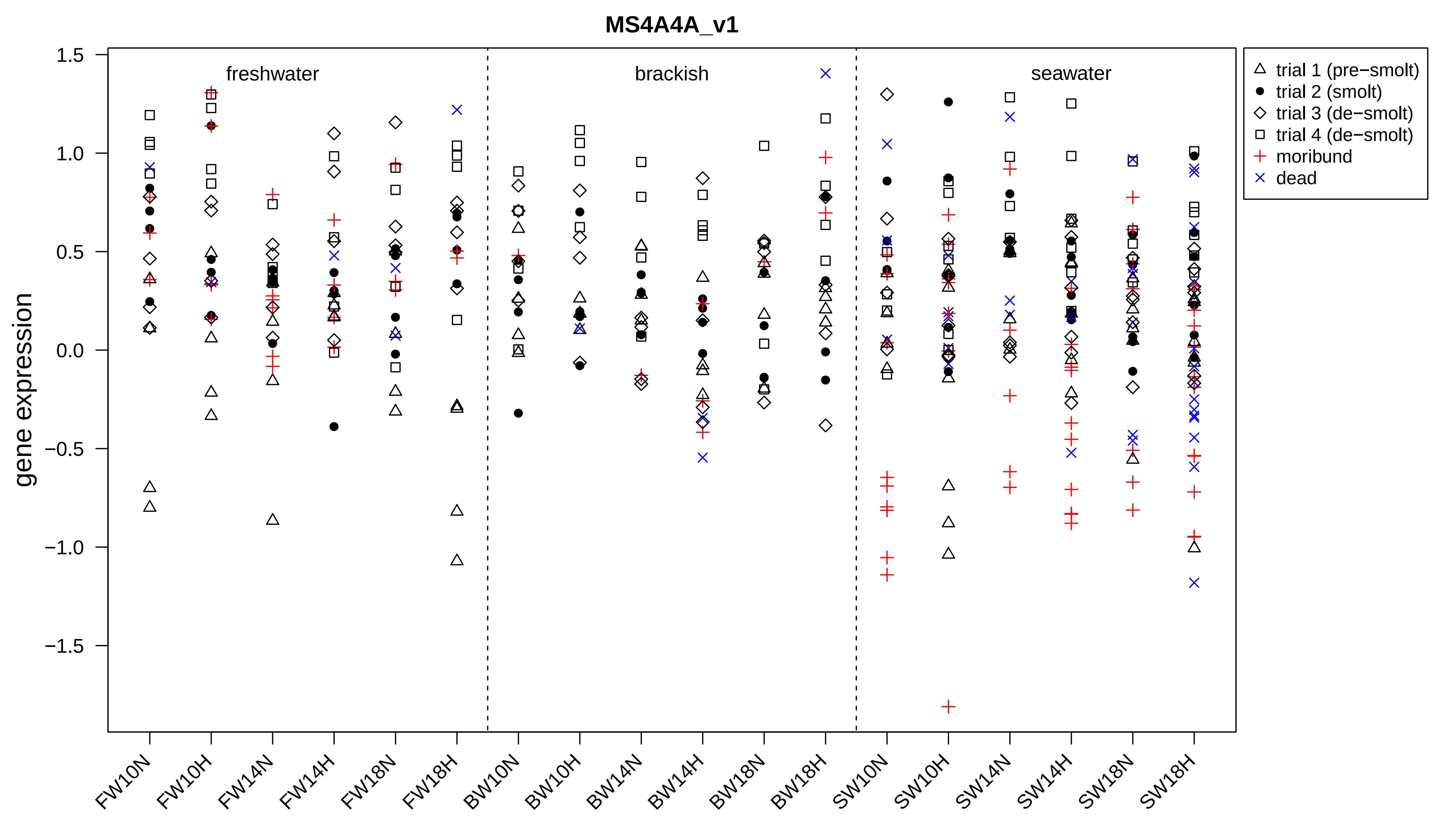
MS4A4A_v1
gene expression
freshwater
brackish
seawater
trial 1 (pre−smolt)
trial 2 (smolt)
trial 3 (de−smolt)
trial 4 (de−smolt)
moribund
dead
1.5
1.0
0.5
0.0
−0.5
−1.0
−1.5
FW10N
FW10H
FW14N
FW14H
FW18N
FW18H
BW10N
BW10H
BW14N
BW14H
BW18N
BW18H
SW10N
SW10H
SW14N
SW14H
SW18N
SW18H

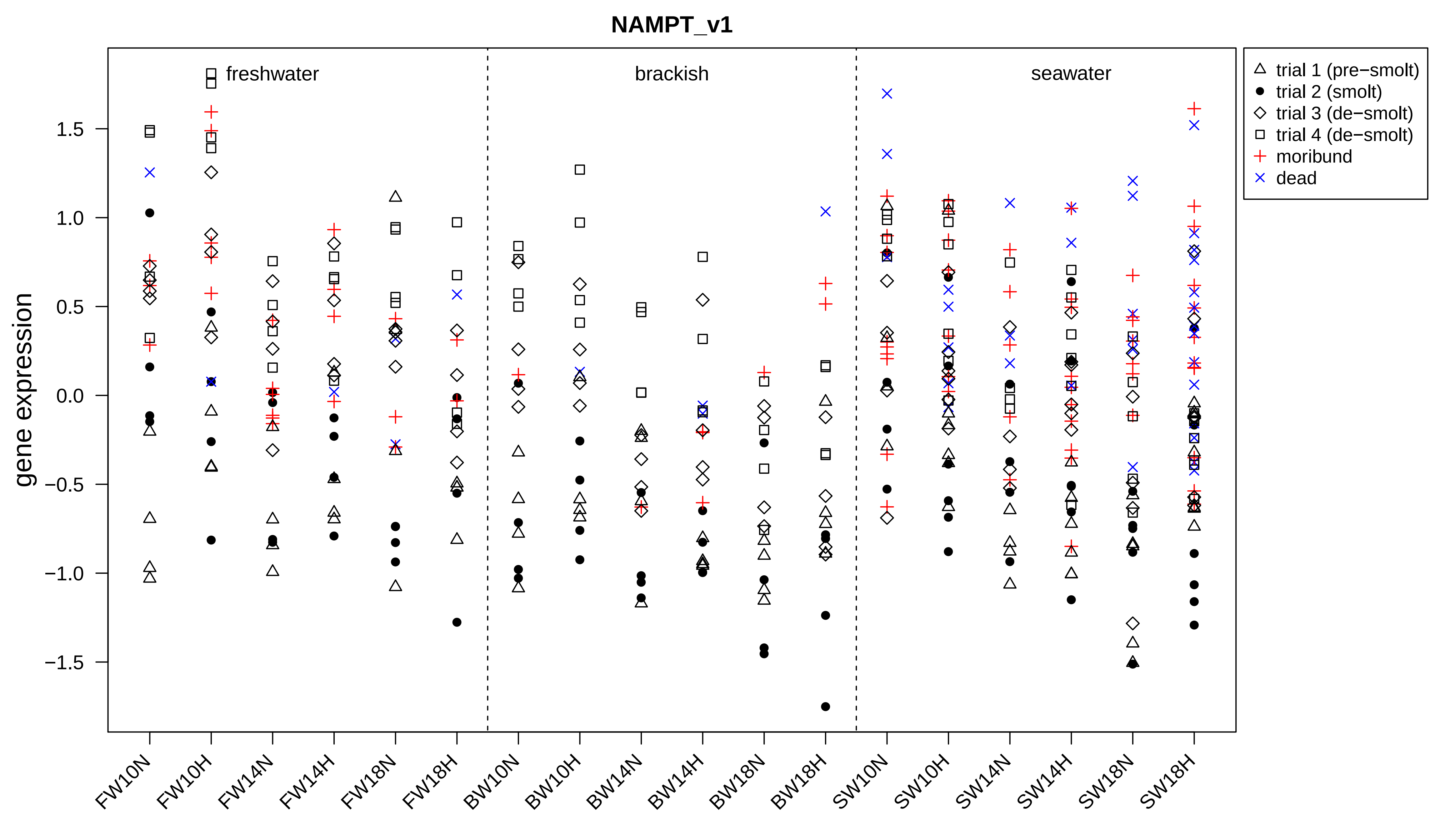

NAMPT_v1
freshwater
brackish
seawater
trial 1 (pre−smolt)
trial 2 (smolt)
trial 3 (de−smolt)
trial 4 (de−smolt)
moribund
dead
gene expression
1.5
1.0
0.5
0.0
−0.5
−1.0
−1.5
FW10N
FW10H
FW14N
FW14H
FW18N
FW18H
BW10N
BW10H
BW14N
BW14H
BW18N
BW18H
SW10N
SW10H
SW14N
SW14H
SW18N
SW18H

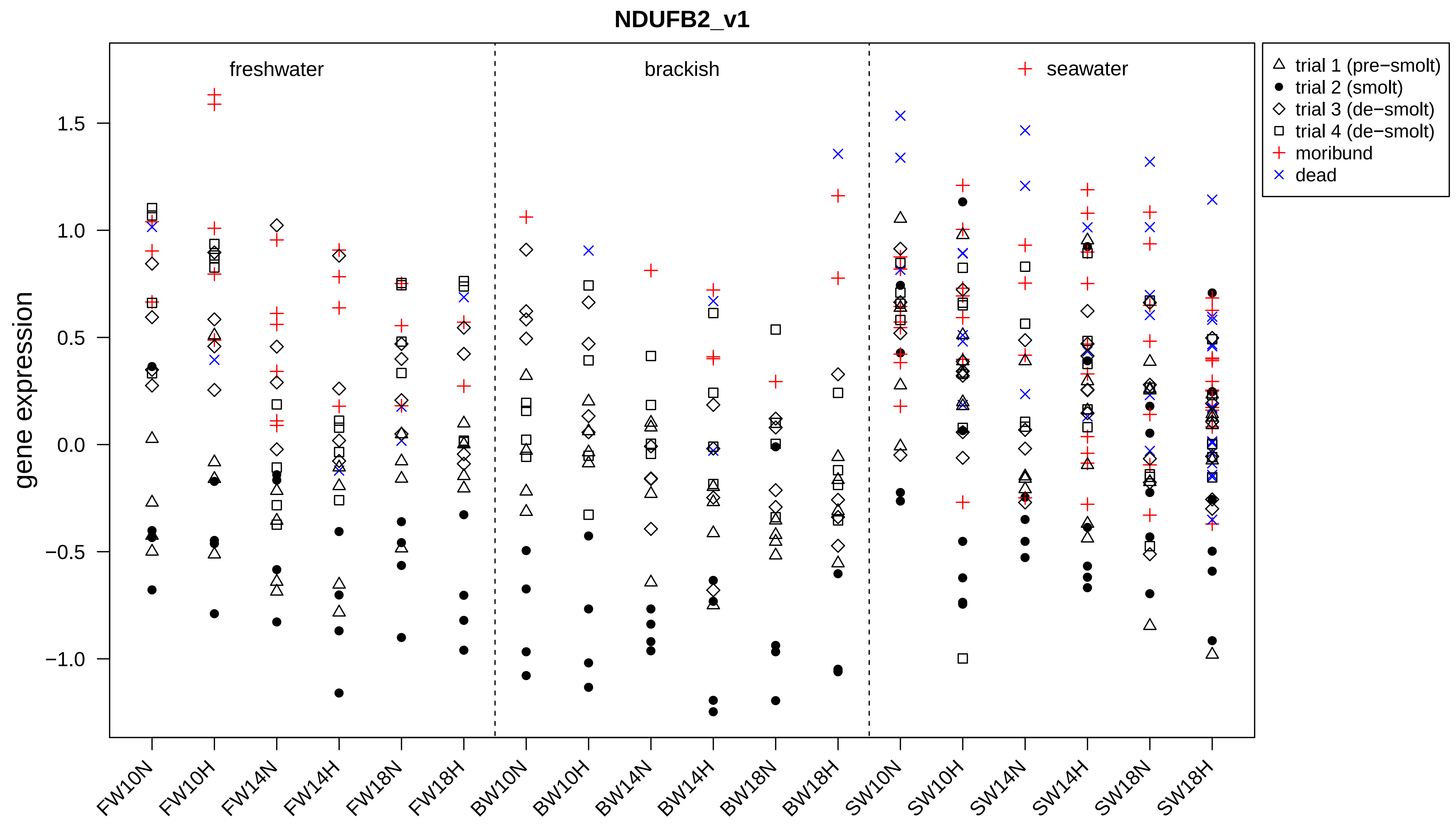

NDUFB2_v1
freshwater
brackish
seawater
gene expression
1.5
1.0
0.5
0.0
−0.5
−1.0
trial 1 (pre−smolt)
trial 2 (smolt)
trial 3 (de−smolt)
trial 4 (de−smolt)
moribund
dead
FW10N
FW10H
FW14N
FW14H
FW18N
FW18H
BW10N
BW10H
BW14N
BW14H
BW18N
BW18H
SW10N
SW10H
SW14N
SW14H
SW18N
SW18H

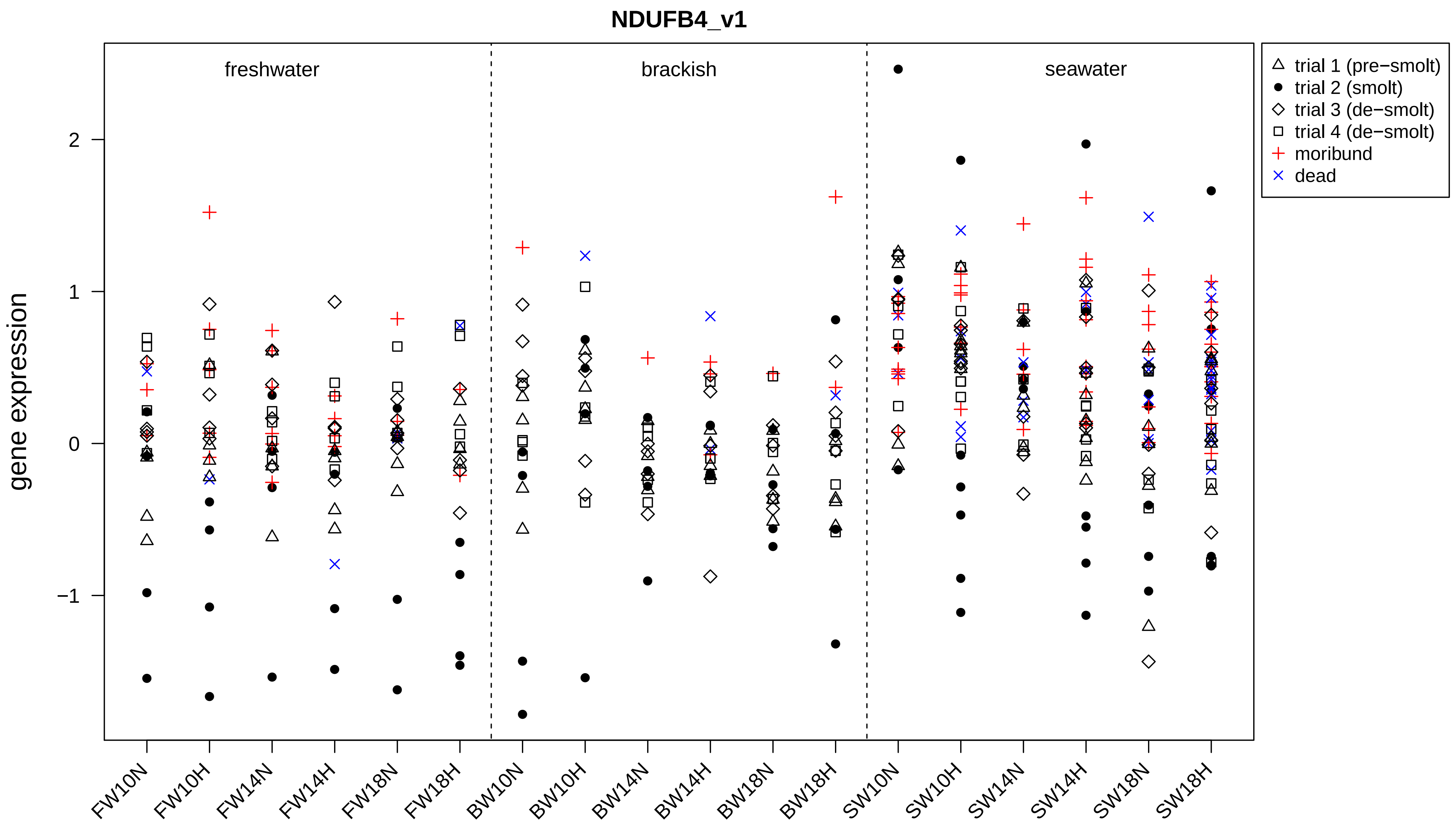
NDUFB4_v1
freshwater
brackish
seawater
gene expression
2
1
0
−1
trial 1 (pre−smolt)
trial 2 (smolt)
trial 3 (de−smolt)
trial 4 (de−smolt)
moribund
dead
FW10N
FW10H
FW14N
FW14H
FW18N
FW18H
BW10N
BW10H
BW14N
BW14H
BW18N
BW18H
SW10N
SW10H
SW14N
SW14H
SW18N
SW18H

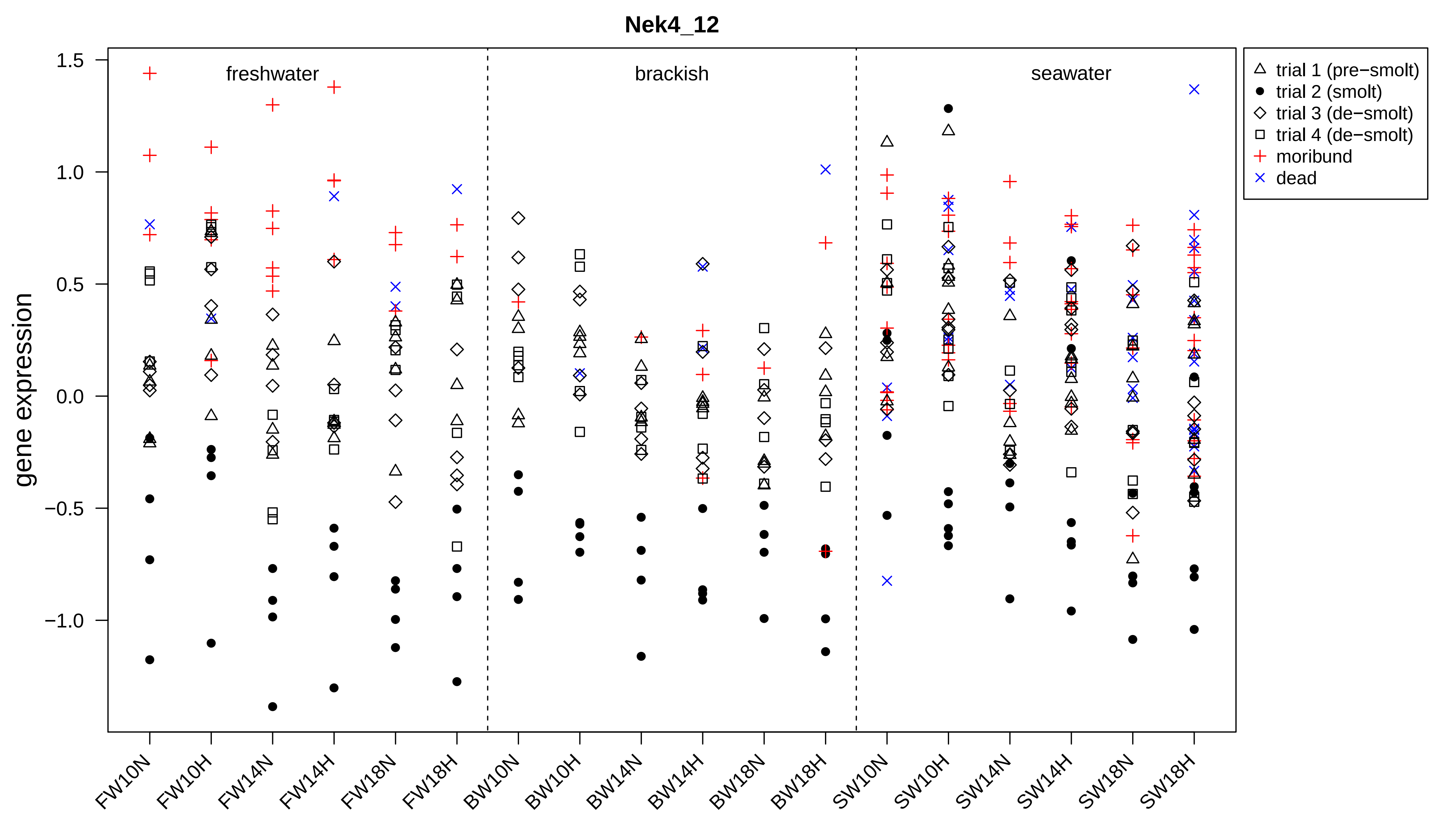

Nek4_12
freshwater
brackish
seawater
gene expression
1.5
1.0
0.5
0.0
−0.5
−1.0
trial 1 (pre−smolt)
trial 2 (smolt)
trial 3 (de−smolt)
trial 4 (de−smolt)
moribund
dead
FW10N
FW10H
FW14N
FW14H
FW18N
FW18H
BW10N
BW10H
BW14N
BW14H
BW18N
BW18H
SW10N
SW10H
SW14N
SW14H
SW18N
SW18H

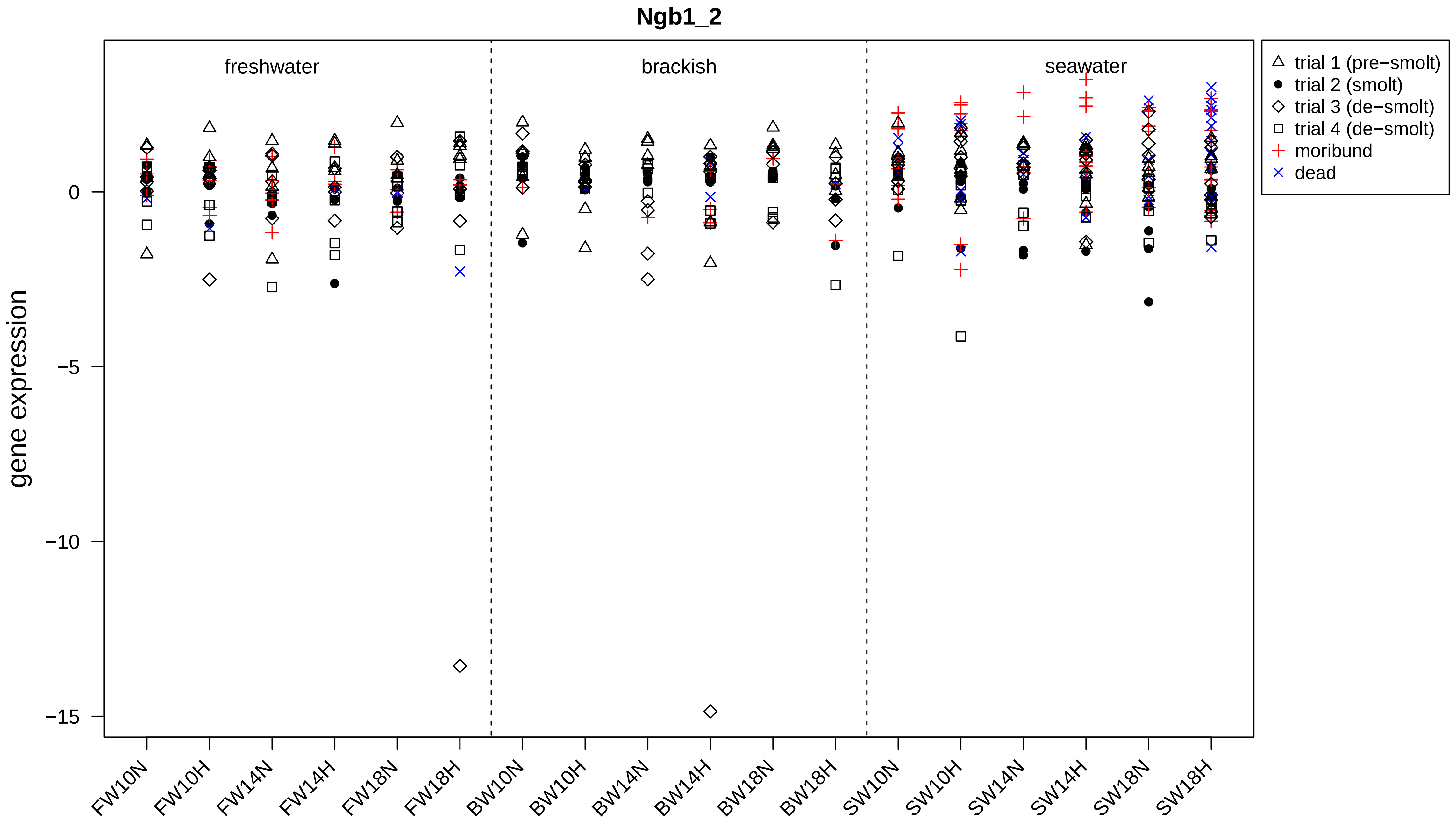
Ngb1_2
freshwater
brackish
seawater
trial 1 (pre−smolt)
trial 2 (smolt)
trial 3 (de−smolt)
trial 4 (de−smolt)
moribund
dead
gene expression
0
−5
−10
−15
FW10N
FW10H
FW14N
FW14H
FW18N
FW18H
BW10N
BW10H
BW14N
BW14H
BW18N
BW18H
SW10N
SW10H
SW14N
SW14H
SW18N
SW18H

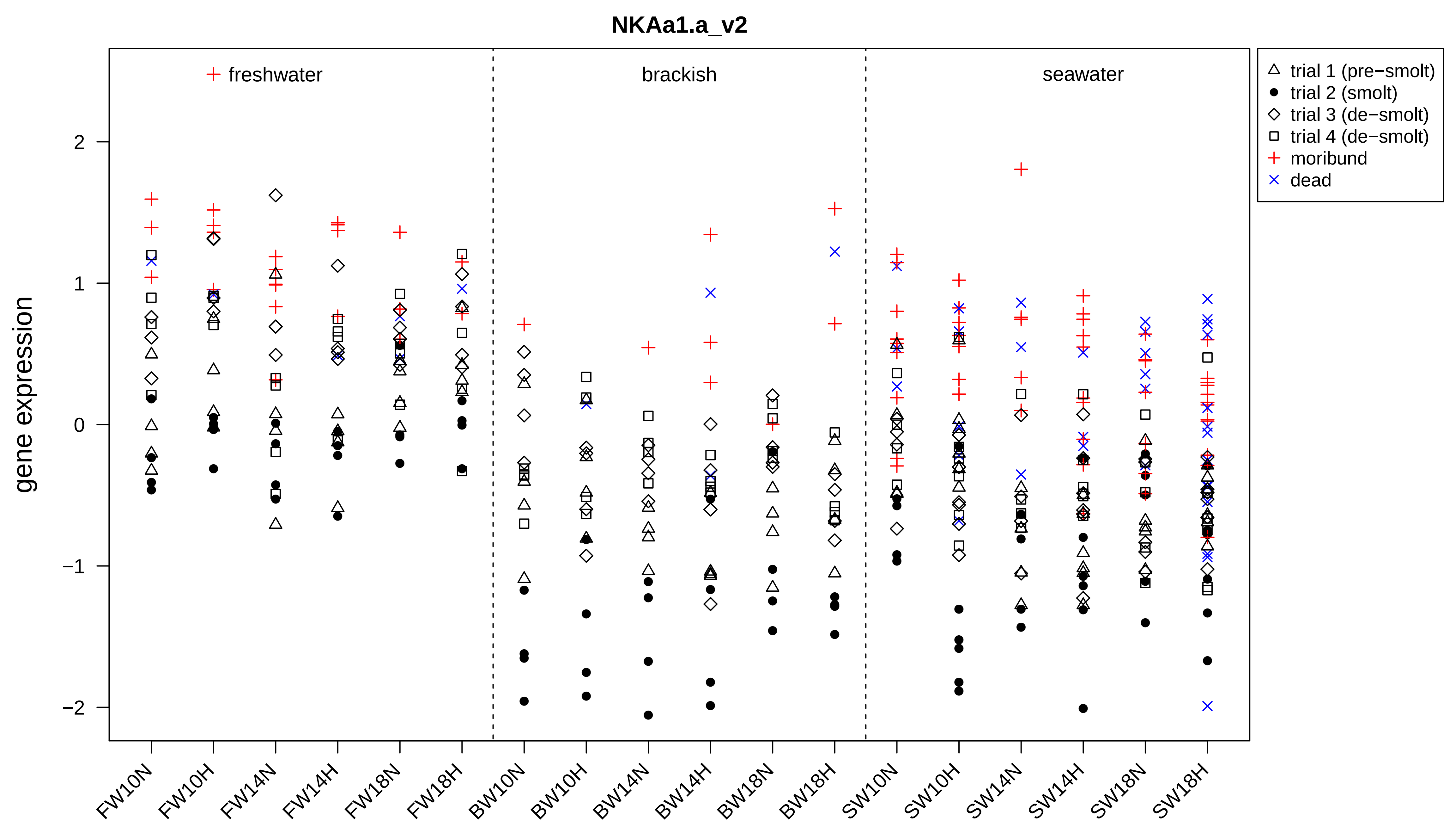
NKAa1.a_v2
gene expression
freshwater
brackish
seawater
trial 1 (pre−smolt)
trial 2 (smolt)
trial 3 (de−smolt)
trial 4 (de−smolt)
moribund
dead
2
1
0
−1
−2
FW10N
FW10H
FW14N
FW14H
FW18N
FW18H
BW10N
BW10H
BW14N
BW14H
BW18N
BW18H
SW10N
SW10H
SW14N
SW14H
SW18N
SW18H

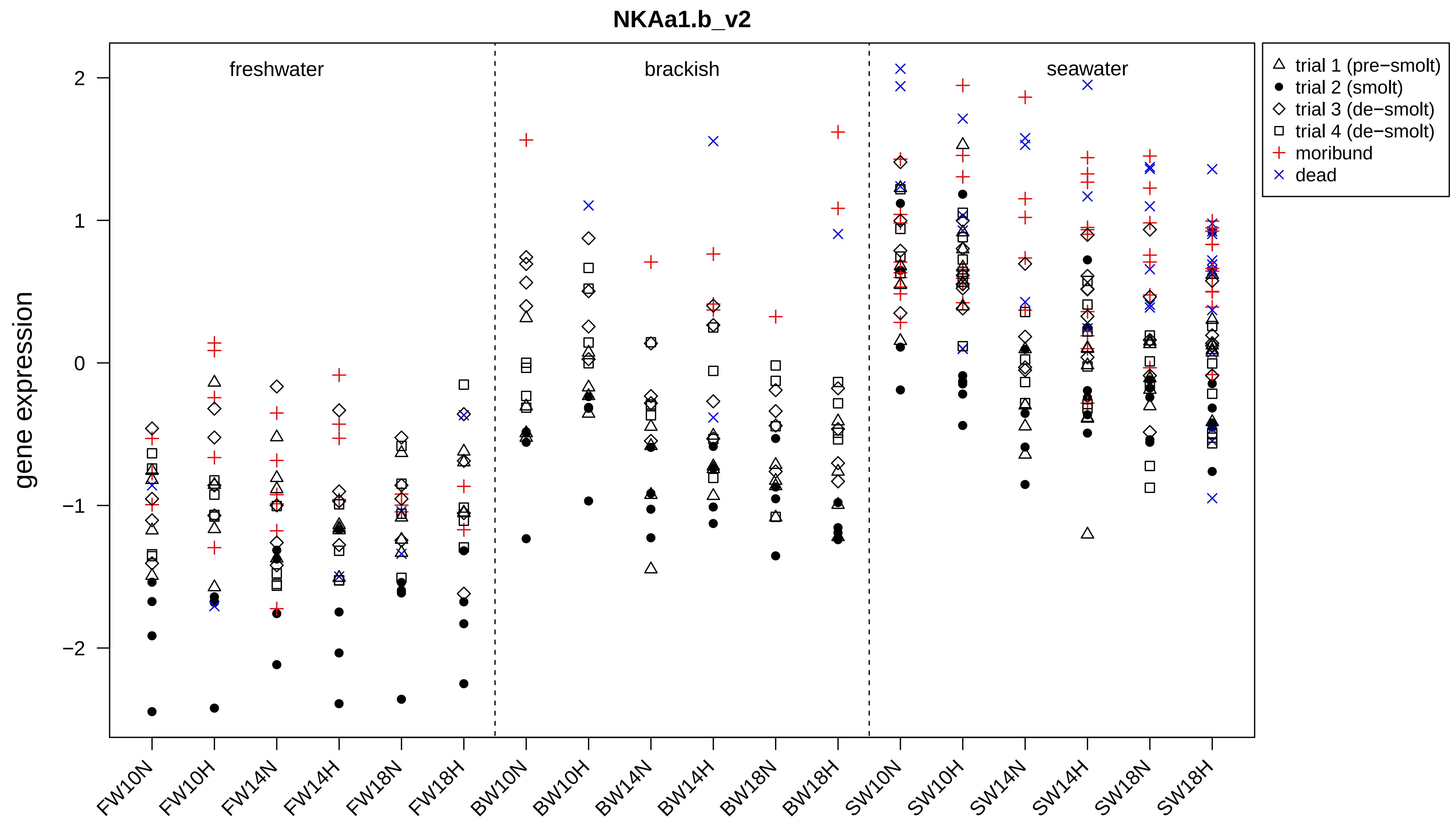

NKAa1.b_v2
gene expression
2
1
0
−1
−2
freshwater
brackish
seawater
trial 1 (pre−smolt)
trial 2 (smolt)
trial 3 (de−smolt)
trial 4 (de−smolt)
moribund
dead
FW10N
FW10H
FW14N
FW14H
FW18N
FW18H
BW10N
BW10H
BW14N
BW14H
BW18N
BW18H
SW10N
SW10H
SW14N
SW14H
SW18N
SW18H

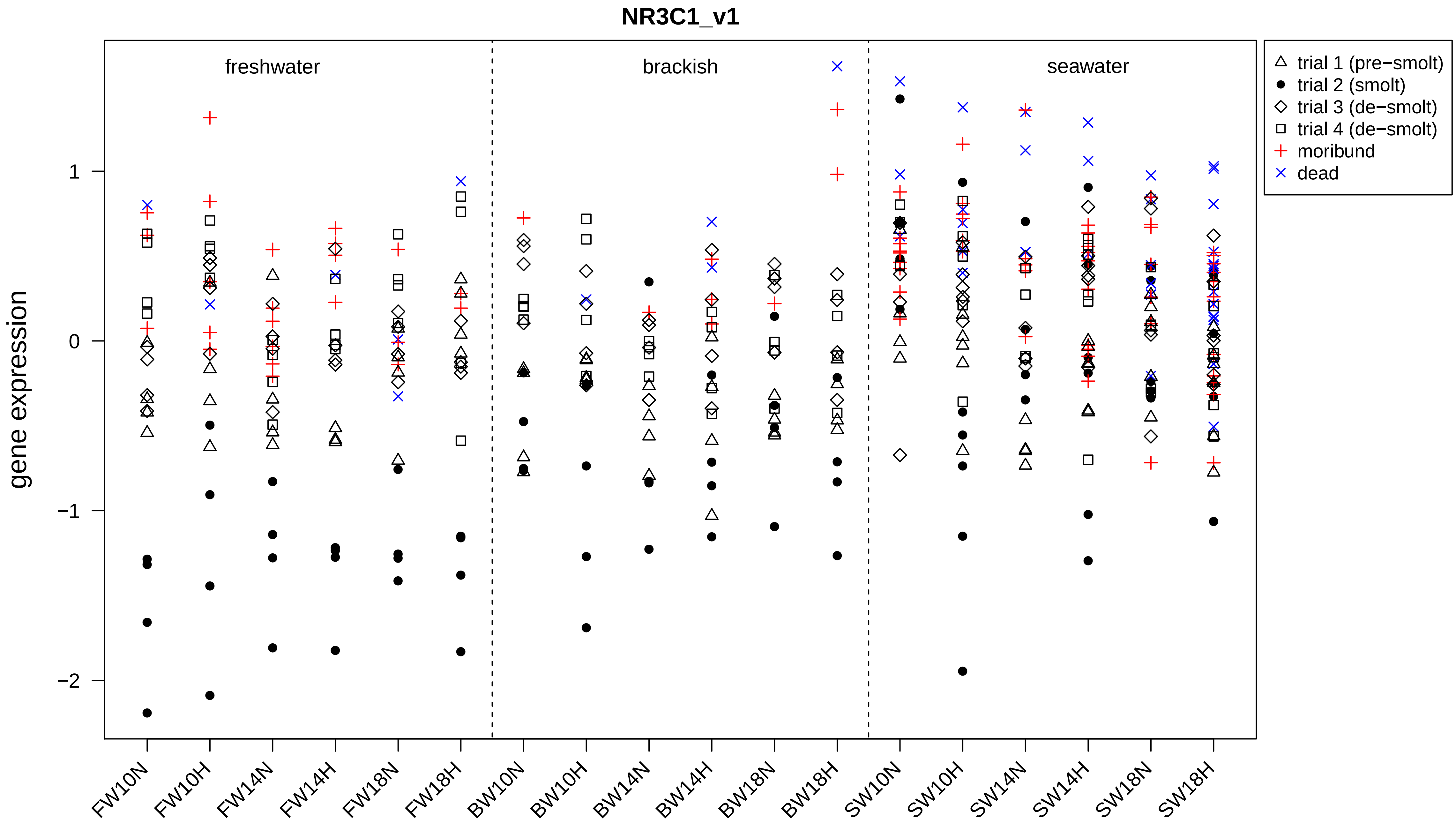
NR3C1_v1
freshwater
brackish
seawater
trial 1 (pre−smolt)
trial 2 (smolt)
trial 3 (de−smolt)
trial 4 (de−smolt)
moribund
dead
gene expression
1
0
−1
−2
FW10N
FW10H
FW14N
FW14H
FW18N
FW18H
BW10N
BW10H
BW14N
BW14H
BW18N
BW18H
SW10N
SW10H
SW14N
SW14H
SW18N
SW18H

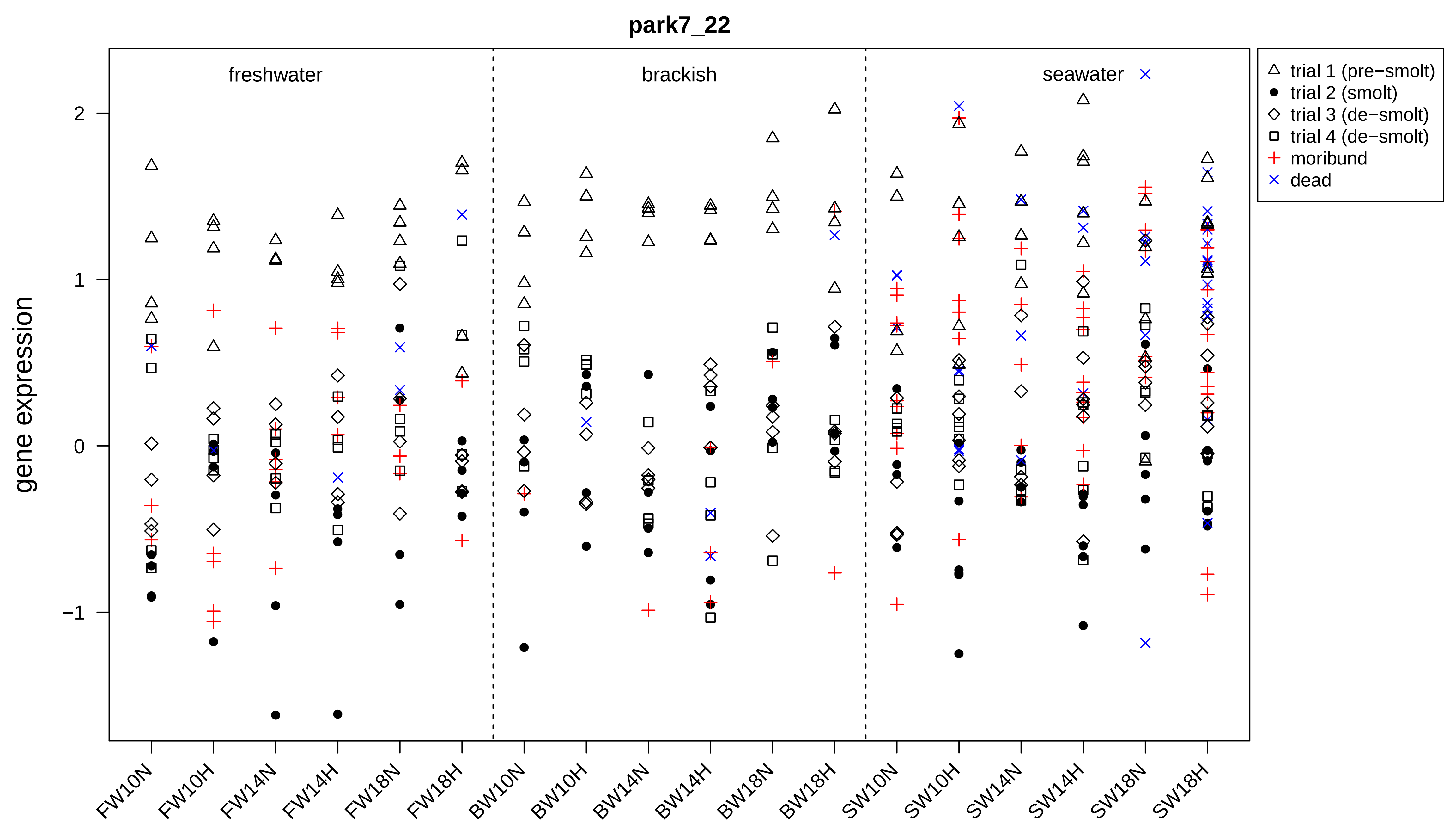
park7_22
freshwater
brackish
seawater
gene expression
2
1
0
−1
trial 1 (pre−smolt)
trial 2 (smolt)
trial 3 (de−smolt)
trial 4 (de−smolt)
moribund
dead
FW10N
FW10H
FW14N
FW14H
FW18N
FW18H
BW10N
BW10H
BW14N
BW14H
BW18N
BW18H
SW10N
SW10H
SW14N
SW14H
SW18N
SW18H

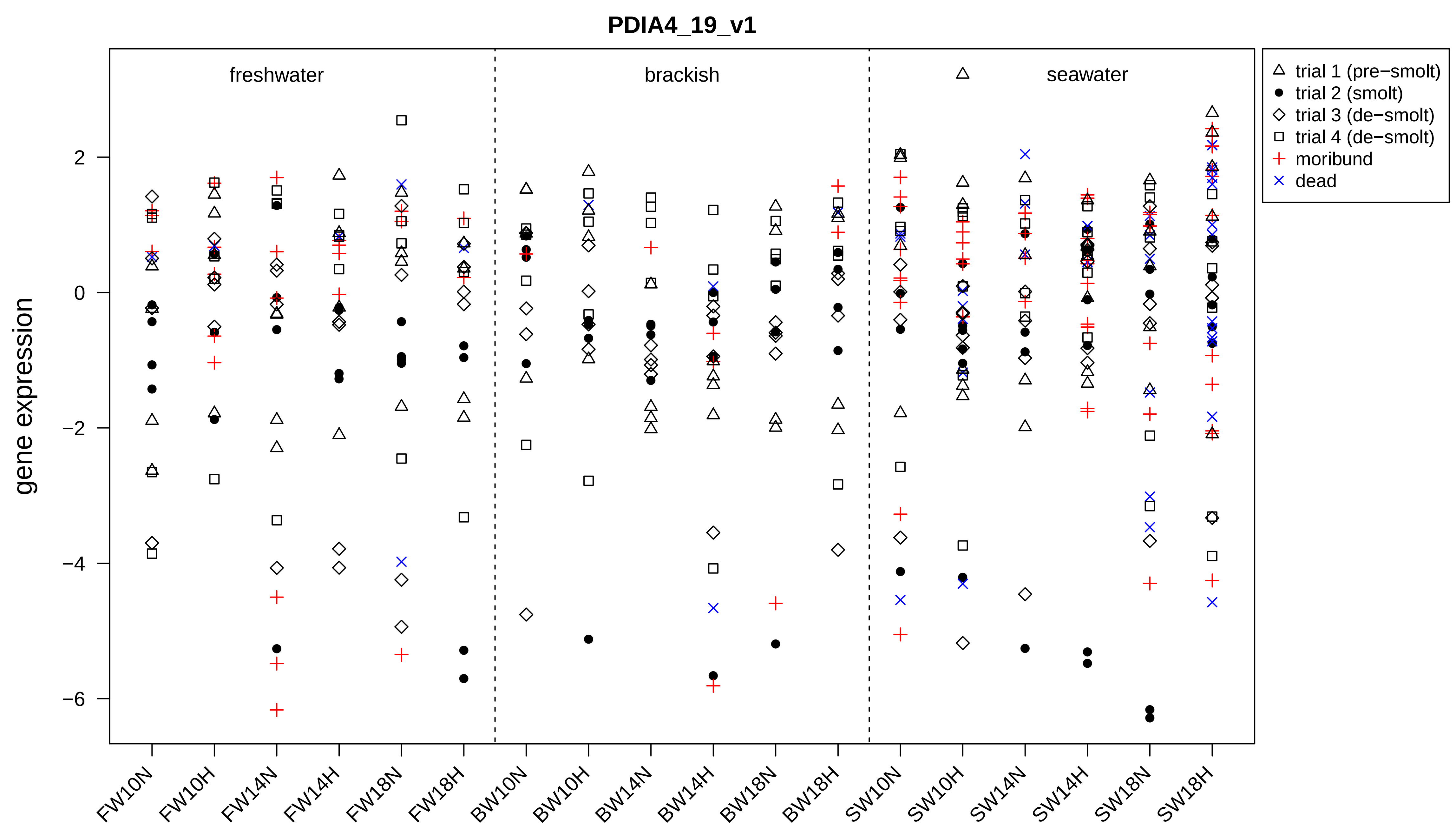

PDIA4_19_v1
freshwater
brackish
seawater
trial 1 (pre−smolt)
trial 2 (smolt)
trial 3 (de−smolt)
trial 4 (de−smolt)
moribund
dead
gene expression
2
0
−2
−4
−6
FW10N
FW10H
FW14N
FW14H
FW18N
FW18H
BW10N
BW10H
BW14N
BW14H
BW18N
BW18H
SW10N
SW10H
SW14N
SW14H
SW18N
SW18H

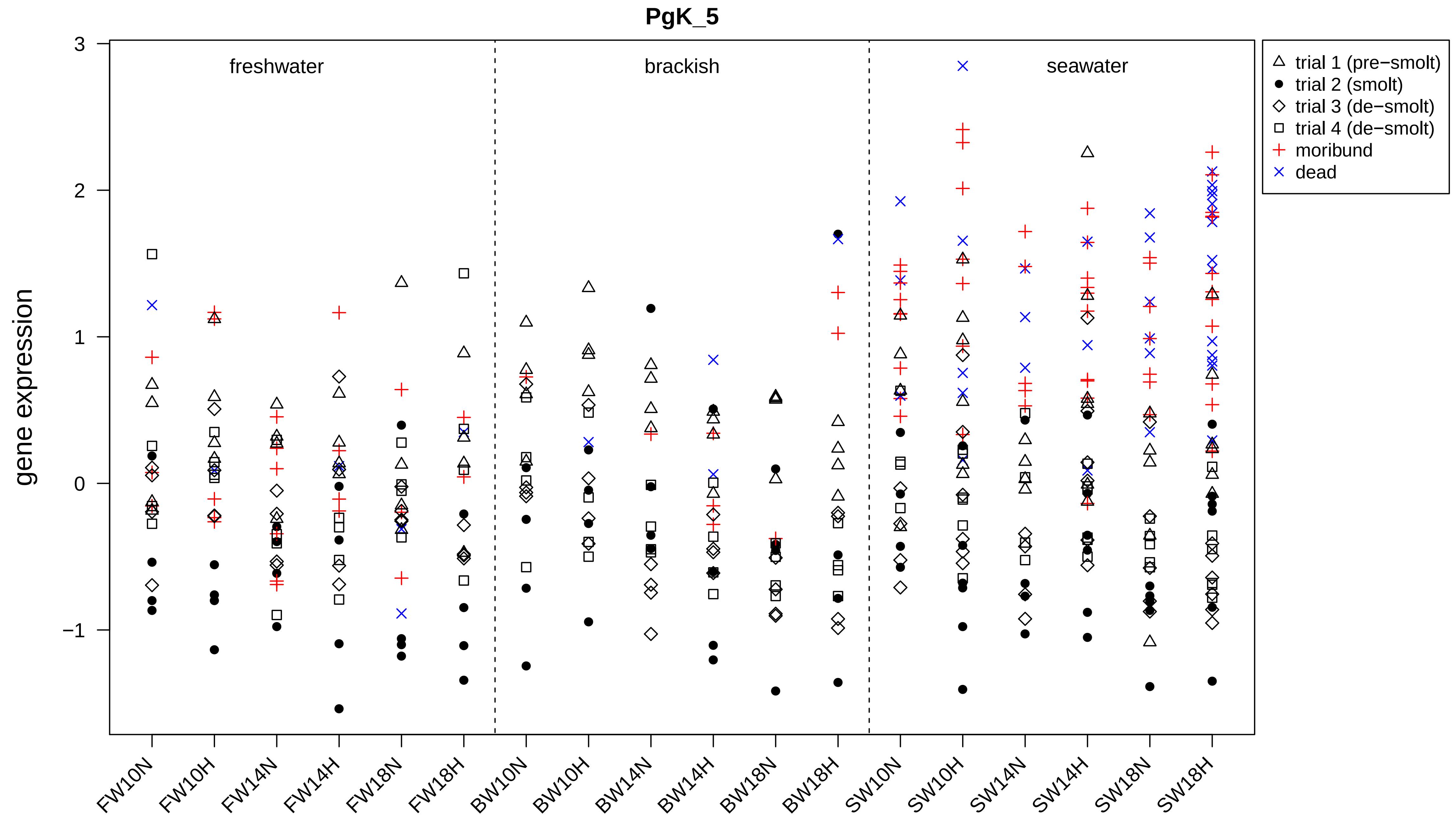
PgK_5
gene expression
3
2
1
0
−1
freshwater
brackish
seawater
trial 1 (pre−smolt)
trial 2 (smolt)
trial 3 (de−smolt)
trial 4 (de−smolt)
moribund
dead
FW10N
FW10H
FW14N
FW14H
FW18N
FW18H
BW10N
BW10H
BW14N
BW14H
BW18N
BW18H
SW10N
SW10H
SW14N
SW14H
SW18N
SW18H

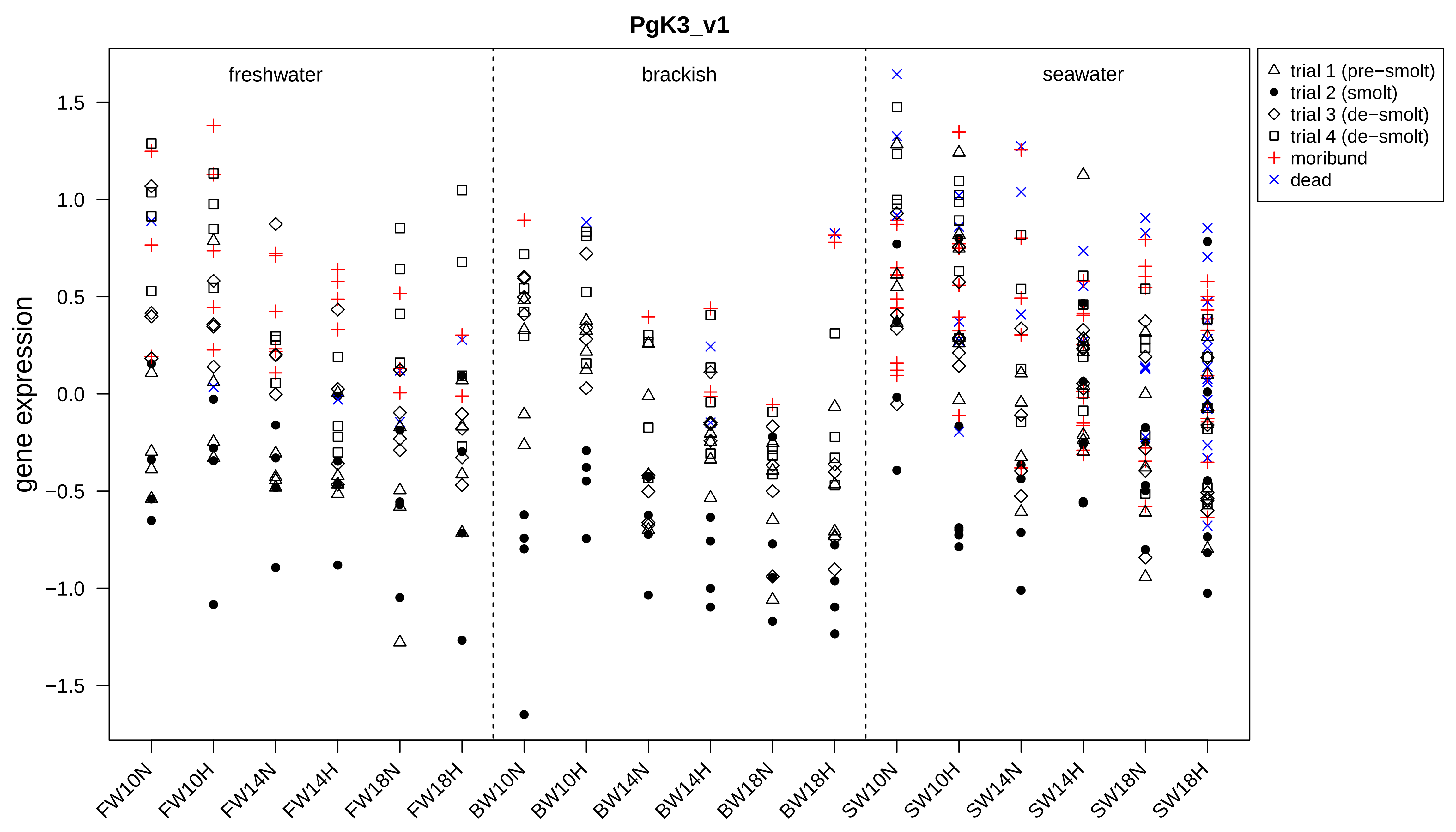
PgK3_v1
freshwater
brackish
seawater
gene expression
1.5
1.0
0.5
0.0
−0.5
−1.0
−1.5
trial 1 (pre−smolt)
trial 2 (smolt)
trial 3 (de−smolt)
trial 4 (de−smolt)
moribund
dead
FW10N
FW10H
FW14N
FW14H
FW18N
FW18H
BW10N
BW10H
BW14N
BW14H
BW18N
BW18H
SW10N
SW10H
SW14N
SW14H
SW18N
SW18H

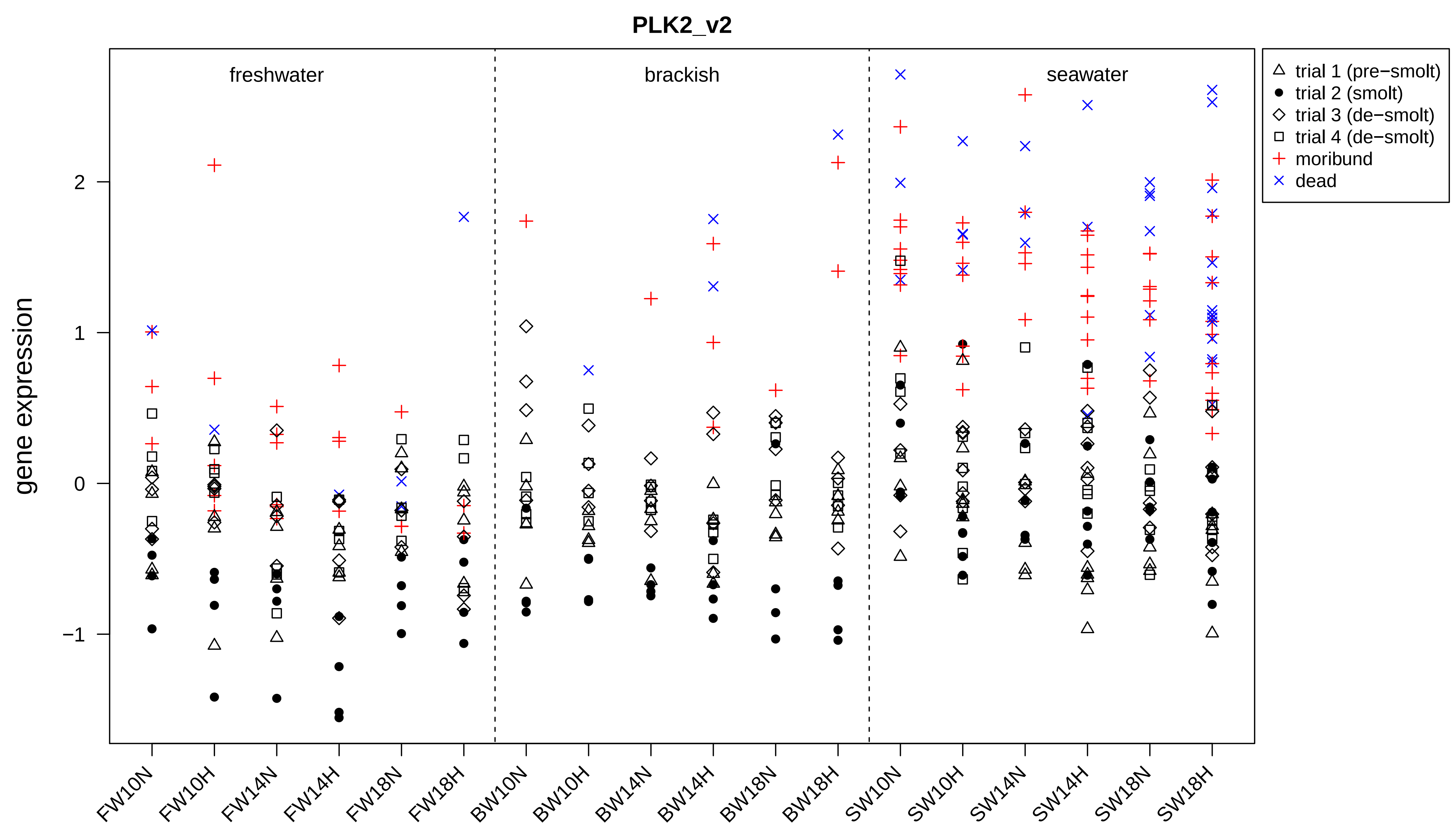
PLK2_v2
freshwater
brackish
seawater
gene expression
2
1
0
−1
FW10N
FW10H
FW14N
FW14H
FW18N
FW18H
BW10N
BW10H
BW14N
BW14H
BW18N
BW18H
SW10N
SW10H
SW14N
SW14H
SW18N
SW18H
trial 1 (pre−smolt)
trial 2 (smolt)
trial 3 (de−smolt)
trial 4 (de−smolt)
moribund
dead

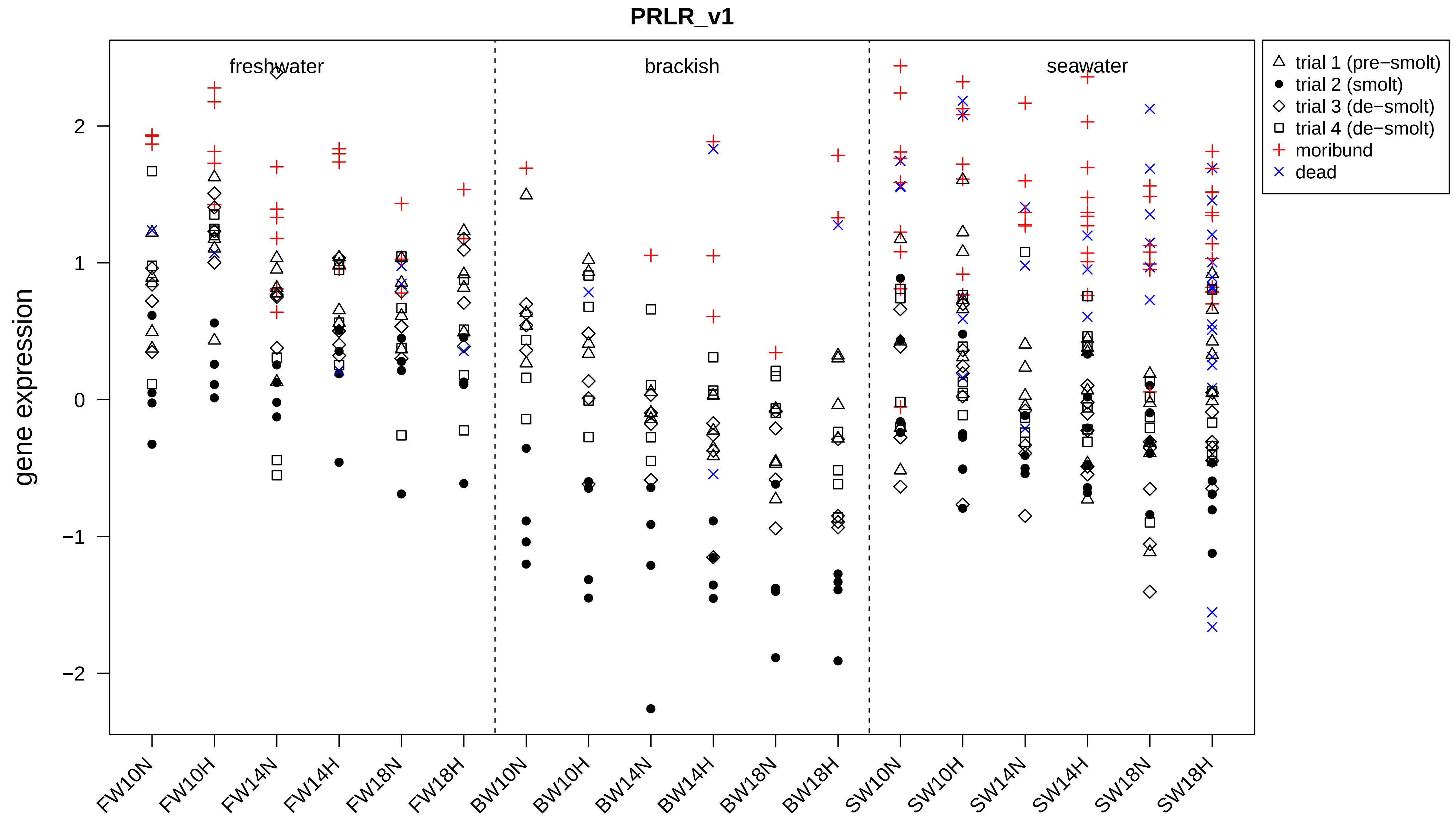
PRLR_v1
freshwater
brackish
seawater
trial 1 (pre−smolt)
trial 2 (smolt)
trial 3 (de−smolt)
trial 4 (de−smolt)
moribund
dead
gene expression
2
1
0
−1
−2
FW10N
FW10H
FW14N
FW14H
FW18N
FW18H
BW10N
BW10H
BW14N
BW14H
BW18N
BW18H
SW10N
SW10H
SW14N
SW14H
SW18N
SW18H

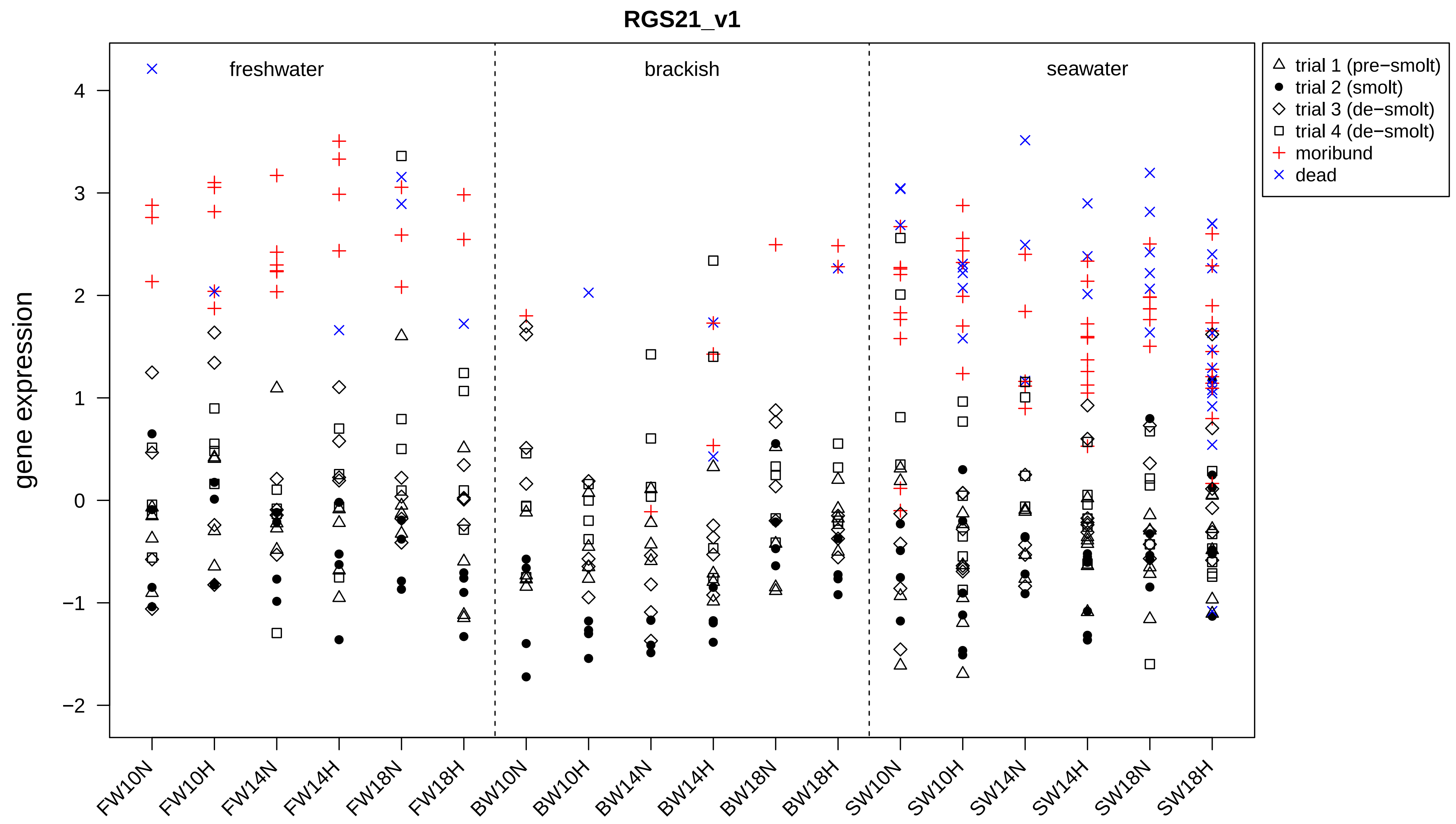
RGS21_v1
freshwater
brackish
seawater
gene expression
4
3
2
1
0
−1
−2
trial 1 (pre−smolt)
trial 2 (smolt)
trial 3 (de−smolt)
trial 4 (de−smolt)
moribund
dead
FW10N
FW10H
FW14N
FW14H
FW18N
FW18H
BW10N
BW10H
BW14N
BW14H
BW18N
BW18H
SW10N
SW10H
SW14N
SW14H
SW18N
SW18H

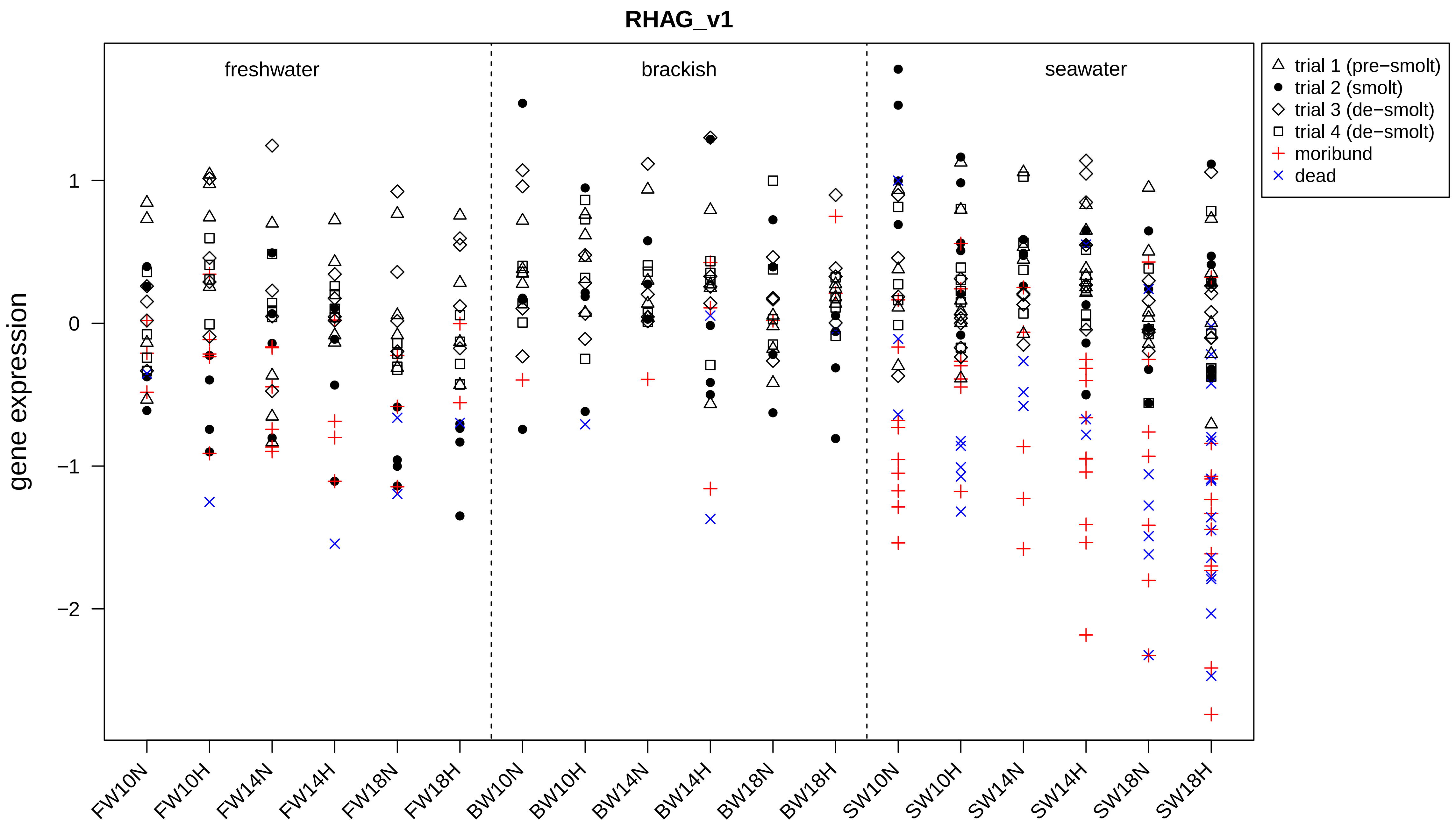

RHAG_v1
freshwater
brackish
seawater
gene expression
1
0
−1
−2
FW10N
FW10H
FW14N
FW14H
FW18N
FW18H
BW10N
BW10H
BW14N
BW14H
BW18N
BW18H
SW10N
SW10H
SW14N
SW14H
SW18N
SW18H
trial 1 (pre−smolt)
trial 2 (smolt)
trial 3 (de−smolt)
trial 4 (de−smolt)
moribund
dead

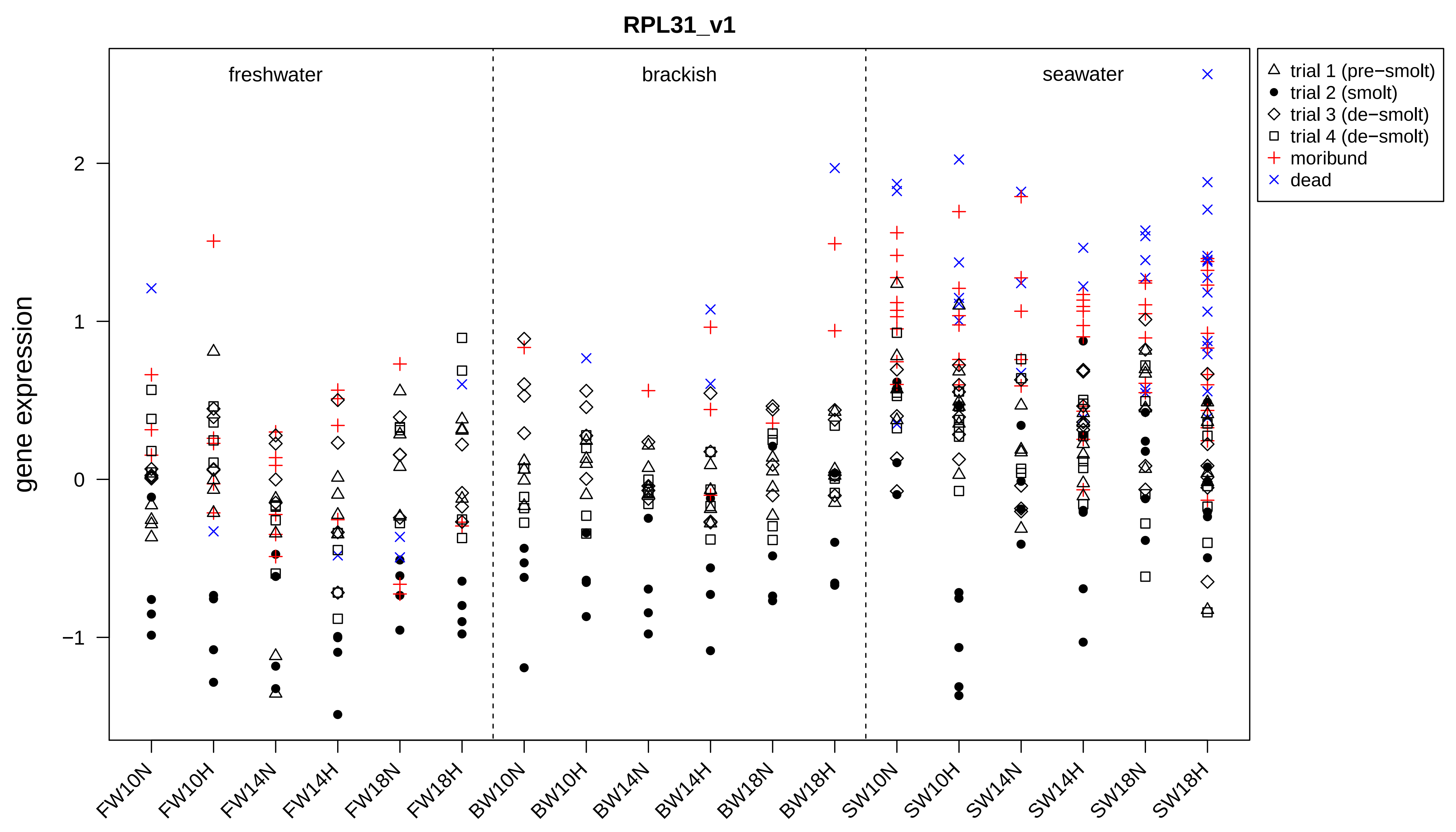
RPL31_v1
gene expression
freshwater
brackish
seawater
2
1
0
−1
trial 1 (pre−smolt)
trial 2 (smolt)
trial 3 (de−smolt)
trial 4 (de−smolt)
moribund
dead
FW10N
FW10H
FW14N
FW14H
FW18N
FW18H
BW10N
BW10H
BW14N
BW14H
BW18N
BW18H
SW10N
SW10H
SW14N
SW14H
SW18N
SW18H

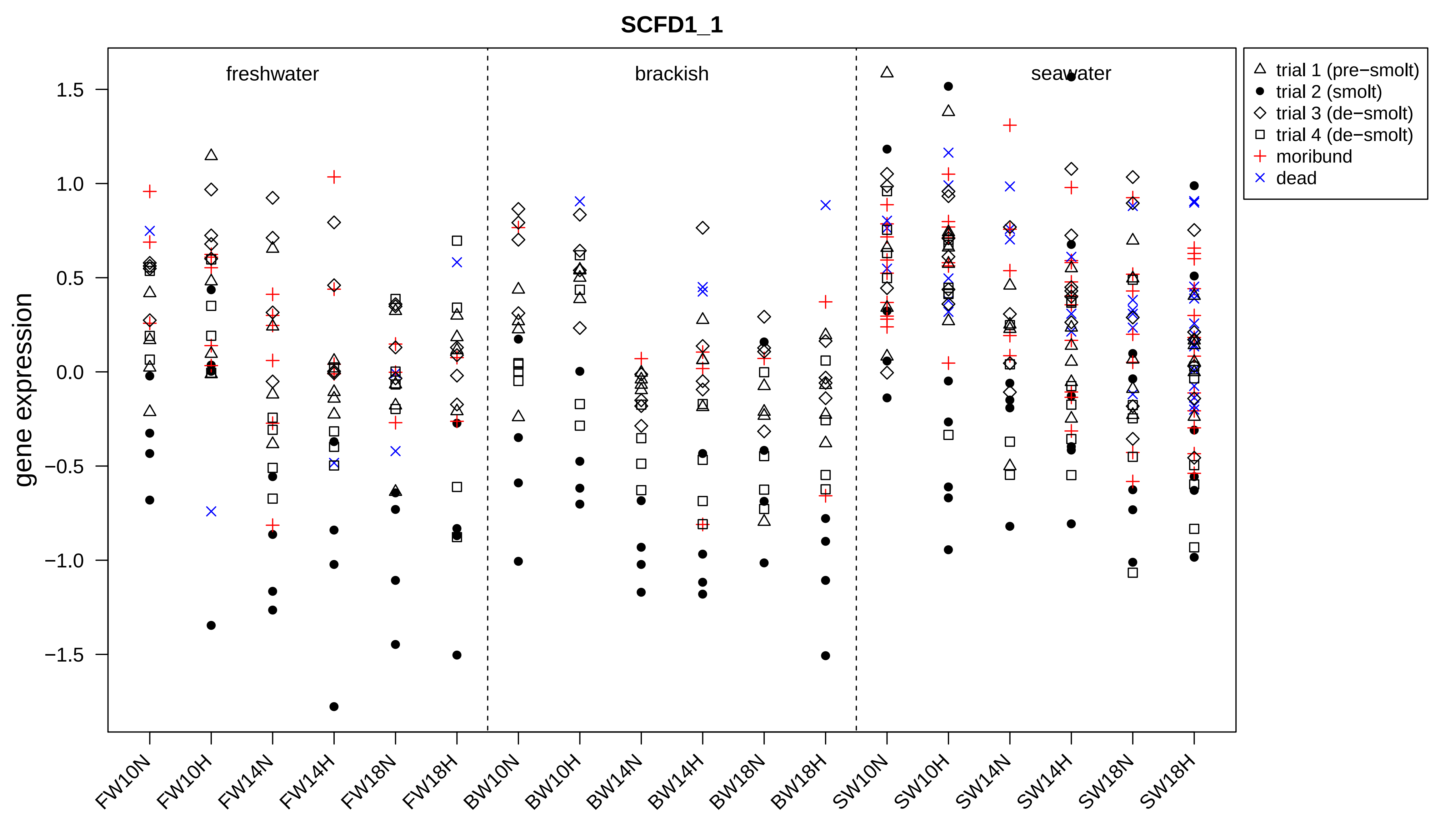

SCFD1_1
gene expression
freshwater
brackish
seawater
1.5
1.0
0.5
0.0
−0.5
−1.0
−1.5
trial 1 (pre−smolt)
trial 2 (smolt)
trial 3 (de−smolt)
trial 4 (de−smolt)
moribund
dead
FW10N
FW10H
FW14N
FW14H
FW18N
FW18H
BW10N
BW10H
BW14N
BW14H
BW18N
BW18H
SW10N
SW10H
SW14N
SW14H
SW18N
SW18H

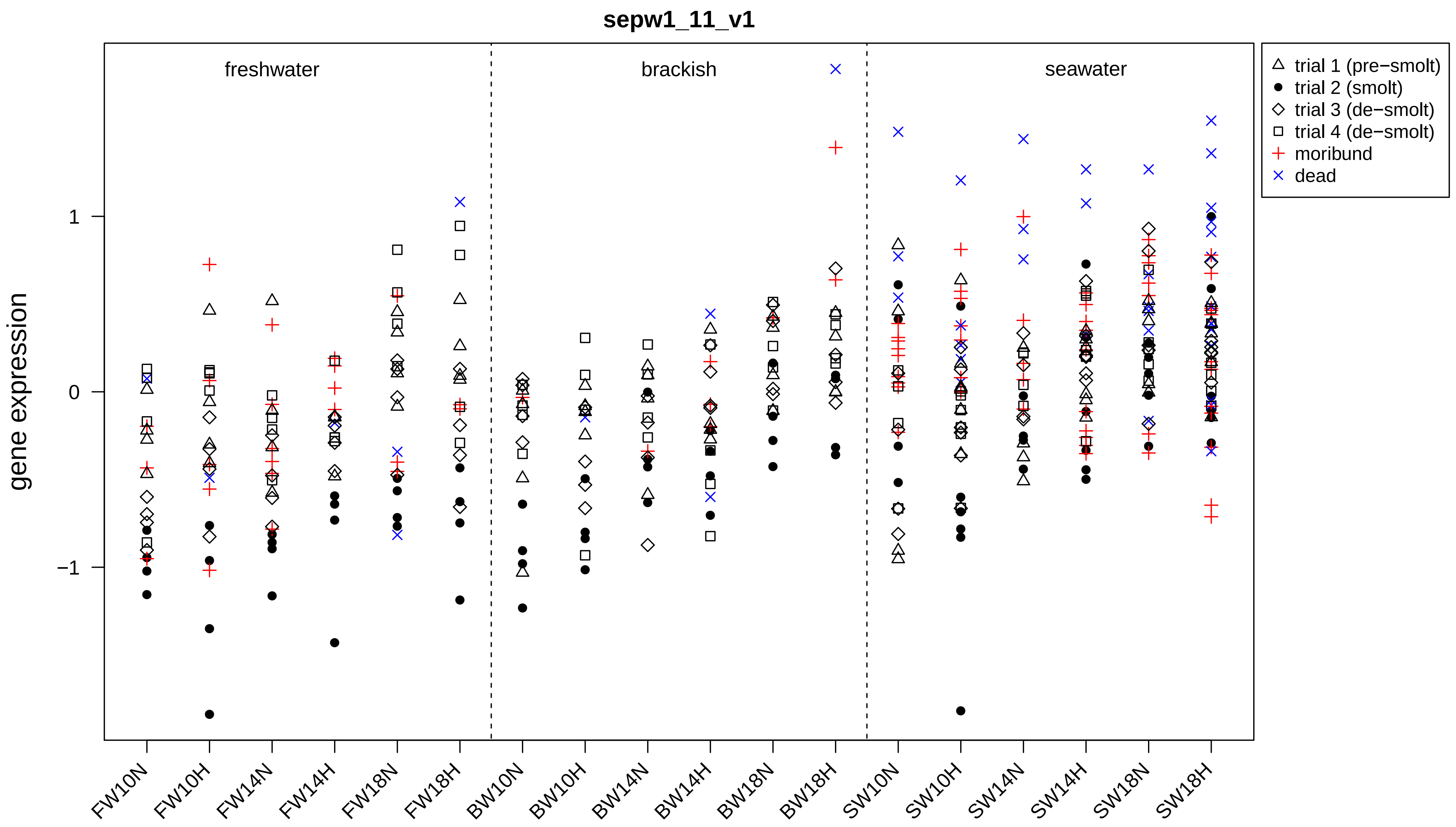
sepw1_11_v1
gene expression
freshwater
brackish
seawater
1
0
−1
FW10N
FW10H
FW14N
FW14H
FW18N
FW18H
BW10N
BW10H
BW14N
BW14H
BW18N
BW18H
SW10N
SW10H
SW14N
SW14H
SW18N
SW18H
trial 1 (pre−smolt)
trial 2 (smolt)
trial 3 (de−smolt)
trial 4 (de−smolt)
moribund
dead

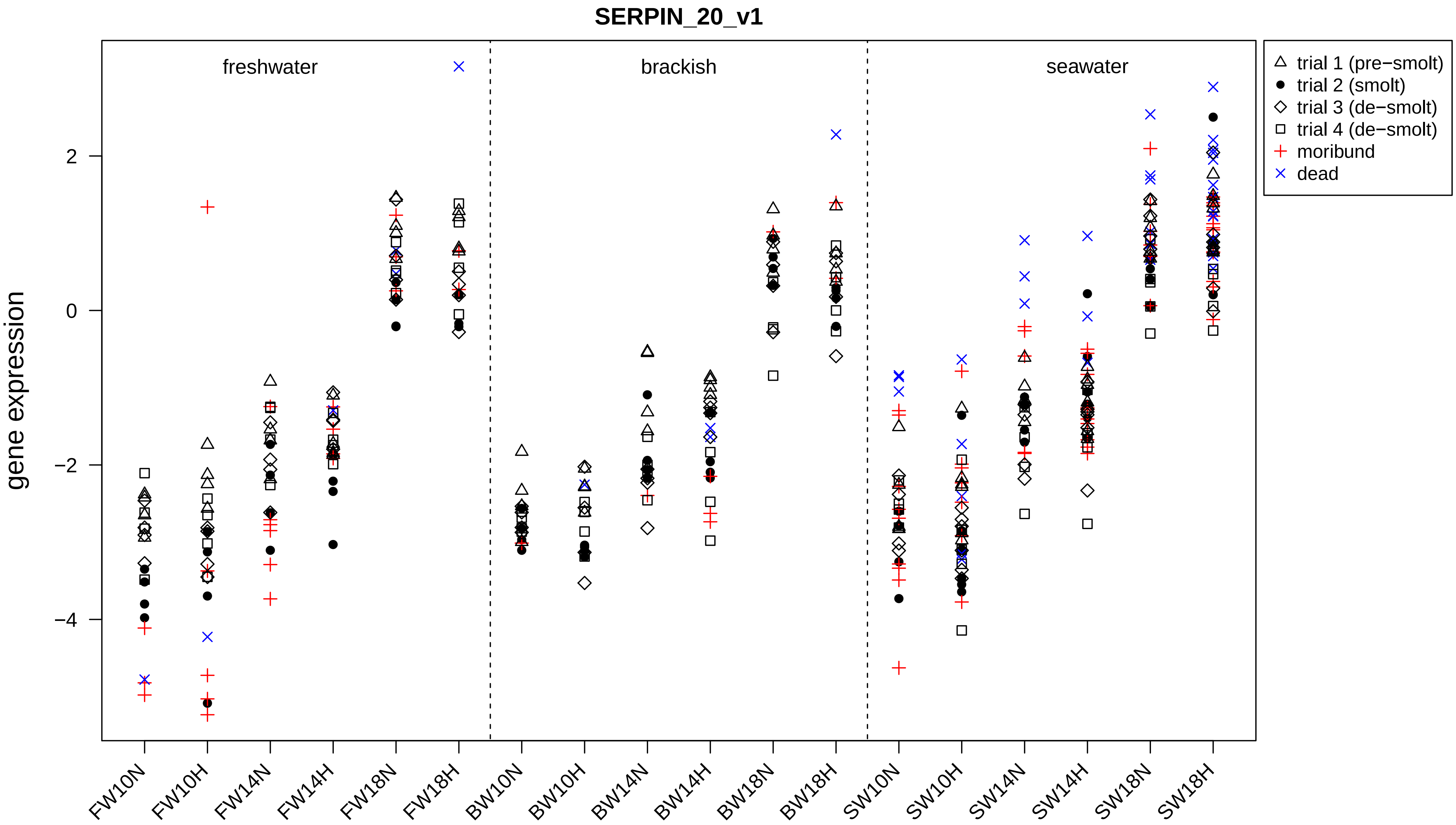

SERPIN_20_v1
freshwater
brackish
seawater
trial 1 (pre−smolt)
trial 2 (smolt)
trial 3 (de−smolt)
trial 4 (de−smolt)
moribund
dead
gene expression
2
0
−2
−4
FW10N
FW10H
FW14N
FW14H
FW18N
FW18H
BW10N
BW10H
BW14N
BW14H
BW18N
BW18H
SW10N
SW10H
SW14N
SW14H
SW18N
SW18H

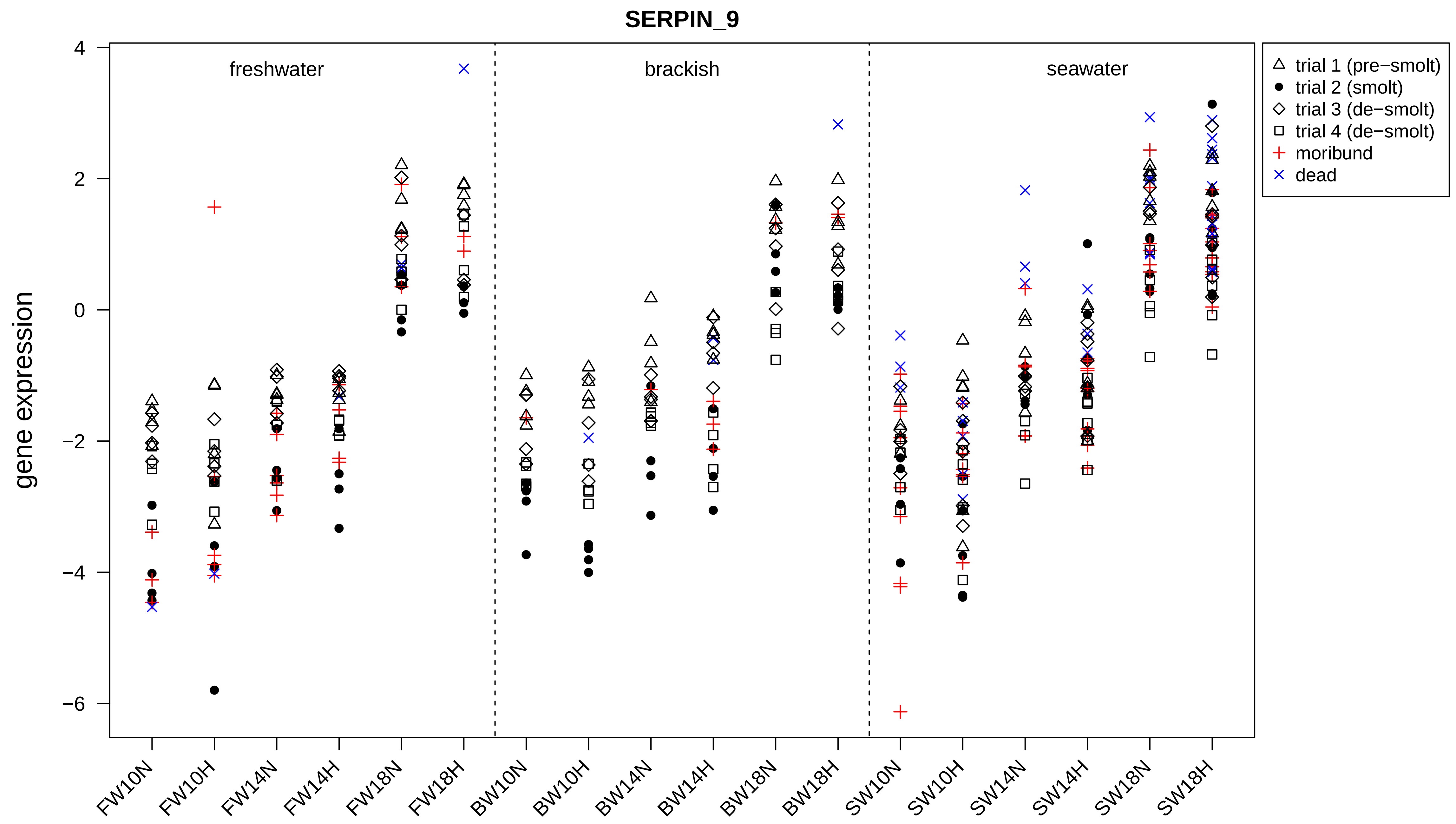
SERPIN_9
gene expression
freshwater
brackish
seawater
trial 1 (pre−smolt)
trial 2 (smolt)
trial 3 (de−smolt)
trial 4 (de−smolt)
moribund
dead
4
2
0
−2
−4
−6
FW10N
FW10H
FW14N
FW14H
FW18N
FW18H
BW10N
BW10H
BW14N
BW14H
BW18N
BW18H
SW10N
SW10H
SW14N
SW14H
SW18N
SW18H

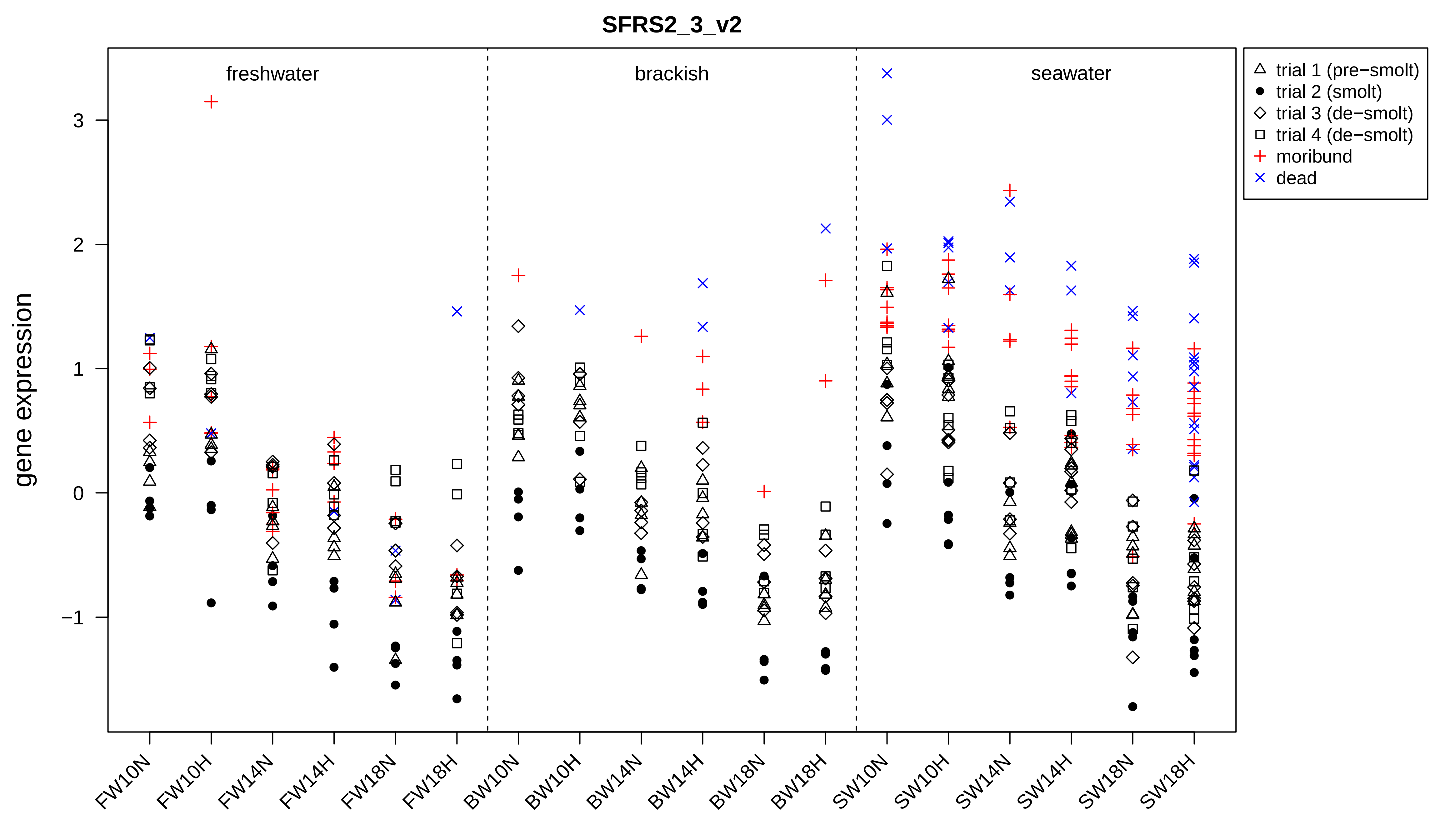
SFRS2_3_v2
gene expression
freshwater
brackish
seawater
trial 1 (pre−smolt)
trial 2 (smolt)
trial 3 (de−smolt)
trial 4 (de−smolt)
moribund
dead
3
2
1
0
−1
FW10N
FW10H
FW14N
FW14H
FW18N
FW18H
BW10N
BW10H
BW14N
BW14H
BW18N
BW18H
SW10N
SW10H
SW14N
SW14H
SW18N
SW18H

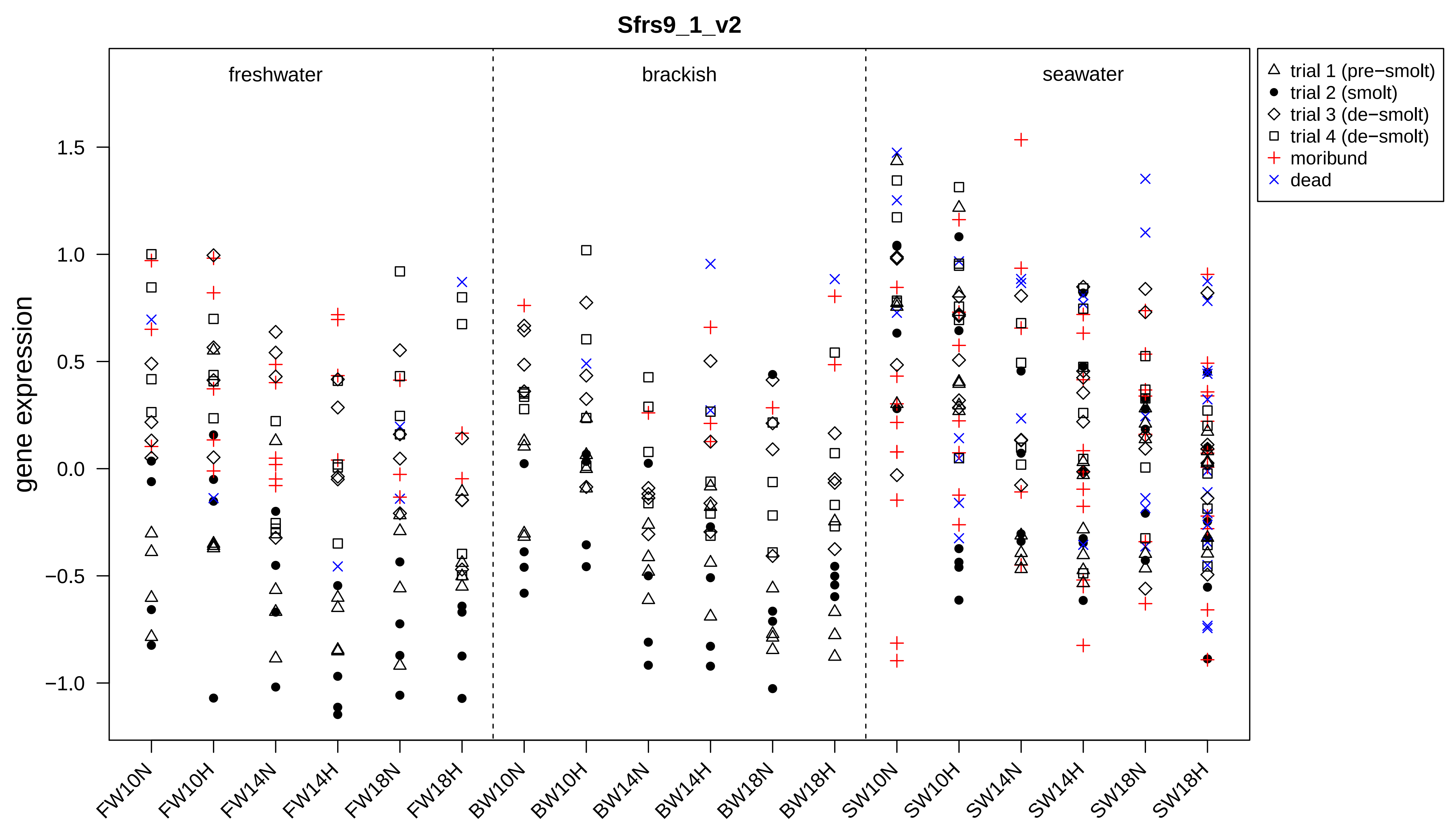

Sfrs9_1_v2
freshwater
brackish
seawater
gene expression
1.5
1.0
0.5
0.0
−0.5
−1.0
FW10N
FW10H
FW14N
FW14H
FW18N
FW18H
BW10N
BW10H
BW14N
BW14H
BW18N
BW18H
SW10N
SW10H
SW14N
SW14H
SW18N
SW18H
trial 1 (pre−smolt)
trial 2 (smolt)
trial 3 (de−smolt)
trial 4 (de−smolt)
moribund
dead

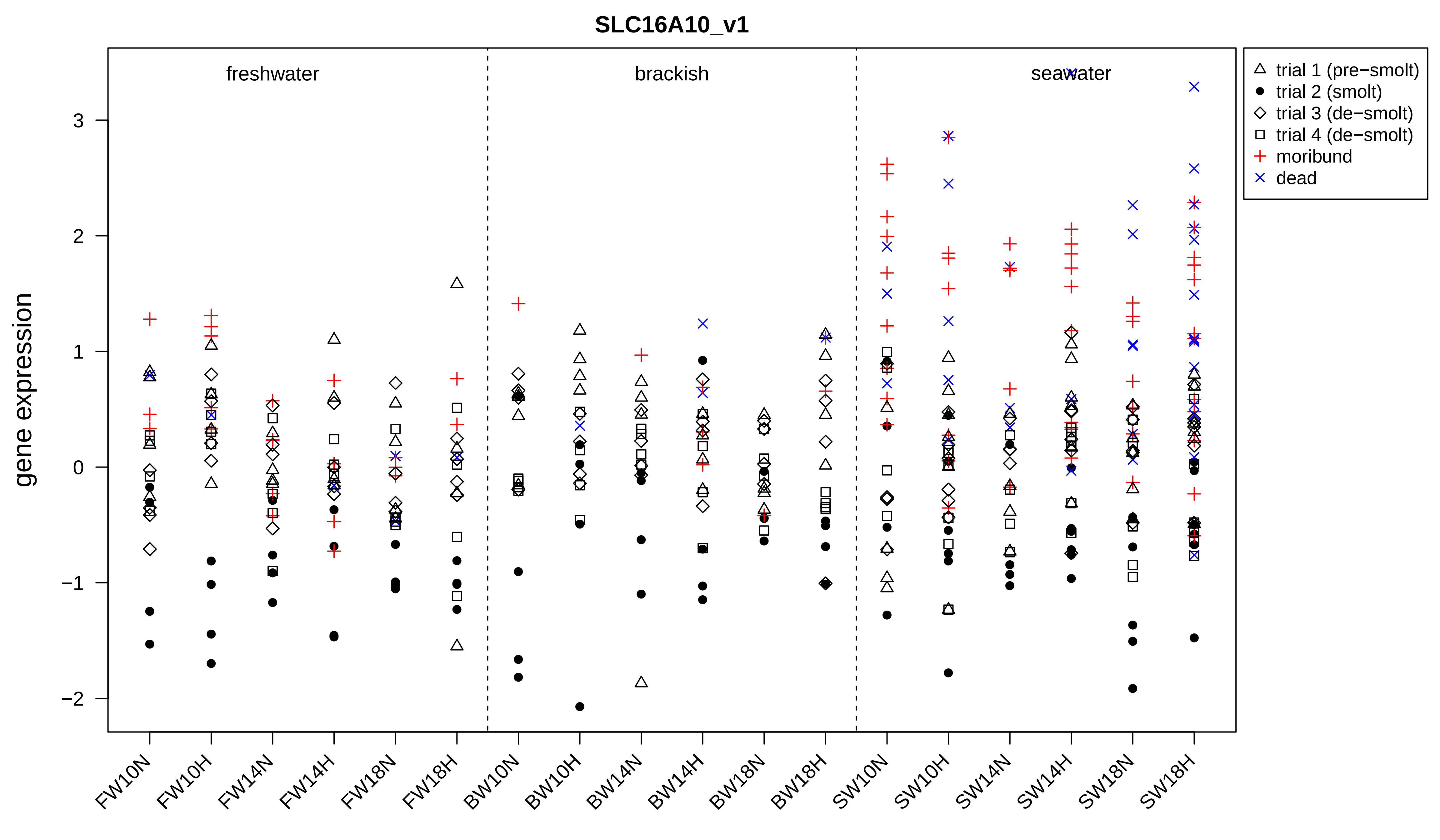

SLC16A10_v1
freshwater
brackish
seawater
gene expression
3
2
1
0
−1
−2
trial 1 (pre−smolt)
trial 2 (smolt)
trial 3 (de−smolt)
trial 4 (de−smolt)
moribund
dead
FW10N
FW10H
FW14N
FW14H
FW18N
FW18H
BW10N
BW10H
BW14N
BW14H
BW18N
BW18H
SW10N
SW10H
SW14N
SW14H
SW18N
SW18H

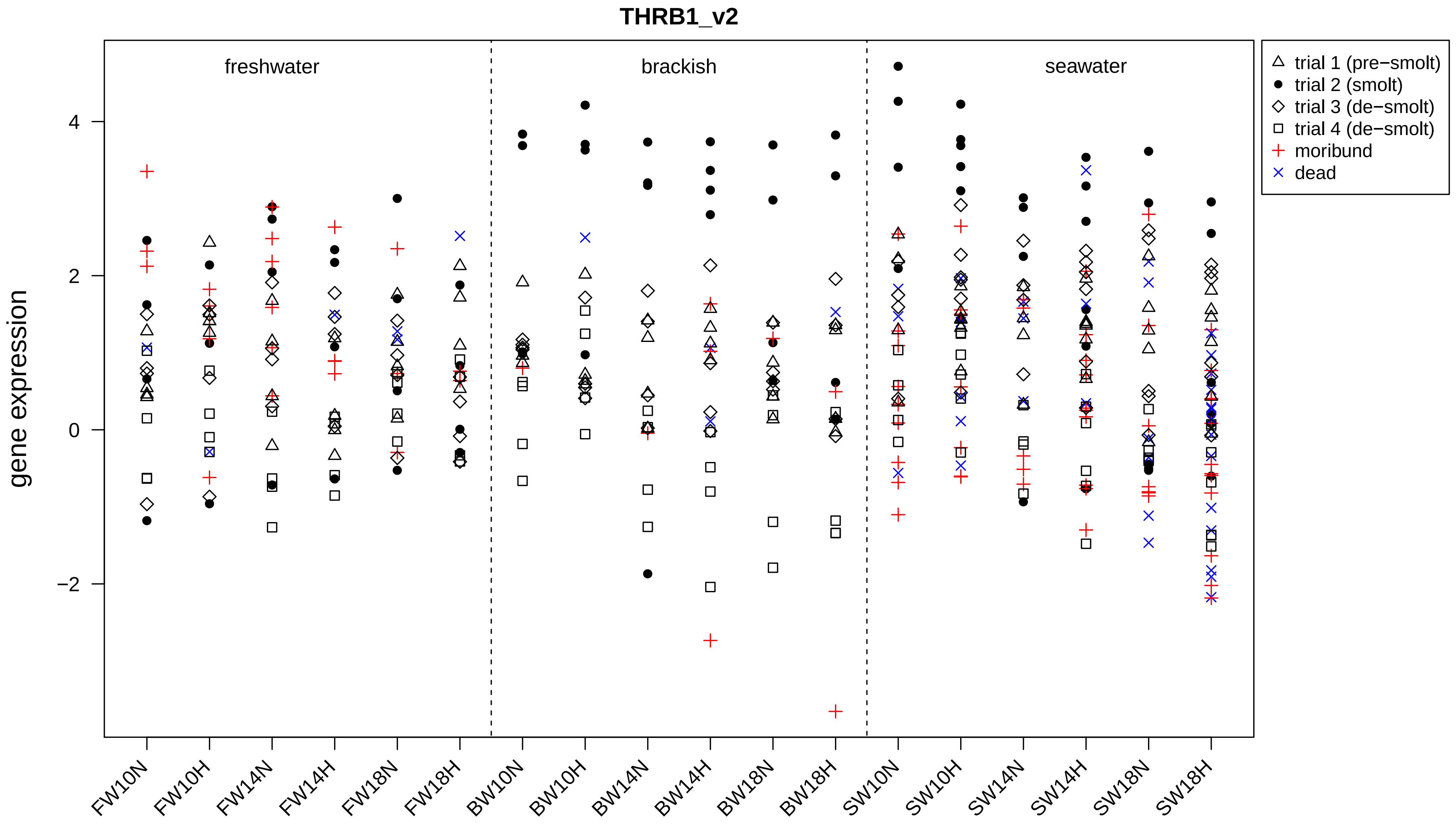
THRB1_v2
gene expression
freshwater
brackish
seawater
4
2
0
−2
trial 1 (pre−smolt)
trial 2 (smolt)
trial 3 (de−smolt)
trial 4 (de−smolt)
moribund
dead
FW10N
FW10H
FW14N
FW14H
FW18N
FW18H
BW10N
BW10H
BW14N
BW14H
BW18N
BW18H
SW10N
SW10H
SW14N
SW14H
SW18N
SW18H

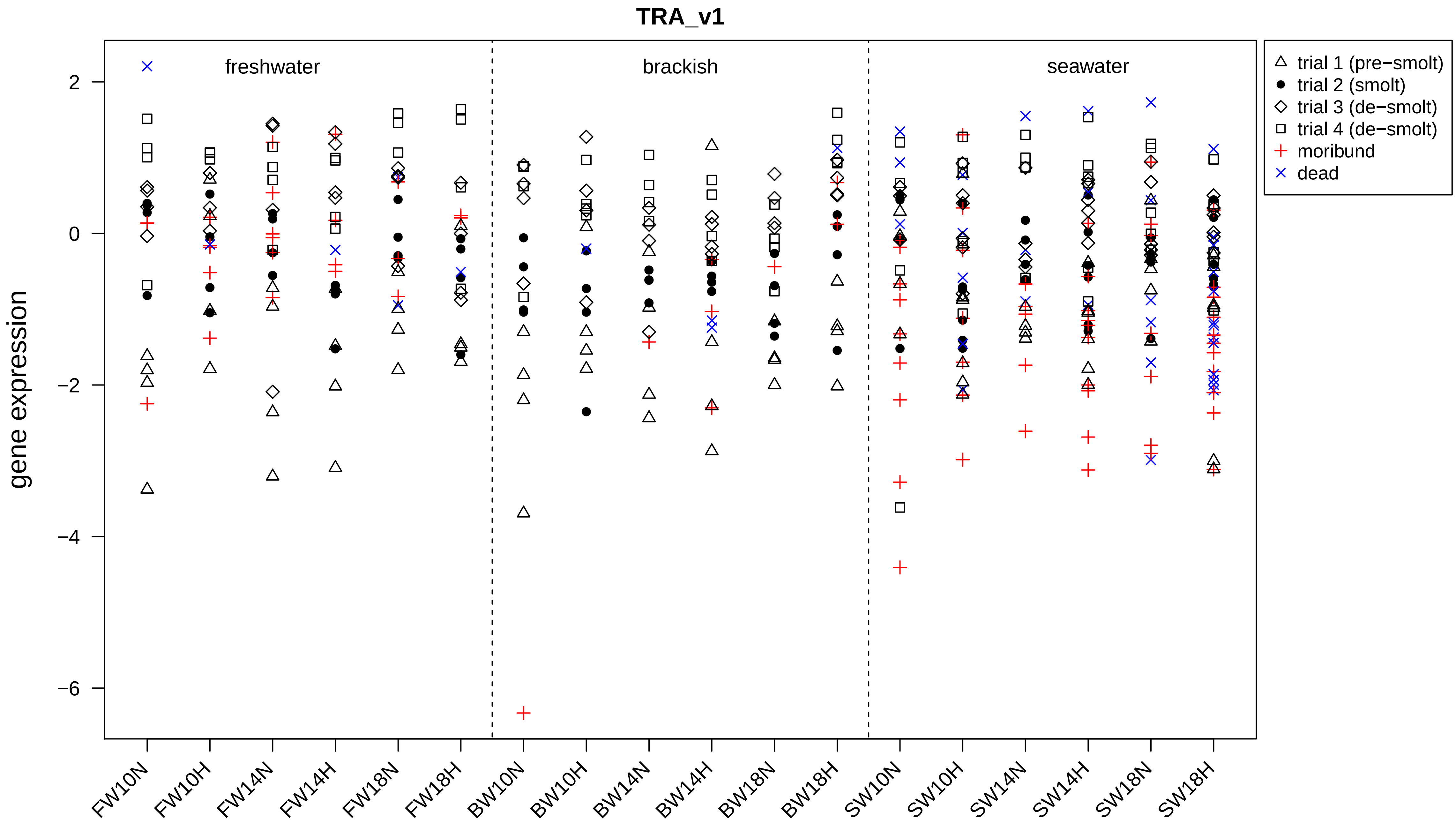
TRA_v1
freshwater
brackish
seawater
trial 1 (pre−smolt)
trial 2 (smolt)
trial 3 (de−smolt)
trial 4 (de−smolt)
moribund
dead
gene expression
2
0
−2
−4
−6
FW10N
FW10H
FW14N
FW14H
FW18N
FW18H
BW10N
BW10H
BW14N
BW14H
BW18N
BW18H
SW10N
SW10H
SW14N
SW14H
SW18N
SW18H

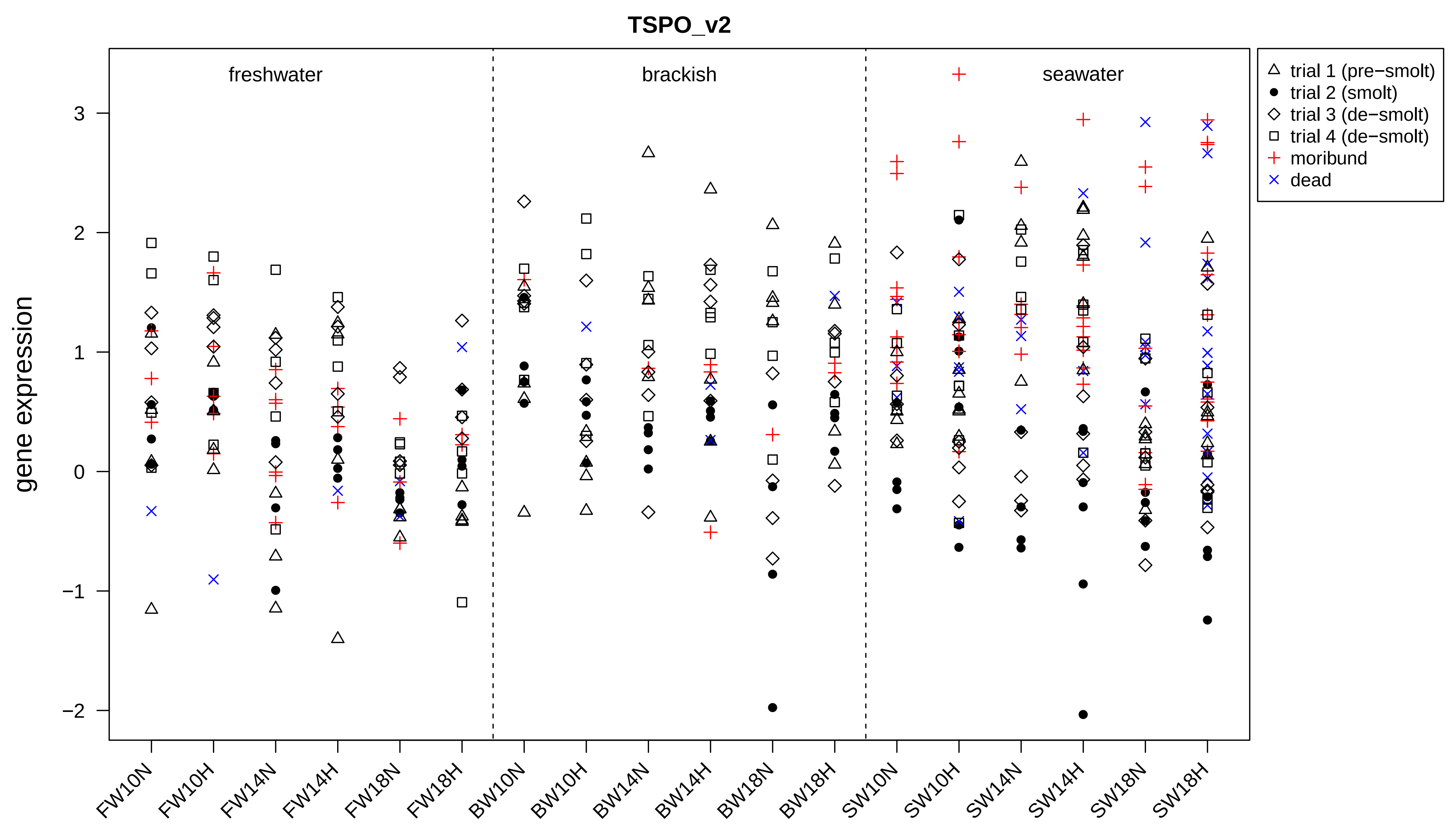
TSPO_v2
freshwater
brackish
seawater
gene expression
3
2
1
0
−1
−2
FW10N
FW10H
FW14N
FW14H
FW18N
FW18H
BW10N
BW10H
BW14N
BW14H
BW18N
BW18H
SW10N
SW10H
SW14N
SW14H
SW18N
SW18H
trial 1 (pre−smolt)
trial 2 (smolt)
trial 3 (de−smolt)
trial 4 (de−smolt)
moribund
dead

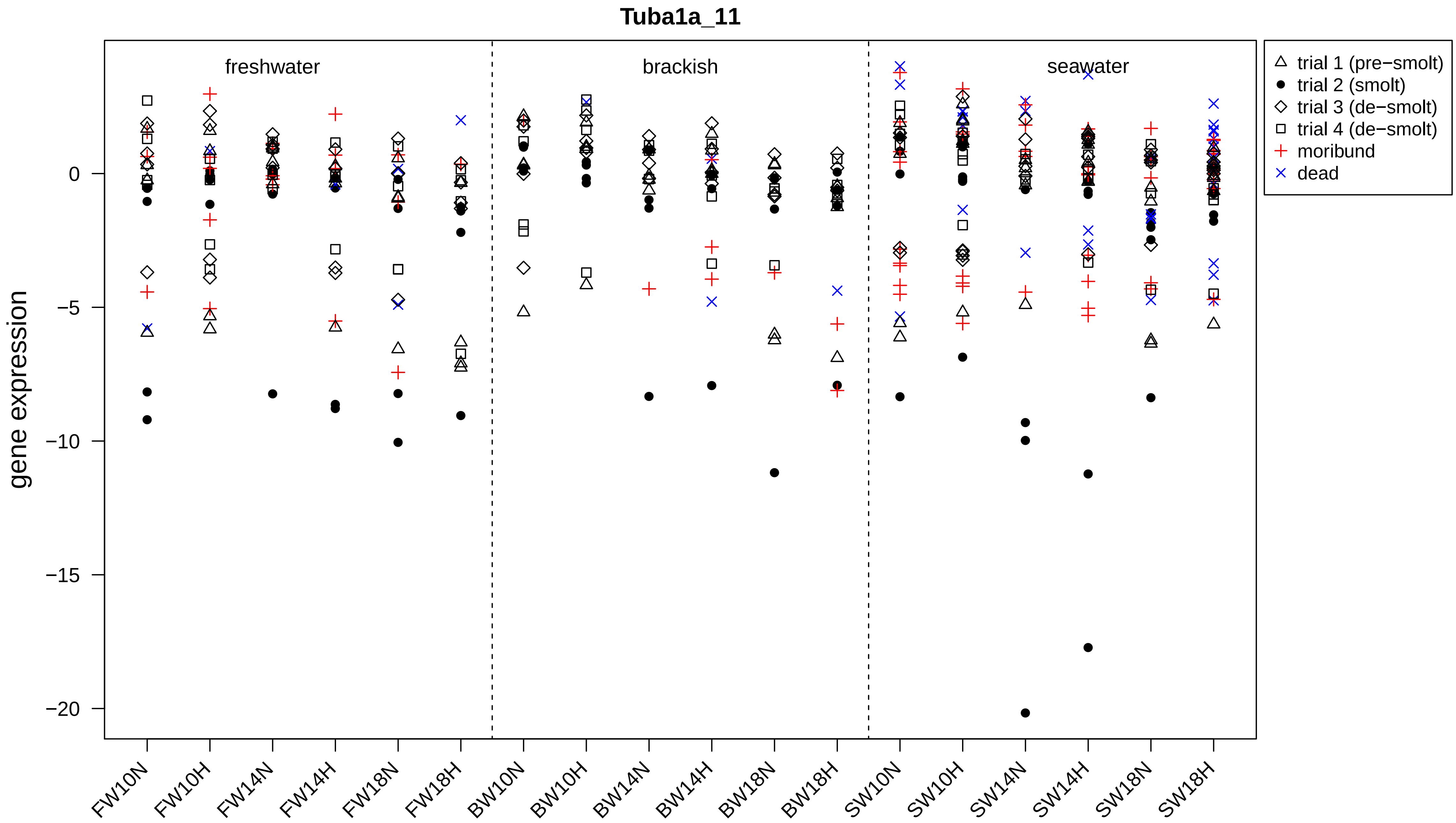
Tuba1a_11
freshwater
brackish
seawater
gene expression
0
−5
−10
−15
−20
FW10N
FW10H
FW14N
FW14H
FW18N
FW18H
BW10N
BW10H
BW14N
BW14H
BW18N
BW18H
SW10N
SW10H
SW14N
SW14H
SW18N
SW18H
trial 1 (pre−smolt)
trial 2 (smolt)
trial 3 (de−smolt)
trial 4 (de−smolt)
moribund
dead

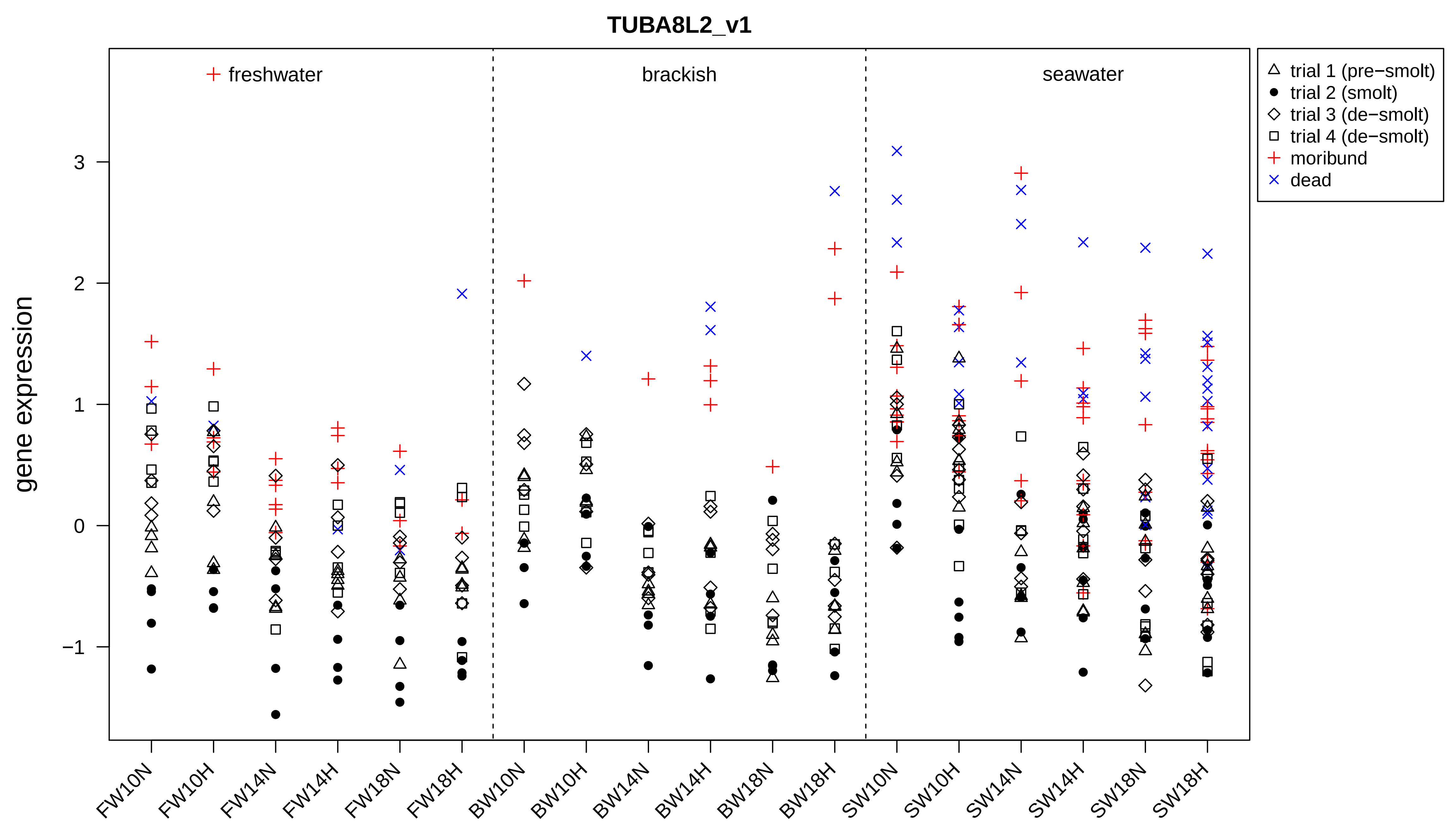
TUBA8L2_v1
gene expression
freshwater
brackish
seawater
trial 1 (pre−smolt)
trial 2 (smolt)
trial 3 (de−smolt)
trial 4 (de−smolt)
moribund
dead
3
2
1
0
−1
FW10N
FW10H
FW14N
FW14H
FW18N
FW18H
BW10N
BW10H
BW14N
BW14H
BW18N
BW18H
SW10N
SW10H
SW14N
SW14H
SW18N
SW18H

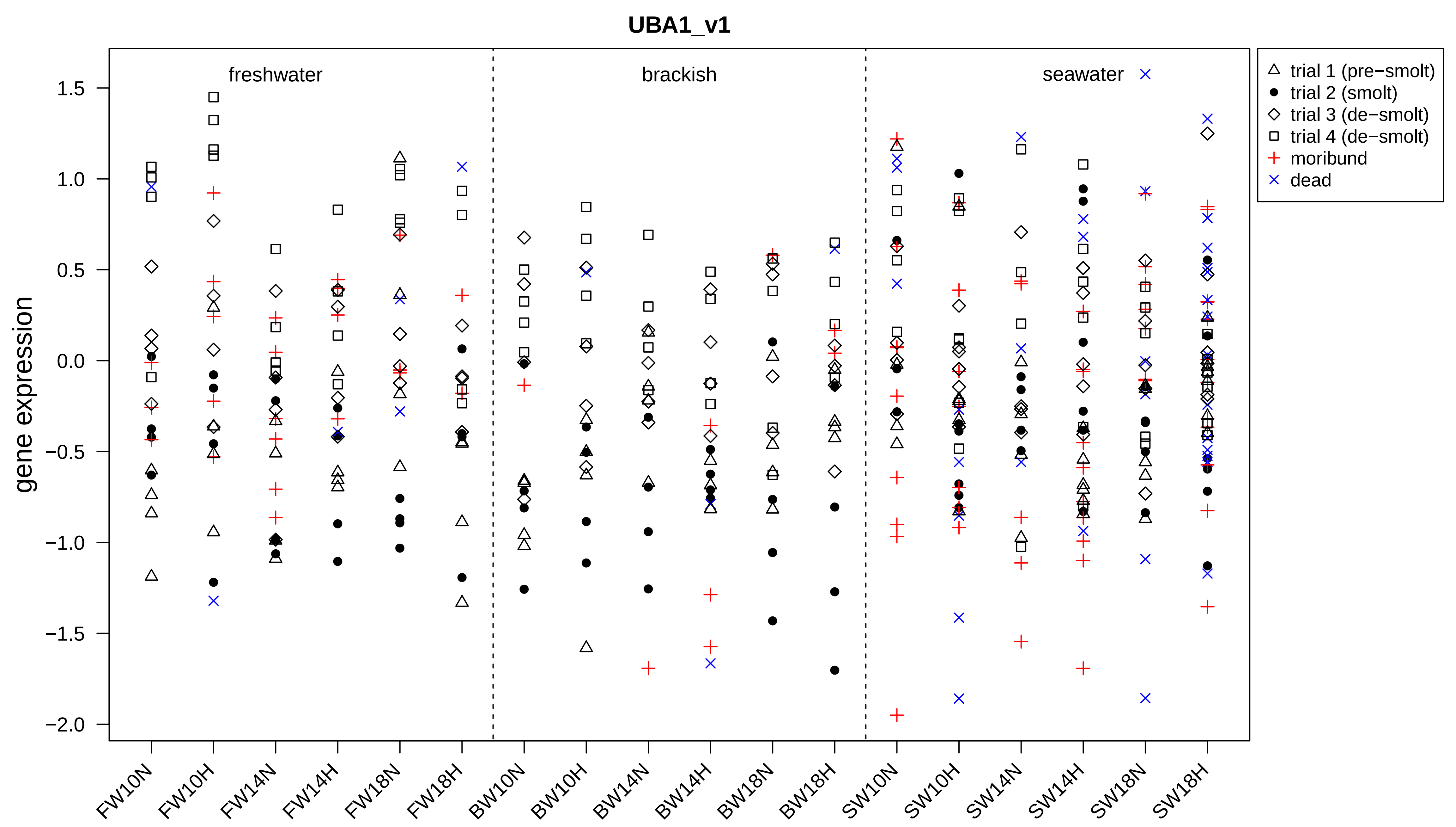

UBA1_v1
gene expression
freshwater
brackish
seawater
trial 1 (pre−smolt)
trial 2 (smolt)
trial 3 (de−smolt)
trial 4 (de−smolt)
moribund
dead
1.5
1.0
0.5
0.0
−0.5
−1.0
−1.5
−2.0
FW10N
FW10H
FW14N
FW14H
FW18N
FW18H
BW10N
BW10H
BW14N
BW14H
BW18N
BW18H
SW10N
SW10H
SW14N
SW14H
SW18N
SW18H

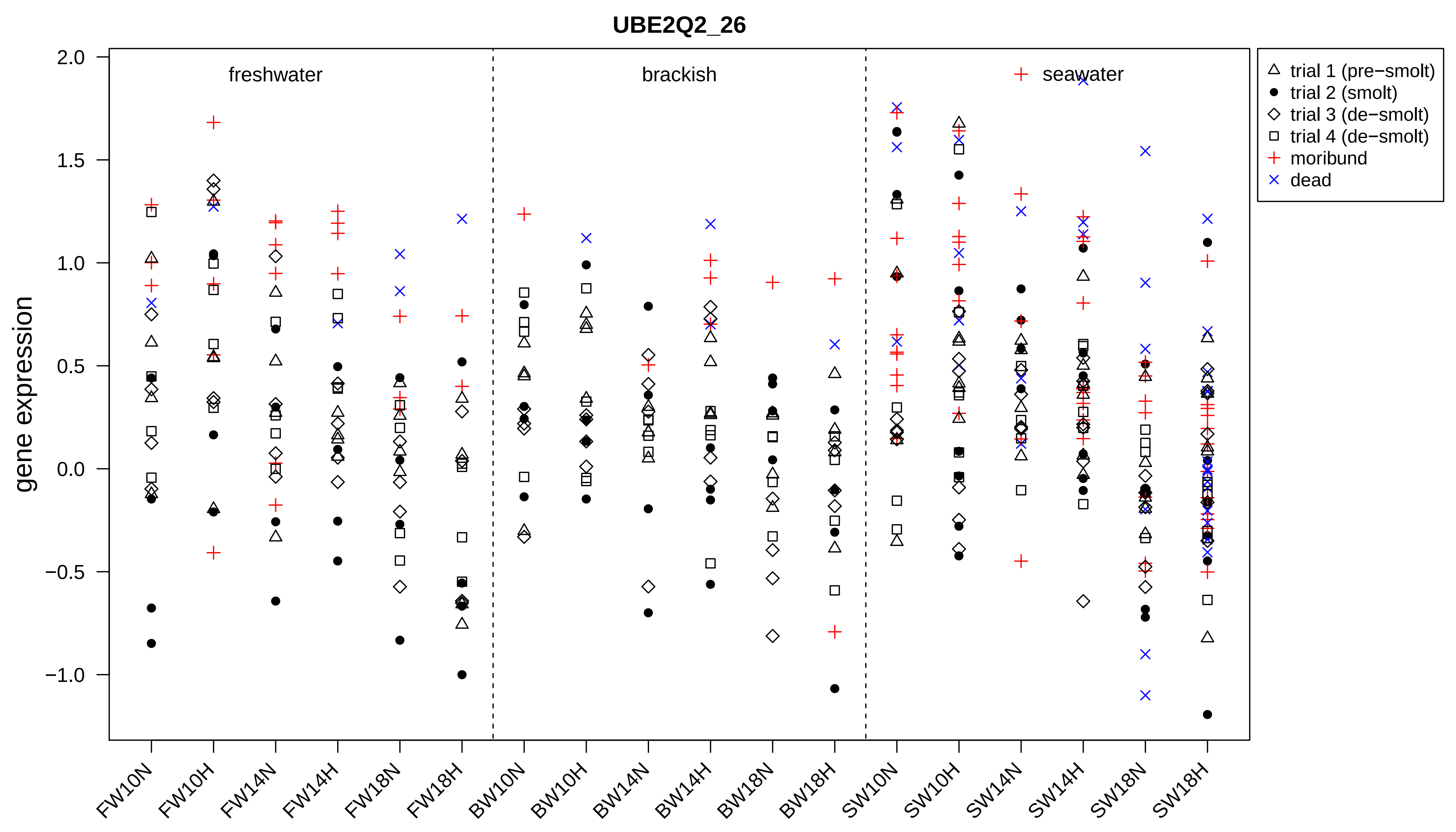
UBE2Q2_26
gene expression
2.0
1.5
1.0
0.5
0.0
−0.5
−1.0
freshwater
brackish
seawater
trial 1 (pre−smolt)
trial 2 (smolt)
trial 3 (de−smolt)
trial 4 (de−smolt)
moribund
dead
FW10N
FW10H
FW14N
FW14H
FW18N
FW18H
BW10N
BW10H
BW14N
BW14H
BW18N
BW18H
SW10N
SW10H
SW14N
SW14H
SW18N
SW18H

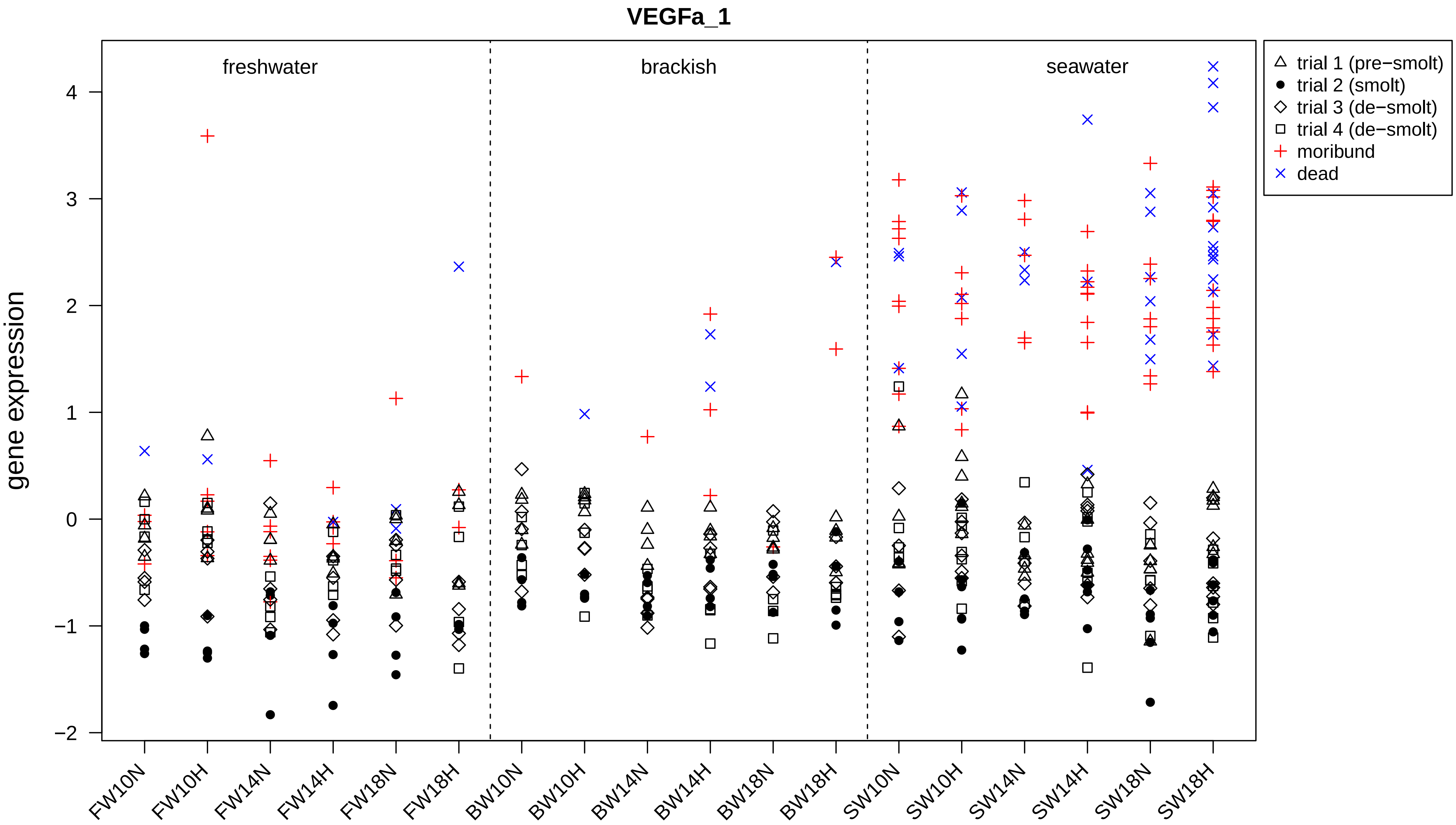
VEGFa_1
freshwater
brackish
seawater
trial 1 (pre−smolt)
trial 2 (smolt)
trial 3 (de−smolt)
trial 4 (de−smolt)
moribund
dead
gene expression
4
3
2
1
0
−1
−2
FW10N
FW10H
FW14N
FW14H
FW18N
FW18H
BW10N
BW10H
BW14N
BW14H
BW18N
BW18H
SW10N
SW10H
SW14N
SW14H
SW18N
SW18H

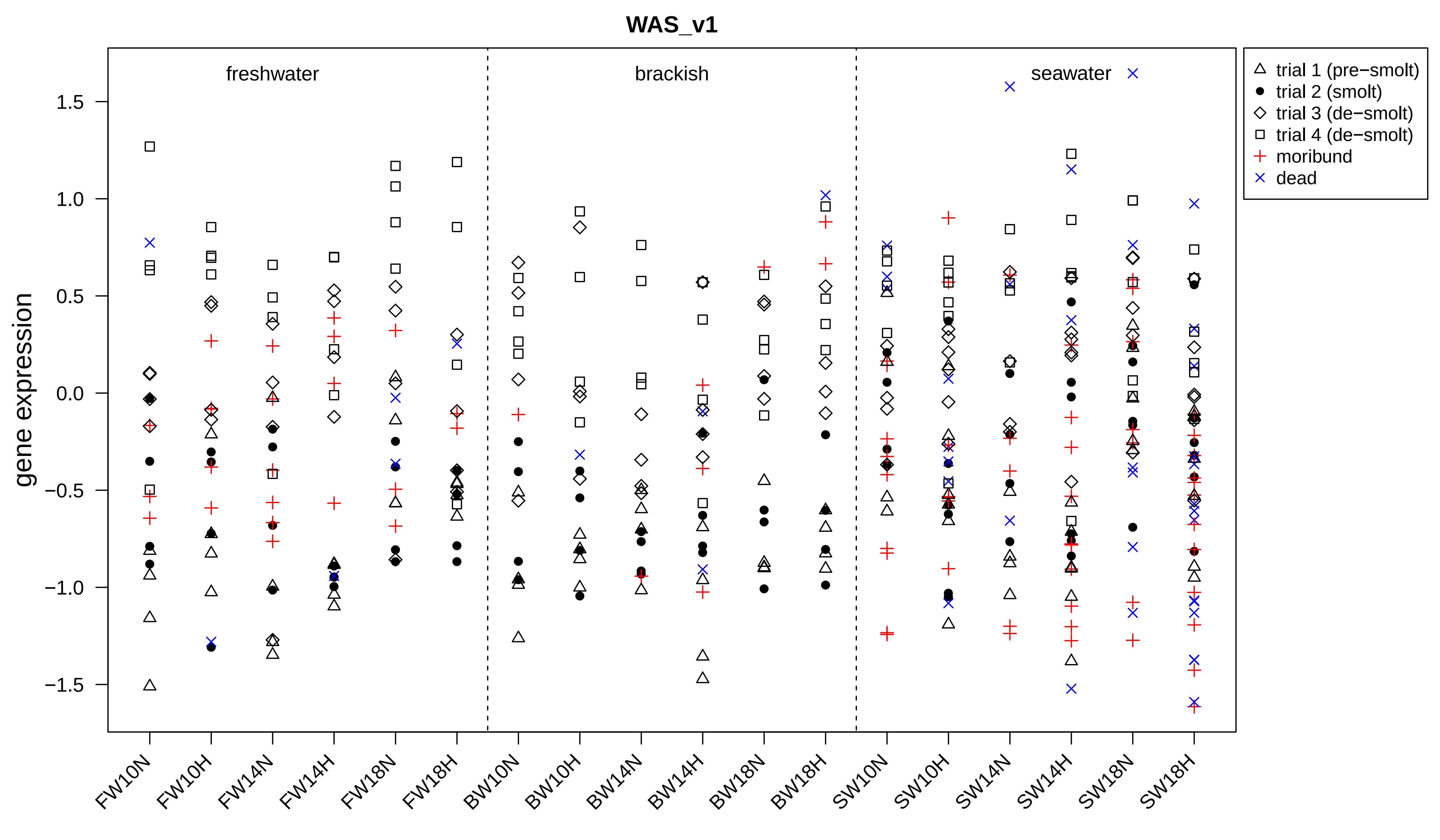
WAS_v1
freshwater
brackish
seawater
gene expression
1.5
1.0
0.5
0.0
−0.5
−1.0
−1.5
trial 1 (pre−smolt)
trial 2 (smolt)
trial 3 (de−smolt)
trial 4 (de−smolt)
moribund
dead
FW10N
FW10H
FW14N
FW14H
FW18N
FW18H
BW10N
BW10H
BW14N
BW14H
BW18N
BW18H
SW10N
SW10H
SW14N
SW14H
SW18N
SW18H

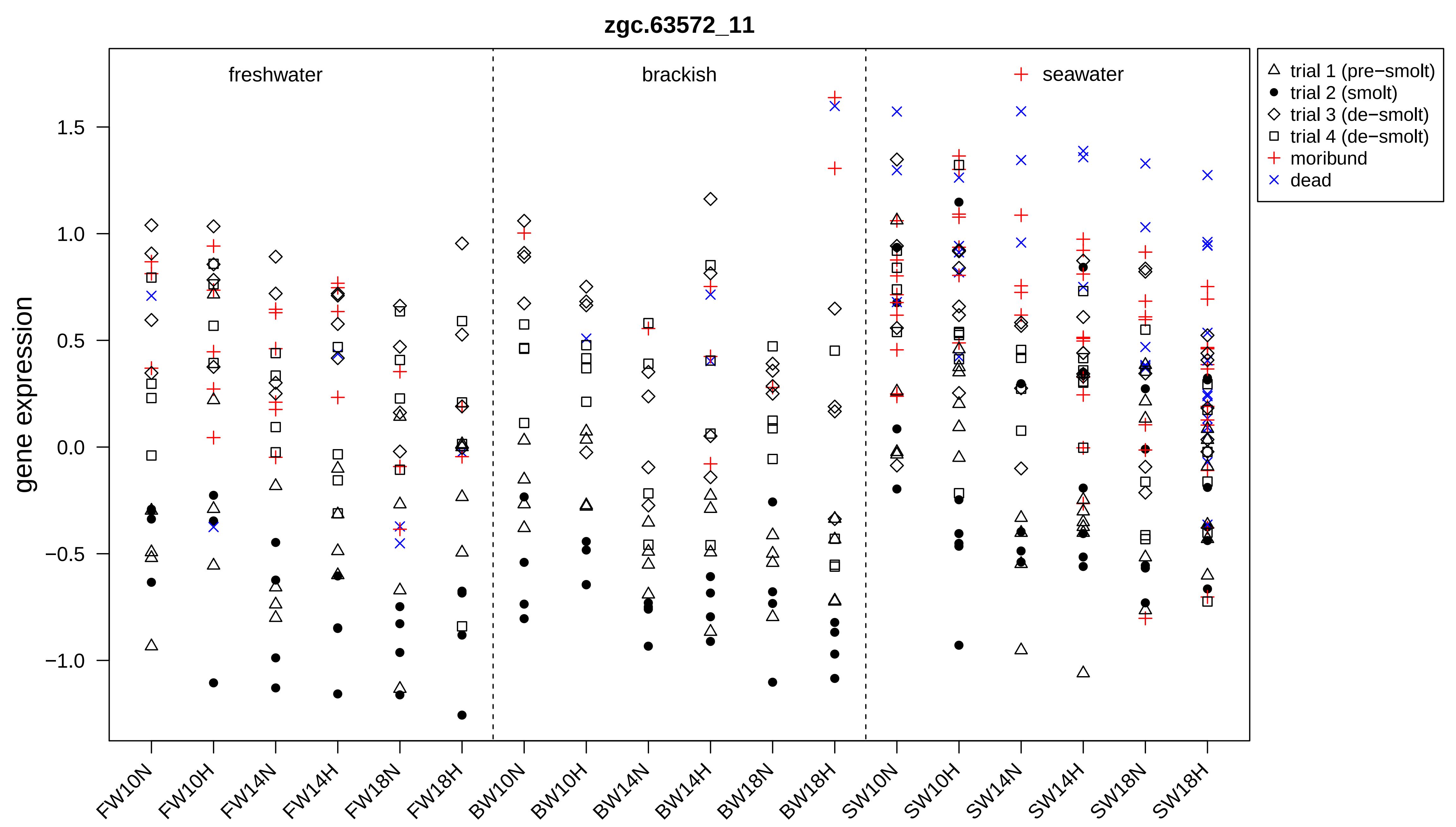

zgc.63572_11
freshwater
brackish
seawater
gene expression
1.5
1.0
0.5
0.0
−0.5
−1.0
FW10N
FW10H
FW14N
FW14H
FW18N
FW18H
BW10N
BW10H
BW14N
BW14H
BW18N
BW18H
SW10N
SW10H
SW14N
SW14H
SW18N
SW18H
trial 1 (pre−smolt)
trial 2 (smolt)
trial 3 (de−smolt)
trial 4 (de−smolt)
moribund
dead

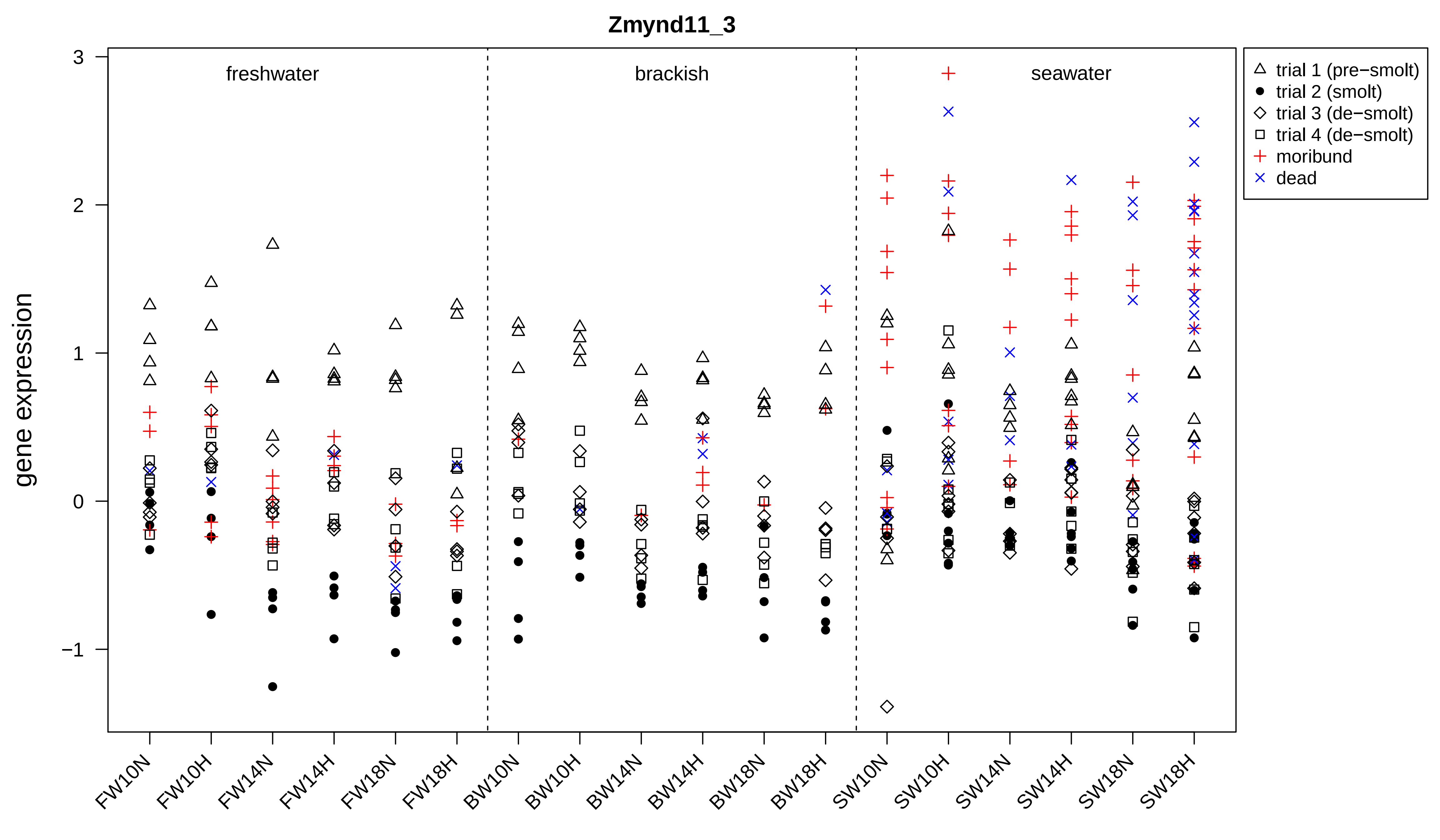

Zmynd11_3
gene expression
freshwater
brackish
seawater
trial 1 (pre−smolt)
trial 2 (smolt)
trial 3 (de−smolt)
trial 4 (de−smolt)
moribund
dead
3
2
1
0
−1
FW10N
FW10H
FW14N
FW14H
FW18N
FW18H
BW10N
BW10H
BW14N
BW14H
BW18N
BW18H
SW10N
SW10H
SW14N
SW14H
SW18N
SW18H
